# Structural basis for peptide substrate specificities of glycosyltransferase GalNAc-T2

**DOI:** 10.1101/2020.06.25.171371

**Authors:** Sai Pooja Mahajan, Yashes Srinivasan, Jason W. Labonte, Matthew P. DeLisa, Jeffrey J. Gray

## Abstract

The polypeptide *N-*acetylgalactosaminyl transferase (GalNAc-T) enzyme family initiates *O*-linked mucin-type glycosylation. The family constitutes 20 isozymes in humans—an unusually large number—unique to O-glycosylation. GalNAc-Ts exhibit both redundancy and finely tuned specificity for a wide range of peptide substrates. In this work, we deciphered the sequence and structural motifs that determine the peptide substrate preferences for the GalNAc-T2 isoform. Our approach involved sampling and characterization of peptide–enzyme conformations obtained from Rosetta Monte Carlo-minimization–based flexible docking. We computationally scanned 19 amino acid residues at positions −1 and +1 of an eight-residue peptide substrate, which comprised a dataset of 361 (19×19) peptides with previously characterized experimental GalNAc-T2 glycosylation efficiencies. The calculations recapitulated experimental specificity data, successfully discriminating between glycosylatable and non-glycosylatable peptides with a probability of 96.5% (ROC-AUC score), a balanced accuracy of 85.5% and a false positive rate of 7.3%. The glycosylatable peptide substrates *viz.* peptides with proline, serine, threonine, and alanine at the −1 position of the peptide preferentially exhibited cognate sequon-like conformations. The preference for specific residues at the −1 position of the peptide was regulated by enzyme residues R362, K363, Q364, H365 and W331, which modulate the pocket size and specific enzyme-peptide interactions. For the +1 position of the peptide, enzyme residues K281 and K363 formed gating interactions with aromatics and glutamines at the +1 position of the peptide, leading to modes of peptide-binding sub-optimal for catalysis. Overall, our work revealed enzyme features that lead to the finely tuned specificity observed for a broad range of peptide substrates for the GalNAc-T2 enzyme. We anticipate that the key sequence and structural motifs can be extended to analyze specificities of other isoforms of the GalNAc-T family and can be used to guide design of variants with tailored specificity.

## Introduction

In higher organisms, O-linked *N*-acetylgalactosamine (GalNAc) glycosylation (or mucin-type glycosylation) is an abundant and essential post-translational modification. This type of glycosylation is initiated by a family of glycosyltransferases (GTs) known as polypeptide *N*-acetylgalactosaminyltransferases or GalNAc-Ts (also referred to as GALNTs). These enzymes transfer a GalNAc sugar from a donor uridine di-phosphate (UDP) nucleotide to the hydroxyl group of a threonine or serine residue of an acceptor peptide. This transfer is the first committed step of mucin-type O-glycosylation, and these enzymes, therefore, define the sites of O-glycosylation. The resulting O-linked GalNAc is further extended to one of the four common core structures, which can be subsequently extended to give mature linear or branched glycans.^1,2^ Aberrant O-glycosylation is a well-known marker of many cancers and has also been linked to developmental and metabolic disorders.^3,4^

In humans, the GalNAc-T family constitutes 20 isozymes. The unusually large number of isoforms for glycosylation is unique to O-glycosylation, and the multiplicity is conserved in mammalian evolution, suggesting that cell or tissue specific isoforms have specialized functions.^5^ The isoforms exhibit specific substrate preferences that vary with isoenzyme surface charge, prior neighboring long-range and short-range glycosylation patterns and the sequence of the acceptor peptide substrate. Over the last two decades, the peptide substrate preferences for a large number of isoforms have been established by *in vitro* studies.^6–8^ The peptide substrate is characterized by a sequence motif (or sequon), Thr/Ser–Pro–X–Pro (T/SPXP), where T/S is the site of glycosylation (position 0). This sequon is the only conserved consensus motif modified by all isoforms except T7 and T10. The proline at the +3 position of the sequon is supported by a conserved structural motif, *viz.,* the “proline pocket” in the enzyme’s peptide binding groove in all isoforms that bind the T/SPXP motif.^9–11^ For the remaining positions in the sequon, most isoforms exhibit overlapping yet selective preferences for different amino acid residues. For example, at the −1 position with respect to the glycosylation site, T1 favors aromatics^12^ and T12 prefers bulky non-polar residues;^13^ whereas T2 exhibits very little to no activity for these amino acids and instead prefers threonine, proline, alanine, and serine. Yet both T1 and T2 glycosylate the sequon TTP^12^ (with threonine at −1 and proline at +1 positions). These observations have led to the hypothesis that GalNAc-Ts exhibit both redundancy and finely tuned specificity for a wide range of peptide substrates.

While there is ample experimental data on the peptide substrate specificities of various isoforms, the molecular basis for observed peptide substrate specificities is not well understood. Computational work, so far, has been focused on understanding the mechanism of sugar transfer,^14,15^ conformational changes in the flexible loop in the catalytic domain,^16^ and the effect of the flexible linker connecting the catalytic and lectin domains.^11^ None of the computational studies so far have examined the amino acid preferences at different positions on the peptide. Computational studies can pinpoint key positions and structural motifs on an isoform that contribute to peptide substrate specificity. These sequence and structural motifs can be studied across isoforms to reveal more general patterns, to modulate enzyme specificity, and to gain insight into the consequences of enzyme and substrate mutations implicated in aberrant glycosylation, (*e.g.*, colorectal cancer associated mutations of GalNAc-T12^17^) paving the way for rational design of specific drugs/inhibitors^18^.

In this work, we seek to understand the sequence and structural motifs that determine the peptide substrate preferences for the GalNAc-T2 isoform. Our immediate goal is to recapitulate experimentally determined specificity in terms of glycosylation efficiency for sequon variations at positions −1 and +1 (19 amino acid residues tested for each position), as reported by Kightlinger *et al.,*^12^ and to understand the structural motifs that best explain experimentally observed trends. To recapitulate experimentally observed specificities for a large dataset, we need an efficient, high-throughput computational method that can capture the key mechanisms of enzymatic catalysis.

Enzymatic catalysis relies primarily on selective transition state stabilization, ground state (reactants) destabilization, dynamics, and active-site gating.^19,20^ In practice, these effects occur at different length- and time-scales and therefore cannot be accurately captured by a single method. Hybrid quantum mechanics/molecular mechanics (QM/MM) simulations have been able to recapitulate catalytic proficiency or mechanistic details for many enzymes such as Kemp eliminases^21^ or glycoside hydrolases^22^ as they are well-suited to characterize the transition state. QM/MM simulations, however, are not suitable to capture binding or dynamics over longer timescales and are prohibitively expensive for a larger dataset. Other factors that determine the stability of the transition state are electrostatic- and shape-complementarity at the peptide-enzyme interface. Electrostatic complementarity can be captured by various computational techniques (*e.g.*, Monte Carlo (MC) or molecular dynamics (MD)-based methods with Poisson-Boltzmann electrostatics or other continuum electrostatics models) at different length-scales. Other effects are determined by the thermodynamics of the enzyme-peptide interactions. To achieve a lower free energy of activation,^19,23^ the enzyme must stabilize the transition state selectively relative to the reactants. Additionally, if the product is too stable in the enzyme’s active site, product release becomes the catalytic rate-limiting step. This thermodynamic description demands the use of methods that capture multiple states (reactants, products and transition states).^24,25^ Furthermore, dynamics is important in many catalytic mechanisms, from small vibrations that lead to rate-promoting motions^26^ to large conformational changes and rearrangements in the molecular structure.^27^ Active site gating is another important mechanism for catalysis by which key residues outside the active site regulate access to the active site.^28,29^ These thermodynamic and kinetic effects, primarily in the nanosecond to microsecond timescales, can be captured faithfully by MD simulations though such simulations can be computationally prohibitive for comparing a large number of substrates. An alternative to MD simulations are Monte Carlo-minimization^30^ (MCM) approaches, which are computationally faster and can be reliably used to determine thermodynamically stable native-like states.

Rosetta-based MCM computational protocols, notably, pepspec,^31^ sequence-tolerance^32,33^ and MFPred^34^ have previously been used for predicting the sequence profiles of peptides recognized by various multi-specific protein recognition domains (PRDs) such as PDZ, SH2, SH3, kinases, and proteases. All protocols rely on MCM sampling and aim to approximate the stabilization of the substrate-bound state or transition state in the enzyme’s peptide binding groove. The transition state is approximated by known cognate sequon conformations in the enzyme’s active site (based on crystal structures and/or homology modeling) with additional constraints to preserve important structural motifs pertaining to the transition state, when available. In the absence of constraints, this approach is equivalent to evaluating the stabilization of the substrate-bound state.^35^ MCM allows for faster sampling facilitating the scanning of a large number of amino acid residues at multiple positions of the peptide substrate. All three protocols achieve impressive accuracy in predicting experimentally observed profiles for many PRDs. However, since all three methods are developed with the broad goal of predicting sequence specificity profiles for a range of PRDs, the accuracy of prediction may not be sufficient to pinpoint subtle differences in specificity for a specific target of interest. For example, the sequence-tolerance protocol pre-calculates the interactions between all interacting residues ignoring changes in conformation of the peptide in the protein’s binding pocket. All three methods struggle to predict specificity for HIV−1 protease, which has a relaxed specificity profile and a preference for small hydrophobic residues, similar to GalNAc-T2. Additionally, all three protocols employ limited backbone sampling, prohibiting the free conformational sampling of the peptide in the binding groove. For the more targeted goal of designing a peptide inhibitor to discriminate between two similar PDZ domains, Zheng *et al.*^36^ employ extensive conformational sampling using the full-fledged flexpepdock protocol^37,38^ along with the CLASSY method to achieve a solution with desired specificity and affinity goals. Similarly, Pethe *et al*.^39^ were able to obtain significantly improved prediction accuracies for proteases (including HIV−1 protease) compared to previous methods (MFPred, sequence tolerance and pepspec) by employing machine learning and a discriminatory score based on geometric features, interface score terms from Rosetta, and electrostatic score terms from Amber. In another work, Pethe *et al.*^40^ used supervised learning on experimentally obtained deep-sequencing data and information from structure-based models to chart the specificity landscape of 3.2 million substrate variants of the viral protease HCV.

Here, we sought to understand specificity determinants for a specific isoform of the GalNAc-T family and to pinpoint sequence and structural motifs in the enzyme that explain fine-tuning of specificity. To this end, we developed a customized Rosetta-based protocol^41,42^ that allowed us to model structures of all 361 peptide sequons (19×19) with the GalNAc-T2 enzyme and computationally determine the sequon preference for the GalNAc-T2 isoform. Our protocol was similar in spirit to earlier protocols in that it docks the peptide substrate into the enzyme’s active site. However, unlike pepspec,^31^ sequence-tolerance^32,33^ and MFPred^34^ and similar to the protocol of Zheng *et al.,*^36^ we allowed fully flexible peptide sampling (as opposed to limited or no backbone sampling) followed by clustering and analysis of the sampled low energy decoys. Our strategy relied on characterizing the peptide binding to the enzyme with a range of structural features at the interface as a function of the amino acid residues at the +1 and −1 positions. Using our methodology, we were able to identify features that recapitulated high-quality experimental specificity data for GalNAc-T2. Extensive peptide backbone sampling revealed that the peptide binding groove of GalNAc-T2 stabilized multiple competing conformations/states – some leading to efficient glycosylation and others hampering it. Furthermore, multiple stable states suggested that kinetics might play an important role in determining specificity. Thus, finely-tuned specificity might be achieved by modulating the relative stability of these states to discriminate between peptide substrates for an isoform and across isoforms. Overall, our work reveals key residues on the enzyme that determine peptide substrate preferences at various sequon positions.

## Results

### Clustering of low interaction energy decoys reveals that peptides exhibit multiple competing low-energy conformations

We studied all 361 (19×19) sequons obtained by scanning 19 amino acids (all amino acids except cysteine) at positions −1 and +1 with respect to the modified threonine (Thr_0_). The experimentally determined glycosylation efficiencies for all of these sequons was determined by Kightlinger *et al*.^12^ and replotted in Figure 1A. For each sequon, we started with the co-crystal structure of the peptide and UDP-sugar bound to the enzyme (pdb ids: 4d0z and 2ffu, respectively).^16^ We mutated the residues at the −1 and +1 position of the peptide to the target sequon and repacked and minimized the nearby side chains to obtain a starting enzyme-peptide configuration for the target sequon. We then subjected this starting structure to MCM sampling of rigid-body displacements and peptide torsion angles in two stages - a low-resolution centroid stage with simulated annealing followed by a high-resolution all-atom stage to generate 2,000 structures (decoys) per sequon (see Methods). In this paper, we use the shorthand notation X_N_ for amino acid ‘X’ (denoted by 1-letter code) and sequon position ‘N’ and a sequon with the shorthand notation X_−1_TX_+1_. For example, P_−1_ denotes amino acid proline at the −1 position, M_+1_ denotes a methionine at the +1 position and P_−1_TM_+1_ denotes a sequon or peptide with P_−1_ and M_+1_. For brevity, we also refer to peptides or sequons containing residue X at position N as “X_N_ peptides” or “X_N_ sequons” respectively.

**Figure 1.**
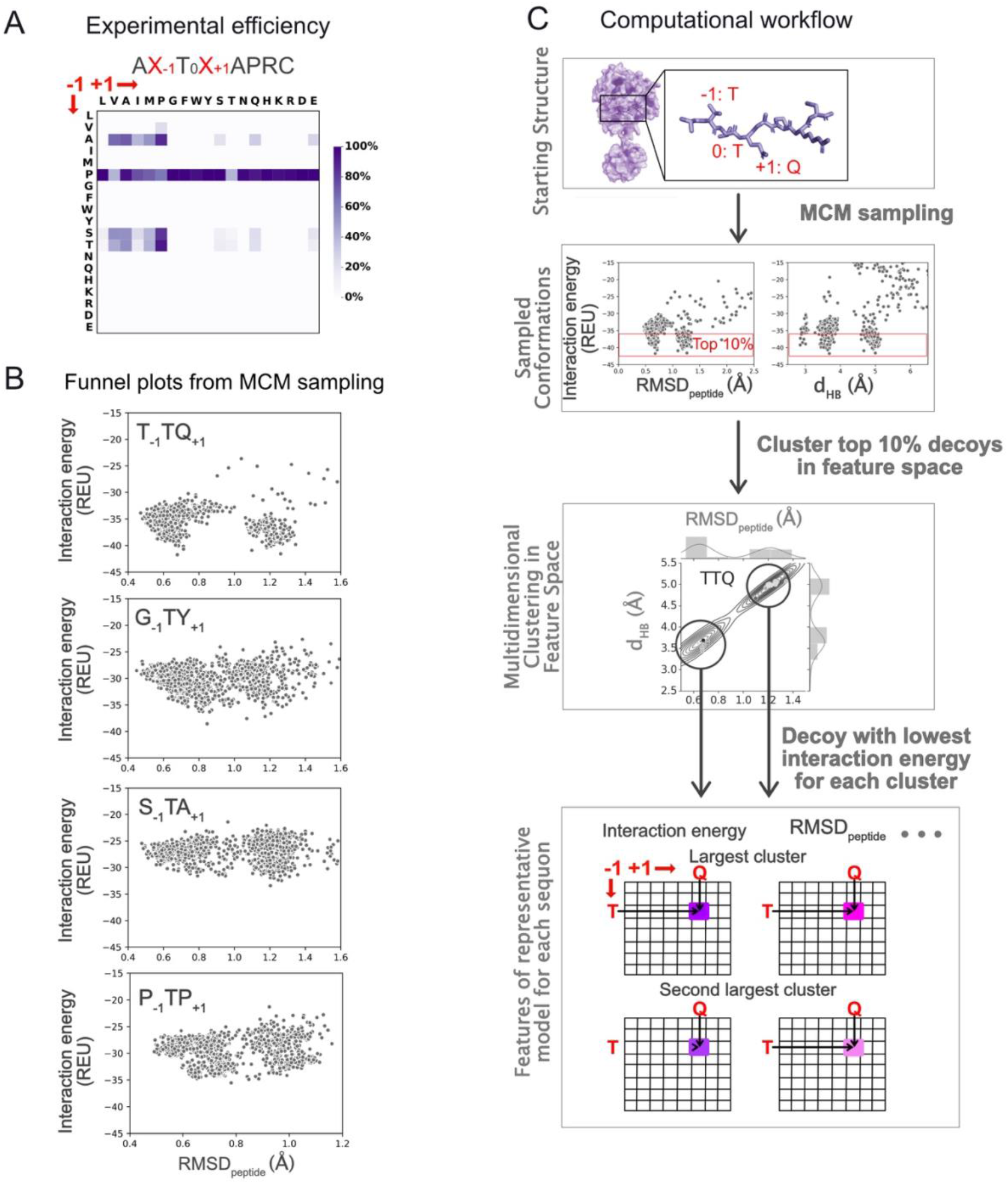
Computational workflow to determine glycosylation efficiency of glycosyltransferase GalNAc-T2 for peptide substrates for an experimentally characterized dataset obtained by scanning 19 amino acid residues (all except cysteine) at positions −1 and +1 of the acceptor peptide. (A) For reference, a replot of the experimentally determined glycosylation efficiencies (efficiency data from Kightlinger *et al.*^12^). (B) Monte Carlo minimization sampling of individual sequons docked to GalNAcT2 result in “funnel plots” like the four shown. Each point represents one structural model, or “decoy,” at its corresponding RMSD from the reference structure and the interaction energy calculated by Rosetta. (C) Steps in the computational workflow to characterize enzyme–peptide interactions for a representative sequon, T_−1_TQ_+1_, with T at the −1 position and Q at the +1 position. For each sequon, we selected the top-10%-scoring decoys (by interaction energy) from MCM sampling and clustered them using three features (see main text for features). For each sequon, we characterized the two largest clusters by the lowest interaction energy belonging to that cluster and then examined which features (or combinations of features) could recapitulate the experimental glycosylation efficiencies (A).

Preliminary analysis of the decoys showed that many peptides exhibited multiple stable states with comparable energies of interaction (interaction energy) between the peptide and enzyme. In Figure *1*B, we show plots of interaction energy vs. distance to the reference crystal structure for four randomly chosen sequons obtained from MCM sampling of the peptide substrate with the respective sequons in the enzyme’s peptide-binding groove. For all four sequons in Figure 1B, we observed multiple clusters of low interaction energy decoys, or “funnels.” For example, for sequon T_−1_TQ_+1_, we observed two distinct funnels at RMSD_peptide_ (the root mean square deviation of C_α_ carbons of the peptide backbone with respect to the peptide in the crystal structure) values of about ~ 0.65 Å and 1.25 Å with comparable lowest interaction energies. Overall, 57% (205/361) of the sequons exhibited two significant clusters. 39/205 (19.0%) and 81/205 (39.5%) sequons exhibited lowest energy states for each cluster within 0.5 and 1.0 Rosetta energy units (or REUs) respectively, of each other, underscoring the importance of considering both states (SI Figure 1).

To characterize multiple low-energy conformations and to construct a more complete picture of the landscape of structural conformations sampled by the peptide substrate in the enzyme cavity, we developed the computational flow summarized in Figure 1C. For each sequon, we selected the top-10%-scoring decoys (by interaction energy) from MCM sampling and then clustered them using three features. The first feature, RMSD_peptide_, characterized decoys on the basis of the similarity of the peptide backbone conformation and position of the peptide in the crystal structure. Next, the distance between the hydroxyl group of T_0_ and the anomeric carbon (C_1_) on the sugar tracked the distance for the new glycosidic linkage. Finally, the distance between the amide group of T_0_ on the peptide and the oxygen of the β-phosphate group of UDP (O_β-PO4_) was a reaction coordinate characterizing a transition-state-stabilizing hydrogen bond between the backbone amide of T_0_ and UDP.^14^

We characterized the lowest-energy decoy for the largest and second-largest clusters obtained for each sequon and plotted heatmaps to show the distribution of the lowest interaction energy (Figure 2A), normalized cluster size (Figure 2B), RMSD_peptide_ (Figure 2C) and *d*_HB_ (Figure 2D) for all 361 sequons (see also SI Figure 2A). We also characterized the clusters by the decoy representing the center of the cluster, the average over all decoys with interaction energies within 1 REU and the average over the five decoys in the cluster with the lowest interaction energies. All strategies resulted in similar heatmaps (SI Figure 3) and hence, going forward, we represented a cluster by the lowest energy decoy belonging to that cluster.

**Figure 2.**
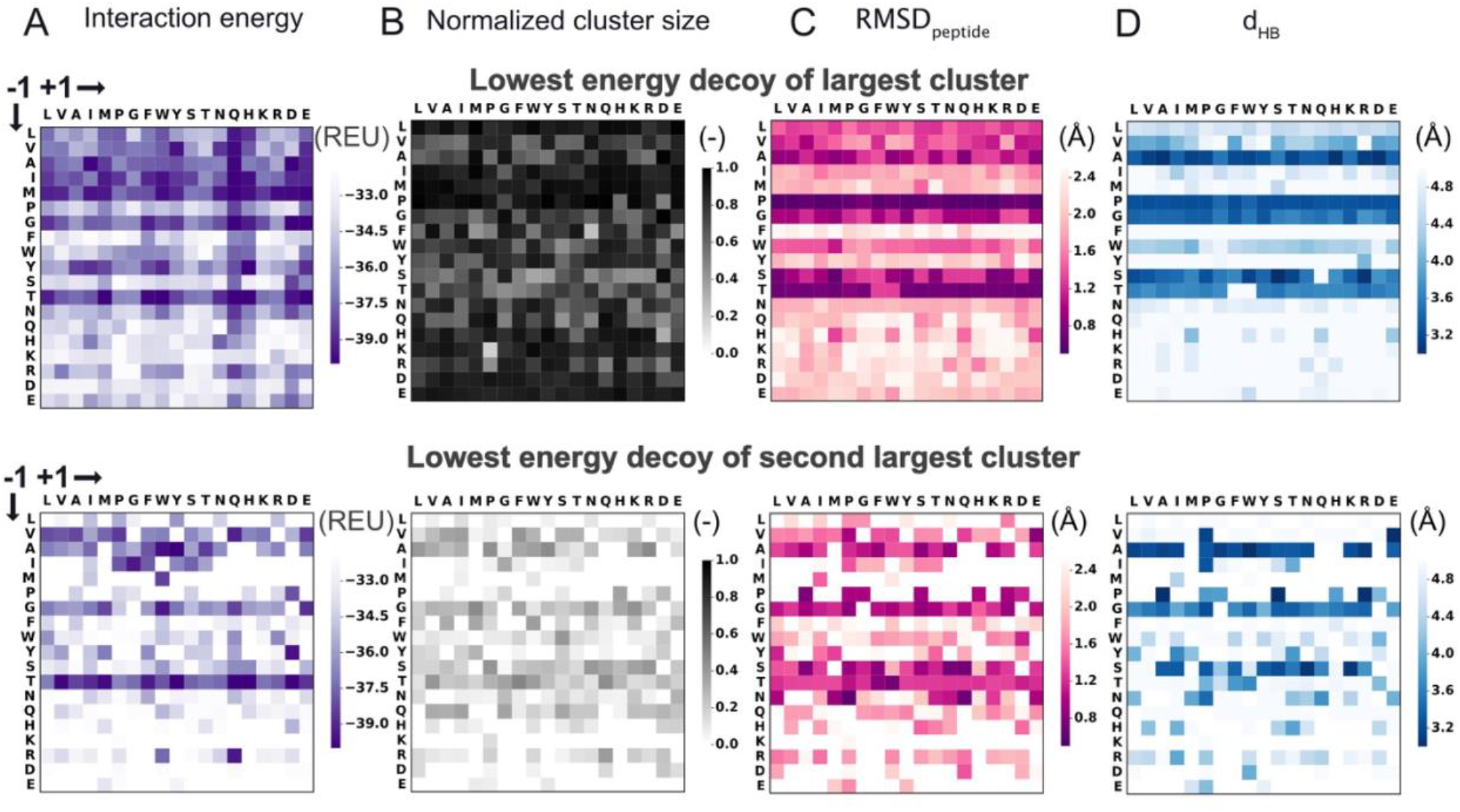
Characterization of the lowest-energy representative conformation for the top two clusters in Rosetta runs. As in Fig. 1A, each heatmap shows results for the 19×19 peptide sequons on a color scale. (A) Interaction energy, (B) normalized cluster size, (C) RMSD_peptide_, and (D) *d*_HB_ of the largest (top) and second-largest (bottom) clusters characterized by the lowest interaction energy decoy for each cluster.

Horizontal stripes emerging across the RMSD_peptide_ and *d*_HB_ heatmaps (Figure 2C-D) suggested that sequons with the same amino acid at the −1 position (horizontal axis) exhibited similar RMSD_peptide_ and *d*_HB_ values. To probe whether the low-energy conformations exhibited by various peptides depends on the identity of the residue in a position-specific manner, we plotted the RMSD_peptide_ and *d*_HB_ of the lowest energy decoy for the two largest clusters for each sequon colored by the amino acid residue at the −1 (Figure 3A) and +1 (Figure 3B) positions. It is apparent in Figure 3A that sequons with the same amino acid residue at the −1 position, especially A, G, T, P, S and V, were grouped or clustered together. The clustering or grouping suggests that sequons with the same amino acid at the −1 position exhibited similar conformations or low-energy states. Similar grouping was not observed for sequons with the same residue at the +1 position (Figure 3B). This high-level analysis of low-energy conformations for the entire dataset suggested that the −1 position plays a dominant role in determining the low-energy conformation(s) exhibited by a sequon and that the +1 position contributed in a secondary capacity.

**Figure 3.**
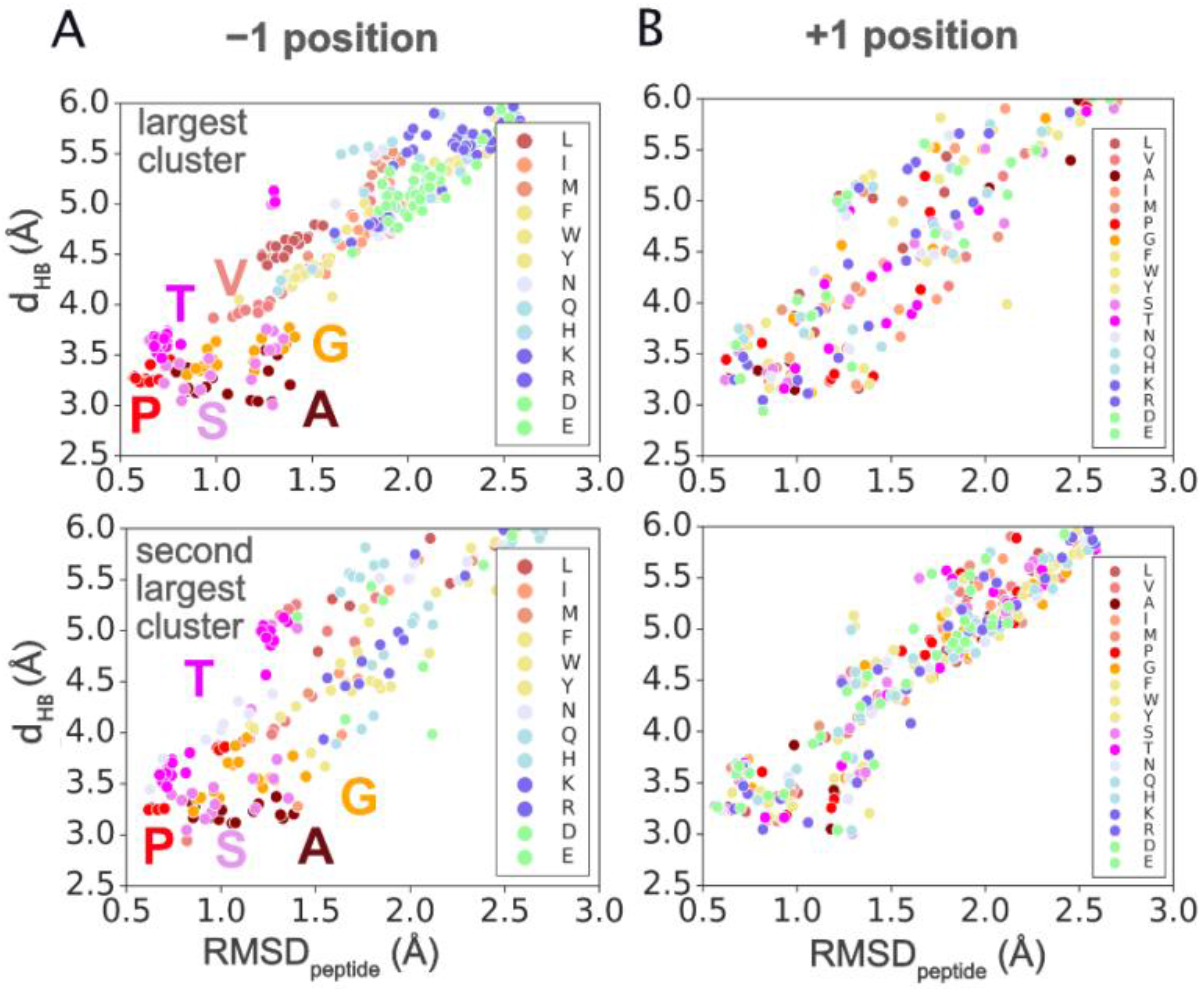
The major determinant of *d*_*HB*_ and RMSD_peptide_ sampled by lowest energy decoys is the amino acid residue at the −1 position. Lowest energy decoys belonging to the largest (top) and second largest clusters (bottom) for all sequons plotted as a function of the RMSD_peptide_ and *d*_*HB*_ and colored by (A) the residue at the −1 position of the sequon and (B) the residue at the +1 position of the sequon.

In the following sections, we present hypotheses to explain how each position (−1, +1) contributed in a characteristic manner to determine the low-energy conformations exhibited by a sequon and how these conformations, in turn, related to experimentally determined specificities. We characterized the low-energy conformations by a selected set of relevant features. To compare our predictions with experiments, we employed logistic regression and, unless otherwise indicated, we labeled all sequons with experimental glycosylation efficiencies greater than 10% (efficiency threshold) as glycosylatable and those with efficiencies less than 10% as un-glycosylatable, in line with previous work^34^. In Table 1 (and SI Table 1), for a chosen value of efficiency threshold, we have tabulated the area-under the curve (AUC) of the receiver operating curve (ROC). Since the dataset of glycosylation efficiency measurements is highly imbalanced with only 46/361 (12.7%) glycosylatable sequons (positive samples) and 315/361 (87.3%) unglycosylatable sequons (negative samples), in Table 1 (and SI Table 2), for each metric, we also report the true positives (TP), true negatives (TN), false positives (FP) and false negatives (FN) and the balanced accuracy (BA = (TPR + TNR) / 2, where the true positive rate TPR = TP / (TP + FN) and the true negative rate TNR = TN / (TN + FP) and false positive rate (FPR = FP / (FP + TN)). Note that a naïve classifier that classifies all 361 sequons as unglycosylatable has an accuracy (ACC = (TP + TN) / (TP + TN + FP + FN)) of 87.3% (= 315/361), a BA of 50% (= (0/46 + 315/315)/2) and an FPR of 0%. Since accuracy (as opposed to balanced accuracy) higher than such a naïve classifier may come at the expense of FPs, we report balanced accuracy instead of accuracy for our dataset.

**Table 1.** Summary of AUC scores, true positives (TP), true negatives (TN), false positives (FP) and false negatives (FN), and balanced accuracy (BA=(TPR+TNR)/2; TPR=TP/(TP+FN) and TNR=TN/(TN+FP)) and false positive rate (FPR=FP/(FP+TN)) for predictions at an experimental glycosylation efficiency threshold of 10%. The calculation of TPs, TNs, FPs, FNs, BA and FPR requires a threshold. Feature thresholds were chosen in two cases (d_HB_ (largest cluster) < 4.0 Å and RMSD_peptide_ (largest cluster) < 1.0 Å) to match criteria discussed in the main text. For all other cases, thresholds were chosen arbitrarily.

| Feature | AUC | Feature threshold | TP | TN | BA (%) | FPR (%) |
| --- | --- | --- | --- | --- | --- | --- |
|  |  |  | FP | FN |  |  |
| $d_{HB}$ (largest cluster) | 0.923 | < 4.0 (Å) | 44 | 259 | 88.9 | 17.8 |
|  |  |  | 56 | 2 |  |  |
| Fraction of decoys ($d_{HB}$ < 4.0 Å) | 0.906 | > 0.70 (-) | 39 | 270 | 85.2 | 14.3 |
|  |  |  | 45 | 7 |  |  |
| RMSD<sub>peptide</sub> (largest cluster) | 0.959 | < 1.0 (Å) | 39 | 287 | 87.9 | 8.9 |
|  |  |  | 28 | 7 |  |  |
| Fraction of decoys (RMSD<sub>peptide</sub> < 1.0 Å) | 0.965 | > 0.53 (-) | 36 | 292 | 85.5 | 7.3 |
|  |  |  | 23 | 10 |  |  |
| Sc (median top 10; RMSD<sub>peptide</sub> < 1.0 Å) | 0.944 | > 0.735 | 42 | 265 | 87.7 | 15.9 |
|  |  |  | 50 | 4 |  |  |
| Interaction Energy (largest cluster) | 0.564 | < -34 (REU) | 36 | 98 | 54.7 | 68.9 |
|  |  |  | 217 | 10 |  |  |
| Interaction Energy ($d_{HB}$ < 4.0 Å) | 0.875 | < -34 (REU) | 42 | 255 | 86.1 | 19.0 |
|  |  |  | 60 | 4 |  |  |
| Interaction Energy (RMSD<sub>peptide</sub> < 1.0 Å) | 0.908 | < -34 (REU) | 39 | 271 | 85.4 | 14.0 |
|  |  |  | 44 | 7 |  |  |

### Recapitulation of amino acid specificity trends for the −1 position

#### Sampling of TS-critical hydrogen bond recapitulates specificity for 84% of the sequons with a false positive rate of 14%

In QM/MM simulations of the glycosylation of the EA2 peptide by GalNAc-T2, Gomez *et al.* characterized a hydrogen bond between the backbone amide of Thr_0_ and the β-phosphate group on UDP ^14^. They proposed that the hydrogen bond stabilizes the transition state (TS) in “a general catalytic strategy used in peptide O-glycosylation by retaining glycosyltransferases”. Hence, our first hypothesis was that successful glycosylation requires a peptide to exhibit a low-energy conformation with *d*_HB_ distances compatible with the proposed hydrogen bond. Thus, in Figure 4A, we show the heatmap of *d*_HB_ for the lowest interaction energy decoys. Applying a 4.0 Å threshold to the 19×19 grid of sequons splits the sequons into those that do not meet this condition (*i.e.,* > 4.0 Å,) and those that exhibited a representative low-energy conformation compatible with hydrogen bonding between the peptide and UDP. When compared with experimental results (Figure 1A), this criterion discriminated well between substrate peptides and non-substrates of the enzyme (Figure 4B) with 44 TPs, 259 TNs, 56 FPs and 2 FNs (Table 1).

**Figure 4.**
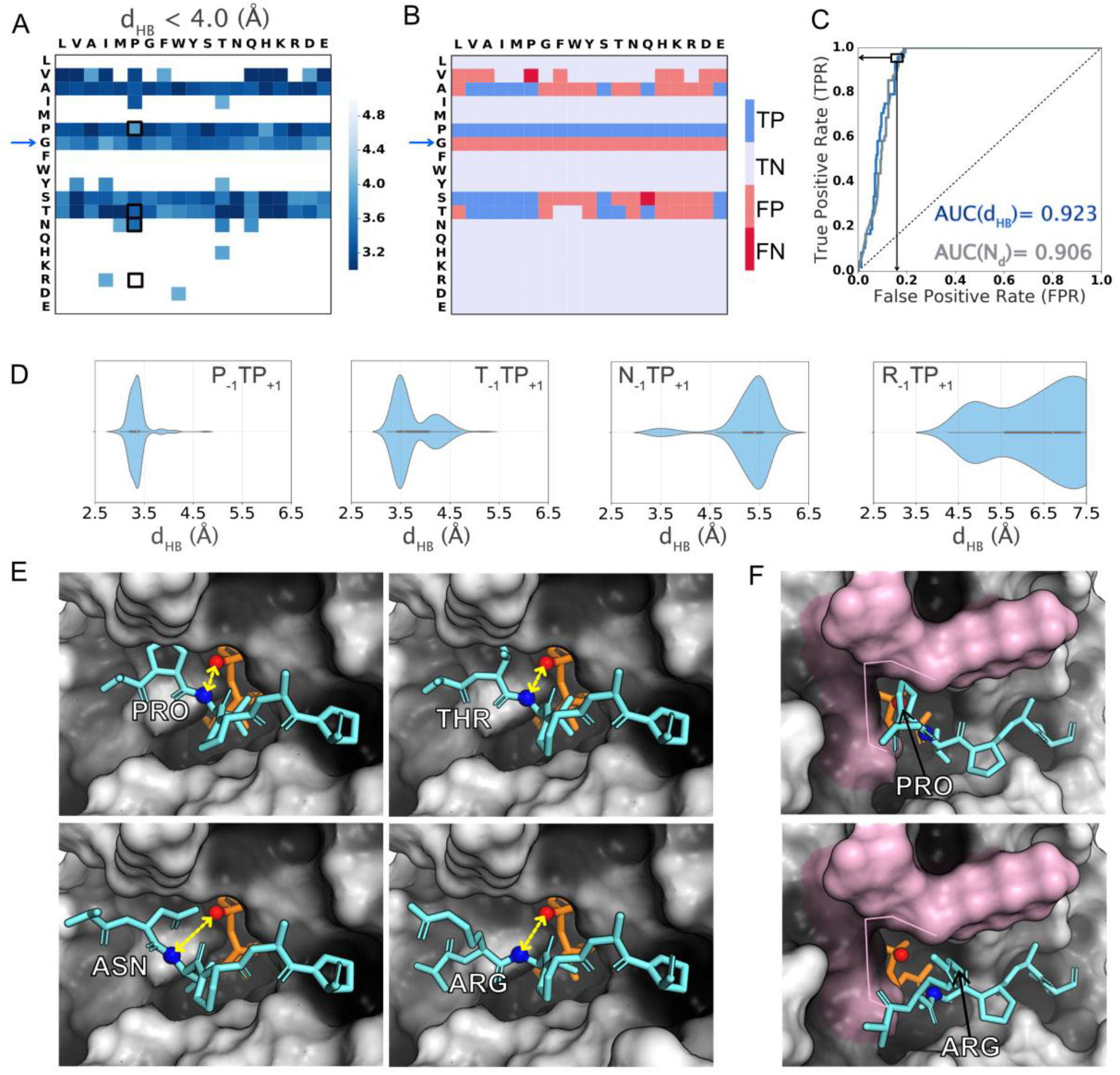
Substrate specificity based on TS stabilizing hydrogen bond criterion with *d*_*HB*_ < 4 Å. (A) Heatmaps of (left panel) *d*_HB_ distances of the lowest-interaction-energy decoy belonging to a cluster with the cluster centroid satisying the criterion. (B) True positives (TP, dark blue), true negatives (TN, light blue), false positives (FP, light red) and false negatives (FN, dark red) predicted based on ***d***_***HB***_ **< 4 Å** threshold applied to the *d*_HB_ value of the lowest-interaction-energy decoy of the largest cluster. (C) ROC curve for *d*_HB_ distances and fraction of decoys (N_d_) satisfying criterion. (D) Violinplot of distribution of *d*_HB_ distances sampled by the top-scoring 10% decoys for four representative sequons (P_−1_TP_+1_, T_−1_TP_+1_, N_−1_TP_+1_ and R_−1_TP_+1_). (E) Lowest interaction energy decoys for four sequons (P_−1_TP_+1_, T_−1_TP_+1_, N_−1_TP_+1_ and R_−1_TP_+1_ – black boxes in the heatmap in (A)). *d*_HB_ is calculated between the amide nitrogen (blue sphere) of T_0_ on peptide(aquamarine) and the O_β-PO4_ (red sphere) on UDP (orange). *d*_HB_ is shown with double-ended yellow arrows. (F) Pocket-like cavity formed by enzyme residues (pink surface) that contacts the amino acid at the −1 position on the peptide.

A ROC analysis (Figure 4C) showed that the *d*_HB_ value of the lowest-energy decoy of the largest cluster correctly distinguished the glycosylatable sequons from the non-glycosylatable sequons with a probability of 92.3% (ROC-AUC value of 0.923; Figure 4C). Setting a threshold of *d*_HB_ < 4 Å, *d*_HB_ classified with a balanced accuracy of 88.9% (TPR = 44/46, TNR = 259/316), accuracy of 84% (=303/316) (SI Table 2) and a false positive rate of 17.8% (56/(56+259)) (Table 1). If instead of *d*_HB_, we used the fraction of the of decoys (N_d_) that satisfied the *d*_HB_ < 4 Å criterion, the probability of correctly classifying a sequon was 90.6% (ROC-AUC was 0.906; Figure 4C). Setting an arbitrary threshold of N_d_ > 0.70, this feature classified with a balanced accuracy of 85.2% (TPR=39/46, TNR=270/316) of the sequons with a false positive rate of 14.3% (45/(45+270)) (Table 1). The large number of FPs for both *d*_HB_ and N_d_ indicates that the of *d*_HB_ < 4 Å criterion has low precision.

#### Large amino acid residues are excluded from GalNAc-T2’s “−1 pocket”

Figure 4A reveals that peptides preferentially exhibited low-energy, highly populated states (higher fraction of decoys) with *d*_HB_ distances compatible with hydrogen-bonding when amino acid residues with smaller side chains such as proline, alanine, glycine, serine, or threonine were present at the −1 position. To understand the structural basis for the observed *d*_HB_ trends, we considered specific sequons and their lowest-energy conformations. In Figure 4D, we show the *d*_HB_ distances sampled by the top 10% low-energy decoys for four representative sequons – P_−1_TP_+1_ and T_−1_TP_+1_ (preferentially sampled conformations with *d*_HB_ < 4.0 Å), and N_−1_TP_+1_ and R_−1_TP_+1_ (higher *d*_HB_ distances). Figure 4E shows the structures of the lowest energy conformation (largest cluster) for these four sequons. The sequons with P_−1_ and T_−1_ fit in the pocket-like cavity in the enzyme’s peptide binding groove (Figure 4E, top panels), whereas sequons with N_−1_ or R_−1_ were excluded from this cavity due to steric hinderance thereby resulting in larger distances (Figure 4E, bottom panels). Hence, the structural basis for peptides to preferentially sample low-energy conformations compatible with sampling of the proposed TS stabilizing hydrogen bond was the *relative size of the side chain of the amino acid at the −1 position and fit into the “−1 pocket” on the enzyme* (highlighted in Figure 4F; discussed in more detail later).

The case of sequon N_−1_TP_+1_ (Also V_−1_TP_+1_; SI Figure 4) is also notable because it exhibited two significant clusters, the smaller one (normalized cluster size ~ 10%) exhibiting distances compatible with hydrogen bonding (Figure 4D) and the larger one (normalized cluster size ~ 90%) with comparable interaction energy exhibiting larger distances. Experimentally this sequon was non-glycosylatable. Sequon T_−1_TP_+1_ which is experimentally glycosylatable, also exhibits two significant clusters. However, the larger cluster exhibits a *d*_HB_ distance compatible with hydrogen bonding (Figure 4D). This suggests that a larger fraction of decoys exhibiting *d*_HB_ compatible with TS-stabilizing hydrogen bond renders a sequon more glycosylatable.

For sequons with larger amino acid residues at the −1 position, characterization of the *d*_HB_ distance correlated with undetectable glycosylation in experimental assays as peptides/sequons with larger amino acids did not meet the hydrogen-bonding criteria and assumed conformations at distances farther from the UDP-GalNAc donor, making the reaction less likely. However, this *d*_HB_ criterion incorrectly classified all G_−1_ sequons. (Figure 4B; blue arrow). Since *d*_HB_ generated some false positives, especially G_−1_ peptides, the ability of the peptide to assume conformations amenable to the formation of the TS-stabilizing hydrogen bonding is a necessary but not sufficient condition to determine specificity.

#### G_−1_ results in distinct low-energy states characterized by higher RMSD_peptide_ values

To probe why G_−1_ peptides may be non-glycosylatable even though they satisfy the *d*_HB_ metric, we examined the joint distribution of the RMSD_peptide_ and *d*_HB_ sampled by the top 10% of decoys for all G_−1_ and P_−1_ peptides (*i.e.*, averaged over all 19 amino acid at the +1 position) (SI Figures 5-9 plots all 19 sequons for G_−1_, S_−1_, A_−1_, T_−1_ and P_−1_.) G_−1_ peptides exhibited two low-energy states (Figure 5A), but P_−1_ peptides primarily exhibited a single, low RMSD_peptide_ state (Figure 5B).

**Figure 5.**
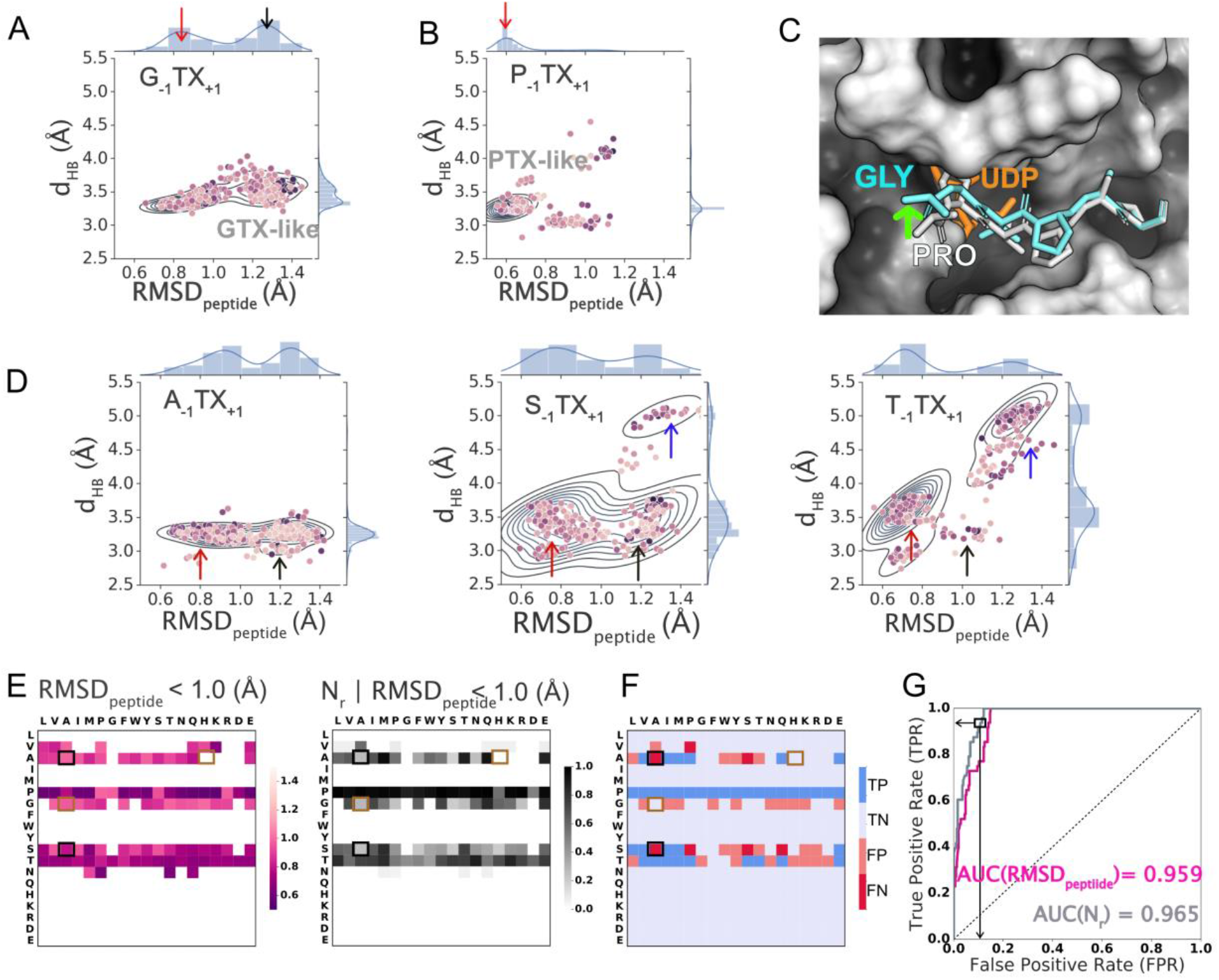
Substrate specificity based on RMSD_peptide_ < 1.0 Å criterion. Joint and marginal probability densities for the top 10% of structures by score (200/2000) for a given amino acid at the −1 position and aggregated over all amino acids at the +1 position. (A) G_−1_ and B) P_−1_ and all amino acid residues at X_+1_; Top 1% (20/2000) decoys per sequon shown as points where darker color indicates lower interaction energy. “GTX-like” state is marked with a black arrrow and “PTX-like state” is marked with a red arrow. (C) Lowest interaction energy decoy in the enzyme’s peptide binding groove for representative sequon P_−1_TP_+1_ (white; RMSD_peptide_ < 1.0 Å) superposed with that for G_−1_TP_+1_ (aquamarine; RMSD_peptide_ > 1.0 Å). (D) Joint and marginal probability densities of Top 10%(200/2000) sequons for all peptides with fixed amino acids A_−1_, S_−1_, T_−1_ and all amino acid residues at X_+1_; Top 1% (20/2000) decoys per sequon shown as points where darker color indicates lower interaction energy. The blue arrow indicates a third state distinct from PTX- and GTX-like states. (E) Heatmap of RMSD_peptide_ (left panel) of the lowest energy decoy per sequon, and fraction of decoys (N_r_) satisfying RMSD criterion (right panel) for RMSD_peptide_ < 1.0 Å. (F) True positives (TP), true negatives (TN), false positives (FP) and false negatives (FN) predicted based on RMSD_peptide_ < 1 Å threshold applied to the lowest-interaction-energy decoy of the largest cluster. (G) ROC curve for RMSD_peptide_ (magenta) and N_r_ (grey) satisfying RMSD_peptide_ < 1.0 Å. Black and brown boxes in (E) and (F) indicate examples of glycosylatable and non-glycosylatables sequons, respectively, that also exhibit the GTX-like states.

#### Amino acid residues with smaller side chains are sub-optimal for −1 pocket of the enzyme

In Figure 5C, we have superposed the lowest-energy decoys for sequons P_−1_TP_+1_ and G_−1_TP_+1_, representative of P_−1_ and G_−1_ peptides, respectively. For G_−1_TP_+1_, the backbone was shifted “up” with respect to that of the P_−1_TP_+1_ backbone (Figure 5C; green arrow). For G_−1_ peptides, the small size of the glycine residue allowed multiple configurations in the −1 pocket of the enzyme, all of which still made the TS stabilizing hydrogen bond (i.e., < 4.0 Å). Consequently, we also observed the “GTX-like state” (defined as RMSD_peptide_ ≥ 1.0 Å and < 4.0 Å and marked in Figure 5 with black arrows) for A_−1_ and S_−1_ (shorter side chains) peptides (Figure 5D) but not for T_−1_ peptides. Instead, T_−1_ peptides exhibited a third state (Figure 5D; blue arrow) which we discuss later. Thus, while the *d*_HB_ metric explained why sequons with larger side chains at the −1 position were non-glycosylatable, the RMSD_peptide_ metric explains why certain sequons with smaller side chains at the −1 position may not be suitable for glycosylation.

#### RMSD_peptide_ metric improves sequon specificity predictions for G_−1_ peptides and recapitulates specificity for 90% of the sequons

P_−1_ peptides, irrespective of the amino acid at the +1 position, experimentally exhibited high glycosylation efficiencies and also primarily exhibited the low-RMSD_peptide_ or the PTX-like state (defined as RMSD_peptide_ < 1.0 Å and < 4.0 Å and marked in Fig. 5 with red arrows). This leads us to hypothesize that besides the TS-stabilizing hydrogen bond (characterized by *d*_HB_), the second factor that determined the glycosylatability of a sequon was the precise positioning of the peptide in the enzyme’s peptide binding groove, *i.e.*, how close the peptide backbone was, spatially and conformationally, to the cognate sequon peptide conformation in the crystal structure. We postulated that the “PTX-like state” (red arrows in Fig. 5D) with RMSD_peptide_ < 1.0 Å leads to successful glycosylation (reactive state) whereas all other conformations or states with RMSD_peptide_ ≥ 1.0 Å (e.g. the GTX-like state) did not lead to glycosylation (non-reactive).

Hence, we used the sampling of the PTX-like state by the top-scoring decoys, quantified by the RMSD_peptide_ of the lowest energy decoy of the largest cluster and the normalized size of the largest cluster, as the second criterion for successful glycosylation that includes multiple low-energy conformations. This criterion improved prediction for sequons that exhibited low-energy conformations in the GTX-like state, (A_−1_TH_+1_, A_−1_TG_+1_, S_−1_TG_+1_, G_−1_TG_+1_, etc.), including G_−1_ peptides and was able to correctly classify many such peptides as non-glycosylatable (Figure 5 E, F). When compared with experimental results, this criterion, based on the RMSD_peptide_ value of the lowest-energy decoy of the largest cluster as a classification metric for glycosylation gives a ROC-AUC value of 0.959 (Figure 4G, Table 1). Setting a threshold of RMSD_peptide_ < 1.0 Å, the criterion correctly classifies 90.3% ((39+287)/361) of sequons with a BA of 87.9% and an FPR of 8.9% (28/(28+287)) (Figure 4F, Table 1). When the fraction of decoys (N_r_) that satisfy the RMSD_peptide_ < 1 Å condition is the classification metric, the ROC-AUC value was 0.965 (Figure 5G). Setting an arbitrary threshold value of N_r_ > 0.53, N_r_ with RMSD_peptide_ < 1 Å correctly classified 90.8% ((36+292)/361) of the sequons with a BA of 85.5% and an FPR rate of 7.3% (23/(23+293)) (Table 1).

Note that for most conformations that satisfy RMSD_peptide_ < 1.0 Å, the condition *d*_*HB*_ < 4.0 Å is also satisfied (Figures 5A, B, D).

However, since both classes of sequons, those that are glycosylatable experimentally (*e.g.* A_−1_TA_+1_ and S_−1_TA_+1_, Figure 5F black boxes) and those that are non-glycosylatable experimentally (*e.g.* G_−1_TA_+1_ and A_−1_TH_+1_Figure 5F brown boxes), exhibited non-reactive states, the criterion based on RMSD_peptide_ was not sufficient to correctly classify all peptides, especially for sequons that exhibited both reactive and non-reactive states with similar interaction energies and/or similar fraction of decoys.

#### Amino acid residue at the −1 position dictates the low-energy conformations and glycosylatibility for the majority of the sequons

The analysis of the low-energy conformations characterized by *d*_HB_ and RMSD_peptide_ lead to the following observations. For the majority of sequons, those with K_−1_, R_−1_, F_−1_, Y_−1_, W_−1_, D_−1_, E_−1_, Q_−1_, N_−1_, H_−1_, I_−1_, M_−1_, L_−1_, or V_−1_, the peptide primarily sampled non-reactive low-energy conformations with *d*_HB_ > 4.0 Å and RMSD_peptide_ > 1.0 Å. For a small fraction of sequons (P_−1_ peptides), the peptide primarily sampled a reactive, cognate-sequon like state (or PTX-like state) with *d*_HB_ < 4.0 Å and RMSD_peptide_ <1.0 Å. For both of these categories that primarily sample one state—either the non-reactive state or the reactive-state—the computational predictions based on either hypothesis (RMSD_peptide_ or *d*_HB_), agreed quite well with experimental data. These observations underscore the importance of the residue at the −1 position in determining the low-energy conformations and, consequently, the glycosylatability for the majority of the sequons (~ 15 × 19 = 285 out of 361 peptides). However, four amino acids (G, A, S, T) at the −1 position are yet ambiguous, showing two states. Sometimes they are classified correctly, but sometimes not.

### Recapitulation of amino acid specificity trends for the +1 position

For G_−1_, A_−1_, S_−1_, T_−1_ sequons (4 × 19 = 76 out of 361), the peptide sampled both reactive and non-reactive states with comparable interaction energies. For many of these sequons, the computational predictions based on the effect of the −1 position did not accurately recapitulate experimental observations. Hence, for G_−1_, A_−1_, S_−1_, T_−1_, to recapitulate experimental glycosylation trends, we must consider the effect of the +1 position.

#### Amino acid at the +1 position confers secondary effects that modulate effects of the −1 position

To investigate the effect of the +1 position for G_−1_, A_−1_, S_−1_ peptides, all of which exhibit the GTX-like state, we considered the variation in sampling of the GTX-like state as a function of the amino acid at the +1 position. Figure 6A shows these fractions for a subset of sequons, viz. the G_−1_, A_−1_, S_−1_, and T_−1_ peptides. For T_−1_ peptides, no sequon exhibited the GTX-like state for a significant fraction of the decoys. For A_−1_, S_−1_ and G_−1_ peptides, G_+1_ and D_+1_ significantly increased the propensity to sample (indicated by a large fraction of decoys) the GTX-like state. Furthermore, for A_−1_ peptides, H_+1_, K_+1_, R_+1_, and S_+1_ also resulted in a large fraction of decoys exhibiting the GTX-like state. The interaction energies of the lowest-energy decoys of the GTX-like state are comparable to those of the PTX-like state (SI Figure 10A), suggesting that such a state could dominate or compete with the PTX-like. Hence, the +1 position, in these specific cases, enhanced the sampling of the non-reactive, GTX-like state, modulating the glycosylatability of a peptide in a capacity secondary to the −1 position.

**Figure 6.**
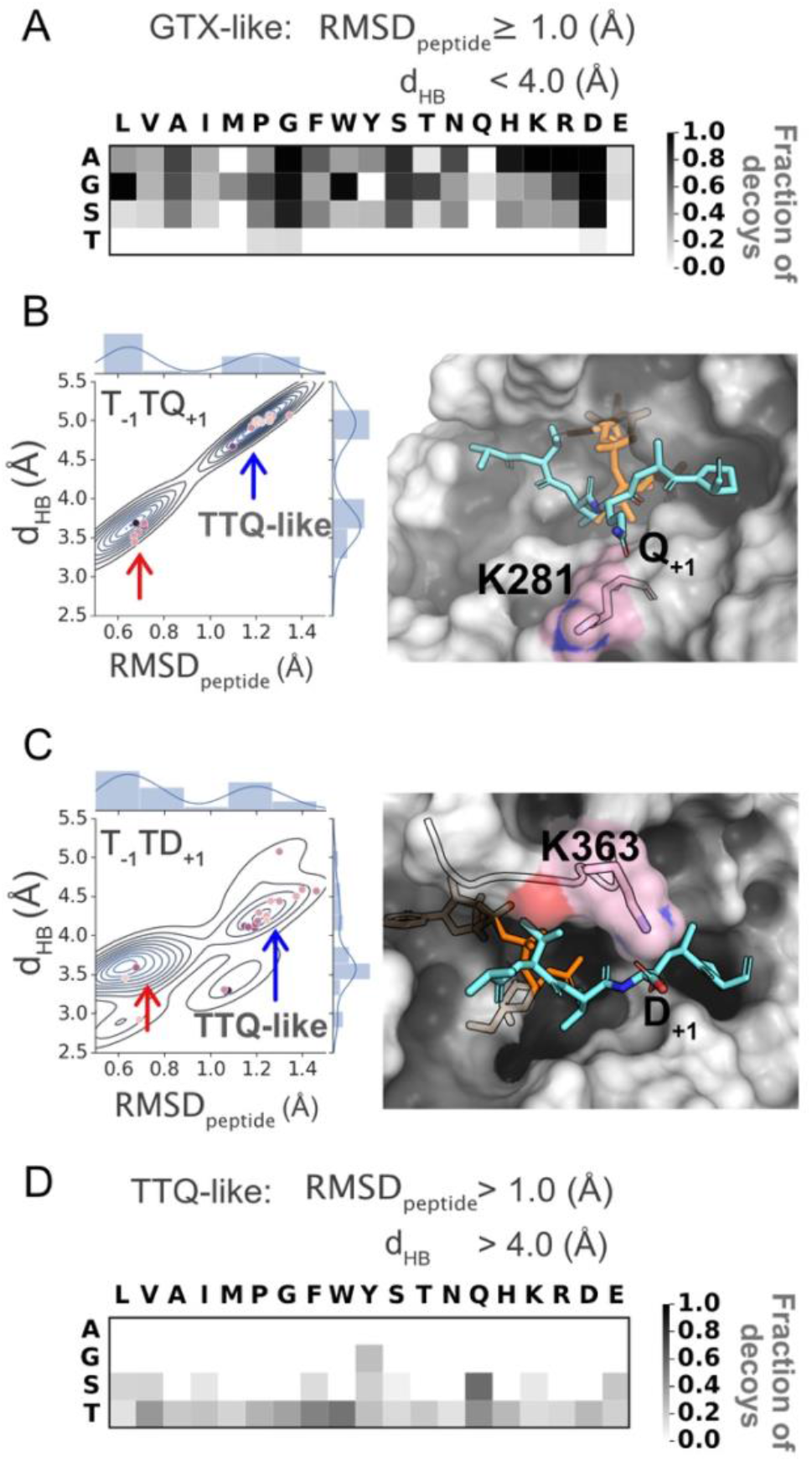
Secondary effects of the amino acid at the +1 position. (A) Fraction of decoys sampling the GTX-like state for sequons with A, G, S, or T at the −1 position. (B) Joint and marginal probability densities of top 10% (200/2000) sequons for T_−1_TQ_+1_ (left panel) and lowest energy decoy for TTQ-like state for sequon T_−1_TQ_+1_ state, where Q_+1_ position interacts with K281. (C) Joint and marginal probability densities of top 10%(200/2000) sequons for T−1TD+1 (left panel) and lowest energy decoy for TTQ-like state for sequon T_−1_TD_+1_ state, where D_+1_ position interacts with K363. (D) Fraction of low-energy decoys in the TTQ-like state. Top 1% (20/2000) decoys per sequon shown as points in (B) and (C) where darker color indicates lower interaction energy. “TTX-like” state is marked with a black arrrow and “PTX-state” is marked with a red arrow.

We used these observations to test a classification based on the sampling of the GTX-like state. While the changes in classification accuracy are negligible, the prediction improves for sequons A_−1_TW_+1_, A_−1_TY_+1_, A_−1_TT_+1_ and T_−1_TG_+1_, summarized in SI Table 2 and SI Figure 11B.

#### Residues glutamine, glutamate, aspartate and the aromatics at the +1 position interact with residues K363/K281 on the enzyme to form competing states

To understand the variation in glycosylatability with the +1 position for T_−1_ peptides, we examined the sequons T_−1_TQ_+1_, T_−1_TF_+1_, T_−1_TY_+1_ and T_−1_TW_+1_. Experimentally, T_−1_TQ_+1_ was glycosylatable with ~20% activity, whereas T_−1_TF_+1_, T_−1_TY_+1_ and T_−1_TW_+1_ were non-glycosylatable. All four sequons exhibited the PTX-like state (red arrow in Figure 6B and SI Figure 12). These sequons additionally exhibited a second, low-energy state with RMSD_peptide_ > 1.0 and > 4.0 Å (blue arrow in Figure 6B and SI Figure 12). In this state, the residue T_−1_ occupied the −1 pocket similar to the PTX-like state, while the residue Q_+1_ interacted with residue K281 on the enzyme, which lies at the rim of the peptide-binding groove (Figure 6B). The interaction between the residues Q_+1_ and K281 pulled the peptide backbone away from the catalysis site (Figure 6B), resulting in a non-reactive state that competed with the reactive PTX-like state. We observed a similar interaction for residue D_+1_ (sequons T_−1_TD_+1_ and P_−1_TD_+1_), however, due to a shorter side chain compared to Q_+1_, it was in a better position to interact with K363 residue (Figure 6C, SI Figure 12D).

In Figure 6D, we show the sampling of the “TTQ-like state” (*d_HB_* > 4.0 and RMSD_peptide_ > 1 Å) for A_−1_, G_−1_, S_−1_, and T_−1_ peptides. The TTQ-like state was observed primarily in S_−1_ and T_−1_ peptides. F_+1_, Y_+1_, W_+1_, M_+1_, I_+1_, E_+1_, Q_+1_, H_+1_, and D_+1_ exhibited highly stabilized TTQ-like states. For non-polar residues such as M_+1_, and I_+1_, the stabilization arose from non-polar interactions of the +1 side chain with the K281 side chain.

For sequons that exhibited both states, we computed the energy difference between the lowest-energy decoys for the two states (SI Figure 10B). Similar to the GTX-like state, the interaction energy of the lowest-energy decoys for the TTQ-like state is comparable to that of the PTX-like state. For sequons T_−1_TD_+1_, T_−1_TW_+1_ and T_−1_TY_+1_, the lowest interaction energy of the TTQ-like state was about 2 REU lower than that of the PTX-like state. For T_−1_TQ_+1_, the difference was small (−0.2 REU), and for T_−1_TE_+1_, the PTX-like state was more stable by 2.4 REU. The relative stabilization of the PTX-like state over the TTQ-like state as measured by interaction energy 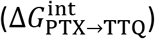 correlated with higher experimental glycosylation efficiencies for sequons T_−1_TQ_+1_ (^~^ 22%) and T_−1_TE_+1_ (~13%) compared to T_−1_TD_+1_ (3%) and T_−1_TX_+1_ (0%), where X was an aromatic residue.

To quantify the interaction energy of different amino acid residues at the +1 position to specific residues on the enzyme, we computed the pairwise energies of interaction between the residue at the +1 position and the enzyme (SI Figure 13) and, as expected, found that the residues that exhibit the TTQ-like state interact favorably with residues K281 or K363 on the enzyme. On the other hand, residues P_+1_ and A_+1_ did not interact with K281 or K363 residues on the enzyme. The lack of interaction with K281 or K363 residues on the enzyme suggested that sequons T_−1_TP_+1_, T_−1_TA_+1_, S_−1_TP_+1_ and S_−1_TA_+1_ had no propensity for the TTQ-like state and may explain the high glycosylation efficiencies observed for these sequons.

With these observations, we tested using TTQ-like state populations for classifying the substrate and non-substrate sequons (classification accuracy in SI Table 2 and SI Figure 11B). While the change in classification accuracy is negligible, the consideration of the TTQ-like state improves predictions for sequons T_−1_TY_+1_, T_−1_TW_+1_ and T_−1_TD_+1_ (SI Figure 11B).

### Characterization of the peptide-enzyme interface

#### Shape complementarity and hydrogen bonding contribute to the finely tuned specificities at the −1 and +1 positions

So far, our analysis focused on analyzing the landscape of low-energy conformations exhibited by the peptides and on recapitulating the experimentally observed specificity trends as a function of the amino acid at the −1 and +1 position of the sequon. In this process, we discovered the dominant modes of interaction between the peptide and the enzyme that lead to reactive (PTX-like) and non-reactive (GTX-like and TTQ-like) conformations. Comparison between experimental data and computational predictions also revealed that a majority of sequons that were glycosylatable exhibited a PTX-like conformation. Next, we characterize the PTX-like state to decipher the structural basis for the variation of specificity within the subset of peptides that exhibited this state.

First, we calculated the shape complementarity,^43^ *S*_*c*_, for the enzyme-peptide interface for all sequons for the ten lowest-interaction-energy decoys that satisfied the RMSD_peptide_ < 1.0 Å criterion (Figure 7A,C). P_−1_ peptides exhibited the highest shape complementarity at the peptide-enzyme interface because it packs against the planar interface formed by a histidine residue at position 365 on the enzyme (Figure 6E). We further characterized the residue-wise and pairwise interaction energies at the interface. The P_−1_ residue exhibited significantly higher attractive van der Waals energy (fa_atr term in Rosetta) with H365 of the enzyme than any other residues at the −1 position (Figure 7B). The P_−1_ residue exhibited generally higher attractive van der Waals energies with all enzyme residues at the interface (SI Figure 14). T_−1_, S_−1_, and A_−1_ residues exhibited energies (SI Figure 14, Figure 7B) and shape complementarities (Figure 7A, C) that varied to a significant extent with the residue at the +1 position. Thus for these sequons, the +1 position may additionally contribute to anchoring the peptide in the binding cavity.

**Figure 7.**
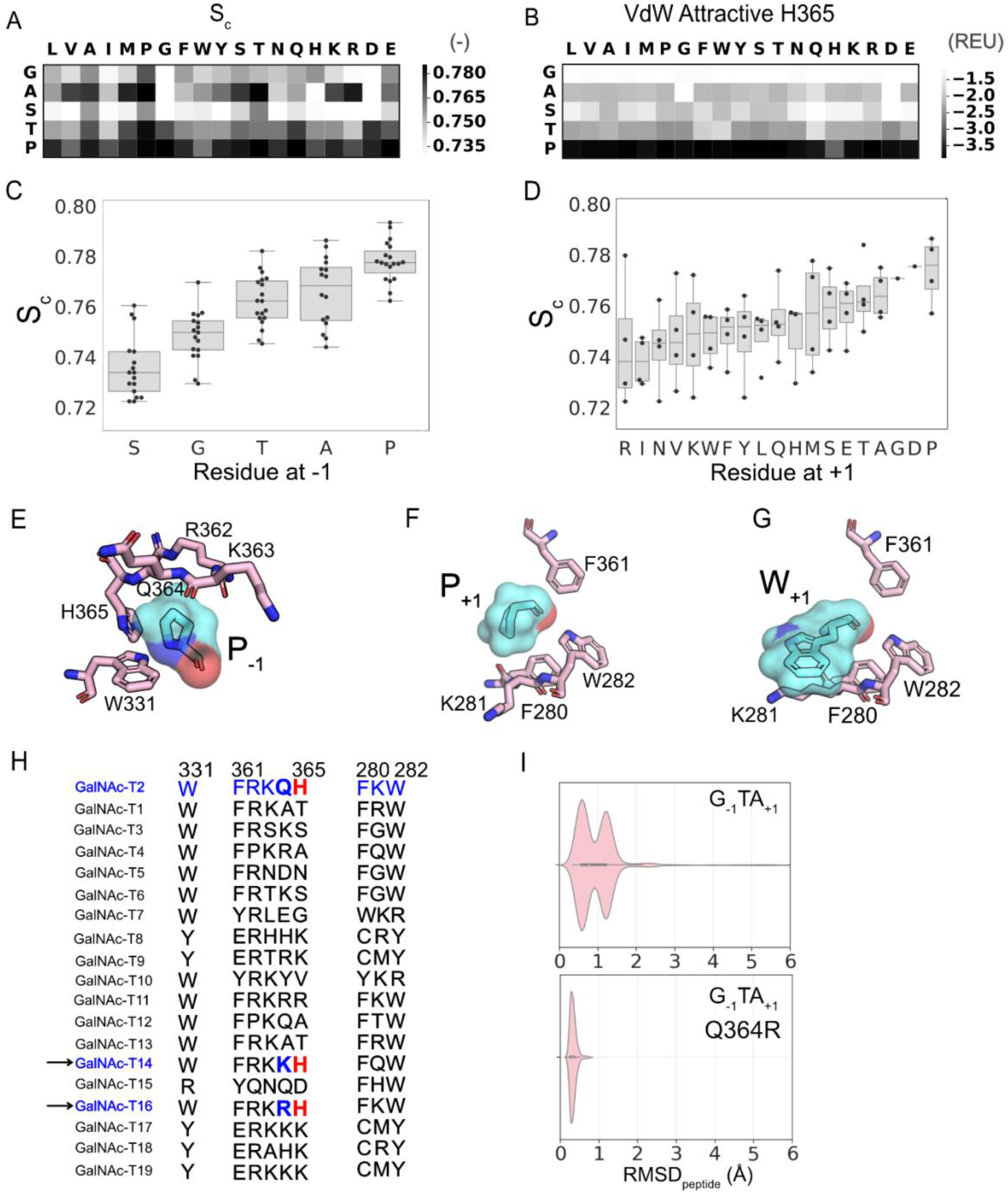
Characterization of enzyme–peptide interactions for top 10 decoys G_−1_, A_−1_, S_−1_, T_−1_, P_−1_ peptides. (A) Median shape complementarity (S_c_). (B) Attractive component of the van der Waals (VdW) potential in Rosetta score function between the residue at the −1 position and H365 on the enzyme. (C) Distribution of median S_c_ values as a function of the residue at −1 position. (D) Distribution of median Sc values as a function of the residue at +1 position. (E) +1 pocket of at the enzyme peptide interface with H365 on the enzyme (pink) interacting with the proline at the −1 position on the peptide (aquamarine). (F) Residues 280, 281, 282 and 361 on the enzyme (pink) interacting with proline at the +1 position on the peptide (aquamarine). (G) Residues 280, 281, 282 and 361 on the enzyme (pink) interacting with tryptophan at the +1 position on the peptide (aquamarine). (H) Multiple sequence alignment of isoform T2 with other isoforms for the residues at the enzyme-peptide interface for +1 and −1 positions on the peptide. (I) Violinplots for RMSD_peptide_ distributions sampled for sequon GTA for isoform T2 (top) and a variant T2-Q364R (bottom). Residue numbering based on GalNAcT-2 Uniprot entry Q10471.

In the +1 position, proline exhibited the highest shape complementarity (Figure 7D) in the “+1 pocket” formed by three aromatics F280, W282 and F361, stabilized by favorable interactions between the partially positively charged proline ring and the partially negatively charged π faces of aromatic side chains (Figure 7F, SI Figure 15). Also, similar to the TTQ-like state, sequons with aromatics, glutamine, glutamate and non-polar residues other than alanine, proline and glycine at the +1 position interacted with K281 on the enzyme (Figure 7G).

Surprisingly, the median *S*_*c*_ of the top ten decoys that satisfied the RMSD_peptide_ < 1.0 Å criterion is a reasonably good classifier of sequon glycosylatability with a ROC-AUC score of 0.944.

For T_−1_ and S_−1_ residues, the PTX-like state was additionally stabilized by a hydrogen bond between the hydroxyl side chain and the backbone carboxyl of R362 on GalNAcT2 (SI Figure 16).

In summary, the shape complementarity and pairwise-energies describe a −1 pocket that is highly specific for the P_−1_ residue, underlying the high experimental glycosylation efficiencies measured for P_−1_ peptides.

#### Sequence motifs at the −1 pocket hint at modes of specificity modulation across isoforms T2, T14 and T16

The −1 pocket on the enzyme plays an important role in screening for optimally-sized side chains at the −1 position of the sequon. This pocket is primarily formed by residues R362, K363, Q364, H365 and W331. These residues determine the size and chemical composition of the −1 pocket. The residues R362, K363, Q364, and H365 reside on the flexible, semi-conserved catalytic loop^44^ of GalNAcT2. This flap-like loop can additionally contribute to the variability of the −1 pocket size across the GalNAc-T isoforms. Among the three isoforms of the GalNAc-T family that show a strong preference for P_−1_ (T2, T14, T16), the H365 residue is conserved (Figure 7H). Residues K363 and Q364 reside at the point of entry for the −1 residue on the peptide. Variation of amino acids at these positions could allow for variation in the size of the amino acid preferred at the −1 position of the sequon. For example, isoforms T14 and T16, which are evolutionary most proximal to T2, have residues lysine or arginine at position 364. Unlike T2, both T14 and T16 prefer G_−1_;^45^ indicative of a −1 pocket suitable for smaller sidechains. In fact, when we repeated MCM sampling of the G_−1_ sequons for the T2 isoform with the Q364R mutation, we observed a complete shift towards conformations with RMSD_peptide_ < 1.0 Å (PTX-like state) and the elimination of the GTX-like state (Figure 7I), suggesting a possible strategy for varying the peptide substrate preference of various isoforms.

#### Energy-based predictors incorrectly classify P_−1_ peptides as non-glycosylatable

Energy is a commonly used metric in determining specificity of peptide substrates (eg. pepspec^31^, sequence_tolerance^32^ and MFPred^34^). In this work, we characterized each cluster by interaction energy and used it as the metric for choosing the top decoys for further analysis. However, for the purpose of prediction of specificity trends for the T2 enzyme, interaction energy by itself was a weaker predictor than other metrics (AUC = 0.566, Table *1*). The significantly lower AUC values based on interaction energy, compared to RMSD_peptide_ (0.959) and fraction of decoys (0.969), were due to the fact that P_−1_ and S_−1_ peptides bind the enzyme with significantly lower interaction energies than A_−1_, G_−1_ or T_−1_ peptides (Figure 2A).

For comparison with other energy based approaches, we applied MFPred^34^ to obtain the specificity profile for GalNAc-T2 (SI Figure 17). MFPred uses a mean-field approach that assumes each residue position is independent. MFPred obtains an AUC score of 0.68 at the −1 position and 0.50 at the +1 position (SI Table 4). While MFPred is able to predict a preference for T_−1_ and S_−1_ residues, it fails to identify P_−1_ and A_−1._ For the +1 position, MFPred performs much worse. For comparison of experimental and MFPred specificity logos, see SI Figure 17. Both of these AUC values as well as the average AUC over all positions predicted by MFPred are lower than those obtained with the *d_HB_* and RMSD_peptide_ criteria (SI Table 4). We note here that in the MFPred study, the addition of structural-motif-preserving constraints improved predictions for some PBDs. For the MFPred scores here, we did not add any constraints. We also note that the improved performance of our approach compared to MFPred is not surprising as MFPred is a general purpose specificity predictor and does not require system-specific features unlike our approach that was specifically developed for elucidating the specificity of the GalNAcT family.

## Discussion

In this work, we attempted to understand the structural basis for the peptide substrate preferences of the T2 isoform of the GalNAc-T family. We expect this work to be useful in understanding how the preference for different peptide substrates is modulated across the 20 isozymes of this family.

We used a flexible backbone protocol with MCM sampling, resulting in more than one low-energy peptide conformation/state in the vicinity of the starting peptide conformation. Most existing protocols for determining peptide specificity for peptide binding domains employ limited backbone sampling, generating ensembles close to the starting structures (pepspec, MFPred) and usually employing additional constraints to sample TS-like conformations. While these studies have been successful at predicting specificity trends, a wealth of information can be garnered from sampling the peptide landscape without imposed constraints. Our work benefitted from the availability of crystal structures for the peptide-enzyme complex but may be less accurate in the absence of crystal structures. Our approach also suffered from inaccuracies in the Rosetta energy function, the limitations of MCM sampling, and the use of implicit solvation models to name a few. We further note that an MD-based simulation, though computationally prohibitive for a large dataset, may be better suited for generating thermodynamically accurate ensembles and for characterizing the density of multiple stable states.

We investigated a range of features to predict the glycosylation efficiency of GalNAcT-2 (Table 1) and found that the features *d*_HB_ and RMSD_peptide_ are able to recapitulate binarized glycosylation specificity with a balanced accuracy of 88.9% and 87.9% and a false positive rate of 17.8% and 8.9% respectively. Alternatively, the fraction of decoys, N_d_ and N_r_, that satisfy criterion d_HB_ < 4.0 Å and RMSD_peptide_ < 1.0 Å respectively, recapitulated specificity with a balanced accuracy of 85.2% and 85.6% and a false positive rate of 14.3% and 7.3% respectively.

Additionally, we found energy-based predictors (based on MFPred and interaction energy in this work) to be poor predictors of specificity, especially in the absence of structural-motif-preserving constraints. This suggests that the stability of the peptide-enzyme complex or the interaction-energy at the interface, by itself, is a weak indicator of efficient catalysis by GalNAcT-2. In fact, since selective stabilization of the transition state over the reactants is important for catalysis, the over-stabilization of the reactant state (indicated by higher interaction energies) may increase the free energy of activation (difference between the energy of reactants and the transition state) thereby slowing or preventing the reaction. The addition of constraints could partially alleviate this issue by restraining the enzyme-peptide complex in a configuration mimicking the transition state. However, the addition of constraints will omit the sampling of potential low energy states that may compete with or hinder the formation of the transition state. Such states can only be identified in protocols that allow flexible sampling of the backbone without constraints.

The −1 position on the peptide strongly determined the glycosylation efficiency. Residues R362, K363, Q364 and H365 on the catalytic loop and residue W331 on the enzyme form the −1 pocket and select for amino acids threonine, proline, serine, alanine or glycine at the sequon’s −1 position. For sequons with residues that did not fit this pocket, the peptide was not able to form a hydrogen bond with UDP that has been proposed to stabilize the TS. We further found that this pocket was especially favorable for recognizing peptides with proline at the −1 position, as demonstrated by highly favorable interactions between H365 and proline and a high degree of shape complementarity at the +1 position irrespective of the amino acid. Hence, by changing the size and other biophysical aspects of this pocket, the specificity for the −1 position can be potentially modulated. These structural and sequence features are especially relevant for specificity modulation across isoforms, as the GalNAc-T family can glycosylate a wide range of amino acids at the −1 position.

We additionally found that residues K281 and K363 acted as gating residues by interacting with peptide amino acid residues, such as Q+1 and D+1, leading to low-energy states that compete with the reactive state. Hence, the specificity for the +1 position may be modulated by altering the lysine residues at positions 281 and 363 on the enzyme. Similar to the −1 position, such variation in specificity for the +1 position is already observed in the GalNAc-T family as certain isoforms (GalNAc-T1^12^, GalNAc-T14^45^) are capable of efficiently glycosylating D_+1_.

Key structural motifs identified in this work may be important for designing more promiscuous forms of the enzyme or tailored forms with specificities different from those seen in the 20 naturally occurring isoforms. Furthermore, since many members of the GalNAc-T family have been associated with various cancers, the sequence and structural motifs identified in this work may help decipher mutations that cause aberrant glycosylation.

## Methods

### Starting structure for enzyme–peptide complex

The primary starting structure of the enzyme–peptide complex was obtained from the crystal structure of the active conformation of GalNAcT2 from the crystal structure of the complex (pdb id: 2ffu). Since the sugar is absent from that structure, we used a second GalNAcT2 structure (pdb id: 4d0z) with bound peptide (mEA2), manganese and UDP-GalNAc-5S. While the sugar bound to UDP in 4d0z has a modification (sulfur instead of oxygen in the ring), it aligns exactly with 2ffu with the additional sugar (SI 12). To generate the starting structure for each sequon, we used the crystal structure of the complex replacing the mEA2 peptide peptide with A_−2_X_−1_T_0_X_+1_A_+2_P_+3_R_+4_C_+5_, where X is any amino acid residue except cysteine. Residues at positions −1 and +1 (denoted by Xs) were mutated to the target sequon for all 361 sequons studied in the work by Kightlinger *et al.*^12^ using Rosetta’s MutateResidue mover followed by side chain repacking and minimization using the PackRotamersMover. No backbone motion is allowed at this stage.

### Rosetta protocol for generating decoys

The glycosylation protocol is based on the flexpepdock^37,38^ protocol with a few modifications. There are two main stages– 1) Low-resolution sampling with the centroid score (united atom) 2) High-resolution refinement with the all-atom ref2015 score function.^46^ In the low-resolution phase, we use simulated annealing for enhanced sampling of the peptide. We vary the temperature from 2.0 to 0.6 in Rosetta temperature units (kT) over 30 Monte Carlo (MC) cycles. For each temperature cycle of simulated annealing, we use 50 inner MC cycles are used for perturbation followed by minimization in rigid body (across enzyme–peptide interface) and torsional (peptide) space. “Small” and “shear” movers from Rosetta are used for torsional sampling of the peptide^47^ with rigid body perturbations using the RigidBodyPerturbMover. The final pose from the low-resolution stage is passed to the high-resolution stage. In the high-resolution stage, the attractive and repulsive potential weights are ramped down and up respectively over 10 outer cycles. Similar to the low-resolution stage, we apply rigid body sampling across the enzyme–peptide interface and torsional sampling of the peptide backbone followed by minimization and Metropolis criterion. Additionally, both rigid body moves (30 cycles) and torsional moves (30 cycles) are accompanied by peptide side chain repacking every cycle and the interface side chains every 3^rd^ cycle^47^. We used the default distance of 8 Å to define the interface. Additionally, the run was terminated if the peptide moved more than 8 Å away from the enzyme–peptide interface. The backbone of the enzyme is fixed throughout sampling. We generated 2000 decoys per sequon. Larger number of decoys (8000) were not found to alter the results.

This protocol is available in the Rosetta suite. See Supplementary Information for the complete list of steps to run the protocol. The protocol is run from command-line as follows:

~~~
>mucintypeglycosylation.<system><compiler><mode> @flags
System=linux; compiler=gcc; mode=release
Where “flags” is a plain text file and contains the following options:
-in:file:s <input pdb file>
-in:file:native <input pdb file>
-nstruct 2000 #no. of decoys
-residue_to_glycosylate 3P #Threonine on peptide chain P
-substrate_type peptide
-glycosylation_low_res_refinement #enables low resolution stage
-tree_type docking
-sugardonor_residue 495 #residue number of UDP
-enable_backbone_moves_pp #enables peptide backbone moves in high resolution
-ex1
-ex2aro
-nevery_interface 3 # pack enzyme peptide interface every 3 MC cycles in high resolution
-ntotal_backbone 30 # run 30 MC cycles in High resolution
-output_distance_metrics #output rmsd, distance, interaction energies to score file
~~~

### Clustering and analysis of decoys

The top 10% decoys (200/2000) were clustered using the dbscan clustering algorithm^48,49^ in sklearn^50^ with parameters set to eps = 0.3 Å (maximum distance between samples for one to be considered in the neighborhood of the other) and min_samples = 10 (number of samples in the neighborhood of a point to be considered a core point).

### Calculation of features

We report two RMSD metrics in this work – RMSD_peptide_ and RMSD_sequon_. Both metrics are calculated over backbone C_α_ atoms only with respect to the backbone of the peptide in the starting structure. For RMSD_peptide_, RMSD is calculated over all peptide positions (8). For RMSD_sequon_, RMSD is calculated for positions −1 to +3 (XTXAP). Shape complementarity^43^ is calculated using PyRosetta^51^ as described in Supplementary Information. The interaction energy at the enzyme–peptide interface is calculated as the difference between the ref2015 score for the bound complex and the ref2015 score for the enzyme (includes the UDP-sugar molecule) and the peptide, separated from the complex without relaxing or repacking side chains.

### Specificity prediction with MFPred

We used MFPred as described:^34^ 1) The starting structure was relaxed. 2) The lowest energy decoy from relax step was used as the starting structure for the FastRelax protocol for each sequon. 3) The lowest energy decoy for each sequon from the FastRelax protocol was processed by the GenMeanFieldMover.

All calculations were performed with Rosetta version [details in Supplementary Information; www.rosettacommons.org/software].^42^

## Supporting information

Additional code and pdb files

## Acknowledgements

We thank Nadine L. Samara at National Institutes of Health for helpful discussions and for critically reading and editing the manuscript. We also thank Weston Kightlinger, Liang Lin and Michael C. Jewett at Northwestern University for sharing data and helpful discussions.

