## Supplementary material for "Structural basis for peptide substrate specificities of glycosyltransferase GalNAc-T2": SI Figures

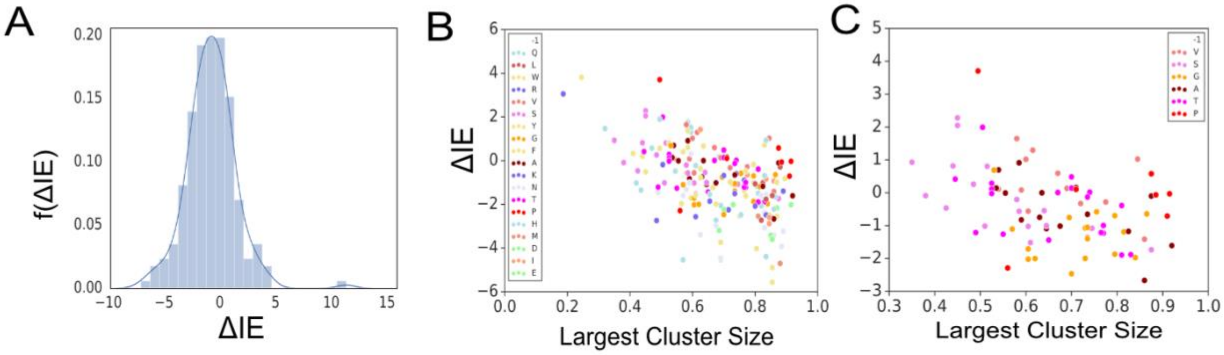


**SI Figure 1. Relation between lowest interaction energies and cluster sizes of top two clusters. A) Distribution of difference between lowest interaction energy of largest and second largest cluster (ΔIE). B) ΔIE versus size of largest cluster for sequons colored by the amino acid at the -1 position. C)**

**ΔIE versus size of largest cluster for a subset of the sequons (A, G, P, T, S at the -1 position) colored by the amino acid residue at the -1 position.**

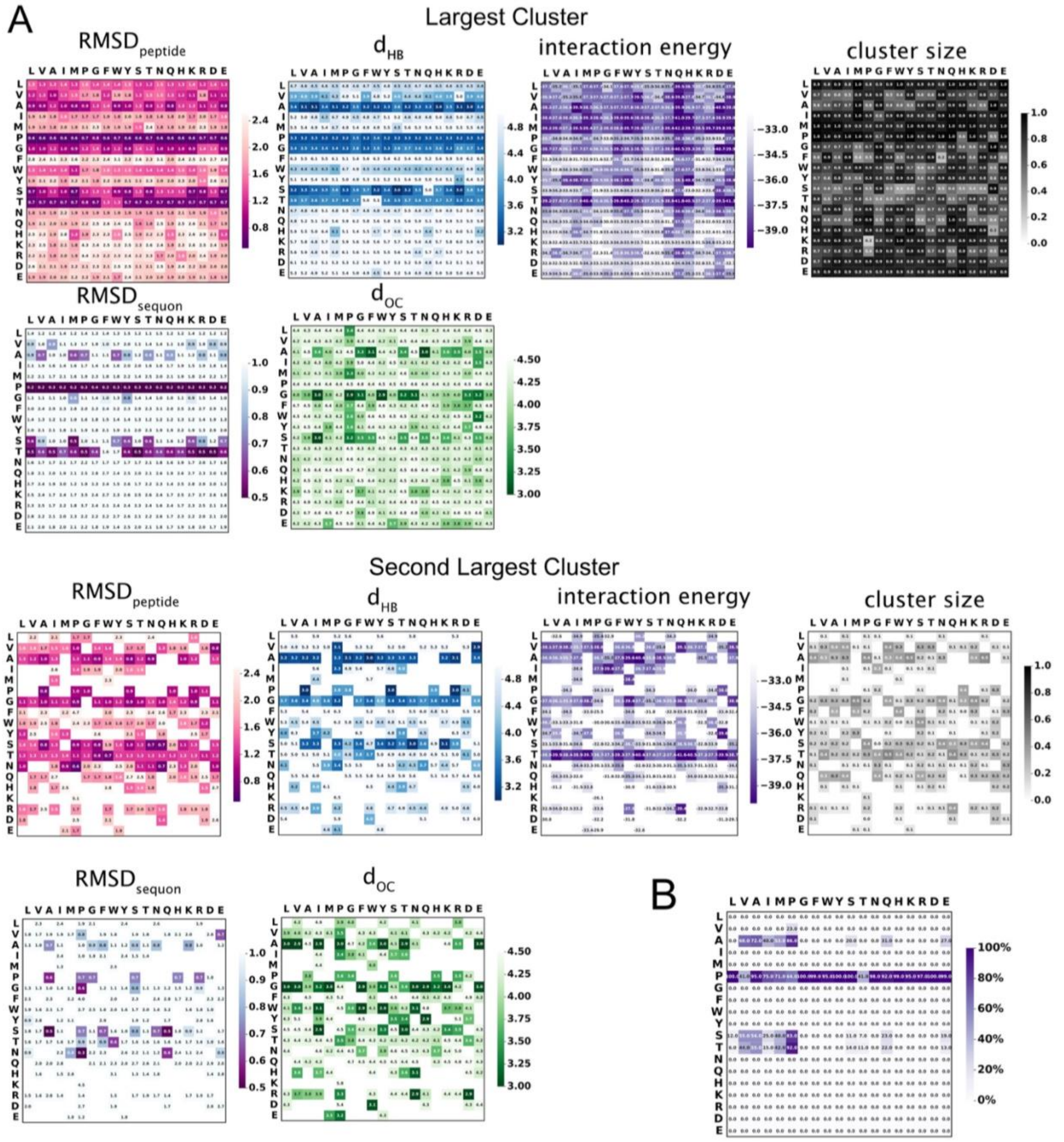


**SI Figure 2. Characterization of lowest energy representative conformation for top two clusters from Rosetta MCM sampling. Largest and second largest clusters characterized by lowest interaction energy**

**representative decoy (from left to right) $RMSD_{peptide}$, $d_{HB}$, interaction energy, normalized cluster size, $RMSD_{sequon}$ and $d_{OC}$**

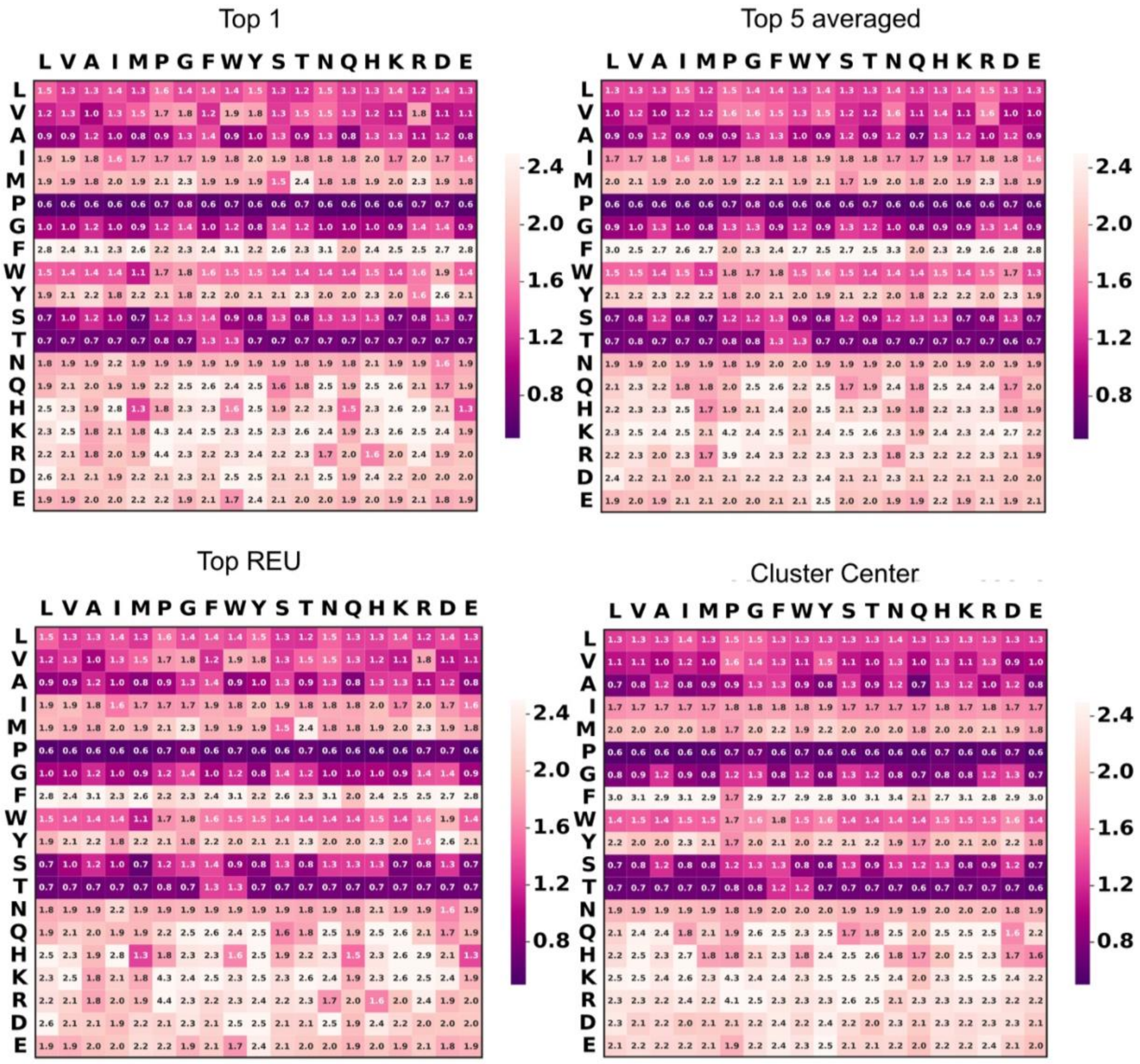


**SI Figure 3. Characterizing cluster by four different metrics - Lowest interaction energy decoy (Top 1), Average of five lowest interaction energy decoys (Top 5), Average of all decoys within 1REU of the**

**lowest interaction energy decoy (Top REU) and the decoy representing the centroid of the cluster (cluster center).**

**SI Table 1. AUC values based on the feature for the lowest interaction energy decoy for binarized experimental data. Efficiency threshold = glycosylation efficiency below which substrate is considered not glycosylated).**

| Efficiency threshold | Feature | AUC |
|---|---|---|
| 0 | interaction energy | 0.57 |
| 0.05 | interaction energy | 0.57 |
| 0.1 | interaction energy | 0.57 |
| 0.2 | interaction energy | 0.55 |
| 0.3 | interaction energy | 0.52 |
| 0.4 | interaction energy | 0.49 |
| 0.5 | interaction energy | 0.46 |
| 0.55 | interaction energy | 0.45 |
| 0 | $RMSD_{peptide}$ | 0.96 |
| 0.05 | $RMSD_{peptide}$ | 0.96 |
| 0.1 | $RMSD_{peptide}$ | 0.96 |
| 0.2 | $RMSD_{peptide}$ | 0.96 |
| 0.3 | $RMSD_{peptide}$ | 0.97 |
| 0.4 | $RMSD_{peptide}$ | 0.97 |
| 0.5 | $RMSD_{peptide}$ | 0.96 |
| 0.55 | $RMSD_{peptide}$ | 0.97 |
| 0 | $d_{HB}$ | 0.93 |
| 0.05 | $d_{HB}$ | 0.93 |
| 0.1 | $d_{HB}$ | 0.92 |
| 0.2 | $d_{HB}$ | 0.92 |
| 0.3 | $d_{HB}$ | 0.95 |
| 0.4 | $d_{HB}$ | 0.95 |
| 0.5 | $d_{HB}$ | 0.95 |
| 0.55 | $d_{HB}$ | 0.95 |
| 0 | $d_{OC}$ | 0.42 |
| 0.05 | $d_{OC}$ | 0.42 |
| 0.1 | $d_{OC}$ | 0.43 |
| 0.2 | $d_{OC}$ | 0.39 |
| 0.3 | $d_{OC}$ | 0.34 |

| | | |
|---|---|---|
| 0.4 | $d_{OC}$ | 0.30 |
| 0.5 | $d_{OC}$ | 0.30 |
| 0.55 | $d_{OC}$ | 0.24 |
| 0 | $RMSD_{sequon}$ | 0.96 |
| 0.05 | $RMSD_{sequon}$ | 0.96 |
| 0.1 | $RMSD_{sequon}$ | 0.96 |
| 0.2 | $RMSD_{sequon}$ | 0.95 |
| 0.3 | $RMSD_{sequon}$ | 0.97 |
| 0.4 | $RMSD_{sequon}$ | 0.97 |
| 0.5 | $RMSD_{sequon}$ | 0.97 |
| 0.55 | $RMSD_{sequon}$ | 0.98 |
| 0 | cluster size | 0.50 |
| 0.05 | cluster size | 0.50 |
| 0.1 | cluster size | 0.50 |
| 0.2 | cluster size | 0.55 |
| 0.3 | cluster size | 0.59 |
| 0.4 | cluster size | 0.59 |
| 0.5 | cluster size | 0.60 |
| 0.55 | cluster size | 0.64 |

**SI Table 2. Summary of AUC scores, True Positives (TP), True Negatives (TN), False Positives (FP) and False Negatives (FN), and Accuracy (ACC), Precision (Prec), Sensitivity (Sens)/True Positive Rate (TPR)/Recall and Specificity (Spec)/True Negative Rate (TNR) for predictions based on features with a range of experimental glycosylation efficiency thresholds below which a peptide is classified as non-glycosylatable. The calculation of TPs, TNs, FPs, FNs, Acc, Prec, Sens and Spec requires a threshold. Feature thresholds were chosen in two cases ($d_{HB}$ (largest cluster) < 4.0 Å and $RMSD_{peptide}$ (largest cluster) < 1.0 Å) to match criteria discussed in the main text. For all other cases, thresholds were chosen arbitrarily.**

| Feature | Experimental glycosylation efficiency threshold (%) | AUC | Feature threshold for calculation of metrics | TP | TN | Acc | Prec |
|---|---|---|---|---|---|---|---|
| | | | | **FP** | **FN** | **Sens** | **Spec** |
| $d_{HB}$ (largest cluster) | 0.10 | 0.923 | <4.0 (Å) | 44 | 259 | 0.84 | 0.44 |
| | | | | 56 | 2 | 0.96 | 0.82 |
| $RMSD_{peptide}$ (largest cluster) | 0.10 | 0.959 | <1.0 (Å) | 39 | 287 | 0.90 | 0.58 |
| | | | | 28 | 7 | 0.85 | 0.91 |
| $RMSD_{sequon}$ (largest cluster) | 0.10 | 0.955 | <0.9 (Å) | 37 | 291 | 0.91 | 0.61 |

| | | | | | | | |
|---|---|---|---|---|---|---|---|
| | | | | 24 | 9 | 0.80 | 0.92 |
| Interaction Energy (largest cluster) | 0.10 | 0.564 | < -34 (REU) | 36 | 98 | 0.37 | 0.14 |
| | | | | 217 | 10 | 0.78 | 0.31 |
| $d_{HB}$ (largest cluster) | 0.55 | 0.954 | <3.3 (Å) | 20 | 322 | 0.95 | 0.57 |
| | | | | 15 | 4 | 0.83 | 0.96 |
| $RMSD_{peptide}$ (largest cluster) | 0.55 | 0.967 | <0.7 (Å) | 16 | 328 | 0.95 | 0.64 |
| | | | | 9 | 8 | 0.67 | 0.97 |
| Fraction of decoys ($d_{HB}$ < 4.0 Å) | 0.10 | 0.908 | >0.70 (-) | 39 | 268 | 0.86 | 0.46 |
| | | | | 45 | 7 | 0.85 | 0.85 |
| Interaction Energy ($d_{HB}$ < 4.0 Å) | 0.10 | 0.875 | < -34 (REU) | 42 | 255 | 0.82 | 0.41 |
| | | | | 60 | 4 | 0.91 | 0.81 |
| Fraction of decoys ($RMSD_{peptide}$ < 1.0 Å) | 0.10 | 0.965 | >0.53 (-) | 36 | 292 | 0.91 | 0.62 |
| | | | | 23 | 10 | 0.78 | 0.93 |
| Interaction Energy ($RMSD_{peptide}$ < 1.0 Å) | 0.10 | 0.908 | < -34 (REU) | 39 | 271 | 0.86 | 0.47 |
| | | | | 44 | 7 | 0.85 | 0.86 |
| Shape Complementarity (median top 10; $RMSD_{peptide}$ < 1.0 Å) | 0.10 | 0.944 | >0.735 | 42 | 265 | 0.85 | 0.46 |
| | | | | 50 | 4 | 0.91 | 0.84 |

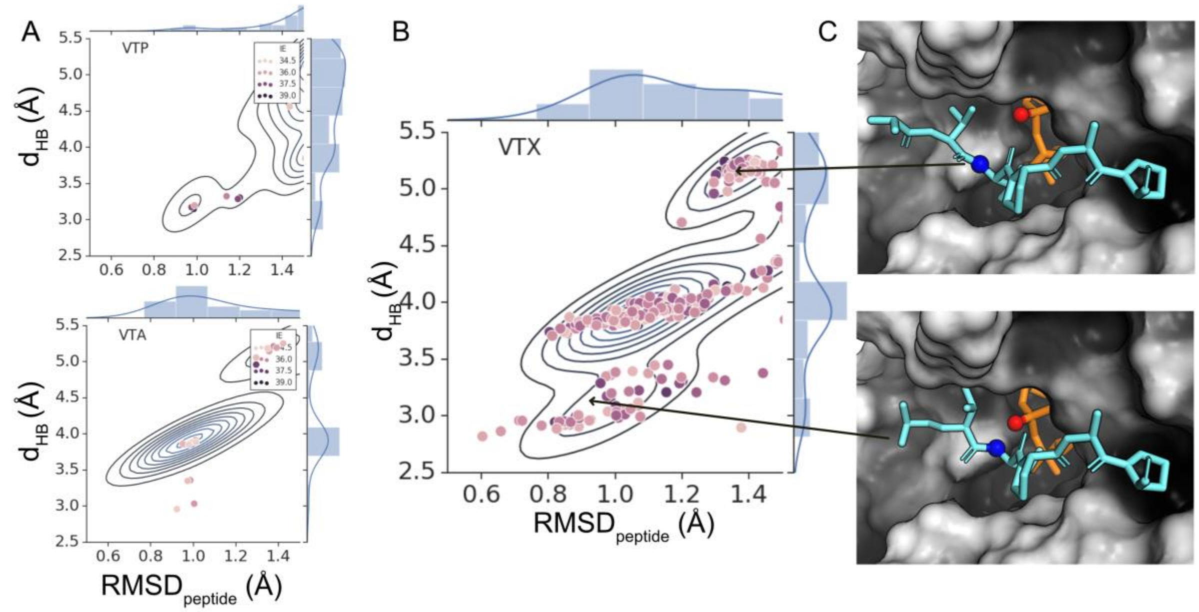


**SI Figure 4. Peptides with valine at the -1 position exhibits both conformations with hydrogen bonding compatible $d_{HB}$ and larger distances. (A) Joint probability distributions for sequons $V_{-1}TP_{+1}$ and $V_{-1}TA_{+1}$. (B) Joint probability distributions for sequons $V_{-1}TX_{+1}$ sequons exhibiting states with multiple states. (C) State with $d_{HB}$ > 4.0 Å [top panel] and state with $d_{HB}$ < 4.0 Å [bottom panel].**

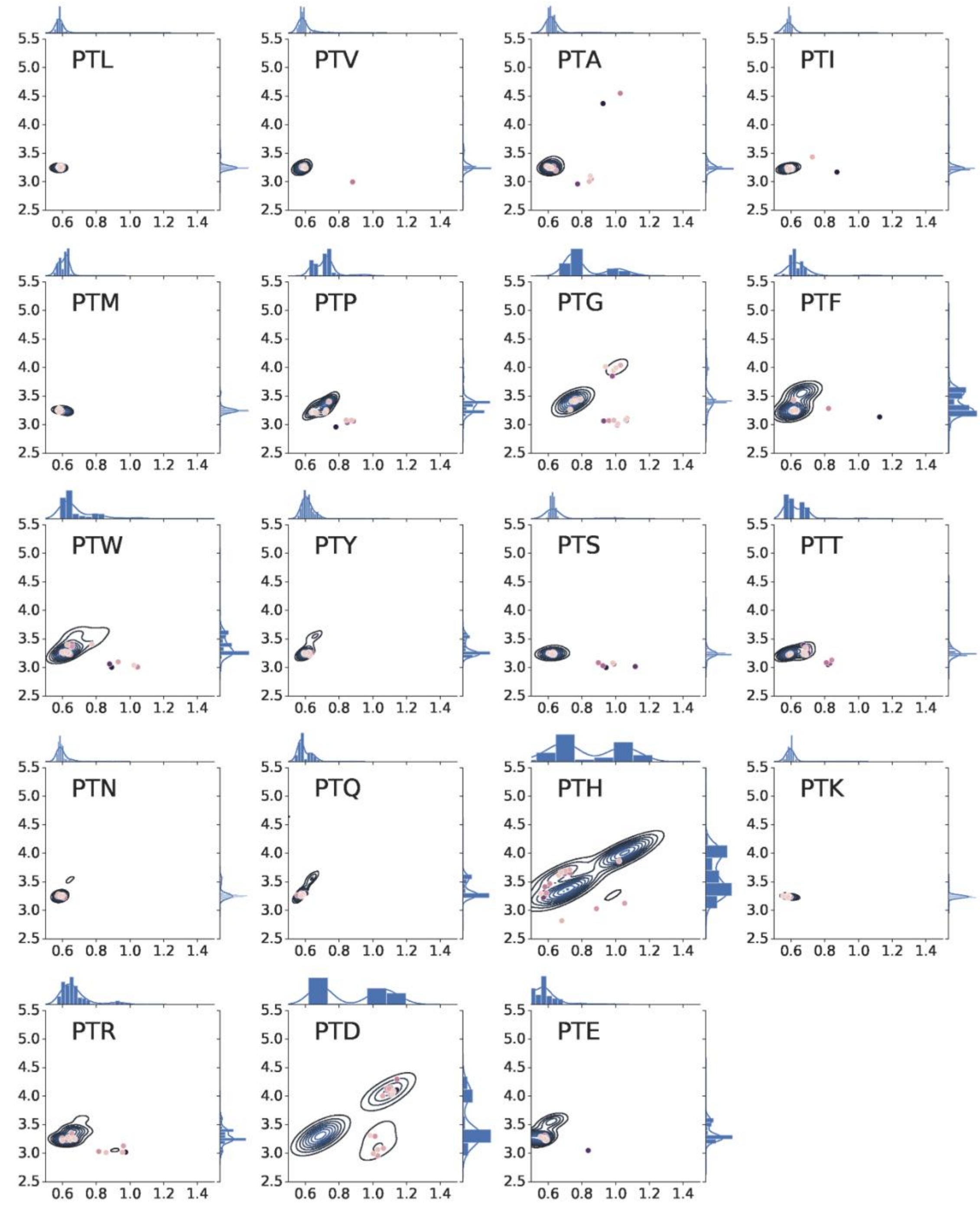


**SI Figure 5. Joint and marginal density plots in $RMSD_{peptide}$ and $d_{HB}$ for the top 1% decoys for all 19 sequons with proline at the -1 position. Top 1% (20/2000) decoys are shown as points. Darker color signifies lower interaction energy.**

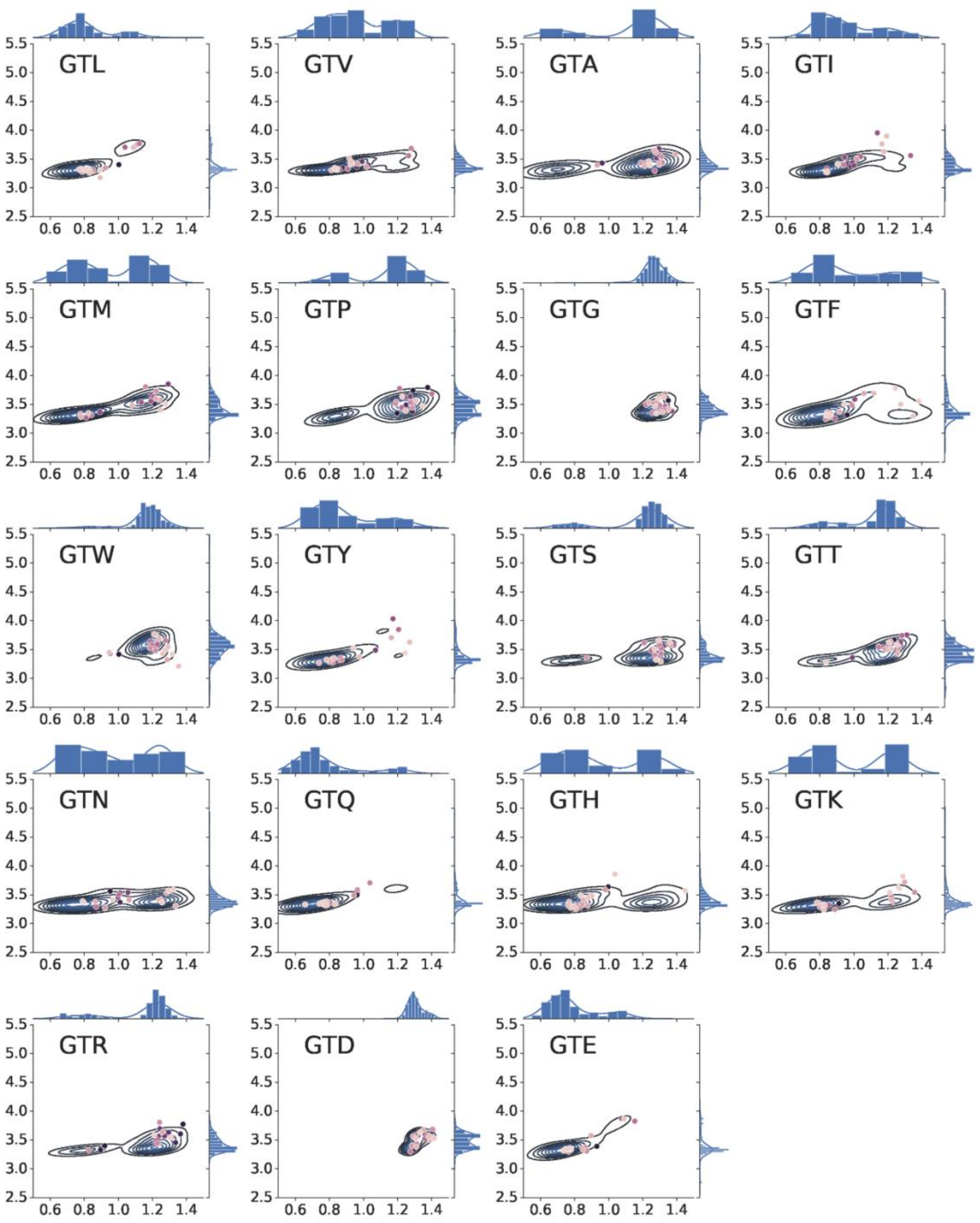


**SI Figure 6. Joint and marginal density plots in $RMSD_{peptide}$ and $d_{HB}$ for the top 1% decoys for all 19 sequons with glycine at the -1 position. Top 1% (20/2000) decoys are shown as points. Darker color signifies lower interaction energy.**

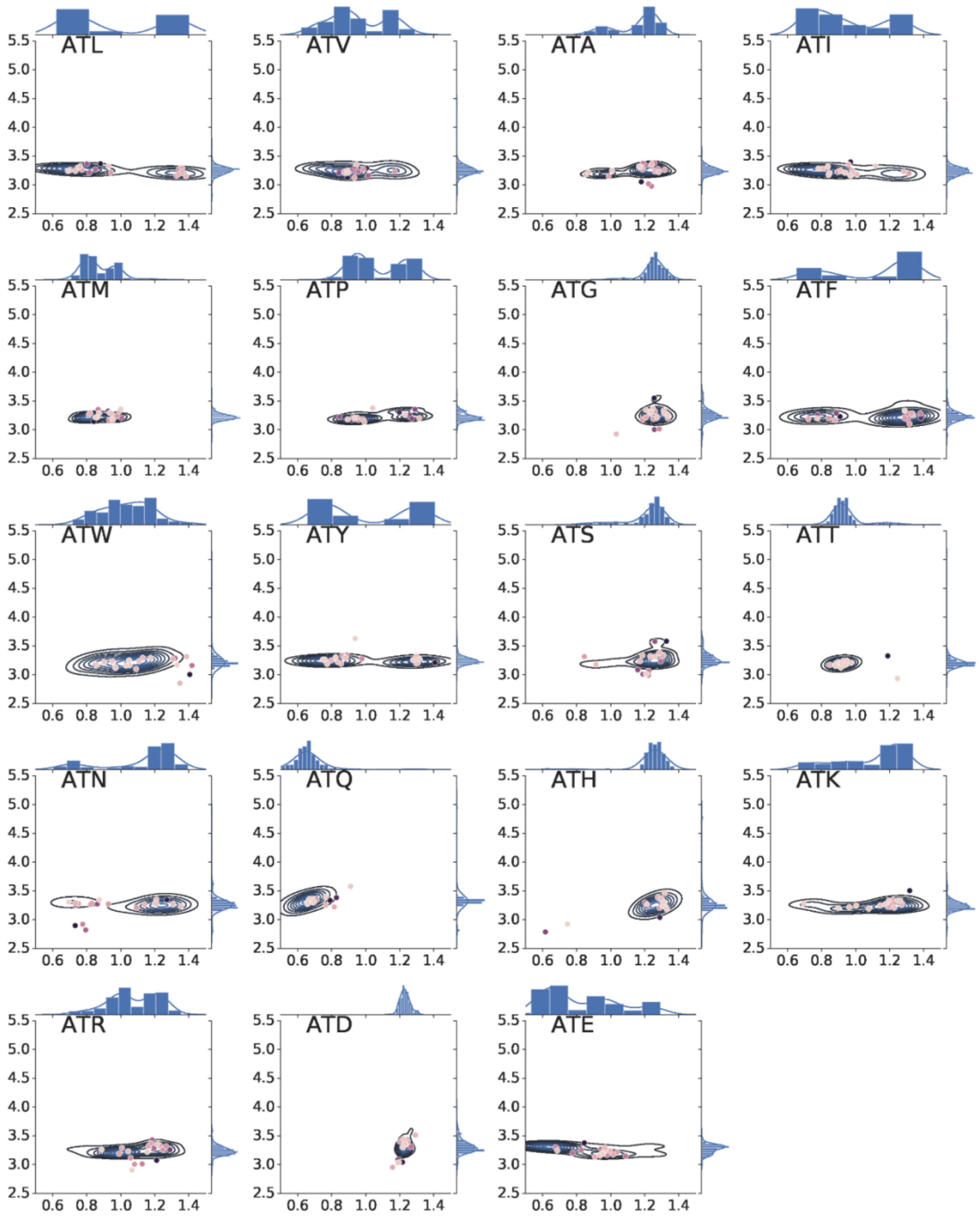


**SI Figure 7. Joint and marginal density plots in RMSD$_{peptide}$ and d$_{HB}$ for the top 1% decoys for all 19 sequons with alanine at the -1 position. Top 1% (20/2000) decoys are shown as points. Darker color signifies lower interaction energy.**

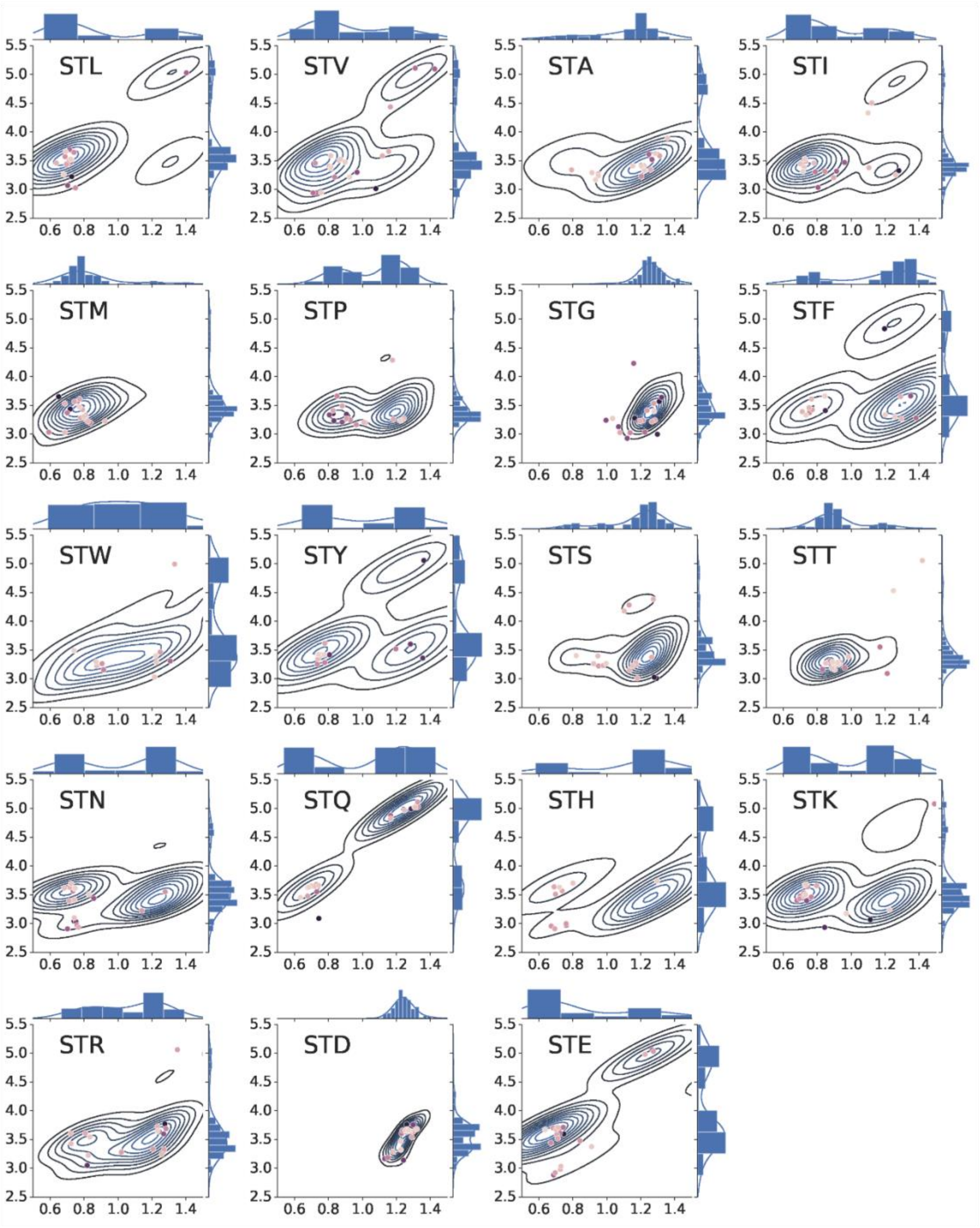


**SI Figure 8. Joint and marginal density plots in $RMSD_{peptide}$ and $d_{HB}$ for the top 1% decoys criterion for all 19 sequons with serine at the -1 position. Top 1% (20/2000) decoys are shown as points. Darker color signifies lower interaction energy.**

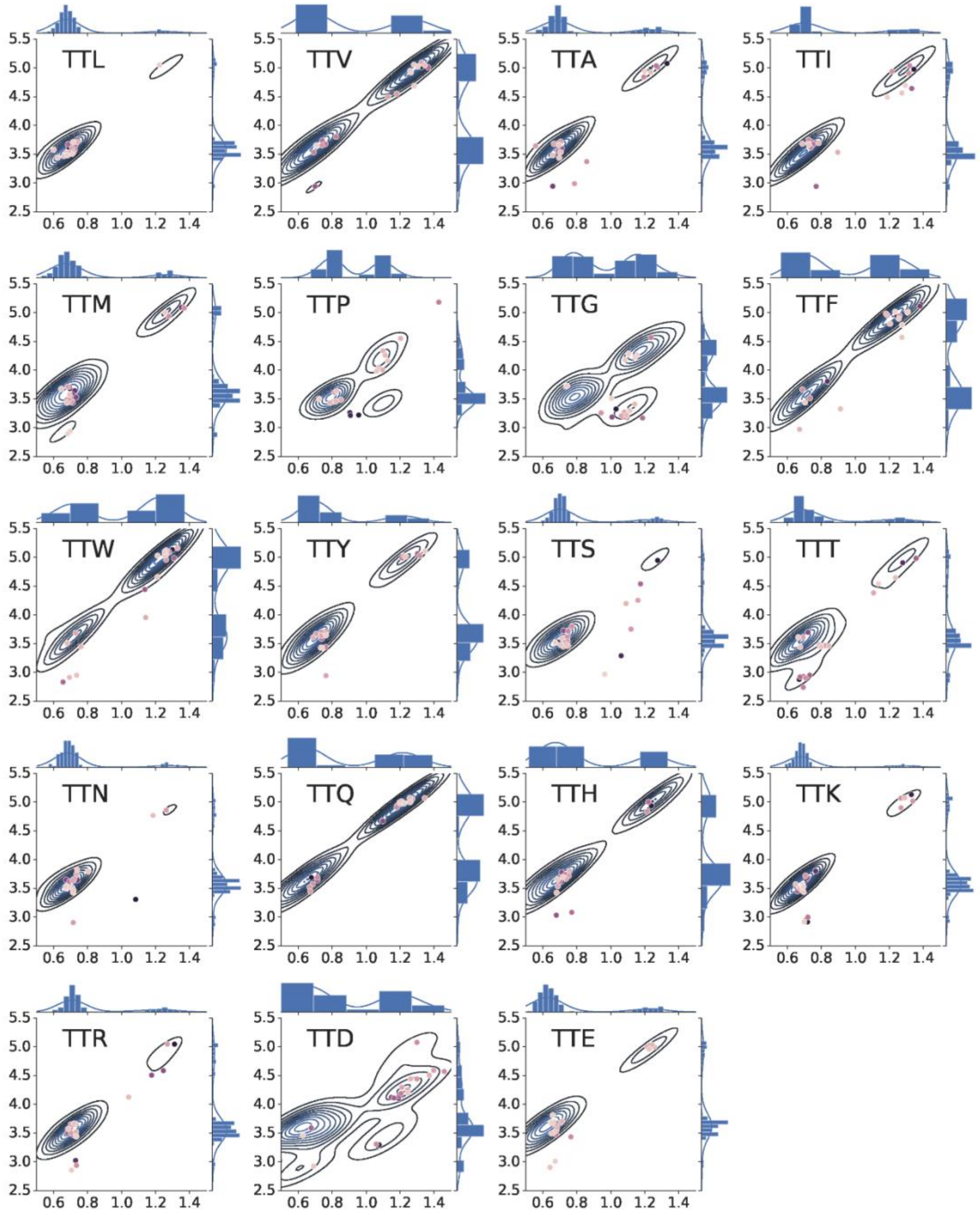


**SI Figure 9. Joint and marginal density plots in RMSD$_{peptide}$ and d$_{HB}$ for the top 1% decoys criterion for all 19 sequons with threonine at the -1 position. Top 1% (20/2000) decoys are shown as points. Darker color signifies lower interaction energy.**

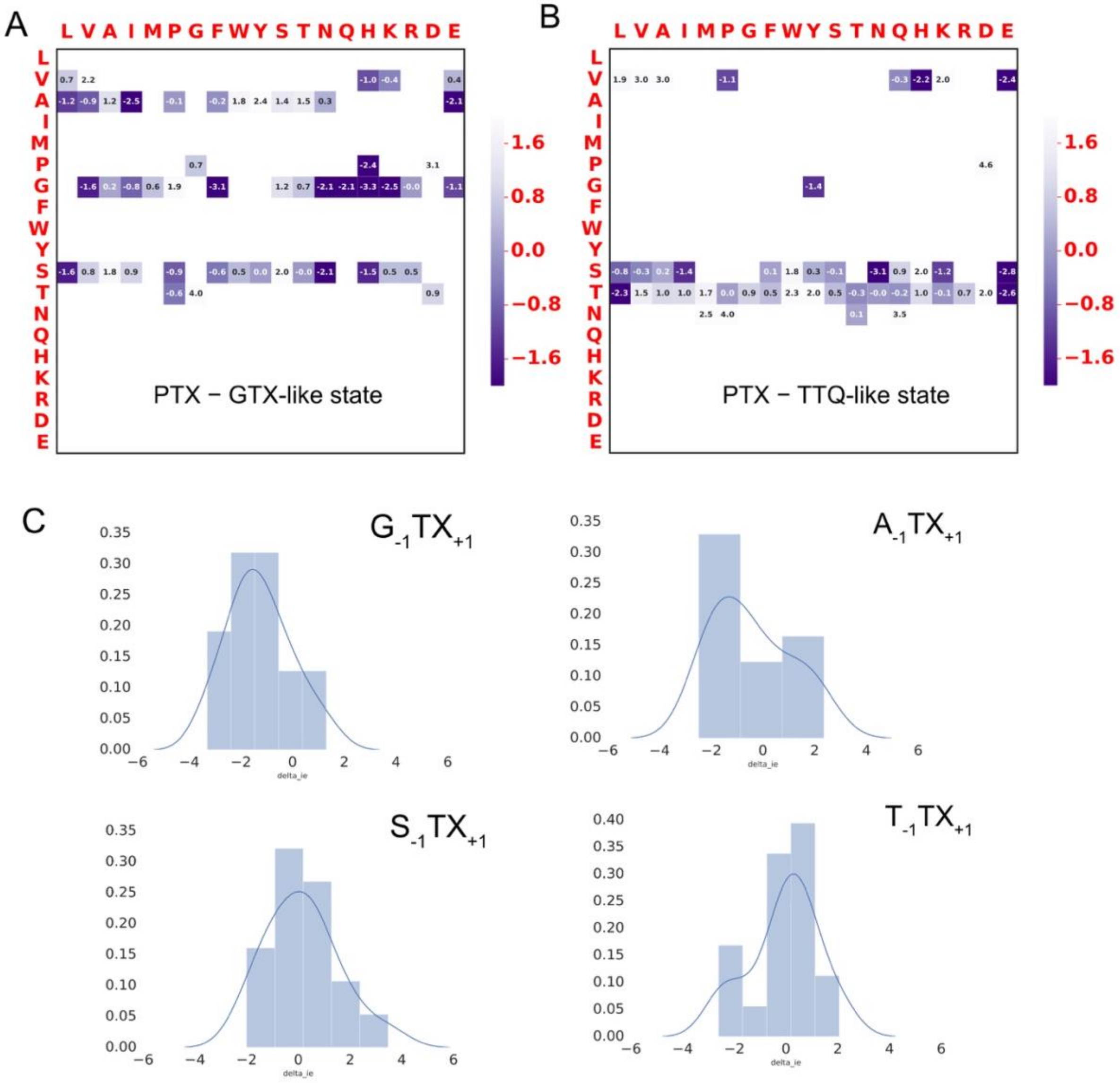


**SI Figure 10. Difference in interaction energies between competing stable states. Difference between lowest energy decoy for the PTX-like state and (A) GTX-like state, and (B) TTQ-like state for sequons that exhibit states—PTX and GTX-like and PTX and TTQ-like—respectively. (C) Distribution of**

**difference in interaction energies between the largest and second largest clusters for peptides with G, A, S, and T at the -1 position and all amino-acids (X) at the +1 position.**

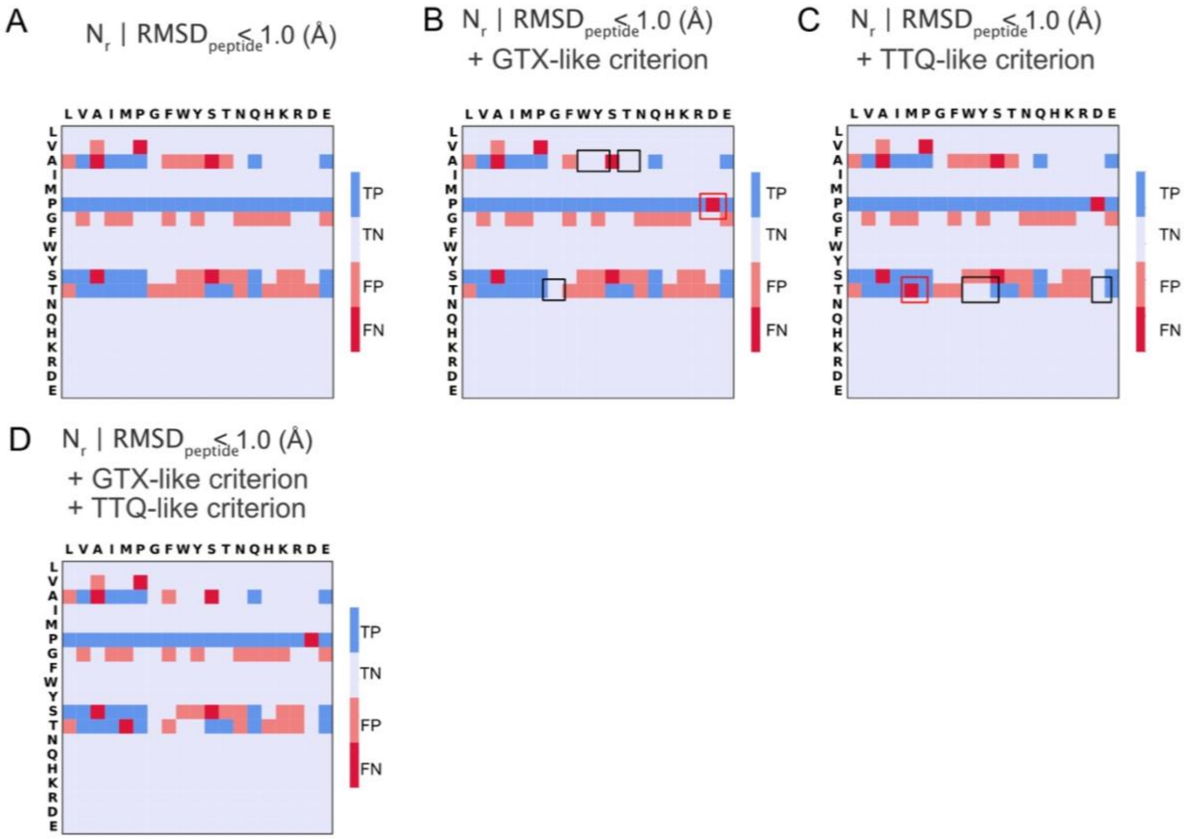


**SI Figure 11. True Positives (TP), True Negatives (TN), False Positives (FP) and False Negatives (FN) predicted by (A) $N_r$ based on the RMSDpeptide < 1.0 Å criterion at a arbitrary chosen True Positive Rate (TPR) of 0.9, (B) $N_r$ based on the RMSDpeptide < 1.0 Å criterion and GTX-like criterion, (C) $N_r$ based on the RMSDpeptide < 1.0 Å criterion and TTQ-like criterion and, (D) $N_r$ based on the RMSDpeptide < 1.0 Å criterion and GTX-like and TTQ-like criterion. Black boxes indicate the sequons for which the prediction improves over the "base" (A) criterion. Red boxes indicate the sequons for which the prediction gets worse over the "base" (A) criterion. GTX-like criterion, here, refers to counting a sequon as "non-glycosylatable" if it satisfies these two conditions: 1. Fraction of decoys that exhibit the GTX-like state is > 0.05 and, 2. The energy of lowest energy decoy for the GTX-like state is lower than the energy of lowest energy decoy for the PTX-like state. TTQ-like criterion, here, refers to counting a sequon as "non-glycosylatable" if it statisfies these two conditions: 1. Fraction of decoys that exhibit the TTQ-like state is > 0.05 and, 2. The energy of lowest energy decoy for the TTQ-like state is lower than the energy of lowest energy decoy for the PTX-like state. The threshold**

fraction of 0.05 is chosen arbitrary for demonstration. The prediction will change based on the choice of this value.

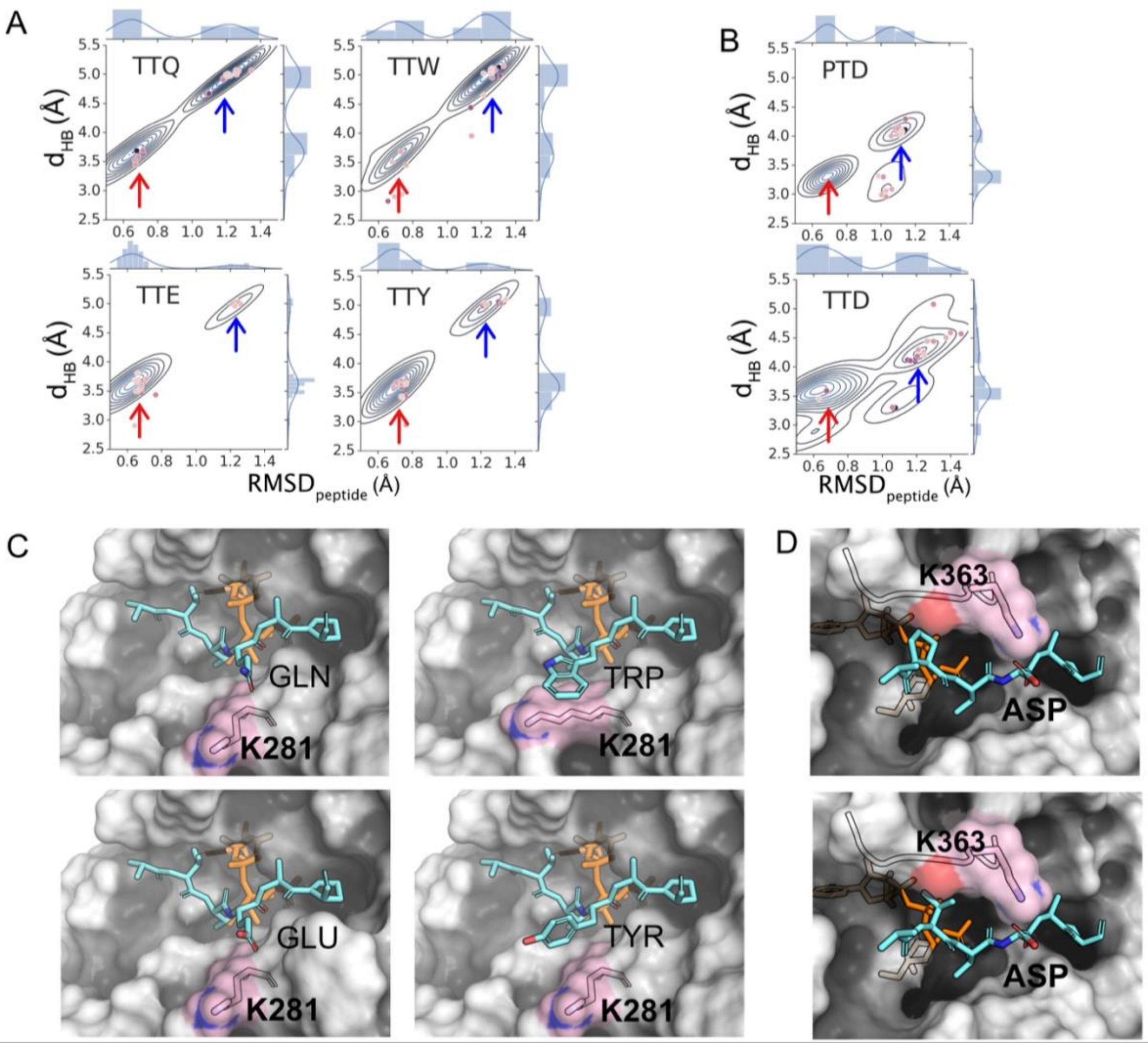


**SI Figure 12. A) Joint and marginal probability densities of Top 10% sequons for peptides with threonine at the −1 position and either glutamine, tryptophan, glutamate, or tyrosine at the +1 position. B) Peptides with proline (top) and threonine (bottom) at the −1 position and amino acids aspartic acid at the +1 position. C) TTQ-like state in which the amino acid at the +1 position interacts with K281. D) Aspartate at the +1 position interacts with K363 in the sequon PTD (top) and TTD (bottom).**

**SI Table 3. Correctness of classifications based on various criterion. The true positives, true negatives, false positives and false negatives are calculated for the $RMSD_{peptide}$ $N_r$ criterion at a True Positive Rate (TPR) of 0.90. TPR of 0.90 was arbitrarily chosen. All subsequent criteria are applied incrementally to $RMSD_{peptide}$ $N_r$ criterion as summarized in SI Figure 10.**

| Criterion | True Positives (#) | True Negatives (#) | False Positives (#) | False Negatives (#) | Balanced Accuracy (%) | FPR (%) |
|---|---|---|---|---|---|---|
| $RMSD_{peptide}$ $N_r$ | 41 | 283 | 32 | 5 | 89.5 | 10.2 |
| $RMSD_{peptide}$ $N_r$ + GTX-like criterion | 40 | 287 | 28 | 6 | 89.0 | 8.9 |
| $RMSD_{peptide}$ $N_r$ + TTQ-like criterion | 39 | 290 | 25 | 7 | 88.4 | 7.9 |
| $RMSD_{peptide}$ $N_r$ + GTX-like criterion + TTQ-like criterion | 39 | 290 | 25 | 7 | 88.4 | 7.9 |

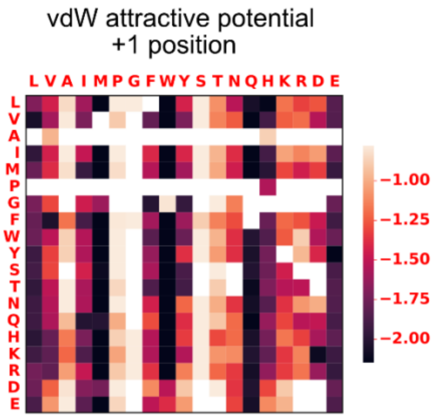


**SI Figure 13. Attractive component of the LJ potential between residue at +1 position and residue K363 on the enzyme for RMSD$_{peptide}$ > 1.0 Å. The median value for the top 20 lowest-scoring decoys is shown.**

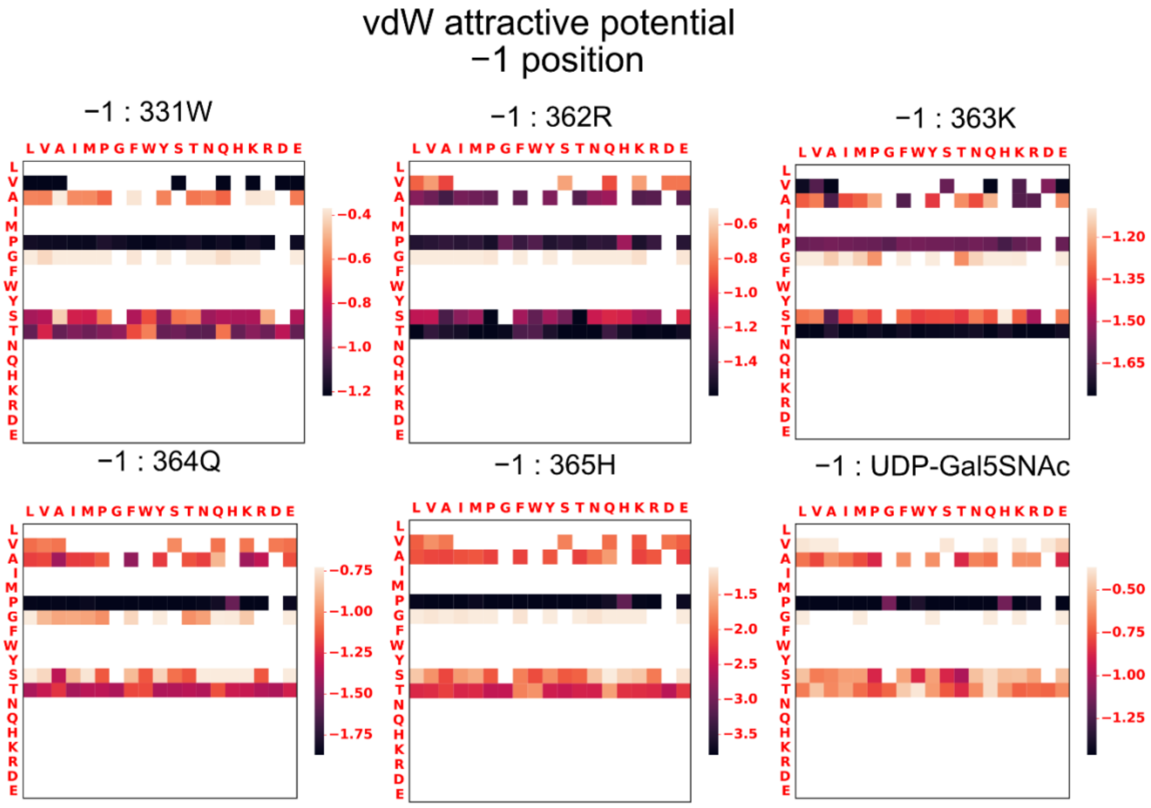


**SI Figure 14. Attractive component of the Lennard-Jones (LJ) potential in REU between residue at -1 position and residues on the enzyme for the PTX-like state (RMSD$_{peptide}$ < 1.0 Å). The median value for the top 20 lowest-scoring decoys is shown.**

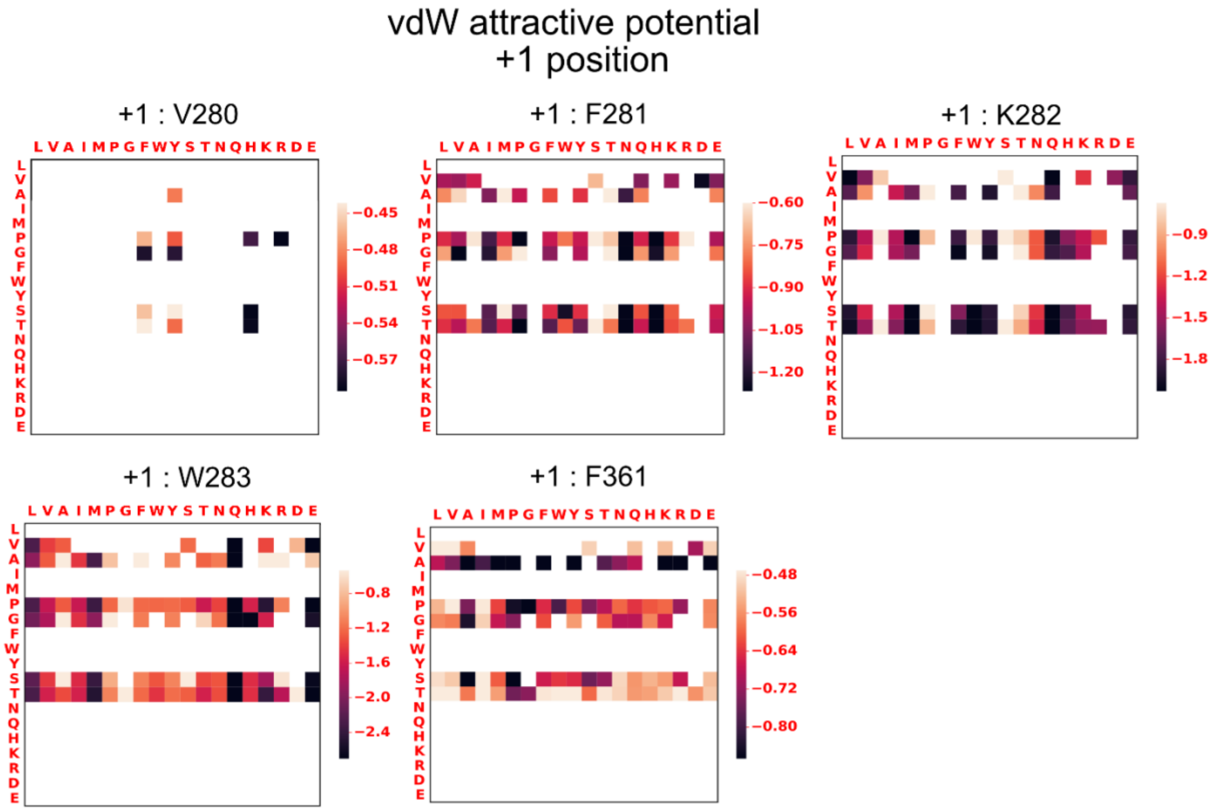


**SI Figure 15. Attractive component of the LJ potential between residue at +1 position and residues on the enzyme for the PTX-like state ($RMSD_{peptide}$ < 1.0 Å). The median value for the top 20 lowest-scoring decoys is shown.**

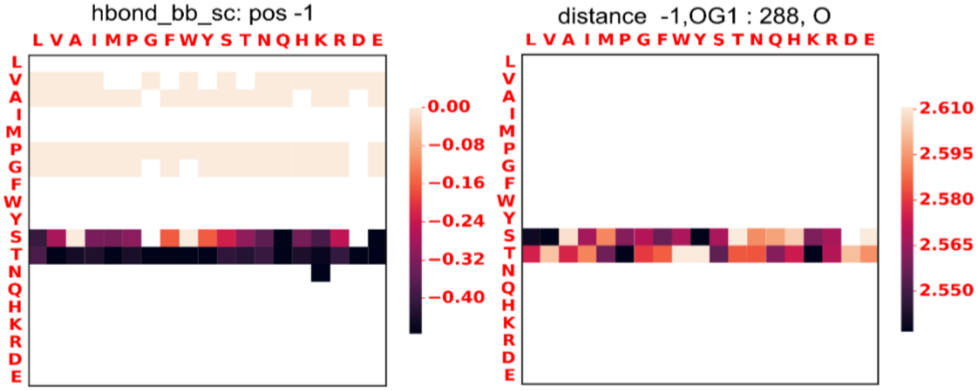


**SI Figure 16. Characterization of hydrogen bond energy (left) and distance (right) between residues Ser/Thr at -1 position (hydroxyl sidechain) and the R288 residue (backbone oxygen) on the enzyme.**

**Values shown for each sequon are the median value of the top 10 lowest interaction energies which satisfy the $RMSD_{peptide}$ < 1.0 Å criterion.**

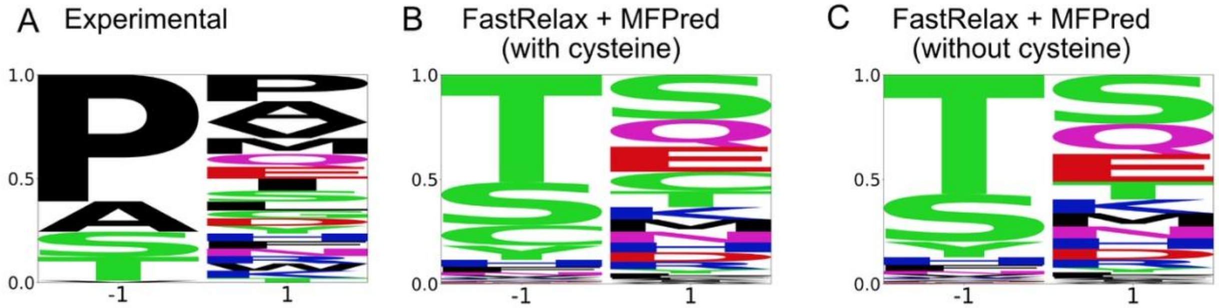


**SI Figure 137. Comparison of experimental specificity profile with specificity profiles predicted by MFPred based on ensembles of top decoy generated by Fast Relax (as outlined in reference[1]). (A) Experimental specificity profile (normalized for 19 amino acids since cysteine was not present in the experimental set). (B) Specificity profile predicted by MFPred (cysteine is included). (C) Specificity profile predicted by MFPred (cysteine is excluded to match experimental setup).**

**SI Table 4. Predictions calculated with the MFPred[1] method. Metrics Cosine Similarity, Frobenius norm, Average Absolute Distance (AAD), Jensen-Shannon Divergence (JSD) and AUC (Area under curve of the ROC curve) were calculated with scripts provided in SI info[1].**

| **Run** | **Metric** | **-1** | **+1** | **Average** |
|---|---|---|---|---|
| FastRelax+MFPred | Cosine | 0.18 | 0.62 | 0.25 |
| FastRelax+MFPred | Frobenius | 0.99 | 0.30 | 1.04 |
| FastRelax+MFPred | AAD | 0.1 | 0.05 | 0.07 |
| FastRelax+MFPred | JSD | 0.82 | 0.29 | 0.56 |
| **FastRelax+MFPred** | **AUC** | **0.68** | **0.5** | **0.59** |

1. Rubenstein, A. B., Pethe, M. A. & Khare, S. D. MFPred: Rapid and accurate prediction of protein-peptide recognition multispecificity using self-consistent mean field theory. *PLoS Comput. Biol.* **13**, e1005614 (2017).
