## Supplementary Information for "Structural basis for peptide substrate specificities of glycosyltransferase GalNAc-T2"

**1. Obtaining starting structure for each sequon**

To obtain starting structures for all sequons, run:

> python3 generate_sequons.py --mutatefile mutatelistall.txt

**2. Generating decoys for each sequon**

>mucintypeglycosylation.linuxgccrelease @flags

Where “flags” is a plain text file and contains the following options:

-in:file:s <input pdb file>

-in:file:native <input pdb file>

-ex1

-ex2aro

-nstruct 2000

-residue_to_glycosylate 3P #Threonine on peptide chain P

-substrate_type peptide

-glycosylation_low_res_refinement #enables low resolution stage

-prevent_anchor_repacking false

-tree_type docking

-sugardonor_residue 495

-nevery_interface 3

-ntotal_backbone 30

-output_distance_metrics

-enable_backbone_moves_pp #enables backbone moves in high resolution

#high resolution is enabled by default

#side chain repacking is always enabled

**3. Calculation of shape complementarity, residue-wise and pair-wise energies**

- **Pyrosetta**(1) **code snippet for shape complementarity calculation**

from pyrosetta import *

from rosetta import *

pyrosetta.init('-include_sugars -mute core')

from rosetta.core.scoring import *

from rosetta.protocols.docking import *

from rosetta.core.pose import *

from rosetta.core.simple_metrics import metrics

def get_sc_for_pose(pose,molecule_1='A',molecule_2='P'):

scC = core.scoring.sc.ShapeComplementarityCalculator()

for ires in range(1,pose.size()+1):

chain = pose.pdb_info().chain(ires)

if chain==molecule_2:

scC.AddResidue(1, pose.residue(ires))

elif chain==molecule_1:

scC.AddResidue(0, pose.residue(ires))

else:

continue

ran = scC.Calc()

results = scC.GetResults()

return results

#infile is the pdb file for a model or decoy

pose = pose_from_pdb(infile)

sc = get_sc_for_pose(pose)

dict_ = {}

dict_['sc']=sc.sc

- **Pyrosetta code snippet for pairwise energy calculation**

from pyrosetta import *

from rosetta import *

pyrosetta.init('-include_sugars','-mute core -mute basic.io.database')

from rosetta.core.scoring import *

from rosetta.protocols.docking import *

sf = ScoreFunction()

sf = get_fa_scorefxn()

def PairwiseEnergyMapForPdbFile(curfile,residues=[498,500]):

#498 is -1 position in the pdb file

#500 is +1 position in the pdb file

curpose = pose_from_pdb(curfile)

sf_pose = sf(curpose)

dict_ = {}

columns=['residue_i','residue_j','score_type','score_value']

for key in columns:

dict_[key] = []

for resi in residues:

score_types = sf.get_nonzero_weighted_scoretypes()

mylist = ['rama','fa_rep','fa_atr','hbond_bb_sc','hbond_sc','fa_sol','fa_intra_rep']

for st in score_types:

strst = str(st)

strst_clean = strst.split('.')[1]

if strst_clean in mylist:

score_tuple = pyrosetta.toolbox.atom_pair_energy._reisude_pair_energies(resi,curpose,sf,st,0.25)

for entry in score_tuple:

dict_['score_type'].append(strst_clean)

dict_['score_value'].append(entry[1])

dict_['residue_j'].append(entry[0])

dict_['residue_i'].append(resi)

return dict_

#curfile is the pdb file for a model or decoy

#498-X-1, 500 is residue X0, 501 is residue X+1

dict_energies = PairwiseEnergyMapForPdbFile(curfile)

- **Pyrosetta code snippet for per-residue energy calculation**

from pyrosetta import *

from rosetta import *

pyrosetta.init('-include_sugars','-mute core -mute basic.io.database')

from rosetta.core.scoring import *

from rosetta.protocols.docking import *

sf = ScoreFunction()

sf = get_fa_scorefxn()

def EnergyMapForPdbFile(curfile,residues=[498,499,500],columns=['residue', 'score_type','score_value']):

#498 is -1 position in the pdb file

#499 is 0 position in the pdb file

#500 is +1 position in the pdb file

columns_total = ['tag','file','score_type','score_value']

dict_ ={}

dict_total = {}

for key in columns_total:

dict_total[key]=[]

for key in columns:

dict_[key]=[]

curpose = pose_from_pdb(curfile)

sf_pose = sf(curpose)

pose_energies = curpose.energies()

total_e = pose_energies.active_total_energies()

score_types = sf.get_nonzero_weighted_scoretypes()

mylist = ['rama','fa_rep','fa_atr','hbond_bb_sc','hbond_sc','fa_sol','fa_intra_rep']

for st in score_types:

for resi in residues:

strst = str(st)

strst_clean = strst.split('.')[1]

if strst_clean in mylist:

value = pose_energies.residue_total_energies(resi)[st]

dict_['score_value'].append(value)

dict_['score_type'].append(strst_clean)

dict_['residue'].append(resi)

return dict_

#curfile is the pdb file for a model or decoy

dict_energies = EnergyMapForPdbFile(curfile)

**4. Software versions and requirements for pyrosetta scripts**

Pyrosetta: Pyrosetta version: PyRosetta-4 2019.

**5. Availability of post-processing scripts and data**

For additional scripts for data processing, see repository on github <https://github.com/heiidii/galnt_paper_repository> - currently private. All data and scripts are available upon request. Git repository will be made public upon publication of work.

All files are available in Supplementary Materials.

1.	Chaudhury S, Lyskov S, Gray JJ. PyRosetta: a script-based interface for implementing molecular modeling algorithms using Rosetta. Bioinformatics [Internet]. 2010 Mar 1 [cited 2018 Apr 12];26(5):689–91. Available from: http://www.ncbi.nlm.nih.gov/pubmed/20061306
