## Additional code and pdb files for "Structural basis for peptide substrate specificities of glycosyltransferase GalNAc-T2": generate_sequons.py

```
from pyrosetta import *
from rosetta import *
pyrosetta.init('-include_sugars')


aa_list2 = ['PRO','LEU',"VAL","ALA","ILE","MET","GLY","PHE","TRP","TYR","SER","THR","ASN","GLN","HIS","LYS","ARG","ASP","GLU"]
aa_list1 = aa_list2
print("X-1: ",aa_list1,"\nLen:",len(aa_list1))
print("X+1: ",aa_list2,"\nLen:",len(aa_list2))
import argparse
from rosetta.core.pose import *
from rosetta.core.scoring import *
import rosetta.protocols.simple_moves as psm
from rosetta.protocols.docking import *
from rosetta.core.kinematics import MoveMap
from rosetta.core.kinematics import FoldTree
from rosetta.core.pack.task import TaskFactory
from rosetta.core.pack.task import operation
from rosetta.core.select import residue_selector as selections
from rosetta.core.select.movemap import *
from rosetta.protocols import minimization_packing as pack_min


sf = ScoreFunction()
sf = get_fa_scorefxn()

def PackAndMinimize(pose,residues):
  
  residueselector = selections.ResidueIndexSelector(residues) 
  tf = TaskFactory()
  tf.push_back(operation.RestrictToRepacking())
  tf.push_back( operation.OperateOnResidueSubset( operation.PreventRepackingRLT() , residueselector , True ))
  packer = pack_min.PackRotamersMover() # uses pack_rotamers_run that uses packing with simualted annealing
  packer.task_factory(tf)
  packer.apply(pose)
  packer_pose = pose.clone()
  return packer_pose

def DumpScoredFile(pose1,name="inputpdb_score.pdb"):
  sf = ScoreFunction()
  sf = get_fa_scorefxn()
  print( "Dumping file")
  pose1.dump_scored_pdb(name,sf)

def MutateResidue(pose_mod,target_index,new_res):
  print( "Mutating residue with index ",target_index," to ",new_res)
  mutate_mover = psm.MutateResidue(target_index,new_res)
  mutate_mover.apply(pose_mod)
  return pose_mod


if __name__ ==  "__main__":
  parser = argparse.ArgumentParser()
  parser.add_argument('--mutatefile',default='mutatelist.txt', help='mutation specs. Single aa fine: target_index new_res inputpdb. All aa: inputpdbfile target_indices separated by spaces')
  parser.add_argument('--partners',default='AB_P', help='partners for docking foldtree')

  args = parser.parse_args()
  partners = args.partners
  pose_ref = Pose()
  if args.mutatefile:
    if os.path.exists(args.mutatefile):
      mf = open(args.mutatefile,'r')
      for line in mf:
        if line.find('#') !=-1: continue
        if len(line.split())<2:
          print( "please specify atleast 2 specs per file eg. inputpdbfile target_index_first {target_index_second...}")
          sys.exit()
        dat = line.split()
        inputpdbfile = dat[0]
        if os.path.exists(inputpdbfile):
          pose_mut = Pose()
          pose_mut = pose_from_pdb(inputpdbfile)
          cleanpose = pose_mut.clone()

          if len(dat[1:])==2:
            target_index = int(dat[1])

            for new_res in aa_list1:
                pose_mut = cleanpose.clone() #resetting to clean pose
                pose_new = MutateResidue(pose_mut,target_index,new_res)
                cleanpose_mut1 = pose_new.clone()

                for new_res_2 in aa_list2:
                    target_index_2 = int(dat[2])
                    pose_new = cleanpose_mut1.clone()
                    pose_new_2 = MutateResidue(pose_new,target_index_2,new_res_2)
                    setup_foldtree(pose_new_2,partners, Vector1([1]))
                    resvec = pyrosetta.rosetta.utility.vector1_unsigned_long()
                    resvec.append(target_index)
                    resvec.append(target_index_2)
                    pack_pose = PackAndMinimize(pose_new,resvec)
                    pdbfile = inputpdbfile.split('.pdb')[0] + "_%s%s_%s%s.pdb" %(dat[1],new_res,dat[2],new_res_2)
                    DumpScoredFile(pack_pose,pdbfile)
```
