## Additional code and pdb files for "Structural basis for peptide substrate specificities of glycosyltransferase GalNAc-T2": StartingStructure.py

```
ATOM      1  N   LYS A   1      -1.474  46.044 -51.944  1.00  0.00           N  
ATOM      2  CA  LYS A   1      -1.864  46.016 -50.504  1.00  0.00           C  
ATOM      3  C   LYS A   1      -1.242  47.188 -49.757  1.00  0.00           C  
ATOM      4  O   LYS A   1      -0.262  47.778 -50.209  1.00  0.00           O  
ATOM      5  CB  LYS A   1      -1.445  44.702 -49.843  1.00  0.00           C  
ATOM      6  CG  LYS A   1      -2.021  43.455 -50.502  1.00  0.00           C  
ATOM      7  CD  LYS A   1      -3.538  43.423 -50.392  1.00  0.00           C  
ATOM      8  CE  LYS A   1      -4.106  42.125 -50.946  1.00  0.00           C  
ATOM      9  NZ  LYS A   1      -5.594  42.103 -50.897  1.00  0.00           N1+
ATOM     10 1H   LYS A   1      -1.891  45.268 -52.418  1.00  0.00           H  
ATOM     11 2H   LYS A   1      -1.792  46.896 -52.359  1.00  0.00           H  
ATOM     12 3H   LYS A   1      -0.479  45.986 -52.022  1.00  0.00           H  
ATOM     13  HA  LYS A   1      -2.951  46.072 -50.438  1.00  0.00           H  
ATOM     14 1HB  LYS A   1      -0.358  44.617 -49.859  1.00  0.00           H  
ATOM     15 2HB  LYS A   1      -1.758  44.703 -48.799  1.00  0.00           H  
ATOM     16 1HG  LYS A   1      -1.741  43.438 -51.556  1.00  0.00           H  
ATOM     17 2HG  LYS A   1      -1.611  42.567 -50.021  1.00  0.00           H  
ATOM     18 1HD  LYS A   1      -3.829  43.520 -49.345  1.00  0.00           H  
ATOM     19 2HD  LYS A   1      -3.961  44.260 -50.947  1.00  0.00           H  
ATOM     20 1HE  LYS A   1      -3.787  42.000 -51.980  1.00  0.00           H  
ATOM     21 2HE  LYS A   1      -3.725  41.284 -50.367  1.00  0.00           H  
ATOM     22 1HZ  LYS A   1      -5.930  41.227 -51.273  1.00  0.00           H  
ATOM     23 2HZ  LYS A   1      -5.901  42.199 -49.939  1.00  0.00           H  
ATOM     24 3HZ  LYS A   1      -5.960  42.867 -51.446  1.00  0.00           H  
ATOM     25  N   VAL A   2      -1.818  47.529 -48.610  1.00  0.00           N  
ATOM     26  CA  VAL A   2      -1.289  48.625 -47.811  1.00  0.00           C  
ATOM     27  C   VAL A   2      -0.737  48.102 -46.494  1.00  0.00           C  
ATOM     28  O   VAL A   2      -1.308  47.196 -45.888  1.00  0.00           O  
ATOM     29  CB  VAL A   2      -2.375  49.684 -47.499  1.00  0.00           C  
ATOM     30  CG1 VAL A   2      -2.895  50.298 -48.792  1.00  0.00           C  
ATOM     31  CG2 VAL A   2      -3.510  49.050 -46.720  1.00  0.00           C  
ATOM     32  H   VAL A   2      -2.631  47.028 -48.281  1.00  0.00           H  
ATOM     33  HA  VAL A   2      -0.494  49.114 -48.377  1.00  0.00           H  
ATOM     34  HB  VAL A   2      -1.930  50.483 -46.907  1.00  0.00           H  
ATOM     35 1HG1 VAL A   2      -3.658  51.042 -48.560  1.00  0.00           H  
ATOM     36 2HG1 VAL A   2      -2.072  50.777 -49.323  1.00  0.00           H  
ATOM     37 3HG1 VAL A   2      -3.328  49.518 -49.417  1.00  0.00           H  
ATOM     38 1HG2 VAL A   2      -4.268  49.802 -46.505  1.00  0.00           H  
ATOM     39 2HG2 VAL A   2      -3.953  48.247 -47.310  1.00  0.00           H  
ATOM     40 3HG2 VAL A   2      -3.126  48.644 -45.784  1.00  0.00           H  
ATOM     41  N   ARG A   3       0.392  48.654 -46.067  1.00  0.00           N  
ATOM     42  CA  ARG A   3       0.978  48.258 -44.796  1.00  0.00           C  
ATOM     43  C   ARG A   3       0.013  48.828 -43.765  1.00  0.00           C  
ATOM     44  O   ARG A   3      -0.519  49.920 -43.959  1.00  0.00           O  
ATOM     45  CB  ARG A   3       2.383  48.809 -44.601  1.00  0.00           C  
ATOM     46  CG  ARG A   3       3.407  48.323 -45.615  1.00  0.00           C  
ATOM     47  CD  ARG A   3       3.767  46.899 -45.392  1.00  0.00           C  
ATOM     48  NE  ARG A   3       4.814  46.457 -46.299  1.00  0.00           N  
ATOM     49  CZ  ARG A   3       5.323  45.209 -46.329  1.00  0.00           C  
ATOM     50  NH1 ARG A   3       4.872  44.295 -45.500  1.00  0.00           N1+
ATOM     51  NH2 ARG A   3       6.276  44.905 -47.193  1.00  0.00           N  
ATOM     52  H   ARG A   3       0.855  49.358 -46.625  1.00  0.00           H  
ATOM     53  HA  ARG A   3       1.066  47.171 -44.779  1.00  0.00           H  
ATOM     54 1HB  ARG A   3       2.357  49.896 -44.654  1.00  0.00           H  
ATOM     55 2HB  ARG A   3       2.746  48.538 -43.610  1.00  0.00           H  
ATOM     56 1HG  ARG A   3       2.997  48.420 -46.621  1.00  0.00           H  
ATOM     57 2HG  ARG A   3       4.314  48.923 -45.533  1.00  0.00           H  
ATOM     58 1HD  ARG A   3       4.123  46.770 -44.371  1.00  0.00           H  
ATOM     59 2HD  ARG A   3       2.890  46.273 -45.553  1.00  0.00           H  
ATOM     60  HE  ARG A   3       5.187  47.133 -46.952  1.00  0.00           H  
ATOM     61 1HH1 ARG A   3       4.143  44.527 -44.839  1.00  0.00           H  
ATOM     62 2HH1 ARG A   3       5.253  43.360 -45.522  1.00  0.00           H  
ATOM     63 1HH2 ARG A   3       6.623  45.609 -47.831  1.00  0.00           H  
ATOM     64 2HH2 ARG A   3       6.657  43.971 -47.216  1.00  0.00           H  
ATOM     65  N   TRP A   4      -0.226  48.105 -42.677  1.00  0.00           N  
ATOM     66  CA  TRP A   4      -1.190  48.586 -41.694  1.00  0.00           C  
ATOM     67  C   TRP A   4      -1.019  50.021 -41.188  1.00  0.00           C  
ATOM     68  O   TRP A   4      -2.011  50.703 -40.954  1.00  0.00           O  
ATOM     69  CB  TRP A   4      -1.296  47.620 -40.502  1.00  0.00           C  
ATOM     70  CG  TRP A   4      -0.107  47.575 -39.604  1.00  0.00           C  
ATOM     71  CD1 TRP A   4       0.926  46.687 -39.652  1.00  0.00           C  
ATOM     72  CD2 TRP A   4       0.180  48.468 -38.522  1.00  0.00           C  
ATOM     73  CE2 TRP A   4       1.407  48.060 -37.959  1.00  0.00           C  
ATOM     74  CE3 TRP A   4      -0.480  49.577 -37.976  1.00  0.00           C  
ATOM     75  NE1 TRP A   4       1.839  46.969 -38.667  1.00  0.00           N  
ATOM     76  CZ2 TRP A   4       1.992  48.722 -36.873  1.00  0.00           C  
ATOM     77  CZ3 TRP A   4       0.101  50.238 -36.897  1.00  0.00           C  
ATOM     78  CH2 TRP A   4       1.326  49.804 -36.358  1.00  0.00           C  
ATOM     79  H   TRP A   4       0.245  47.227 -42.513  1.00  0.00           H  
ATOM     80  HA  TRP A   4      -2.167  48.651 -42.172  1.00  0.00           H  
ATOM     81 1HB  TRP A   4      -2.155  47.892 -39.888  1.00  0.00           H  
ATOM     82 2HB  TRP A   4      -1.464  46.607 -40.868  1.00  0.00           H  
ATOM     83  HD1 TRP A   4       1.013  45.871 -40.368  1.00  0.00           H  
ATOM     84  HE1 TRP A   4       2.692  46.456 -38.491  1.00  0.00           H  
ATOM     85  HE3 TRP A   4      -1.428  49.917 -38.391  1.00  0.00           H  
ATOM     86  HZ2 TRP A   4       2.941  48.402 -36.442  1.00  0.00           H  
ATOM     87  HZ3 TRP A   4      -0.418  51.100 -36.478  1.00  0.00           H  
ATOM     88  HH2 TRP A   4       1.754  50.342 -35.512  1.00  0.00           H  
ATOM     89  N   PRO A   5       0.227  50.507 -41.026  1.00  0.00           N  
ATOM     90  CA  PRO A   5       0.377  51.883 -40.537  1.00  0.00           C  
ATOM     91  C   PRO A   5      -0.254  52.915 -41.466  1.00  0.00           C  
ATOM     92  O   PRO A   5      -0.599  54.022 -41.041  1.00  0.00           O  
ATOM     93  CB  PRO A   5       1.891  52.058 -40.446  1.00  0.00           C  
ATOM     94  CG  PRO A   5       2.388  50.665 -40.224  1.00  0.00           C  
ATOM     95  CD  PRO A   5       1.546  49.872 -41.187  1.00  0.00           C  
ATOM     96  HA  PRO A   5      -0.088  51.966 -39.544  1.00  0.00           H  
ATOM     97 1HB  PRO A   5       2.274  52.512 -41.371  1.00  0.00           H  
ATOM     98 2HB  PRO A   5       2.141  52.743 -39.623  1.00  0.00           H  
ATOM     99 1HG  PRO A   5       3.467  50.610 -40.430  1.00  0.00           H  
ATOM    100 2HG  PRO A   5       2.250  50.375 -39.173  1.00  0.00           H  
ATOM    101 1HD  PRO A   5       1.944  49.989 -42.206  1.00  0.00           H  
ATOM    102 2HD  PRO A   5       1.548  48.813 -40.890  1.00  0.00           H  
ATOM    103  N   ASP A   6      -0.397  52.548 -42.735  1.00  0.00           N  
ATOM    104  CA  ASP A   6      -0.972  53.447 -43.727  1.00  0.00           C  
ATOM    105  C   ASP A   6      -2.469  53.250 -43.954  1.00  0.00           C  
ATOM    106  O   ASP A   6      -3.074  53.944 -44.770  1.00  0.00           O  
ATOM    107  CB  ASP A   6      -0.238  53.278 -45.060  1.00  0.00           C  
ATOM    108  CG  ASP A   6       1.213  53.738 -45.000  1.00  0.00           C  
ATOM    109  OD1 ASP A   6       1.466  54.765 -44.416  1.00  0.00           O  
ATOM    110  OD2 ASP A   6       2.053  53.057 -45.538  1.00  0.00           O1-
ATOM    111  H   ASP A   6      -0.102  51.626 -43.019  1.00  0.00           H  
ATOM    112  HA  ASP A   6      -0.865  54.471 -43.369  1.00  0.00           H  
ATOM    113 1HB  ASP A   6      -0.260  52.230 -45.357  1.00  0.00           H  
ATOM    114 2HB  ASP A   6      -0.753  53.849 -45.833  1.00  0.00           H  
ATOM    115  N   PHE A   7      -3.075  52.308 -43.236  1.00  0.00           N  
ATOM    116  CA  PHE A   7      -4.505  52.076 -43.396  1.00  0.00           C  
ATOM    117  C   PHE A   7      -5.268  53.283 -42.852  1.00  0.00           C  
ATOM    118  O   PHE A   7      -4.955  53.791 -41.775  1.00  0.00           O  
ATOM    119  CB  PHE A   7      -4.938  50.802 -42.668  1.00  0.00           C  
ATOM    120  CG  PHE A   7      -6.391  50.464 -42.851  1.00  0.00           C  
ATOM    121  CD1 PHE A   7      -6.844  49.903 -44.035  1.00  0.00           C  
ATOM    122  CD2 PHE A   7      -7.307  50.708 -41.839  1.00  0.00           C  
ATOM    123  CE1 PHE A   7      -8.180  49.592 -44.204  1.00  0.00           C  
ATOM    124  CE2 PHE A   7      -8.643  50.398 -42.004  1.00  0.00           C  
ATOM    125  CZ  PHE A   7      -9.079  49.839 -43.189  1.00  0.00           C  
ATOM    126  H   PHE A   7      -2.556  51.746 -42.576  1.00  0.00           H  
ATOM    127  HA  PHE A   7      -4.718  51.952 -44.459  1.00  0.00           H  
ATOM    128 1HB  PHE A   7      -4.346  49.960 -43.025  1.00  0.00           H  
ATOM    129 2HB  PHE A   7      -4.745  50.909 -41.602  1.00  0.00           H  
ATOM    130  HD1 PHE A   7      -6.132  49.706 -44.837  1.00  0.00           H  
ATOM    131  HD2 PHE A   7      -6.962  51.150 -40.903  1.00  0.00           H  
ATOM    132  HE1 PHE A   7      -8.522  49.151 -45.140  1.00  0.00           H  
ATOM    133  HE2 PHE A   7      -9.353  50.594 -41.201  1.00  0.00           H  
ATOM    134  HZ  PHE A   7     -10.133  49.595 -43.321  1.00  0.00           H  
ATOM    135  N   ASN A   8      -6.254  53.751 -43.611  1.00  0.00           N  
ATOM    136  CA  ASN A   8      -7.055  54.911 -43.215  1.00  0.00           C  
ATOM    137  C   ASN A   8      -8.108  54.506 -42.194  1.00  0.00           C  
ATOM    138  O   ASN A   8      -9.264  54.267 -42.539  1.00  0.00           O  
ATOM    139  CB  ASN A   8      -7.703  55.561 -44.424  1.00  0.00           C  
ATOM    140  CG  ASN A   8      -8.378  56.862 -44.088  1.00  0.00           C  
ATOM    141  ND2 ASN A   8      -8.509  57.721 -45.066  1.00  0.00           N  
ATOM    142  OD1 ASN A   8      -8.781  57.088 -42.941  1.00  0.00           O  
ATOM    143  H   ASN A   8      -6.454  53.291 -44.487  1.00  0.00           H  
ATOM    144  HA  ASN A   8      -6.390  55.659 -42.782  1.00  0.00           H  
ATOM    145 1HB  ASN A   8      -6.946  55.745 -45.188  1.00  0.00           H  
ATOM    146 2HB  ASN A   8      -8.441  54.882 -44.851  1.00  0.00           H  
ATOM    147 1HD2 ASN A   8      -8.951  58.604 -44.901  1.00  0.00           H  
ATOM    148 2HD2 ASN A   8      -8.168  57.496 -45.978  1.00  0.00           H  
ATOM    149  N   GLN A   9      -7.712  54.444 -40.929  1.00  0.00           N  
ATOM    150  CA  GLN A   9      -8.638  54.041 -39.884  1.00  0.00           C  
ATOM    151  C   GLN A   9      -9.783  55.023 -39.703  1.00  0.00           C  
ATOM    152  O   GLN A   9     -10.906  54.622 -39.413  1.00  0.00           O  
ATOM    153  CB  GLN A   9      -7.891  53.856 -38.566  1.00  0.00           C  
ATOM    154  CG  GLN A   9      -7.296  55.119 -37.993  1.00  0.00           C  
ATOM    155  CD  GLN A   9      -6.452  54.832 -36.770  1.00  0.00           C  
ATOM    156  NE2 GLN A   9      -5.231  55.361 -36.753  1.00  0.00           N  
ATOM    157  OE1 GLN A   9      -6.895  54.141 -35.850  1.00  0.00           O  
ATOM    158  H   GLN A   9      -6.760  54.676 -40.681  1.00  0.00           H  
ATOM    159  HA  GLN A   9      -9.089  53.090 -40.169  1.00  0.00           H  
ATOM    160 1HB  GLN A   9      -8.569  53.443 -37.819  1.00  0.00           H  
ATOM    161 2HB  GLN A   9      -7.080  53.142 -38.705  1.00  0.00           H  
ATOM    162 1HG  GLN A   9      -6.666  55.587 -38.750  1.00  0.00           H  
ATOM    163 2HG  GLN A   9      -8.104  55.794 -37.711  1.00  0.00           H  
ATOM    164 1HE2 GLN A   9      -4.630  55.204 -35.969  1.00  0.00           H  
ATOM    165 2HE2 GLN A   9      -4.915  55.914 -37.522  1.00  0.00           H  
ATOM    166  N   GLU A  10      -9.505  56.311 -39.881  1.00  0.00           N  
ATOM    167  CA  GLU A  10     -10.537  57.330 -39.728  1.00  0.00           C  
ATOM    168  C   GLU A  10     -11.683  57.127 -40.715  1.00  0.00           C  
ATOM    169  O   GLU A  10     -12.854  57.200 -40.344  1.00  0.00           O  
ATOM    170  CB  GLU A  10      -9.935  58.724 -39.915  1.00  0.00           C  
ATOM    171  CG  GLU A  10      -9.019  59.170 -38.783  1.00  0.00           C  
ATOM    172  CD  GLU A  10      -8.373  60.503 -39.044  1.00  0.00           C  
ATOM    173  OE1 GLU A  10      -8.555  61.030 -40.115  1.00  0.00           O  
ATOM    174  OE2 GLU A  10      -7.697  60.994 -38.171  1.00  0.00           O1-
ATOM    175  H   GLU A  10      -8.567  56.594 -40.125  1.00  0.00           H  
ATOM    176  HA  GLU A  10     -10.949  57.258 -38.721  1.00  0.00           H  
ATOM    177 1HB  GLU A  10      -9.360  58.752 -40.841  1.00  0.00           H  
ATOM    178 2HB  GLU A  10     -10.736  59.458 -40.005  1.00  0.00           H  
ATOM    179 1HG  GLU A  10      -9.600  59.235 -37.864  1.00  0.00           H  
ATOM    180 2HG  GLU A  10      -8.245  58.418 -38.639  1.00  0.00           H  
ATOM    181  N   ALA A  11     -11.342  56.870 -41.974  1.00  0.00           N  
ATOM    182  CA  ALA A  11     -12.353  56.657 -43.002  1.00  0.00           C  
ATOM    183  C   ALA A  11     -13.117  55.360 -42.739  1.00  0.00           C  
ATOM    184  O   ALA A  11     -14.340  55.303 -42.883  1.00  0.00           O  
ATOM    185  CB  ALA A  11     -11.696  56.610 -44.375  1.00  0.00           C  
ATOM    186  H   ALA A  11     -10.366  56.821 -42.226  1.00  0.00           H  
ATOM    187  HA  ALA A  11     -13.050  57.494 -42.968  1.00  0.00           H  
ATOM    188 1HB  ALA A  11     -12.458  56.450 -45.138  1.00  0.00           H  
ATOM    189 2HB  ALA A  11     -11.183  57.552 -44.566  1.00  0.00           H  
ATOM    190 3HB  ALA A  11     -10.976  55.794 -44.405  1.00  0.00           H  
ATOM    191  N   TYR A  12     -12.384  54.322 -42.350  1.00  0.00           N  
ATOM    192  CA  TYR A  12     -12.979  53.016 -42.072  1.00  0.00           C  
ATOM    193  C   TYR A  12     -13.974  53.074 -40.912  1.00  0.00           C  
ATOM    194  O   TYR A  12     -15.140  52.709 -41.066  1.00  0.00           O  
ATOM    195  CB  TYR A  12     -11.884  51.989 -41.775  1.00  0.00           C  
ATOM    196  CG  TYR A  12     -12.414  50.617 -41.422  1.00  0.00           C  
ATOM    197  CD1 TYR A  12     -13.288  49.968 -42.282  1.00  0.00           C  
ATOM    198  CD2 TYR A  12     -12.026  50.007 -40.238  1.00  0.00           C  
ATOM    199  CE1 TYR A  12     -13.772  48.715 -41.960  1.00  0.00           C  
ATOM    200  CE2 TYR A  12     -12.509  48.754 -39.916  1.00  0.00           C  
ATOM    201  CZ  TYR A  12     -13.379  48.109 -40.772  1.00  0.00           C  
ATOM    202  OH  TYR A  12     -13.861  46.861 -40.450  1.00  0.00           O  
ATOM    203  H   TYR A  12     -11.387  54.441 -42.244  1.00  0.00           H  
ATOM    204  HA  TYR A  12     -13.522  52.688 -42.958  1.00  0.00           H  
ATOM    205 1HB  TYR A  12     -11.233  51.889 -42.645  1.00  0.00           H  
ATOM    206 2HB  TYR A  12     -11.271  52.340 -40.946  1.00  0.00           H  
ATOM    207  HD1 TYR A  12     -13.593  50.448 -43.212  1.00  0.00           H  
ATOM    208  HD2 TYR A  12     -11.339  50.517 -39.562  1.00  0.00           H  
ATOM    209  HE1 TYR A  12     -14.459  48.206 -42.635  1.00  0.00           H  
ATOM    210  HE2 TYR A  12     -12.205  48.275 -38.985  1.00  0.00           H  
ATOM    211  HH  TYR A  12     -14.449  46.557 -41.146  1.00  0.00           H  
ATOM    212  N   VAL A  13     -13.513  53.535 -39.754  1.00  0.00           N  
ATOM    213  CA  VAL A  13     -14.361  53.627 -38.565  1.00  0.00           C  
ATOM    214  C   VAL A  13     -15.416  54.732 -38.655  1.00  0.00           C  
ATOM    215  O   VAL A  13     -16.525  54.586 -38.137  1.00  0.00           O  
ATOM    216  CB  VAL A  13     -13.486  53.875 -37.322  1.00  0.00           C  
ATOM    217  CG1 VAL A  13     -14.356  54.132 -36.101  1.00  0.00           C  
ATOM    218  CG2 VAL A  13     -12.567  52.686 -37.090  1.00  0.00           C  
ATOM    219  H   VAL A  13     -12.550  53.830 -39.694  1.00  0.00           H  
ATOM    220  HA  VAL A  13     -14.902  52.687 -38.452  1.00  0.00           H  
ATOM    221  HB  VAL A  13     -12.887  54.772 -37.484  1.00  0.00           H  
ATOM    222 1HG1 VAL A  13     -13.721  54.305 -35.232  1.00  0.00           H  
ATOM    223 2HG1 VAL A  13     -14.979  55.009 -36.275  1.00  0.00           H  
ATOM    224 3HG1 VAL A  13     -14.990  53.265 -35.918  1.00  0.00           H  
ATOM    225 1HG2 VAL A  13     -11.950  52.868 -36.210  1.00  0.00           H  
ATOM    226 2HG2 VAL A  13     -13.166  51.788 -36.934  1.00  0.00           H  
ATOM    227 3HG2 VAL A  13     -11.925  52.546 -37.960  1.00  0.00           H  
ATOM    228  N   GLY A  14     -15.071  55.830 -39.319  1.00  0.00           N  
ATOM    229  CA  GLY A  14     -15.995  56.945 -39.443  1.00  0.00           C  
ATOM    230  C   GLY A  14     -17.345  56.614 -40.053  1.00  0.00           C  
ATOM    231  O   GLY A  14     -18.342  57.268 -39.749  1.00  0.00           O  
ATOM    232  H   GLY A  14     -14.158  55.899 -39.746  1.00  0.00           H  
ATOM    233 1HA  GLY A  14     -16.180  57.376 -38.458  1.00  0.00           H  
ATOM    234 2HA  GLY A  14     -15.544  57.726 -40.055  1.00  0.00           H  
ATOM    235  N   GLY A  15     -17.385  55.594 -40.904  1.00  0.00           N  
ATOM    236  CA  GLY A  15     -18.632  55.222 -41.551  1.00  0.00           C  
ATOM    237  C   GLY A  15     -19.646  54.453 -40.724  1.00  0.00           C  
ATOM    238  O   GLY A  15     -20.671  54.023 -41.259  1.00  0.00           O  
ATOM    239  H   GLY A  15     -16.548  55.067 -41.108  1.00  0.00           H  
ATOM    240 1HA  GLY A  15     -19.139  56.120 -41.904  1.00  0.00           H  
ATOM    241 2HA  GLY A  15     -18.419  54.609 -42.425  1.00  0.00           H  
ATOM    242  N   THR A  16     -19.391  54.280 -39.430  1.00  0.00           N  
ATOM    243  CA  THR A  16     -20.319  53.541 -38.577  1.00  0.00           C  
ATOM    244  C   THR A  16     -20.337  54.042 -37.137  1.00  0.00           C  
ATOM    245  O   THR A  16     -20.783  53.329 -36.235  1.00  0.00           O  
ATOM    246  CB  THR A  16     -19.972  52.033 -38.541  1.00  0.00           C  
ATOM    247  CG2 THR A  16     -20.004  51.443 -39.946  1.00  0.00           C  
ATOM    248  OG1 THR A  16     -18.667  51.854 -37.976  1.00  0.00           O  
ATOM    249  H   THR A  16     -18.547  54.660 -39.026  1.00  0.00           H  
ATOM    250  HA  THR A  16     -21.324  53.650 -38.983  1.00  0.00           H  
ATOM    251  HB  THR A  16     -20.695  51.508 -37.917  1.00  0.00           H  
ATOM    252  HG1 THR A  16     -18.298  52.711 -37.746  1.00  0.00           H  
ATOM    253 1HG2 THR A  16     -19.759  50.382 -39.900  1.00  0.00           H  
ATOM    254 2HG2 THR A  16     -21.000  51.567 -40.370  1.00  0.00           H  
ATOM    255 3HG2 THR A  16     -19.276  51.957 -40.572  1.00  0.00           H  
ATOM    256  N   MET A  17     -19.868  55.265 -36.916  1.00  0.00           N  
ATOM    257  CA  MET A  17     -19.836  55.806 -35.563  1.00  0.00           C  
ATOM    258  C   MET A  17     -21.224  56.048 -34.986  1.00  0.00           C  
ATOM    259  O   MET A  17     -22.197  56.239 -35.720  1.00  0.00           O  
ATOM    260  CB  MET A  17     -19.031  57.104 -35.547  1.00  0.00           C  
ATOM    261  CG  MET A  17     -17.541  56.924 -35.803  1.00  0.00           C  
ATOM    262  SD  MET A  17     -16.648  58.490 -35.839  1.00  0.00           S  
ATOM    263  CE  MET A  17     -16.732  58.968 -34.116  1.00  0.00           C  
ATOM    264  H   MET A  17     -19.530  55.831 -37.680  1.00  0.00           H  
ATOM    265  HA  MET A  17     -19.367  55.072 -34.908  1.00  0.00           H  
ATOM    266 1HB  MET A  17     -19.420  57.783 -36.304  1.00  0.00           H  
ATOM    267 2HB  MET A  17     -19.147  57.592 -34.578  1.00  0.00           H  
ATOM    268 1HG  MET A  17     -17.110  56.299 -35.022  1.00  0.00           H  
ATOM    269 2HG  MET A  17     -17.395  56.422 -36.759  1.00  0.00           H  
ATOM    270 1HE  MET A  17     -16.222  59.921 -33.977  1.00  0.00           H  
ATOM    271 2HE  MET A  17     -17.777  59.068 -33.816  1.00  0.00           H  
ATOM    272 3HE  MET A  17     -16.250  58.206 -33.503  1.00  0.00           H  
ATOM    273  N   VAL A  18     -21.306  56.023 -33.660  1.00  0.00           N  
ATOM    274  CA  VAL A  18     -22.567  56.236 -32.961  1.00  0.00           C  
ATOM    275  C   VAL A  18     -23.014  57.683 -33.124  1.00  0.00           C  
ATOM    276  O   VAL A  18     -22.244  58.612 -32.881  1.00  0.00           O  
ATOM    277  CB  VAL A  18     -22.421  55.904 -31.464  1.00  0.00           C  
ATOM    278  CG1 VAL A  18     -23.702  56.241 -30.716  1.00  0.00           C  
ATOM    279  CG2 VAL A  18     -22.066  54.435 -31.292  1.00  0.00           C  
ATOM    280  H   VAL A  18     -20.469  55.850 -33.122  1.00  0.00           H  
ATOM    281  HA  VAL A  18     -23.314  55.561 -33.381  1.00  0.00           H  
ATOM    282  HB  VAL A  18     -21.629  56.522 -31.040  1.00  0.00           H  
ATOM    283 1HG1 VAL A  18     -23.581  56.001 -29.660  1.00  0.00           H  
ATOM    284 2HG1 VAL A  18     -23.917  57.304 -30.823  1.00  0.00           H  
ATOM    285 3HG1 VAL A  18     -24.527  55.660 -31.127  1.00  0.00           H  
ATOM    286 1HG2 VAL A  18     -21.963  54.208 -30.231  1.00  0.00           H  
ATOM    287 2HG2 VAL A  18     -22.855  53.817 -31.721  1.00  0.00           H  
ATOM    288 3HG2 VAL A  18     -21.125  54.226 -31.800  1.00  0.00           H  
ATOM    289  N   ARG A  19     -24.260  57.868 -33.538  1.00  0.00           N  
ATOM    290  CA  ARG A  19     -24.798  59.207 -33.725  1.00  0.00           C  
ATOM    291  C   ARG A  19     -25.103  59.838 -32.375  1.00  0.00           C  
ATOM    292  O   ARG A  19     -25.405  59.138 -31.409  1.00  0.00           O  
ATOM    293  CB  ARG A  19     -26.062  59.172 -34.571  1.00  0.00           C  
ATOM    294  CG  ARG A  19     -25.851  58.759 -36.019  1.00  0.00           C  
ATOM    295  CD  ARG A  19     -27.135  58.696 -36.765  1.00  0.00           C  
ATOM    296  NE  ARG A  19     -27.994  57.629 -36.276  1.00  0.00           N  
ATOM    297  CZ  ARG A  19     -29.270  57.437 -36.662  1.00  0.00           C  
ATOM    298  NH1 ARG A  19     -29.820  58.247 -37.540  1.00  0.00           N1+
ATOM    299  NH2 ARG A  19     -29.970  56.436 -36.158  1.00  0.00           N  
ATOM    300  H   ARG A  19     -24.850  57.070 -33.727  1.00  0.00           H  
ATOM    301  HA  ARG A  19     -24.055  59.808 -34.249  1.00  0.00           H  
ATOM    302 1HB  ARG A  19     -26.775  58.476 -34.131  1.00  0.00           H  
ATOM    303 2HB  ARG A  19     -26.526  60.158 -34.573  1.00  0.00           H  
ATOM    304 1HG  ARG A  19     -25.203  59.483 -36.513  1.00  0.00           H  
ATOM    305 2HG  ARG A  19     -25.386  57.773 -36.053  1.00  0.00           H  
ATOM    306 1HD  ARG A  19     -27.667  59.640 -36.653  1.00  0.00           H  
ATOM    307 2HD  ARG A  19     -26.934  58.515 -37.820  1.00  0.00           H  
ATOM    308  HE  ARG A  19     -27.606  56.986 -35.599  1.00  0.00           H  
ATOM    309 1HH1 ARG A  19     -29.285  59.012 -37.925  1.00  0.00           H  
ATOM    310 2HH1 ARG A  19     -30.777  58.103 -37.829  1.00  0.00           H  
ATOM    311 1HH2 ARG A  19     -29.547  55.813 -35.483  1.00  0.00           H  
ATOM    312 2HH2 ARG A  19     -30.926  56.292 -36.447  1.00  0.00           H  
ATOM    313  N   SER A  20     -25.018  61.162 -32.306  1.00  0.00           N  
ATOM    314  CA  SER A  20     -25.292  61.860 -31.059  1.00  0.00           C  
ATOM    315  C   SER A  20     -26.697  61.540 -30.571  1.00  0.00           C  
ATOM    316  O   SER A  20     -27.651  61.542 -31.350  1.00  0.00           O  
ATOM    317  CB  SER A  20     -25.135  63.356 -31.247  1.00  0.00           C  
ATOM    318  OG  SER A  20     -25.477  64.048 -30.077  1.00  0.00           O  
ATOM    319  H   SER A  20     -24.760  61.697 -33.123  1.00  0.00           H  
ATOM    320  HA  SER A  20     -24.576  61.522 -30.308  1.00  0.00           H  
ATOM    321 1HB  SER A  20     -24.104  63.583 -31.518  1.00  0.00           H  
ATOM    322 2HB  SER A  20     -25.769  63.689 -32.067  1.00  0.00           H  
ATOM    323  HG  SER A  20     -26.248  63.601 -29.721  1.00  0.00           H  
ATOM    324  N   GLY A  21     -26.815  61.253 -29.279  1.00  0.00           N  
ATOM    325  CA  GLY A  21     -28.110  60.932 -28.706  1.00  0.00           C  
ATOM    326  C   GLY A  21     -28.435  59.449 -28.737  1.00  0.00           C  
ATOM    327  O   GLY A  21     -29.352  58.996 -28.050  1.00  0.00           O  
ATOM    328  H   GLY A  21     -26.000  61.255 -28.682  1.00  0.00           H  
ATOM    329 1HA  GLY A  21     -28.146  61.273 -27.671  1.00  0.00           H  
ATOM    330 2HA  GLY A  21     -28.890  61.468 -29.247  1.00  0.00           H  
ATOM    331  N   GLN A  22     -27.685  58.687 -29.527  1.00  0.00           N  
ATOM    332  CA  GLN A  22     -27.918  57.252 -29.635  1.00  0.00           C  
ATOM    333  C   GLN A  22     -27.160  56.442 -28.585  1.00  0.00           C  
ATOM    334  O   GLN A  22     -26.084  56.833 -28.135  1.00  0.00           O  
ATOM    335  CB  GLN A  22     -27.532  56.766 -31.034  1.00  0.00           C  
ATOM    336  CG  GLN A  22     -28.322  57.413 -32.159  1.00  0.00           C  
ATOM    337  CD  GLN A  22     -29.807  57.115 -32.069  1.00  0.00           C  
ATOM    338  NE2 GLN A  22     -30.618  58.165 -32.018  1.00  0.00           N  
ATOM    339  OE1 GLN A  22     -30.220  55.952 -32.047  1.00  0.00           O  
ATOM    340  H   GLN A  22     -26.940  59.107 -30.064  1.00  0.00           H  
ATOM    341  HA  GLN A  22     -28.976  57.058 -29.458  1.00  0.00           H  
ATOM    342 1HB  GLN A  22     -26.474  56.963 -31.209  1.00  0.00           H  
ATOM    343 2HB  GLN A  22     -27.679  55.687 -31.098  1.00  0.00           H  
ATOM    344 1HG  GLN A  22     -28.187  58.494 -32.108  1.00  0.00           H  
ATOM    345 2HG  GLN A  22     -27.954  57.035 -33.112  1.00  0.00           H  
ATOM    346 1HE2 GLN A  22     -31.608  58.030 -31.959  1.00  0.00           H  
ATOM    347 2HE2 GLN A  22     -30.241  59.091 -32.039  1.00  0.00           H  
ATOM    348  N   ASP A  23     -27.745  55.310 -28.204  1.00  0.00           N  
ATOM    349  CA  ASP A  23     -27.177  54.412 -27.199  1.00  0.00           C  
ATOM    350  C   ASP A  23     -25.850  53.820 -27.667  1.00  0.00           C  
ATOM    351  O   ASP A  23     -25.796  53.101 -28.662  1.00  0.00           O  
ATOM    352  CB  ASP A  23     -28.187  53.296 -26.910  1.00  0.00           C  
ATOM    353  CG  ASP A  23     -27.750  52.372 -25.791  1.00  0.00           C  
ATOM    354  OD1 ASP A  23     -28.571  51.521 -25.393  1.00  0.00           O  
ATOM    355  OD2 ASP A  23     -26.604  52.485 -25.311  1.00  0.00           O1-
ATOM    356  H   ASP A  23     -28.624  55.070 -28.637  1.00  0.00           H  
ATOM    357  HA  ASP A  23     -26.996  54.983 -26.288  1.00  0.00           H  
ATOM    358 1HB  ASP A  23     -29.148  53.736 -26.641  1.00  0.00           H  
ATOM    359 2HB  ASP A  23     -28.339  52.701 -27.811  1.00  0.00           H  
ATOM    360  N   PRO A  24     -24.754  54.118 -26.950  1.00  0.00           N  
ATOM    361  CA  PRO A  24     -23.441  53.590 -27.340  1.00  0.00           C  
ATOM    362  C   PRO A  24     -23.218  52.104 -27.063  1.00  0.00           C  
ATOM    363  O   PRO A  24     -22.168  51.566 -27.407  1.00  0.00           O  
ATOM    364  CB  PRO A  24     -22.492  54.447 -26.496  1.00  0.00           C  
ATOM    365  CG  PRO A  24     -23.291  54.793 -25.286  1.00  0.00           C  
ATOM    366  CD  PRO A  24     -24.697  54.958 -25.800  1.00  0.00           C  
ATOM    367  HA  PRO A  24     -23.307  53.718 -28.424  1.00  0.00           H  
ATOM    368 1HB  PRO A  24     -21.582  53.877 -26.257  1.00  0.00           H  
ATOM    369 2HB  PRO A  24     -22.176  55.332 -27.067  1.00  0.00           H  
ATOM    370 1HG  PRO A  24     -23.207  53.996 -24.533  1.00  0.00           H  
ATOM    371 2HG  PRO A  24     -22.900  55.711 -24.822  1.00  0.00           H  
ATOM    372 1HD  PRO A  24     -25.410  54.623 -25.033  1.00  0.00           H  
ATOM    373 2HD  PRO A  24     -24.872  56.013 -26.059  1.00  0.00           H  
ATOM    374  N   TYR A  25     -24.202  51.448 -26.452  1.00  0.00           N  
ATOM    375  CA  TYR A  25     -24.103  50.021 -26.124  1.00  0.00           C  
ATOM    376  C   TYR A  25     -25.037  49.142 -26.949  1.00  0.00           C  
ATOM    377  O   TYR A  25     -24.900  47.920 -26.954  1.00  0.00           O  
ATOM    378  CB  TYR A  25     -24.381  49.807 -24.634  1.00  0.00           C  
ATOM    379  CG  TYR A  25     -23.390  50.499 -23.724  1.00  0.00           C  
ATOM    380  CD1 TYR A  25     -23.674  51.761 -23.221  1.00  0.00           C  
ATOM    381  CD2 TYR A  25     -22.199  49.872 -23.391  1.00  0.00           C  
ATOM    382  CE1 TYR A  25     -22.770  52.392 -22.390  1.00  0.00           C  
ATOM    383  CE2 TYR A  25     -21.295  50.504 -22.559  1.00  0.00           C  
ATOM    384  CZ  TYR A  25     -21.577  51.759 -22.059  1.00  0.00           C  
ATOM    385  OH  TYR A  25     -20.677  52.388 -21.231  1.00  0.00           O  
ATOM    386  H   TYR A  25     -25.042  51.952 -26.208  1.00  0.00           H  
ATOM    387  HA  TYR A  25     -23.079  49.695 -26.305  1.00  0.00           H  
ATOM    388 1HB  TYR A  25     -25.379  50.175 -24.393  1.00  0.00           H  
ATOM    389 2HB  TYR A  25     -24.362  48.741 -24.410  1.00  0.00           H  
ATOM    390  HD1 TYR A  25     -24.611  52.254 -23.483  1.00  0.00           H  
ATOM    391  HD2 TYR A  25     -21.976  48.881 -23.786  1.00  0.00           H  
ATOM    392  HE1 TYR A  25     -22.993  53.383 -21.995  1.00  0.00           H  
ATOM    393  HE2 TYR A  25     -20.358  50.011 -22.298  1.00  0.00           H  
ATOM    394  HH  TYR A  25     -19.912  51.822 -21.103  1.00  0.00           H  
ATOM    395  N   ALA A  26     -25.986  49.764 -27.638  1.00  0.00           N  
ATOM    396  CA  ALA A  26     -26.982  49.038 -28.421  1.00  0.00           C  
ATOM    397  C   ALA A  26     -26.469  48.009 -29.426  1.00  0.00           C  
ATOM    398  O   ALA A  26     -26.924  46.860 -29.443  1.00  0.00           O  
ATOM    399  CB  ALA A  26     -27.889  50.038 -29.137  1.00  0.00           C  
ATOM    400  H   ALA A  26     -26.016  50.774 -27.618  1.00  0.00           H  
ATOM    401  HA  ALA A  26     -27.577  48.435 -27.736  1.00  0.00           H  
ATOM    402 1HB  ALA A  26     -28.634  49.499 -29.722  1.00  0.00           H  
ATOM    403 2HB  ALA A  26     -28.391  50.666 -28.400  1.00  0.00           H  
ATOM    404 3HB  ALA A  26     -27.291  50.663 -29.797  1.00  0.00           H  
ATOM    405  N   ARG A  27     -25.521  48.416 -30.259  1.00  0.00           N  
ATOM    406  CA  ARG A  27     -25.003  47.532 -31.296  1.00  0.00           C  
ATOM    407  C   ARG A  27     -24.049  46.422 -30.871  1.00  0.00           C  
ATOM    408  O   ARG A  27     -24.104  45.320 -31.421  1.00  0.00           O  
ATOM    409  CB  ARG A  27     -24.295  48.375 -32.346  1.00  0.00           C  
ATOM    410  CG  ARG A  27     -25.210  49.250 -33.188  1.00  0.00           C  
ATOM    411  CD  ARG A  27     -24.443  50.074 -34.157  1.00  0.00           C  
ATOM    412  NE  ARG A  27     -25.314  50.913 -34.964  1.00  0.00           N  
ATOM    413  CZ  ARG A  27     -24.885  51.834 -35.849  1.00  0.00           C  
ATOM    414  NH1 ARG A  27     -23.596  52.022 -36.028  1.00  0.00           N1+
ATOM    415  NH2 ARG A  27     -25.759  52.548 -36.536  1.00  0.00           N  
ATOM    416  H   ARG A  27     -25.149  49.352 -30.179  1.00  0.00           H  
ATOM    417  HA  ARG A  27     -25.839  46.987 -31.734  1.00  0.00           H  
ATOM    418 1HB  ARG A  27     -23.570  49.027 -31.861  1.00  0.00           H  
ATOM    419 2HB  ARG A  27     -23.745  47.723 -33.026  1.00  0.00           H  
ATOM    420 1HG  ARG A  27     -25.903  48.620 -33.746  1.00  0.00           H  
ATOM    421 2HG  ARG A  27     -25.772  49.920 -32.537  1.00  0.00           H  
ATOM    422 1HD  ARG A  27     -23.752  50.721 -33.617  1.00  0.00           H  
ATOM    423 2HD  ARG A  27     -23.882  49.422 -34.825  1.00  0.00           H  
ATOM    424  HE  ARG A  27     -26.313  50.798 -34.853  1.00  0.00           H  
ATOM    425 1HH1 ARG A  27     -22.928  51.476 -35.503  1.00  0.00           H  
ATOM    426 2HH1 ARG A  27     -23.274  52.712 -36.691  1.00  0.00           H  
ATOM    427 1HH2 ARG A  27     -26.750  52.404 -36.398  1.00  0.00           H  
ATOM    428 2HH2 ARG A  27     -25.437  53.238 -37.199  1.00  0.00           H  
ATOM    429  N   ASN A  28     -23.183  46.696 -29.900  1.00  0.00           N  
ATOM    430  CA  ASN A  28     -22.185  45.706 -29.496  1.00  0.00           C  
ATOM    431  C   ASN A  28     -22.206  45.213 -28.058  1.00  0.00           C  
ATOM    432  O   ASN A  28     -21.470  44.293 -27.709  1.00  0.00           O  
ATOM    433  CB  ASN A  28     -20.804  46.256 -29.801  1.00  0.00           C  
ATOM    434  CG  ASN A  28     -20.571  46.447 -31.274  1.00  0.00           C  
ATOM    435  ND2 ASN A  28     -20.664  47.672 -31.726  1.00  0.00           N  
ATOM    436  OD1 ASN A  28     -20.306  45.485 -32.004  1.00  0.00           O  
ATOM    437  H   ASN A  28     -23.205  47.591 -29.431  1.00  0.00           H  
ATOM    438  HA  ASN A  28     -22.337  44.801 -30.087  1.00  0.00           H  
ATOM    439 1HB  ASN A  28     -20.674  47.215 -29.297  1.00  0.00           H  
ATOM    440 2HB  ASN A  28     -20.047  45.576 -29.411  1.00  0.00           H  
ATOM    441 1HD2 ASN A  28     -20.519  47.859 -32.698  1.00  0.00           H  
ATOM    442 2HD2 ASN A  28     -20.880  48.420 -31.100  1.00  0.00           H  
ATOM    443  N   LYS A  29     -23.035  45.820 -27.224  1.00  0.00           N  
ATOM    444  CA  LYS A  29     -23.088  45.454 -25.816  1.00  0.00           C  
ATOM    445  C   LYS A  29     -21.770  45.808 -25.117  1.00  0.00           C  
ATOM    446  O   LYS A  29     -21.379  45.192 -24.131  1.00  0.00           O  
ATOM    447  CB  LYS A  29     -23.445  43.971 -25.641  1.00  0.00           C  
ATOM    448  CG  LYS A  29     -24.890  43.662 -26.018  1.00  0.00           C  
ATOM    449  CD  LYS A  29     -25.868  44.479 -25.174  1.00  0.00           C  
ATOM    450  CE  LYS A  29     -27.312  44.218 -25.579  1.00  0.00           C  
ATOM    451  NZ  LYS A  29     -28.273  45.035 -24.777  1.00  0.00           N1+
ATOM    452  H   LYS A  29     -23.644  46.550 -27.565  1.00  0.00           H  
ATOM    453  HA  LYS A  29     -23.860  46.053 -25.332  1.00  0.00           H  
ATOM    454 1HB  LYS A  29     -22.786  43.362 -26.260  1.00  0.00           H  
ATOM    455 2HB  LYS A  29     -23.285  43.678 -24.603  1.00  0.00           H  
ATOM    456 1HG  LYS A  29     -25.050  43.895 -27.071  1.00  0.00           H  
ATOM    457 2HG  LYS A  29     -25.087  42.602 -25.864  1.00  0.00           H  
ATOM    458 1HD  LYS A  29     -25.744  44.221 -24.121  1.00  0.00           H  
ATOM    459 2HD  LYS A  29     -25.655  45.542 -25.297  1.00  0.00           H  
ATOM    460 1HE  LYS A  29     -27.443  44.456 -26.633  1.00  0.00           H  
ATOM    461 2HE  LYS A  29     -27.545  43.163 -25.436  1.00  0.00           H  
ATOM    462 1HZ  LYS A  29     -29.217  44.834 -25.075  1.00  0.00           H  
ATOM    463 2HZ  LYS A  29     -28.173  44.809 -23.797  1.00  0.00           H  
ATOM    464 3HZ  LYS A  29     -28.078  46.017 -24.917  1.00  0.00           H  
ATOM    465  N   PHE A  30     -21.081  46.792 -25.687  1.00  0.00           N  
ATOM    466  CA  PHE A  30     -19.871  47.379 -25.122  1.00  0.00           C  
ATOM    467  C   PHE A  30     -19.881  48.788 -25.699  1.00  0.00           C  
ATOM    468  O   PHE A  30     -20.542  49.029 -26.712  1.00  0.00           O  
ATOM    469  CB  PHE A  30     -18.563  46.592 -25.416  1.00  0.00           C  
ATOM    470  CG  PHE A  30     -18.159  46.488 -26.873  1.00  0.00           C  
ATOM    471  CD1 PHE A  30     -17.926  47.618 -27.651  1.00  0.00           C  
ATOM    472  CD2 PHE A  30     -17.902  45.233 -27.428  1.00  0.00           C  
ATOM    473  CE1 PHE A  30     -17.437  47.498 -28.958  1.00  0.00           C  
ATOM    474  CE2 PHE A  30     -17.416  45.102 -28.727  1.00  0.00           C  
ATOM    475  CZ  PHE A  30     -17.182  46.236 -29.495  1.00  0.00           C  
ATOM    476  H   PHE A  30     -21.433  47.143 -26.566  1.00  0.00           H  
ATOM    477  HA  PHE A  30     -19.976  47.410 -24.036  1.00  0.00           H  
ATOM    478 1HB  PHE A  30     -17.732  47.059 -24.889  1.00  0.00           H  
ATOM    479 2HB  PHE A  30     -18.659  45.574 -25.041  1.00  0.00           H  
ATOM    480  HD1 PHE A  30     -18.127  48.604 -27.232  1.00  0.00           H  
ATOM    481  HD2 PHE A  30     -18.086  44.344 -26.823  1.00  0.00           H  
ATOM    482  HE1 PHE A  30     -17.256  48.391 -29.556  1.00  0.00           H  
ATOM    483  HE2 PHE A  30     -17.218  44.114 -29.141  1.00  0.00           H  
ATOM    484  HZ  PHE A  30     -16.804  46.142 -30.512  1.00  0.00           H  
ATOM    485  N   ASN A  31     -19.203  49.726 -25.047  1.00  0.00           N  
ATOM    486  CA  ASN A  31     -19.230  51.109 -25.501  1.00  0.00           C  
ATOM    487  C   ASN A  31     -18.475  51.365 -26.798  1.00  0.00           C  
ATOM    488  O   ASN A  31     -17.243  51.400 -26.819  1.00  0.00           O  
ATOM    489  CB  ASN A  31     -18.697  52.016 -24.399  1.00  0.00           C  
ATOM    490  CG  ASN A  31     -19.075  53.454 -24.617  1.00  0.00           C  
ATOM    491  ND2 ASN A  31     -19.723  54.053 -23.627  1.00  0.00           N  
ATOM    492  OD1 ASN A  31     -18.787  54.026 -25.666  1.00  0.00           O  
ATOM    493  H   ASN A  31     -18.662  49.487 -24.229  1.00  0.00           H  
ATOM    494  HA  ASN A  31     -20.264  51.381 -25.719  1.00  0.00           H  
ATOM    495 1HB  ASN A  31     -19.088  51.687 -23.436  1.00  0.00           H  
ATOM    496 2HB  ASN A  31     -17.610  51.936 -24.356  1.00  0.00           H  
ATOM    497 1HD2 ASN A  31     -20.000  55.010 -23.717  1.00  0.00           H  
ATOM    498 2HD2 ASN A  31     -19.935  53.548 -22.791  1.00  0.00           H  
ATOM    499  N   GLN A  32     -19.221  51.566 -27.881  1.00  0.00           N  
ATOM    500  CA  GLN A  32     -18.595  51.794 -29.174  1.00  0.00           C  
ATOM    501  C   GLN A  32     -17.950  53.171 -29.262  1.00  0.00           C  
ATOM    502  O   GLN A  32     -16.963  53.355 -29.970  1.00  0.00           O  
ATOM    503  CB  GLN A  32     -19.607  51.627 -30.310  1.00  0.00           C  
ATOM    504  CG  GLN A  32     -18.934  51.523 -31.670  1.00  0.00           C  
ATOM    505  CD  GLN A  32     -19.903  51.614 -32.827  1.00  0.00           C  
ATOM    506  NE2 GLN A  32     -19.607  52.498 -33.774  1.00  0.00           N  
ATOM    507  OE1 GLN A  32     -20.903  50.893 -32.878  1.00  0.00           O  
ATOM    508  H   GLN A  32     -20.229  51.563 -27.816  1.00  0.00           H  
ATOM    509  HA  GLN A  32     -17.802  51.058 -29.308  1.00  0.00           H  
ATOM    510 1HB  GLN A  32     -20.202  50.730 -30.138  1.00  0.00           H  
ATOM    511 2HB  GLN A  32     -20.291  52.476 -30.316  1.00  0.00           H  
ATOM    512 1HG  GLN A  32     -18.216  52.336 -31.772  1.00  0.00           H  
ATOM    513 2HG  GLN A  32     -18.423  50.563 -31.738  1.00  0.00           H  
ATOM    514 1HE2 GLN A  32     -20.210  52.605 -34.565  1.00  0.00           H  
ATOM    515 2HE2 GLN A  32     -18.782  53.057 -33.696  1.00  0.00           H  
ATOM    516  N   VAL A  33     -18.505  54.144 -28.548  1.00  0.00           N  
ATOM    517  CA  VAL A  33     -17.932  55.482 -28.576  1.00  0.00           C  
ATOM    518  C   VAL A  33     -16.501  55.419 -28.050  1.00  0.00           C  
ATOM    519  O   VAL A  33     -15.587  55.998 -28.638  1.00  0.00           O  
ATOM    520  CB  VAL A  33     -18.774  56.467 -27.732  1.00  0.00           C  
ATOM    521  CG1 VAL A  33     -18.041  57.795 -27.577  1.00  0.00           C  
ATOM    522  CG2 VAL A  33     -20.114  56.696 -28.411  1.00  0.00           C  
ATOM    523  H   VAL A  33     -19.323  53.971 -27.982  1.00  0.00           H  
ATOM    524  HA  VAL A  33     -17.925  55.834 -29.607  1.00  0.00           H  
ATOM    525  HB  VAL A  33     -18.931  56.041 -26.741  1.00  0.00           H  
ATOM    526 1HG1 VAL A  33     -18.646  58.478 -26.980  1.00  0.00           H  
ATOM    527 2HG1 VAL A  33     -17.086  57.628 -27.079  1.00  0.00           H  
ATOM    528 3HG1 VAL A  33     -17.867  58.232 -28.560  1.00  0.00           H  
ATOM    529 1HG2 VAL A  33     -20.709  57.389 -27.818  1.00  0.00           H  
ATOM    530 2HG2 VAL A  33     -19.952  57.114 -29.404  1.00  0.00           H  
ATOM    531 3HG2 VAL A  33     -20.644  55.747 -28.500  1.00  0.00           H  
ATOM    532  N   GLU A  34     -16.303  54.690 -26.955  1.00  0.00           N  
ATOM    533  CA  GLU A  34     -14.973  54.558 -26.377  1.00  0.00           C  
ATOM    534  C   GLU A  34     -14.049  53.688 -27.230  1.00  0.00           C  
ATOM    535  O   GLU A  34     -12.849  53.931 -27.294  1.00  0.00           O  
ATOM    536  CB  GLU A  34     -15.078  54.011 -24.952  1.00  0.00           C  
ATOM    537  CG  GLU A  34     -15.722  55.008 -23.995  1.00  0.00           C  
ATOM    538  CD  GLU A  34     -14.899  56.280 -23.839  1.00  0.00           C  
ATOM    539  OE1 GLU A  34     -15.499  57.363 -23.659  1.00  0.00           O  
ATOM    540  OE2 GLU A  34     -13.651  56.196 -23.884  1.00  0.00           O1-
ATOM    541  H   GLU A  34     -17.080  54.219 -26.515  1.00  0.00           H  
ATOM    542  HA  GLU A  34     -14.510  55.545 -26.344  1.00  0.00           H  
ATOM    543 1HB  GLU A  34     -15.666  53.094 -24.957  1.00  0.00           H  
ATOM    544 2HB  GLU A  34     -14.082  53.760 -24.585  1.00  0.00           H  
ATOM    545 1HG  GLU A  34     -16.712  55.268 -24.369  1.00  0.00           H  
ATOM    546 2HG  GLU A  34     -15.844  54.534 -23.022  1.00  0.00           H  
ATOM    547  N   SER A  35     -14.599  52.673 -27.884  1.00  0.00           N  
ATOM    548  CA  SER A  35     -13.772  51.834 -28.746  1.00  0.00           C  
ATOM    549  C   SER A  35     -13.317  52.670 -29.949  1.00  0.00           C  
ATOM    550  O   SER A  35     -12.165  52.585 -30.383  1.00  0.00           O  
ATOM    551  CB  SER A  35     -14.543  50.612 -29.206  1.00  0.00           C  
ATOM    552  OG  SER A  35     -13.751  49.800 -30.030  1.00  0.00           O  
ATOM    553  H   SER A  35     -15.585  52.469 -27.798  1.00  0.00           H  
ATOM    554  HA  SER A  35     -12.919  51.479 -28.167  1.00  0.00           H  
ATOM    555 1HB  SER A  35     -14.872  50.042 -28.338  1.00  0.00           H  
ATOM    556 2HB  SER A  35     -15.433  50.926 -29.748  1.00  0.00           H  
ATOM    557  HG  SER A  35     -13.770  50.207 -30.899  1.00  0.00           H  
ATOM    558  N   ASP A  36     -14.229  53.481 -30.477  1.00  0.00           N  
ATOM    559  CA  ASP A  36     -13.934  54.316 -31.639  1.00  0.00           C  
ATOM    560  C   ASP A  36     -12.932  55.440 -31.359  1.00  0.00           C  
ATOM    561  O   ASP A  36     -12.313  55.971 -32.284  1.00  0.00           O  
ATOM    562  CB  ASP A  36     -15.231  54.924 -32.178  1.00  0.00           C  
ATOM    563  CG  ASP A  36     -16.132  53.897 -32.851  1.00  0.00           C  
ATOM    564  OD1 ASP A  36     -15.663  52.822 -33.139  1.00  0.00           O  
ATOM    565  OD2 ASP A  36     -17.281  54.198 -33.070  1.00  0.00           O1-
ATOM    566  H   ASP A  36     -15.150  53.520 -30.064  1.00  0.00           H  
ATOM    567  HA  ASP A  36     -13.471  53.693 -32.404  1.00  0.00           H  
ATOM    568 1HB  ASP A  36     -15.782  55.390 -31.361  1.00  0.00           H  
ATOM    569 2HB  ASP A  36     -14.994  55.706 -32.900  1.00  0.00           H  
ATOM    570  N   LYS A  37     -12.771  55.804 -30.091  1.00  0.00           N  
ATOM    571  CA  LYS A  37     -11.835  56.868 -29.734  1.00  0.00           C  
ATOM    572  C   LYS A  37     -10.389  56.412 -29.830  1.00  0.00           C  
ATOM    573  O   LYS A  37      -9.490  57.224 -30.047  1.00  0.00           O  
ATOM    574  CB  LYS A  37     -12.088  57.357 -28.307  1.00  0.00           C  
ATOM    575  CG  LYS A  37     -13.319  58.221 -28.138  1.00  0.00           C  
ATOM    576  CD  LYS A  37     -13.427  58.710 -26.701  1.00  0.00           C  
ATOM    577  CE  LYS A  37     -14.612  59.641 -26.511  1.00  0.00           C  
ATOM    578  NZ  LYS A  37     -14.726  60.069 -25.087  1.00  0.00           N1+
ATOM    579  H   LYS A  37     -13.296  55.345 -29.361  1.00  0.00           H  
ATOM    580  HA  LYS A  37     -11.984  57.703 -30.418  1.00  0.00           H  
ATOM    581 1HB  LYS A  37     -12.194  56.500 -27.641  1.00  0.00           H  
ATOM    582 2HB  LYS A  37     -11.230  57.935 -27.963  1.00  0.00           H  
ATOM    583 1HG  LYS A  37     -13.256  59.078 -28.811  1.00  0.00           H  
ATOM    584 2HG  LYS A  37     -14.206  57.644 -28.395  1.00  0.00           H  
ATOM    585 1HD  LYS A  37     -13.542  57.854 -26.034  1.00  0.00           H  
ATOM    586 2HD  LYS A  37     -12.516  59.242 -26.428  1.00  0.00           H  
ATOM    587 1HE  LYS A  37     -14.489  60.519 -27.143  1.00  0.00           H  
ATOM    588 2HE  LYS A  37     -15.527  59.131 -26.809  1.00  0.00           H  
ATOM    589 1HZ  LYS A  37     -15.519  60.687 -24.984  1.00  0.00           H  
ATOM    590 2HZ  LYS A  37     -14.852  59.257 -24.500  1.00  0.00           H  
ATOM    591 3HZ  LYS A  37     -13.884  60.555 -24.813  1.00  0.00           H  
ATOM    592  N   LEU A  38     -10.171  55.112 -29.671  1.00  0.00           N  
ATOM    593  CA  LEU A  38      -8.826  54.546 -29.697  1.00  0.00           C  
ATOM    594  C   LEU A  38      -8.190  54.376 -31.066  1.00  0.00           C  
ATOM    595  O   LEU A  38      -8.857  54.030 -32.036  1.00  0.00           O  
ATOM    596  CB  LEU A  38      -8.852  53.178 -29.004  1.00  0.00           C  
ATOM    597  CG  LEU A  38      -9.229  53.192 -27.517  1.00  0.00           C  
ATOM    598  CD1 LEU A  38      -9.317  51.762 -27.002  1.00  0.00           C  
ATOM    599  CD2 LEU A  38      -8.194  53.991 -26.739  1.00  0.00           C  
ATOM    600  H   LEU A  38     -10.960  54.499 -29.528  1.00  0.00           H  
ATOM    601  HA  LEU A  38      -8.157  55.219 -29.163  1.00  0.00           H  
ATOM    602 1HB  LEU A  38      -9.567  52.540 -29.521  1.00  0.00           H  
ATOM    603 2HB  LEU A  38      -7.864  52.725 -29.092  1.00  0.00           H  
ATOM    604  HG  LEU A  38     -10.210  53.652 -27.395  1.00  0.00           H  
ATOM    605 1HD1 LEU A  38      -9.585  51.772 -25.945  1.00  0.00           H  
ATOM    606 2HD1 LEU A  38     -10.078  51.219 -27.563  1.00  0.00           H  
ATOM    607 3HD1 LEU A  38      -8.353  51.270 -27.127  1.00  0.00           H  
ATOM    608 1HD2 LEU A  38      -8.463  54.002 -25.682  1.00  0.00           H  
ATOM    609 2HD2 LEU A  38      -7.213  53.531 -26.859  1.00  0.00           H  
ATOM    610 3HD2 LEU A  38      -8.165  55.013 -27.116  1.00  0.00           H  
ATOM    611  N   ARG A  39      -6.885  54.614 -31.131  1.00  0.00           N  
ATOM    612  CA  ARG A  39      -6.152  54.440 -32.373  1.00  0.00           C  
ATOM    613  C   ARG A  39      -5.969  52.938 -32.536  1.00  0.00           C  
ATOM    614  O   ARG A  39      -5.937  52.208 -31.546  1.00  0.00           O  
ATOM    615  CB  ARG A  39      -4.805  55.148 -32.346  1.00  0.00           C  
ATOM    616  CG  ARG A  39      -4.879  56.657 -32.174  1.00  0.00           C  
ATOM    617  CD  ARG A  39      -5.402  57.324 -33.394  1.00  0.00           C  
ATOM    618  NE  ARG A  39      -5.394  58.772 -33.266  1.00  0.00           N  
ATOM    619  CZ  ARG A  39      -5.827  59.626 -34.214  1.00  0.00           C  
ATOM    620  NH1 ARG A  39      -6.297  59.163 -35.350  1.00  0.00           N1+
ATOM    621  NH2 ARG A  39      -5.778  60.930 -34.001  1.00  0.00           N  
ATOM    622  H   ARG A  39      -6.390  54.921 -30.306  1.00  0.00           H  
ATOM    623  HA  ARG A  39      -6.739  54.873 -33.185  1.00  0.00           H  
ATOM    624 1HB  ARG A  39      -4.203  54.752 -31.530  1.00  0.00           H  
ATOM    625 2HB  ARG A  39      -4.270  54.948 -33.274  1.00  0.00           H  
ATOM    626 1HG  ARG A  39      -5.542  56.896 -31.342  1.00  0.00           H  
ATOM    627 2HG  ARG A  39      -3.883  57.049 -31.969  1.00  0.00           H  
ATOM    628 1HD  ARG A  39      -4.783  57.054 -34.249  1.00  0.00           H  
ATOM    629 2HD  ARG A  39      -6.427  57.003 -33.573  1.00  0.00           H  
ATOM    630  HE  ARG A  39      -5.039  59.166 -32.405  1.00  0.00           H  
ATOM    631 1HH1 ARG A  39      -6.335  58.167 -35.513  1.00  0.00           H  
ATOM    632 2HH1 ARG A  39      -6.622  59.803 -36.061  1.00  0.00           H  
ATOM    633 1HH2 ARG A  39      -5.416  61.286 -33.127  1.00  0.00           H  
ATOM    634 2HH2 ARG A  39      -6.102  61.570 -34.711  1.00  0.00           H  
ATOM    635  N   MET A  40      -5.859  52.483 -33.779  1.00  0.00           N  
ATOM    636  CA  MET A  40      -5.725  51.061 -34.081  1.00  0.00           C  
ATOM    637  C   MET A  40      -4.487  50.407 -33.485  1.00  0.00           C  
ATOM    638  O   MET A  40      -4.446  49.189 -33.313  1.00  0.00           O  
ATOM    639  CB  MET A  40      -5.729  50.853 -35.597  1.00  0.00           C  
ATOM    640  CG  MET A  40      -4.555  51.506 -36.310  1.00  0.00           C  
ATOM    641  SD  MET A  40      -4.750  51.484 -38.103  1.00  0.00           S  
ATOM    642  CE  MET A  40      -3.385  52.550 -38.605  1.00  0.00           C  
ATOM    643  H   MET A  40      -5.867  53.148 -34.539  1.00  0.00           H  
ATOM    644  HA  MET A  40      -6.575  50.534 -33.648  1.00  0.00           H  
ATOM    645 1HB  MET A  40      -5.710  49.787 -35.819  1.00  0.00           H  
ATOM    646 2HB  MET A  40      -6.650  51.258 -36.018  1.00  0.00           H  
ATOM    647 1HG  MET A  40      -4.460  52.541 -35.981  1.00  0.00           H  
ATOM    648 2HG  MET A  40      -3.635  50.981 -36.051  1.00  0.00           H  
ATOM    649 1HE  MET A  40      -3.371  52.636 -39.692  1.00  0.00           H  
ATOM    650 2HE  MET A  40      -3.514  53.539 -38.164  1.00  0.00           H  
ATOM    651 3HE  MET A  40      -2.444  52.120 -38.263  1.00  0.00           H  
ATOM    652  N   ASP A  41      -3.479  51.218 -33.178  1.00  0.00           N  
ATOM    653  CA  ASP A  41      -2.235  50.717 -32.614  1.00  0.00           C  
ATOM    654  C   ASP A  41      -1.959  51.321 -31.245  1.00  0.00           C  
ATOM    655  O   ASP A  41      -0.804  51.442 -30.830  1.00  0.00           O  
ATOM    656  CB  ASP A  41      -1.079  51.022 -33.571  1.00  0.00           C  
ATOM    657  CG  ASP A  41      -0.914  52.507 -33.844  1.00  0.00           C  
ATOM    658  OD1 ASP A  41      -1.858  53.285 -33.583  1.00  0.00           O  
ATOM    659  OD2 ASP A  41       0.167  52.895 -34.335  1.00  0.00           O1-
ATOM    660  H   ASP A  41      -3.582  52.209 -33.340  1.00  0.00           H  
ATOM    661  HA  ASP A  41      -2.321  49.637 -32.491  1.00  0.00           H  
ATOM    662 1HB  ASP A  41      -0.148  50.638 -33.151  1.00  0.00           H  
ATOM    663 2HB  ASP A  41      -1.246  50.509 -34.519  1.00  0.00           H  
ATOM    664  N   ARG A  42      -3.025  51.680 -30.538  1.00  0.00           N  
ATOM    665  CA  ARG A  42      -2.884  52.287 -29.221  1.00  0.00           C  
ATOM    666  C   ARG A  42      -2.012  51.449 -28.299  1.00  0.00           C  
ATOM    667  O   ARG A  42      -2.080  50.217 -28.295  1.00  0.00           O  
ATOM    668  CB  ARG A  42      -4.249  52.480 -28.577  1.00  0.00           C  
ATOM    669  CG  ARG A  42      -4.222  53.131 -27.203  1.00  0.00           C  
ATOM    670  CD  ARG A  42      -3.933  54.586 -27.290  1.00  0.00           C  
ATOM    671  NE  ARG A  42      -3.964  55.224 -25.984  1.00  0.00           N  
ATOM    672  CZ  ARG A  42      -3.702  56.527 -25.763  1.00  0.00           C  
ATOM    673  NH1 ARG A  42      -3.392  57.315 -26.768  1.00  0.00           N1+
ATOM    674  NH2 ARG A  42      -3.756  57.013 -24.535  1.00  0.00           N  
ATOM    675  H   ARG A  42      -3.950  51.532 -30.914  1.00  0.00           H  
ATOM    676  HA  ARG A  42      -2.396  53.255 -29.338  1.00  0.00           H  
ATOM    677 1HB  ARG A  42      -4.871  53.099 -29.222  1.00  0.00           H  
ATOM    678 2HB  ARG A  42      -4.743  51.514 -28.474  1.00  0.00           H  
ATOM    679 1HG  ARG A  42      -5.190  53.002 -26.720  1.00  0.00           H  
ATOM    680 2HG  ARG A  42      -3.447  52.664 -26.594  1.00  0.00           H  
ATOM    681 1HD  ARG A  42      -2.943  54.735 -27.719  1.00  0.00           H  
ATOM    682 2HD  ARG A  42      -4.678  55.067 -27.923  1.00  0.00           H  
ATOM    683  HE  ARG A  42      -4.199  54.650 -25.186  1.00  0.00           H  
ATOM    684 1HH1 ARG A  42      -3.350  56.944 -27.707  1.00  0.00           H  
ATOM    685 2HH1 ARG A  42      -3.195  58.291 -26.603  1.00  0.00           H  
ATOM    686 1HH2 ARG A  42      -3.995  56.407 -23.762  1.00  0.00           H  
ATOM    687 2HH2 ARG A  42      -3.560  57.989 -24.369  1.00  0.00           H  
ATOM    688  N   ALA A  43      -1.186  52.132 -27.516  1.00  0.00           N  
ATOM    689  CA  ALA A  43      -0.291  51.465 -26.586  1.00  0.00           C  
ATOM    690  C   ALA A  43      -1.053  50.872 -25.414  1.00  0.00           C  
ATOM    691  O   ALA A  43      -2.043  51.435 -24.951  1.00  0.00           O  
ATOM    692  CB  ALA A  43       0.758  52.451 -26.081  1.00  0.00           C  
ATOM    693  H   ALA A  43      -1.179  53.141 -27.567  1.00  0.00           H  
ATOM    694  HA  ALA A  43       0.207  50.654 -27.119  1.00  0.00           H  
ATOM    695 1HB  ALA A  43       1.427  51.946 -25.385  1.00  0.00           H  
ATOM    696 2HB  ALA A  43       1.333  52.834 -26.924  1.00  0.00           H  
ATOM    697 3HB  ALA A  43       0.264  53.278 -25.574  1.00  0.00           H  
ATOM    698  N   ILE A  44      -0.587  49.722 -24.943  1.00  0.00           N  
ATOM    699  CA  ILE A  44      -1.192  49.050 -23.803  1.00  0.00           C  
ATOM    700  C   ILE A  44      -0.081  48.674 -22.827  1.00  0.00           C  
ATOM    701  O   ILE A  44       1.074  48.498 -23.225  1.00  0.00           O  
ATOM    702  CB  ILE A  44      -1.975  47.795 -24.232  1.00  0.00           C  
ATOM    703  CG1 ILE A  44      -1.038  46.783 -24.897  1.00  0.00           C  
ATOM    704  CG2 ILE A  44      -3.110  48.172 -25.172  1.00  0.00           C  
ATOM    705  CD1 ILE A  44      -1.684  45.446 -25.178  1.00  0.00           C  
ATOM    706  H   ILE A  44       0.213  49.302 -25.393  1.00  0.00           H  
ATOM    707  HA  ILE A  44      -1.901  49.732 -23.335  1.00  0.00           H  
ATOM    708  HB  ILE A  44      -2.391  47.308 -23.352  1.00  0.00           H  
ATOM    709 1HG1 ILE A  44      -0.672  47.191 -25.839  1.00  0.00           H  
ATOM    710 2HG1 ILE A  44      -0.171  46.614 -24.257  1.00  0.00           H  
ATOM    711 1HG2 ILE A  44      -3.652  47.274 -25.466  1.00  0.00           H  
ATOM    712 2HG2 ILE A  44      -3.789  48.856 -24.665  1.00  0.00           H  
ATOM    713 3HG2 ILE A  44      -2.702  48.657 -26.059  1.00  0.00           H  
ATOM    714 1HD1 ILE A  44      -0.958  44.783 -25.649  1.00  0.00           H  
ATOM    715 2HD1 ILE A  44      -2.026  45.003 -24.242  1.00  0.00           H  
ATOM    716 3HD1 ILE A  44      -2.533  45.586 -25.846  1.00  0.00           H  
ATOM    717  N   PRO A  45      -0.410  48.561 -21.534  1.00  0.00           N  
ATOM    718  CA  PRO A  45       0.595  48.207 -20.530  1.00  0.00           C  
ATOM    719  C   PRO A  45       1.117  46.779 -20.661  1.00  0.00           C  
ATOM    720  O   PRO A  45       0.381  45.877 -21.059  1.00  0.00           O  
ATOM    721  CB  PRO A  45      -0.172  48.396 -19.217  1.00  0.00           C  
ATOM    722  CG  PRO A  45      -1.601  48.214 -19.599  1.00  0.00           C  
ATOM    723  CD  PRO A  45      -1.688  48.768 -20.996  1.00  0.00           C  
ATOM    724  HA  PRO A  45       1.452  48.891 -20.620  1.00  0.00           H  
ATOM    725 1HB  PRO A  45       0.168  47.660 -18.474  1.00  0.00           H  
ATOM    726 2HB  PRO A  45       0.036  49.393 -18.801  1.00  0.00           H  
ATOM    727 1HG  PRO A  45      -1.876  47.151 -19.547  1.00  0.00           H  
ATOM    728 2HG  PRO A  45      -2.254  48.746 -18.891  1.00  0.00           H  
ATOM    729 1HD  PRO A  45      -2.441  48.205 -21.568  1.00  0.00           H  
ATOM    730 2HD  PRO A  45      -1.952  49.835 -20.953  1.00  0.00           H  
ATOM    731  N   ASP A  46       2.391  46.583 -20.333  1.00  0.00           N  
ATOM    732  CA  ASP A  46       2.997  45.254 -20.373  1.00  0.00           C  
ATOM    733  C   ASP A  46       2.592  44.601 -19.054  1.00  0.00           C  
ATOM    734  O   ASP A  46       3.130  44.918 -17.992  1.00  0.00           O  
ATOM    735  CB  ASP A  46       4.524  45.362 -20.466  1.00  0.00           C  
ATOM    736  CG  ASP A  46       5.200  44.014 -20.690  1.00  0.00           C  
ATOM    737  OD1 ASP A  46       4.559  42.970 -20.449  1.00  0.00           O  
ATOM    738  OD2 ASP A  46       6.381  44.002 -21.099  1.00  0.00           O1-
ATOM    739  H   ASP A  46       2.953  47.372 -20.050  1.00  0.00           H  
ATOM    740  HA  ASP A  46       2.631  44.733 -21.258  1.00  0.00           H  
ATOM    741 1HB  ASP A  46       4.793  46.029 -21.286  1.00  0.00           H  
ATOM    742 2HB  ASP A  46       4.915  45.799 -19.546  1.00  0.00           H  
ATOM    743  N   THR A  47       1.635  43.686 -19.134  1.00  0.00           N  
ATOM    744  CA  THR A  47       1.104  43.003 -17.964  1.00  0.00           C  
ATOM    745  C   THR A  47       1.839  41.712 -17.594  1.00  0.00           C  
ATOM    746  O   THR A  47       1.442  41.019 -16.657  1.00  0.00           O  
ATOM    747  CB  THR A  47      -0.389  42.687 -18.172  1.00  0.00           C  
ATOM    748  CG2 THR A  47      -1.181  43.967 -18.385  1.00  0.00           C  
ATOM    749  OG1 THR A  47      -0.545  41.837 -19.316  1.00  0.00           O  
ATOM    750  H   THR A  47       1.266  43.459 -20.046  1.00  0.00           H  
ATOM    751  HA  THR A  47       1.193  43.668 -17.105  1.00  0.00           H  
ATOM    752  HB  THR A  47      -0.776  42.168 -17.295  1.00  0.00           H  
ATOM    753  HG1 THR A  47      -1.476  41.641 -19.443  1.00  0.00           H  
ATOM    754 1HG2 THR A  47      -2.233  43.724 -18.530  1.00  0.00           H  
ATOM    755 2HG2 THR A  47      -1.073  44.611 -17.512  1.00  0.00           H  
ATOM    756 3HG2 THR A  47      -0.804  44.486 -19.266  1.00  0.00           H  
ATOM    757  N   ARG A  48       2.903  41.397 -18.326  1.00  0.00           N  
ATOM    758  CA  ARG A  48       3.677  40.185 -18.069  1.00  0.00           C  
ATOM    759  C   ARG A  48       4.488  40.239 -16.783  1.00  0.00           C  
ATOM    760  O   ARG A  48       4.896  41.307 -16.327  1.00  0.00           O  
ATOM    761  CB  ARG A  48       4.651  39.912 -19.215  1.00  0.00           C  
ATOM    762  CG  ARG A  48       4.029  39.604 -20.560  1.00  0.00           C  
ATOM    763  CD  ARG A  48       5.130  39.450 -21.591  1.00  0.00           C  
ATOM    764  NE  ARG A  48       5.941  40.663 -21.670  1.00  0.00           N  
ATOM    765  CZ  ARG A  48       7.122  40.739 -22.274  1.00  0.00           C  
ATOM    766  NH1 ARG A  48       7.786  41.892 -22.295  1.00  0.00           N1+
ATOM    767  NH2 ARG A  48       7.647  39.663 -22.845  1.00  0.00           N  
ATOM    768  H   ARG A  48       3.183  42.010 -19.078  1.00  0.00           H  
ATOM    769  HA  ARG A  48       2.988  39.344 -17.992  1.00  0.00           H  
ATOM    770 1HB  ARG A  48       5.300  40.775 -19.355  1.00  0.00           H  
ATOM    771 2HB  ARG A  48       5.287  39.064 -18.957  1.00  0.00           H  
ATOM    772 1HG  ARG A  48       3.457  38.677 -20.493  1.00  0.00           H  
ATOM    773 2HG  ARG A  48       3.366  40.420 -20.850  1.00  0.00           H  
ATOM    774 1HD  ARG A  48       5.774  38.615 -21.315  1.00  0.00           H  
ATOM    775 2HD  ARG A  48       4.689  39.259 -22.569  1.00  0.00           H  
ATOM    776  HE  ARG A  48       5.577  41.500 -21.235  1.00  0.00           H  
ATOM    777 1HH1 ARG A  48       7.392  42.709 -21.851  1.00  0.00           H  
ATOM    778 2HH1 ARG A  48       8.683  41.949 -22.753  1.00  0.00           H  
ATOM    779 1HH2 ARG A  48       7.148  38.785 -22.822  1.00  0.00           H  
ATOM    780 2HH2 ARG A  48       8.544  39.723 -23.304  1.00  0.00           H  
ATOM    781  N   HIS A  49       4.731  39.061 -16.222  1.00  0.00           N  
ATOM    782  CA  HIS A  49       5.532  38.914 -15.014  1.00  0.00           C  
ATOM    783  C   HIS A  49       6.957  39.305 -15.413  1.00  0.00           C  
ATOM    784  O   HIS A  49       7.355  39.103 -16.560  1.00  0.00           O  
ATOM    785  CB  HIS A  49       5.484  37.486 -14.460  1.00  0.00           C  
ATOM    786  CG  HIS A  49       6.133  37.340 -13.119  1.00  0.00           C  
ATOM    787  CD2 HIS A  49       5.604  37.266 -11.875  1.00  0.00           C  
ATOM    788  ND1 HIS A  49       7.500  37.250 -12.960  1.00  0.00           N  
ATOM    789  CE1 HIS A  49       7.784  37.128 -11.675  1.00  0.00           C  
ATOM    790  NE2 HIS A  49       6.651  37.135 -10.996  1.00  0.00           N  
ATOM    791  H   HIS A  49       4.340  38.237 -16.656  1.00  0.00           H  
ATOM    792  HA  HIS A  49       5.119  39.543 -14.226  1.00  0.00           H  
ATOM    793 1HB  HIS A  49       4.446  37.164 -14.373  1.00  0.00           H  
ATOM    794 2HB  HIS A  49       5.980  36.810 -15.155  1.00  0.00           H  
ATOM    795  HD2 HIS A  49       4.545  37.303 -11.619  1.00  0.00           H  
ATOM    796  HE1 HIS A  49       8.783  37.038 -11.248  1.00  0.00           H  
ATOM    797  HE2 HIS A  49       6.563  37.057  -9.993  1.00  0.00           H  
ATOM    798  N   ASP A  50       7.723  39.867 -14.481  1.00  0.00           N  
ATOM    799  CA  ASP A  50       9.089  40.279 -14.791  1.00  0.00           C  
ATOM    800  C   ASP A  50       9.969  39.167 -15.358  1.00  0.00           C  
ATOM    801  O   ASP A  50      10.847  39.427 -16.183  1.00  0.00           O  
ATOM    802  CB  ASP A  50       9.752  40.845 -13.533  1.00  0.00           C  
ATOM    803  CG  ASP A  50       9.200  42.206 -13.133  1.00  0.00           C  
ATOM    804  OD1 ASP A  50       8.525  42.810 -13.933  1.00  0.00           O  
ATOM    805  OD2 ASP A  50       9.458  42.630 -12.032  1.00  0.00           O1-
ATOM    806  H   ASP A  50       7.364  40.014 -13.549  1.00  0.00           H  
ATOM    807  HA  ASP A  50       9.052  41.051 -15.559  1.00  0.00           H  
ATOM    808 1HB  ASP A  50       9.608  40.153 -12.703  1.00  0.00           H  
ATOM    809 2HB  ASP A  50      10.826  40.939 -13.698  1.00  0.00           H  
ATOM    810  N   GLN A  51       9.741  37.933 -14.924  1.00  0.00           N  
ATOM    811  CA  GLN A  51      10.558  36.820 -15.402  1.00  0.00           C  
ATOM    812  C   GLN A  51      10.358  36.490 -16.875  1.00  0.00           C  
ATOM    813  O   GLN A  51      11.170  35.782 -17.467  1.00  0.00           O  
ATOM    814  CB  GLN A  51      10.271  35.571 -14.564  1.00  0.00           C  
ATOM    815  CG  GLN A  51      10.735  35.671 -13.120  1.00  0.00           C  
ATOM    816  CD  GLN A  51      10.404  34.426 -12.319  1.00  0.00           C  
ATOM    817  NE2 GLN A  51      10.160  34.604 -11.026  1.00  0.00           N  
ATOM    818  OE1 GLN A  51      10.367  33.316 -12.857  1.00  0.00           O  
ATOM    819  H   GLN A  51       9.001  37.751 -14.260  1.00  0.00           H  
ATOM    820  HA  GLN A  51      11.608  37.094 -15.303  1.00  0.00           H  
ATOM    821 1HB  GLN A  51       9.199  35.374 -14.560  1.00  0.00           H  
ATOM    822 2HB  GLN A  51      10.761  34.709 -15.016  1.00  0.00           H  
ATOM    823 1HG  GLN A  51      11.816  35.809 -13.106  1.00  0.00           H  
ATOM    824 2HG  GLN A  51      10.243  36.522 -12.649  1.00  0.00           H  
ATOM    825 1HE2 GLN A  51       9.935  33.819 -10.446  1.00  0.00           H  
ATOM    826 2HE2 GLN A  51      10.200  35.522 -10.630  1.00  0.00           H  
ATOM    827  N   CYS A  52       9.291  37.009 -17.471  1.00 17.94           N  
ATOM    828  CA  CYS A  52       9.004  36.736 -18.874  1.00 19.31           C  
ATOM    829  C   CYS A  52      10.030  37.287 -19.845  1.00 21.63           C  
ATOM    830  O   CYS A  52      10.329  36.668 -20.866  1.00 22.43           O  
ATOM    831  CB  CYS A  52       7.631  37.287 -19.247  1.00 17.59           C  
ATOM    832  SG  CYS A  52       6.291  36.372 -18.444  1.00 13.62           S  
ATOM    833  H   CYS A  52       8.664  37.603 -16.948  1.00  0.00           H  
ATOM    834  HA  CYS A  52       9.001  35.657 -19.024  1.00  0.00           H  
ATOM    835 1HB  CYS A  52       7.570  38.337 -18.958  1.00  0.00           H  
ATOM    836 2HB  CYS A  52       7.501  37.237 -20.328  1.00  0.00           H  
ATOM    837  N   GLN A  53      10.567  38.454 -19.528  1.00  0.00           N  
ATOM    838  CA  GLN A  53      11.545  39.079 -20.397  1.00  0.00           C  
ATOM    839  C   GLN A  53      12.865  38.315 -20.449  1.00  0.00           C  
ATOM    840  O   GLN A  53      13.742  38.635 -21.250  1.00  0.00           O  
ATOM    841  CB  GLN A  53      11.799  40.519 -19.943  1.00  0.00           C  
ATOM    842  CG  GLN A  53      10.607  41.445 -20.114  1.00  0.00           C  
ATOM    843  CD  GLN A  53      10.892  42.852 -19.623  1.00  0.00           C  
ATOM    844  NE2 GLN A  53       9.860  43.525 -19.128  1.00  0.00           N  
ATOM    845  OE1 GLN A  53      12.028  43.329 -19.687  1.00  0.00           O  
ATOM    846  H   GLN A  53      10.298  38.921 -18.673  1.00  0.00           H  
ATOM    847  HA  GLN A  53      11.159  39.067 -21.417  1.00  0.00           H  
ATOM    848 1HB  GLN A  53      12.080  40.524 -18.890  1.00  0.00           H  
ATOM    849 2HB  GLN A  53      12.633  40.936 -20.506  1.00  0.00           H  
ATOM    850 1HG  GLN A  53      10.351  41.498 -21.172  1.00  0.00           H  
ATOM    851 2HG  GLN A  53       9.767  41.048 -19.546  1.00  0.00           H  
ATOM    852 1HE2 GLN A  53       9.986  44.458 -18.788  1.00  0.00           H  
ATOM    853 2HE2 GLN A  53       8.955  43.100 -19.094  1.00  0.00           H  
ATOM    854  N   ARG A  54      12.993  37.290 -19.611  1.00  0.00           N  
ATOM    855  CA  ARG A  54      14.209  36.488 -19.578  1.00  0.00           C  
ATOM    856  C   ARG A  54      13.983  35.063 -20.077  1.00  0.00           C  
ATOM    857  O   ARG A  54      14.942  34.325 -20.313  1.00  0.00           O  
ATOM    858  CB  ARG A  54      14.766  36.437 -18.163  1.00  0.00           C  
ATOM    859  CG  ARG A  54      15.221  37.777 -17.606  1.00  0.00           C  
ATOM    860  CD  ARG A  54      15.778  37.642 -16.236  1.00  0.00           C  
ATOM    861  NE  ARG A  54      16.212  38.921 -15.698  1.00  0.00           N  
ATOM    862  CZ  ARG A  54      16.781  39.089 -14.488  1.00  0.00           C  
ATOM    863  NH1 ARG A  54      16.976  38.052 -13.704  1.00  0.00           N1+
ATOM    864  NH2 ARG A  54      17.142  40.296 -14.090  1.00  0.00           N  
ATOM    865  H   ARG A  54      12.235  37.060 -18.984  1.00  0.00           H  
ATOM    866  HA  ARG A  54      14.943  36.946 -20.242  1.00  0.00           H  
ATOM    867 1HB  ARG A  54      14.008  36.040 -17.488  1.00  0.00           H  
ATOM    868 2HB  ARG A  54      15.619  35.760 -18.132  1.00  0.00           H  
ATOM    869 1HG  ARG A  54      15.994  38.195 -18.251  1.00  0.00           H  
ATOM    870 2HG  ARG A  54      14.373  38.461 -17.566  1.00  0.00           H  
ATOM    871 1HD  ARG A  54      15.017  37.234 -15.572  1.00  0.00           H  
ATOM    872 2HD  ARG A  54      16.637  36.972 -16.257  1.00  0.00           H  
ATOM    873  HE  ARG A  54      16.078  39.742 -16.273  1.00  0.00           H  
ATOM    874 1HH1 ARG A  54      16.700  37.129 -14.009  1.00  0.00           H  
ATOM    875 2HH1 ARG A  54      17.402  38.178 -12.798  1.00  0.00           H  
ATOM    876 1HH2 ARG A  54      16.991  41.094 -14.693  1.00  0.00           H  
ATOM    877 2HH2 ARG A  54      17.567  40.422 -13.184  1.00  0.00           H  
ATOM    878  N   LYS A  55      12.721  34.673 -20.231  1.00  0.00           N  
ATOM    879  CA  LYS A  55      12.399  33.336 -20.721  1.00  0.00           C  
ATOM    880  C   LYS A  55      12.671  33.270 -22.218  1.00  0.00           C  
ATOM    881  O   LYS A  55      12.381  34.216 -22.947  1.00  0.00           O  
ATOM    882  CB  LYS A  55      10.942  32.977 -20.423  1.00  0.00           C  
ATOM    883  CG  LYS A  55      10.554  31.555 -20.806  1.00  0.00           C  
ATOM    884  CD  LYS A  55       9.124  31.242 -20.391  1.00  0.00           C  
ATOM    885  CE  LYS A  55       8.735  29.821 -20.771  1.00  0.00           C  
ATOM    886  NZ  LYS A  55       7.338  29.500 -20.371  1.00  0.00           N1+
ATOM    887  H   LYS A  55      11.969  35.309 -20.008  1.00  0.00           H  
ATOM    888  HA  LYS A  55      13.028  32.615 -20.197  1.00  0.00           H  
ATOM    889 1HB  LYS A  55      10.747  33.102 -19.358  1.00  0.00           H  
ATOM    890 2HB  LYS A  55      10.282  33.660 -20.959  1.00  0.00           H  
ATOM    891 1HG  LYS A  55      10.646  31.431 -21.886  1.00  0.00           H  
ATOM    892 2HG  LYS A  55      11.227  30.850 -20.319  1.00  0.00           H  
ATOM    893 1HD  LYS A  55       9.024  31.361 -19.311  1.00  0.00           H  
ATOM    894 2HD  LYS A  55       8.443  31.938 -20.880  1.00  0.00           H  
ATOM    895 1HE  LYS A  55       8.830  29.694 -21.849  1.00  0.00           H  
ATOM    896 2HE  LYS A  55       9.410  29.116 -20.284  1.00  0.00           H  
ATOM    897 1HZ  LYS A  55       7.120  28.551 -20.640  1.00  0.00           H  
ATOM    898 2HZ  LYS A  55       7.244  29.597 -19.370  1.00  0.00           H  
ATOM    899 3HZ  LYS A  55       6.702  30.135 -20.832  1.00  0.00           H  
ATOM    900  N   GLN A  56      13.229  32.152 -22.674  1.00  0.00           N  
ATOM    901  CA  GLN A  56      13.554  32.000 -24.087  1.00  0.00           C  
ATOM    902  C   GLN A  56      12.724  30.952 -24.819  1.00  0.00           C  
ATOM    903  O   GLN A  56      12.028  30.141 -24.208  1.00  0.00           O  
ATOM    904  CB  GLN A  56      15.038  31.659 -24.250  1.00  0.00           C  
ATOM    905  CG  GLN A  56      15.995  32.649 -23.596  1.00  0.00           C  
ATOM    906  CD  GLN A  56      15.762  34.079 -24.048  1.00  0.00           C  
ATOM    907  NE2 GLN A  56      15.653  34.991 -23.089  1.00  0.00           N  
ATOM    908  OE1 GLN A  56      15.691  34.362 -25.245  1.00  0.00           O  
ATOM    909  H   GLN A  56      13.434  31.394 -22.038  1.00  0.00           H  
ATOM    910  HA  GLN A  56      13.351  32.944 -24.591  1.00  0.00           H  
ATOM    911 1HB  GLN A  56      15.235  30.677 -23.823  1.00  0.00           H  
ATOM    912 2HB  GLN A  56      15.286  31.612 -25.310  1.00  0.00           H  
ATOM    913 1HG  GLN A  56      15.858  32.609 -22.515  1.00  0.00           H  
ATOM    914 2HG  GLN A  56      17.018  32.374 -23.853  1.00  0.00           H  
ATOM    915 1HE2 GLN A  56      15.499  35.951 -23.326  1.00  0.00           H  
ATOM    916 2HE2 GLN A  56      15.724  34.720 -22.129  1.00  0.00           H  
ATOM    917  N   TRP A  57      12.814  30.991 -26.143  1.00  0.00           N  
ATOM    918  CA  TRP A  57      12.115  30.064 -27.023  1.00  0.00           C  
ATOM    919  C   TRP A  57      13.030  29.787 -28.211  1.00  0.00           C  
ATOM    920  O   TRP A  57      13.959  30.551 -28.467  1.00  0.00           O  
ATOM    921  CB  TRP A  57      10.786  30.669 -27.489  1.00  0.00           C  
ATOM    922  CG  TRP A  57      10.900  32.080 -27.991  1.00  0.00           C  
ATOM    923  CD1 TRP A  57      11.427  32.492 -29.181  1.00  0.00           C  
ATOM    924  CD2 TRP A  57      10.499  33.268 -27.297  1.00  0.00           C  
ATOM    925  CE2 TRP A  57      10.817  34.365 -28.128  1.00  0.00           C  
ATOM    926  CE3 TRP A  57       9.904  33.511 -26.049  1.00  0.00           C  
ATOM    927  NE1 TRP A  57      11.381  33.864 -29.273  1.00  0.00           N  
ATOM    928  CZ2 TRP A  57      10.557  35.688 -27.756  1.00  0.00           C  
ATOM    929  CZ3 TRP A  57       9.647  34.828 -25.678  1.00  0.00           C  
ATOM    930  CH2 TRP A  57       9.973  35.899 -26.530  1.00  0.00           C  
ATOM    931  H   TRP A  57      13.400  31.706 -26.548  1.00  0.00           H  
ATOM    932  HA  TRP A  57      11.907  29.150 -26.466  1.00  0.00           H  
ATOM    933 1HB  TRP A  57      10.369  30.058 -28.290  1.00  0.00           H  
ATOM    934 2HB  TRP A  57      10.073  30.660 -26.665  1.00  0.00           H  
ATOM    935  HD1 TRP A  57      11.826  31.830 -29.948  1.00  0.00           H  
ATOM    936  HE1 TRP A  57      11.710  34.412 -30.054  1.00  0.00           H  
ATOM    937  HE3 TRP A  57       9.648  32.685 -25.386  1.00  0.00           H  
ATOM    938  HZ2 TRP A  57      10.799  36.531 -28.403  1.00  0.00           H  
ATOM    939  HZ3 TRP A  57       9.184  35.008 -24.707  1.00  0.00           H  
ATOM    940  HH2 TRP A  57       9.759  36.918 -26.207  1.00  0.00           H  
ATOM    941  N   ARG A  58      12.777  28.698 -28.928  1.00  0.00           N  
ATOM    942  CA  ARG A  58      13.613  28.345 -30.069  1.00  0.00           C  
ATOM    943  C   ARG A  58      13.489  29.368 -31.187  1.00  0.00           C  
ATOM    944  O   ARG A  58      12.473  30.052 -31.312  1.00  0.00           O  
ATOM    945  CB  ARG A  58      13.233  26.959 -30.589  1.00  0.00           C  
ATOM    946  CG  ARG A  58      11.901  26.907 -31.299  1.00  0.00           C  
ATOM    947  CD  ARG A  58      11.428  25.475 -31.388  1.00  0.00           C  
ATOM    948  NE  ARG A  58      11.194  24.936 -30.053  1.00  0.00           N  
ATOM    949  CZ  ARG A  58      11.044  23.644 -29.780  1.00  0.00           C  
ATOM    950  NH1 ARG A  58      11.104  22.745 -30.752  1.00  0.00           N1+
ATOM    951  NH2 ARG A  58      10.837  23.254 -28.529  1.00  0.00           N  
ATOM    952  H   ARG A  58      11.999  28.103 -28.685  1.00  0.00           H  
ATOM    953  HA  ARG A  58      14.653  28.326 -29.743  1.00  0.00           H  
ATOM    954 1HB  ARG A  58      13.998  26.608 -31.280  1.00  0.00           H  
ATOM    955 2HB  ARG A  58      13.199  26.255 -29.757  1.00  0.00           H  
ATOM    956 1HG  ARG A  58      11.170  27.496 -30.745  1.00  0.00           H  
ATOM    957 2HG  ARG A  58      12.009  27.316 -32.304  1.00  0.00           H  
ATOM    958 1HD  ARG A  58      10.499  25.433 -31.956  1.00  0.00           H  
ATOM    959 2HD  ARG A  58      12.186  24.873 -31.888  1.00  0.00           H  
ATOM    960  HE  ARG A  58      11.142  25.590 -29.284  1.00  0.00           H  
ATOM    961 1HH1 ARG A  58      11.263  23.041 -31.705  1.00  0.00           H  
ATOM    962 2HH1 ARG A  58      10.989  21.764 -30.541  1.00  0.00           H  
ATOM    963 1HH2 ARG A  58      10.793  23.939 -27.787  1.00  0.00           H  
ATOM    964 2HH2 ARG A  58      10.722  22.275 -28.319  1.00  0.00           H  
ATOM    965  N   VAL A  59      14.533  29.470 -32.001  1.00  0.00           N  
ATOM    966  CA  VAL A  59      14.534  30.416 -33.107  1.00  0.00           C  
ATOM    967  C   VAL A  59      13.666  29.922 -34.257  1.00  0.00           C  
ATOM    968  O   VAL A  59      12.963  30.706 -34.893  1.00  0.00           O  
ATOM    969  CB  VAL A  59      15.971  30.644 -33.612  1.00  0.00           C  
ATOM    970  CG1 VAL A  59      15.965  31.514 -34.860  1.00  0.00           C  
ATOM    971  CG2 VAL A  59      16.811  31.279 -32.515  1.00  0.00           C  
ATOM    972  H   VAL A  59      15.342  28.883 -31.855  1.00  0.00           H  
ATOM    973  HA  VAL A  59      14.135  31.366 -32.750  1.00  0.00           H  
ATOM    974  HB  VAL A  59      16.404  29.683 -33.893  1.00  0.00           H  
ATOM    975 1HG1 VAL A  59      16.989  31.664 -35.204  1.00  0.00           H  
ATOM    976 2HG1 VAL A  59      15.388  31.023 -35.644  1.00  0.00           H  
ATOM    977 3HG1 VAL A  59      15.516  32.479 -34.629  1.00  0.00           H  
ATOM    978 1HG2 VAL A  59      17.826  31.437 -32.879  1.00  0.00           H  
ATOM    979 2HG2 VAL A  59      16.375  32.237 -32.231  1.00  0.00           H  
ATOM    980 3HG2 VAL A  59      16.835  30.620 -31.647  1.00  0.00           H  
ATOM    981  N   ASP A  60      13.709  28.618 -34.513  1.00  0.00           N  
ATOM    982  CA  ASP A  60      12.935  28.035 -35.599  1.00  0.00           C  
ATOM    983  C   ASP A  60      11.498  27.698 -35.222  1.00  0.00           C  
ATOM    984  O   ASP A  60      11.077  26.545 -35.319  1.00  0.00           O  
ATOM    985  CB  ASP A  60      13.626  26.773 -36.139  1.00  0.00           C  
ATOM    986  CG  ASP A  60      13.815  25.701 -35.077  1.00  0.00           C  
ATOM    987  OD1 ASP A  60      13.748  24.501 -35.427  1.00  0.00           O  
ATOM    988  OD2 ASP A  60      14.046  26.050 -33.900  1.00  0.00           O1-
ATOM    989  H   ASP A  60      14.289  28.017 -33.945  1.00  0.00           H  
ATOM    990  HA  ASP A  60      12.867  28.766 -36.406  1.00  0.00           H  
ATOM    991 1HB  ASP A  60      13.034  26.354 -36.954  1.00  0.00           H  
ATOM    992 2HB  ASP A  60      14.602  27.038 -36.545  1.00  0.00           H  
ATOM    993  N   LEU A  61      10.741  28.703 -34.796  1.00  0.00           N  
ATOM    994  CA  LEU A  61       9.345  28.484 -34.438  1.00  0.00           C  
ATOM    995  C   LEU A  61       8.514  28.422 -35.714  1.00  0.00           C  
ATOM    996  O   LEU A  61       8.816  29.105 -36.693  1.00  0.00           O  
ATOM    997  CB  LEU A  61       8.832  29.620 -33.550  1.00  0.00           C  
ATOM    998  CG  LEU A  61       9.378  29.656 -32.120  1.00  0.00           C  
ATOM    999  CD1 LEU A  61       8.888  30.904 -31.410  1.00  0.00           C  
ATOM   1000  CD2 LEU A  61       8.922  28.403 -31.381  1.00  0.00           C  
ATOM   1001  H   LEU A  61      11.130  29.632 -34.716  1.00  0.00           H  
ATOM   1002  HA  LEU A  61       9.274  27.549 -33.883  1.00  0.00           H  
ATOM   1003 1HB  LEU A  61       9.083  30.570 -34.020  1.00  0.00           H  
ATOM   1004 2HB  LEU A  61       7.746  29.546 -33.484  1.00  0.00           H  
ATOM   1005  HG  LEU A  61      10.468  29.691 -32.148  1.00  0.00           H  
ATOM   1006 1HD1 LEU A  61       9.281  30.922 -30.393  1.00  0.00           H  
ATOM   1007 2HD1 LEU A  61       9.232  31.787 -31.947  1.00  0.00           H  
ATOM   1008 3HD1 LEU A  61       7.799  30.900 -31.377  1.00  0.00           H  
ATOM   1009 1HD2 LEU A  61       9.308  28.421 -30.362  1.00  0.00           H  
ATOM   1010 2HD2 LEU A  61       7.833  28.371 -31.356  1.00  0.00           H  
ATOM   1011 3HD2 LEU A  61       9.299  27.519 -31.896  1.00  0.00           H  
ATOM   1012  N   PRO A  62       7.456  27.598 -35.721  1.00  0.00           N  
ATOM   1013  CA  PRO A  62       6.597  27.473 -36.901  1.00  0.00           C  
ATOM   1014  C   PRO A  62       5.684  28.688 -37.047  1.00  0.00           C  
ATOM   1015  O   PRO A  62       5.180  29.215 -36.051  1.00  0.00           O  
ATOM   1016  CB  PRO A  62       5.798  26.198 -36.615  1.00  0.00           C  
ATOM   1017  CG  PRO A  62       5.738  26.130 -35.127  1.00  0.00           C  
ATOM   1018  CD  PRO A  62       7.056  26.699 -34.673  1.00  0.00           C  
ATOM   1019  HA  PRO A  62       7.226  27.357 -37.795  1.00  0.00           H  
ATOM   1020 1HB  PRO A  62       4.803  26.265 -37.079  1.00  0.00           H  
ATOM   1021 2HB  PRO A  62       6.303  25.329 -37.061  1.00  0.00           H  
ATOM   1022 1HG  PRO A  62       4.878  26.706 -34.755  1.00  0.00           H  
ATOM   1023 2HG  PRO A  62       5.590  25.090 -34.799  1.00  0.00           H  
ATOM   1024 1HD  PRO A  62       6.913  27.252 -33.733  1.00  0.00           H  
ATOM   1025 2HD  PRO A  62       7.781  25.883 -34.536  1.00  0.00           H  
ATOM   1026  N   ALA A  63       5.479  29.135 -38.282  1.00  0.00           N  
ATOM   1027  CA  ALA A  63       4.605  30.275 -38.536  1.00  0.00           C  
ATOM   1028  C   ALA A  63       3.172  29.837 -38.269  1.00  0.00           C  
ATOM   1029  O   ALA A  63       2.869  28.643 -38.282  1.00  0.00           O  
ATOM   1030  CB  ALA A  63       4.750  30.741 -39.976  1.00  0.00           C  
ATOM   1031  H   ALA A  63       5.934  28.681 -39.061  1.00  0.00           H  
ATOM   1032  HA  ALA A  63       4.903  31.086 -37.871  1.00  0.00           H  
ATOM   1033 1HB  ALA A  63       4.092  31.592 -40.152  1.00  0.00           H  
ATOM   1034 2HB  ALA A  63       5.784  31.036 -40.160  1.00  0.00           H  
ATOM   1035 3HB  ALA A  63       4.480  29.929 -40.650  1.00  0.00           H  
ATOM   1036  N   THR A  64       2.285  30.799 -38.038  1.00  0.00           N  
ATOM   1037  CA  THR A  64       0.895  30.466 -37.753  1.00  0.00           C  
ATOM   1038  C   THR A  64      -0.088  31.223 -38.637  1.00  0.00           C  
ATOM   1039  O   THR A  64       0.220  32.294 -39.169  1.00  0.00           O  
ATOM   1040  CB  THR A  64       0.554  30.781 -36.269  1.00  0.00           C  
ATOM   1041  CG2 THR A  64       0.623  32.279 -36.016  1.00  0.00           C  
ATOM   1042  OG1 THR A  64      -0.768  30.320 -35.960  1.00  0.00           O  
ATOM   1043  H   THR A  64       2.565  31.770 -38.056  1.00  0.00           H  
ATOM   1044  HA  THR A  64       0.752  29.399 -37.925  1.00  0.00           H  
ATOM   1045  HB  THR A  64       1.265  30.274 -35.617  1.00  0.00           H  
ATOM   1046  HG1 THR A  64      -1.156  29.916 -36.740  1.00  0.00           H  
ATOM   1047 1HG2 THR A  64       0.383  32.484 -34.973  1.00  0.00           H  
ATOM   1048 2HG2 THR A  64       1.628  32.638 -36.235  1.00  0.00           H  
ATOM   1049 3HG2 THR A  64      -0.092  32.789 -36.660  1.00  0.00           H  
ATOM   1050  N   SER A  65      -1.265  30.632 -38.810  1.00  0.00           N  
ATOM   1051  CA  SER A  65      -2.343  31.255 -39.560  1.00  0.00           C  
ATOM   1052  C   SER A  65      -3.287  31.721 -38.462  1.00  0.00           C  
ATOM   1053  O   SER A  65      -3.772  30.909 -37.679  1.00  0.00           O  
ATOM   1054  CB  SER A  65      -3.017  30.290 -40.516  1.00  0.00           C  
ATOM   1055  OG  SER A  65      -2.127  29.866 -41.511  1.00  0.00           O  
ATOM   1056  H   SER A  65      -1.412  29.719 -38.403  1.00  0.00           H  
ATOM   1057  HA  SER A  65      -1.918  32.032 -40.197  1.00  0.00           H  
ATOM   1058 1HB  SER A  65      -3.385  29.427 -39.961  1.00  0.00           H  
ATOM   1059 2HB  SER A  65      -3.877  30.775 -40.975  1.00  0.00           H  
ATOM   1060  HG  SER A  65      -2.516  30.138 -42.346  1.00  0.00           H  
ATOM   1061  N   VAL A  66      -3.526  33.024 -38.387  1.00  0.00           N  
ATOM   1062  CA  VAL A  66      -4.416  33.565 -37.367  1.00  0.00           C  
ATOM   1063  C   VAL A  66      -5.836  33.624 -37.910  1.00  0.00           C  
ATOM   1064  O   VAL A  66      -6.101  34.292 -38.909  1.00  0.00           O  
ATOM   1065  CB  VAL A  66      -3.964  34.974 -36.939  1.00  0.00           C  
ATOM   1066  CG1 VAL A  66      -4.927  35.553 -35.913  1.00  0.00           C  
ATOM   1067  CG2 VAL A  66      -2.551  34.916 -36.379  1.00  0.00           C  
ATOM   1068  H   VAL A  66      -3.089  33.656 -39.043  1.00  0.00           H  
ATOM   1069  HA  VAL A  66      -4.383  32.910 -36.496  1.00  0.00           H  
ATOM   1070  HB  VAL A  66      -3.985  35.632 -37.807  1.00  0.00           H  
ATOM   1071 1HG1 VAL A  66      -4.592  36.549 -35.622  1.00  0.00           H  
ATOM   1072 2HG1 VAL A  66      -5.924  35.619 -36.347  1.00  0.00           H  
ATOM   1073 3HG1 VAL A  66      -4.954  34.909 -35.035  1.00  0.00           H  
ATOM   1074 1HG2 VAL A  66      -2.237  35.915 -36.079  1.00  0.00           H  
ATOM   1075 2HG2 VAL A  66      -2.529  34.253 -35.514  1.00  0.00           H  
ATOM   1076 3HG2 VAL A  66      -1.872  34.537 -37.143  1.00  0.00           H  
ATOM   1077  N   VAL A  67      -6.739  32.908 -37.248  1.00  0.00           N  
ATOM   1078  CA  VAL A  67      -8.139  32.852 -37.655  1.00  0.00           C  
ATOM   1079  C   VAL A  67      -8.999  33.669 -36.707  1.00  0.00           C  
ATOM   1080  O   VAL A  67      -9.052  33.397 -35.509  1.00  0.00           O  
ATOM   1081  CB  VAL A  67      -8.640  31.395 -37.676  1.00  0.00           C  
ATOM   1082  CG1 VAL A  67     -10.109  31.343 -38.068  1.00  0.00           C  
ATOM   1083  CG2 VAL A  67      -7.795  30.572 -38.637  1.00  0.00           C  
ATOM   1084  H   VAL A  67      -6.442  32.386 -36.436  1.00  0.00           H  
ATOM   1085  HA  VAL A  67      -8.221  33.246 -38.668  1.00  0.00           H  
ATOM   1086  HB  VAL A  67      -8.559  30.979 -36.672  1.00  0.00           H  
ATOM   1087 1HG1 VAL A  67     -10.446  30.306 -38.077  1.00  0.00           H  
ATOM   1088 2HG1 VAL A  67     -10.698  31.909 -37.346  1.00  0.00           H  
ATOM   1089 3HG1 VAL A  67     -10.236  31.774 -39.060  1.00  0.00           H  
ATOM   1090 1HG2 VAL A  67      -8.154  29.543 -38.646  1.00  0.00           H  
ATOM   1091 2HG2 VAL A  67      -7.871  30.992 -39.640  1.00  0.00           H  
ATOM   1092 3HG2 VAL A  67      -6.754  30.590 -38.313  1.00  0.00           H  
ATOM   1093  N   ILE A  68      -9.669  34.673 -37.263  1.00  0.00           N  
ATOM   1094  CA  ILE A  68     -10.544  35.546 -36.494  1.00  0.00           C  
ATOM   1095  C   ILE A  68     -11.910  35.561 -37.160  1.00  0.00           C  
ATOM   1096  O   ILE A  68     -12.048  36.077 -38.269  1.00  0.00           O  
ATOM   1097  CB  ILE A  68      -9.982  36.976 -36.400  1.00  0.00           C  
ATOM   1098  CG1 ILE A  68      -8.598  36.965 -35.747  1.00  0.00           C  
ATOM   1099  CG2 ILE A  68     -10.934  37.872 -35.622  1.00  0.00           C  
ATOM   1100  CD1 ILE A  68      -7.895  38.302 -35.782  1.00  0.00           C  
ATOM   1101  H   ILE A  68      -9.565  34.831 -38.255  1.00  0.00           H  
ATOM   1102  HA  ILE A  68     -10.581  35.181 -35.468  1.00  0.00           H  
ATOM   1103  HB  ILE A  68      -9.854  37.383 -37.403  1.00  0.00           H  
ATOM   1104 1HG1 ILE A  68      -8.689  36.654 -34.707  1.00  0.00           H  
ATOM   1105 2HG1 ILE A  68      -7.964  36.234 -36.251  1.00  0.00           H  
ATOM   1106 1HG2 ILE A  68     -10.521  38.879 -35.565  1.00  0.00           H  
ATOM   1107 2HG2 ILE A  68     -11.899  37.903 -36.127  1.00  0.00           H  
ATOM   1108 3HG2 ILE A  68     -11.065  37.475 -34.615  1.00  0.00           H  
ATOM   1109 1HD1 ILE A  68      -6.920  38.214 -35.301  1.00  0.00           H  
ATOM   1110 2HD1 ILE A  68      -7.761  38.616 -36.818  1.00  0.00           H  
ATOM   1111 3HD1 ILE A  68      -8.494  39.042 -35.253  1.00  0.00           H  
ATOM   1112  N   THR A  69     -12.914  34.979 -36.509  1.00  0.00           N  
ATOM   1113  CA  THR A  69     -14.251  35.012 -37.091  1.00  0.00           C  
ATOM   1114  C   THR A  69     -14.958  36.201 -36.462  1.00  0.00           C  
ATOM   1115  O   THR A  69     -14.584  36.654 -35.379  1.00  0.00           O  
ATOM   1116  CB  THR A  69     -15.079  33.719 -36.814  1.00  0.00           C  
ATOM   1117  CG2 THR A  69     -14.181  32.498 -36.833  1.00  0.00           C  
ATOM   1118  OG1 THR A  69     -15.737  33.810 -35.541  1.00  0.00           O  
ATOM   1119  H   THR A  69     -12.774  34.516 -35.623  1.00  0.00           H  
ATOM   1120  HA  THR A  69     -14.156  35.108 -38.173  1.00  0.00           H  
ATOM   1121  HB  THR A  69     -15.849  33.609 -37.578  1.00  0.00           H  
ATOM   1122  HG1 THR A  69     -15.525  34.651 -35.131  1.00  0.00           H  
ATOM   1123 1HG2 THR A  69     -14.775  31.606 -36.638  1.00  0.00           H  
ATOM   1124 2HG2 THR A  69     -13.706  32.412 -37.810  1.00  0.00           H  
ATOM   1125 3HG2 THR A  69     -13.414  32.598 -36.065  1.00  0.00           H  
ATOM   1126  N   PHE A  70     -15.970  36.726 -37.139  1.00  0.00           N  
ATOM   1127  CA  PHE A  70     -16.685  37.858 -36.579  1.00  0.00           C  
ATOM   1128  C   PHE A  70     -18.074  38.008 -37.179  1.00  0.00           C  
ATOM   1129  O   PHE A  70     -18.300  37.683 -38.343  1.00  0.00           O  
ATOM   1130  CB  PHE A  70     -15.867  39.152 -36.769  1.00  0.00           C  
ATOM   1131  CG  PHE A  70     -15.772  39.626 -38.200  1.00  0.00           C  
ATOM   1132  CD1 PHE A  70     -16.683  40.555 -38.701  1.00  0.00           C  
ATOM   1133  CD2 PHE A  70     -14.769  39.152 -39.043  1.00  0.00           C  
ATOM   1134  CE1 PHE A  70     -16.596  41.005 -40.020  1.00  0.00           C  
ATOM   1135  CE2 PHE A  70     -14.673  39.595 -40.362  1.00  0.00           C  
ATOM   1136  CZ  PHE A  70     -15.588  40.524 -40.852  1.00  0.00           C  
ATOM   1137  H   PHE A  70     -16.255  36.358 -38.035  1.00  0.00           H  
ATOM   1138  HA  PHE A  70     -16.825  37.686 -35.511  1.00  0.00           H  
ATOM   1139 1HB  PHE A  70     -16.312  39.954 -36.180  1.00  0.00           H  
ATOM   1140 2HB  PHE A  70     -14.854  38.999 -36.401  1.00  0.00           H  
ATOM   1141  HD1 PHE A  70     -17.472  40.931 -38.049  1.00  0.00           H  
ATOM   1142  HD2 PHE A  70     -14.054  38.425 -38.659  1.00  0.00           H  
ATOM   1143  HE1 PHE A  70     -17.315  41.732 -40.395  1.00  0.00           H  
ATOM   1144  HE2 PHE A  70     -13.883  39.215 -41.010  1.00  0.00           H  
ATOM   1145  HZ  PHE A  70     -15.518  40.872 -41.882  1.00  0.00           H  
ATOM   1146  N   HIS A  71     -19.004  38.477 -36.358  1.00  0.00           N  
ATOM   1147  CA  HIS A  71     -20.374  38.696 -36.792  1.00  0.00           C  
ATOM   1148  C   HIS A  71     -20.824  40.027 -36.215  1.00  0.00           C  
ATOM   1149  O   HIS A  71     -21.037  40.146 -35.010  1.00  0.00           O  
ATOM   1150  CB  HIS A  71     -21.303  37.567 -36.334  1.00  0.00           C  
ATOM   1151  CG  HIS A  71     -22.703  37.693 -36.849  1.00  0.00           C  
ATOM   1152  CD2 HIS A  71     -23.858  38.018 -36.222  1.00  0.00           C  
ATOM   1153  ND1 HIS A  71     -23.032  37.473 -38.170  1.00  0.00           N  
ATOM   1154  CE1 HIS A  71     -24.331  37.656 -38.332  1.00  0.00           C  
ATOM   1155  NE2 HIS A  71     -24.854  37.988 -37.166  1.00  0.00           N  
ATOM   1156  H   HIS A  71     -18.751  38.688 -35.403  1.00  0.00           H  
ATOM   1157  HA  HIS A  71     -20.416  38.693 -37.881  1.00  0.00           H  
ATOM   1158 1HB  HIS A  71     -20.902  36.608 -36.667  1.00  0.00           H  
ATOM   1159 2HB  HIS A  71     -21.340  37.545 -35.246  1.00  0.00           H  
ATOM   1160  HD1 HIS A  71     -22.417  37.143 -38.886  1.00  0.00           H  
ATOM   1161  HD2 HIS A  71     -24.092  38.274 -35.188  1.00  0.00           H  
ATOM   1162  HE1 HIS A  71     -24.792  37.528 -39.312  1.00  0.00           H  
ATOM   1163  N   ASN A  72     -20.961  41.023 -37.084  1.00  0.00           N  
ATOM   1164  CA  ASN A  72     -21.378  42.360 -36.672  1.00  0.00           C  
ATOM   1165  C   ASN A  72     -20.516  42.921 -35.554  1.00  0.00           C  
ATOM   1166  O   ASN A  72     -21.016  43.324 -34.502  1.00  0.00           O  
ATOM   1167  CB  ASN A  72     -22.837  42.351 -36.253  1.00  0.00           C  
ATOM   1168  CG  ASN A  72     -23.761  41.996 -37.385  1.00  0.00           C  
ATOM   1169  ND2 ASN A  72     -24.644  41.061 -37.147  1.00  0.00           N  
ATOM   1170  OD1 ASN A  72     -23.674  42.570 -38.477  1.00  0.00           O  
ATOM   1171  H   ASN A  72     -20.769  40.846 -38.059  1.00  0.00           H  
ATOM   1172  HA  ASN A  72     -21.260  43.036 -37.520  1.00  0.00           H  
ATOM   1173 1HB  ASN A  72     -22.979  41.632 -35.445  1.00  0.00           H  
ATOM   1174 2HB  ASN A  72     -23.112  43.333 -35.870  1.00  0.00           H  
ATOM   1175 1HD2 ASN A  72     -25.285  40.784 -37.863  1.00  0.00           H  
ATOM   1176 2HD2 ASN A  72     -24.679  40.623 -36.249  1.00  0.00           H  
ATOM   1177  N   GLU A  73     -19.212  42.939 -35.801  1.00  0.00           N  
ATOM   1178  CA  GLU A  73     -18.251  43.458 -34.851  1.00  0.00           C  
ATOM   1179  C   GLU A  73     -18.099  44.956 -35.080  1.00  0.00           C  
ATOM   1180  O   GLU A  73     -18.332  45.450 -36.181  1.00  0.00           O  
ATOM   1181  CB  GLU A  73     -16.893  42.777 -35.051  1.00  0.00           C  
ATOM   1182  CG  GLU A  73     -15.849  43.229 -34.053  1.00  0.00           C  
ATOM   1183  CD  GLU A  73     -16.291  42.941 -32.641  1.00  0.00           C  
ATOM   1184  OE1 GLU A  73     -16.240  41.759 -32.235  1.00  0.00           O  
ATOM   1185  OE2 GLU A  73     -16.717  43.890 -31.948  1.00  0.00           O1-
ATOM   1186  H   GLU A  73     -18.884  42.578 -36.686  1.00  0.00           H  
ATOM   1187  HA  GLU A  73     -18.608  43.242 -33.843  1.00  0.00           H  
ATOM   1188 1HB  GLU A  73     -17.012  41.697 -34.963  1.00  0.00           H  
ATOM   1189 2HB  GLU A  73     -16.526  42.985 -36.056  1.00  0.00           H  
ATOM   1190 1HG  GLU A  73     -14.914  42.709 -34.261  1.00  0.00           H  
ATOM   1191 2HG  GLU A  73     -15.674  44.296 -34.184  1.00  0.00           H  
ATOM   1192  N   ALA A  74     -17.724  45.683 -34.036  1.00  0.00           N  
ATOM   1193  CA  ALA A  74     -17.514  47.115 -34.180  1.00  0.00           C  
ATOM   1194  C   ALA A  74     -16.239  47.260 -35.011  1.00  0.00           C  
ATOM   1195  O   ALA A  74     -15.219  46.643 -34.703  1.00  0.00           O  
ATOM   1196  CB  ALA A  74     -17.336  47.766 -32.811  1.00  0.00           C  
ATOM   1197  H   ALA A  74     -17.580  45.254 -33.133  1.00  0.00           H  
ATOM   1198  HA  ALA A  74     -18.394  47.540 -34.661  1.00  0.00           H  
ATOM   1199 1HB  ALA A  74     -17.180  48.837 -32.935  1.00  0.00           H  
ATOM   1200 2HB  ALA A  74     -18.229  47.596 -32.209  1.00  0.00           H  
ATOM   1201 3HB  ALA A  74     -16.473  47.329 -32.311  1.00  0.00           H  
ATOM   1202  N   ARG A  75     -16.297  48.060 -36.071  1.00  0.00           N  
ATOM   1203  CA  ARG A  75     -15.132  48.254 -36.932  1.00  0.00           C  
ATOM   1204  C   ARG A  75     -13.849  48.617 -36.190  1.00  0.00           C  
ATOM   1205  O   ARG A  75     -12.790  48.067 -36.485  1.00  0.00           O  
ATOM   1206  CB  ARG A  75     -15.425  49.314 -37.997  1.00  0.00           C  
ATOM   1207  CG  ARG A  75     -16.339  48.819 -39.104  1.00  0.00           C  
ATOM   1208  CD  ARG A  75     -16.397  49.800 -40.267  1.00  0.00           C  
ATOM   1209  NE  ARG A  75     -17.239  49.294 -41.349  1.00  0.00           N  
ATOM   1210  CZ  ARG A  75     -17.426  49.919 -42.508  1.00  0.00           C  
ATOM   1211  NH1 ARG A  75     -16.829  51.081 -42.746  1.00  0.00           N1+
ATOM   1212  NH2 ARG A  75     -18.214  49.380 -43.432  1.00  0.00           N  
ATOM   1213  H   ARG A  75     -17.156  48.543 -36.291  1.00  0.00           H  
ATOM   1214  HA  ARG A  75     -14.910  47.310 -37.431  1.00  0.00           H  
ATOM   1215 1HB  ARG A  75     -15.888  50.181 -37.529  1.00  0.00           H  
ATOM   1216 2HB  ARG A  75     -14.488  49.646 -38.446  1.00  0.00           H  
ATOM   1217 1HG  ARG A  75     -15.972  47.864 -39.479  1.00  0.00           H  
ATOM   1218 2HG  ARG A  75     -17.349  48.692 -38.713  1.00  0.00           H  
ATOM   1219 1HD  ARG A  75     -16.808  50.748 -39.923  1.00  0.00           H  
ATOM   1220 2HD  ARG A  75     -15.392  49.961 -40.658  1.00  0.00           H  
ATOM   1221  HE  ARG A  75     -17.711  48.411 -41.204  1.00  0.00           H  
ATOM   1222 1HH1 ARG A  75     -16.230  51.495 -42.044  1.00  0.00           H  
ATOM   1223 2HH1 ARG A  75     -16.973  51.552 -43.627  1.00  0.00           H  
ATOM   1224 1HH2 ARG A  75     -18.671  48.496 -43.253  1.00  0.00           H  
ATOM   1225 2HH2 ARG A  75     -18.356  49.853 -44.312  1.00  0.00           H  
ATOM   1226  N   SER A  76     -13.927  49.540 -35.234  1.00  0.00           N  
ATOM   1227  CA  SER A  76     -12.726  49.924 -34.500  1.00  0.00           C  
ATOM   1228  C   SER A  76     -12.109  48.746 -33.751  1.00  0.00           C  
ATOM   1229  O   SER A  76     -10.891  48.587 -33.737  1.00  0.00           O  
ATOM   1230  CB  SER A  76     -13.054  51.035 -33.521  1.00  0.00           C  
ATOM   1231  OG  SER A  76     -13.946  50.590 -32.537  1.00  0.00           O  
ATOM   1232  H   SER A  76     -14.806  49.982 -35.005  1.00  0.00           H  
ATOM   1233  HA  SER A  76     -11.981  50.275 -35.215  1.00  0.00           H  
ATOM   1234 1HB  SER A  76     -12.136  51.388 -33.051  1.00  0.00           H  
ATOM   1235 2HB  SER A  76     -13.491  51.875 -34.059  1.00  0.00           H  
ATOM   1236  HG  SER A  76     -14.725  51.146 -32.615  1.00  0.00           H  
ATOM   1237  N   ALA A  77     -12.949  47.918 -33.140  1.00  0.00           N  
ATOM   1238  CA  ALA A  77     -12.469  46.762 -32.390  1.00  0.00           C  
ATOM   1239  C   ALA A  77     -11.863  45.699 -33.304  1.00  0.00           C  
ATOM   1240  O   ALA A  77     -10.828  45.115 -32.983  1.00  0.00           O  
ATOM   1241  CB  ALA A  77     -13.611  46.164 -31.572  1.00  0.00           C  
ATOM   1242  H   ALA A  77     -13.942  48.091 -33.195  1.00  0.00           H  
ATOM   1243  HA  ALA A  77     -11.684  47.101 -31.714  1.00  0.00           H  
ATOM   1244 1HB  ALA A  77     -13.246  45.302 -31.015  1.00  0.00           H  
ATOM   1245 2HB  ALA A  77     -13.991  46.913 -30.877  1.00  0.00           H  
ATOM   1246 3HB  ALA A  77     -14.412  45.852 -32.241  1.00  0.00           H  
ATOM   1247  N   LEU A  78     -12.504  45.453 -34.442  1.00  0.00           N  
ATOM   1248  CA  LEU A  78     -12.013  44.453 -35.391  1.00  0.00           C  
ATOM   1249  C   LEU A  78     -10.660  44.886 -35.948  1.00  0.00           C  
ATOM   1250  O   LEU A  78      -9.704  44.103 -35.978  1.00  0.00           O  
ATOM   1251  CB  LEU A  78     -13.021  44.271 -36.535  1.00  0.00           C  
ATOM   1252  CG  LEU A  78     -12.663  43.290 -37.655  1.00  0.00           C  
ATOM   1253  CD1 LEU A  78     -12.423  41.892 -37.088  1.00  0.00           C  
ATOM   1254  CD2 LEU A  78     -13.794  43.263 -38.671  1.00  0.00           C  
ATOM   1255  H   LEU A  78     -13.348  45.964 -34.659  1.00  0.00           H  
ATOM   1256  HA  LEU A  78     -11.899  43.505 -34.868  1.00  0.00           H  
ATOM   1257 1HB  LEU A  78     -13.966  43.930 -36.113  1.00  0.00           H  
ATOM   1258 2HB  LEU A  78     -13.187  45.238 -37.009  1.00  0.00           H  
ATOM   1259  HG  LEU A  78     -11.741  43.612 -38.140  1.00  0.00           H  
ATOM   1260 1HD1 LEU A  78     -12.169  41.209 -37.898  1.00  0.00           H  
ATOM   1261 2HD1 LEU A  78     -11.601  41.927 -36.373  1.00  0.00           H  
ATOM   1262 3HD1 LEU A  78     -13.325  41.543 -36.588  1.00  0.00           H  
ATOM   1263 1HD2 LEU A  78     -13.546  42.567 -39.473  1.00  0.00           H  
ATOM   1264 2HD2 LEU A  78     -14.714  42.942 -38.183  1.00  0.00           H  
ATOM   1265 3HD2 LEU A  78     -13.933  44.261 -39.087  1.00  0.00           H  
ATOM   1266  N   LEU A  79     -10.577  46.138 -36.382  1.00  0.00           N  
ATOM   1267  CA  LEU A  79      -9.333  46.670 -36.928  1.00  0.00           C  
ATOM   1268  C   LEU A  79      -8.210  46.605 -35.894  1.00  0.00           C  
ATOM   1269  O   LEU A  79      -7.091  46.189 -36.201  1.00  0.00           O  
ATOM   1270  CB  LEU A  79      -9.535  48.120 -37.370  1.00  0.00           C  
ATOM   1271  CG  LEU A  79      -8.271  48.884 -37.775  1.00  0.00           C  
ATOM   1272  CD1 LEU A  79      -7.642  48.230 -38.993  1.00  0.00           C  
ATOM   1273  CD2 LEU A  79      -8.627  50.337 -38.069  1.00  0.00           C  
ATOM   1274  H   LEU A  79     -11.389  46.737 -36.335  1.00  0.00           H  
ATOM   1275  HA  LEU A  79      -9.052  46.072 -37.794  1.00  0.00           H  
ATOM   1276 1HB  LEU A  79     -10.211  48.132 -38.224  1.00  0.00           H  
ATOM   1277 2HB  LEU A  79     -10.002  48.671 -36.555  1.00  0.00           H  
ATOM   1278  HG  LEU A  79      -7.547  48.846 -36.962  1.00  0.00           H  
ATOM   1279 1HD1 LEU A  79      -6.743  48.778 -39.276  1.00  0.00           H  
ATOM   1280 2HD1 LEU A  79      -7.378  47.199 -38.757  1.00  0.00           H  
ATOM   1281 3HD1 LEU A  79      -8.351  48.244 -39.820  1.00  0.00           H  
ATOM   1282 1HD2 LEU A  79      -7.728  50.881 -38.356  1.00  0.00           H  
ATOM   1283 2HD2 LEU A  79      -9.351  50.376 -38.883  1.00  0.00           H  
ATOM   1284 3HD2 LEU A  79      -9.059  50.793 -37.178  1.00  0.00           H  
ATOM   1285  N   ARG A  80      -8.512  47.017 -34.669  1.00  0.00           N  
ATOM   1286  CA  ARG A  80      -7.519  47.017 -33.601  1.00  0.00           C  
ATOM   1287  C   ARG A  80      -7.007  45.603 -33.309  1.00  0.00           C  
ATOM   1288  O   ARG A  80      -5.833  45.413 -32.993  1.00  0.00           O  
ATOM   1289  CB  ARG A  80      -8.121  47.657 -32.343  1.00  0.00           C  
ATOM   1290  CG  ARG A  80      -7.113  48.023 -31.267  1.00  0.00           C  
ATOM   1291  CD  ARG A  80      -7.667  49.122 -30.359  1.00  0.00           C  
ATOM   1292  NE  ARG A  80      -8.936  48.748 -29.736  1.00  0.00           N  
ATOM   1293  CZ  ARG A  80     -10.100  49.353 -29.960  1.00  0.00           C  
ATOM   1294  NH1 ARG A  80     -11.191  48.933 -29.334  1.00  0.00           N1+
ATOM   1295  NH2 ARG A  80     -10.177  50.373 -30.807  1.00  0.00           N  
ATOM   1296  H   ARG A  80      -9.449  47.337 -34.470  1.00  0.00           H  
ATOM   1297  HA  ARG A  80      -6.661  47.607 -33.925  1.00  0.00           H  
ATOM   1298 1HB  ARG A  80      -8.654  48.565 -32.617  1.00  0.00           H  
ATOM   1299 2HB  ARG A  80      -8.846  46.974 -31.898  1.00  0.00           H  
ATOM   1300 1HG  ARG A  80      -6.892  47.143 -30.661  1.00  0.00           H  
ATOM   1301 2HG  ARG A  80      -6.196  48.381 -31.735  1.00  0.00           H  
ATOM   1302 1HD  ARG A  80      -6.952  49.331 -29.564  1.00  0.00           H  
ATOM   1303 2HD  ARG A  80      -7.835  50.025 -30.943  1.00  0.00           H  
ATOM   1304  HE  ARG A  80      -8.929  47.972 -29.087  1.00  0.00           H  
ATOM   1305 1HH1 ARG A  80     -11.136  48.157 -28.690  1.00  0.00           H  
ATOM   1306 2HH1 ARG A  80     -12.076  49.389 -29.501  1.00  0.00           H  
ATOM   1307 1HH2 ARG A  80      -9.348  50.696 -31.287  1.00  0.00           H  
ATOM   1308 2HH2 ARG A  80     -11.064  50.827 -30.971  1.00  0.00           H  
ATOM   1309  N   THR A  81      -7.879  44.605 -33.425  1.00  0.00           N  
ATOM   1310  CA  THR A  81      -7.454  43.229 -33.185  1.00  0.00           C  
ATOM   1311  C   THR A  81      -6.413  42.851 -34.238  1.00  0.00           C  
ATOM   1312  O   THR A  81      -5.341  42.331 -33.921  1.00  0.00           O  
ATOM   1313  CB  THR A  81      -8.640  42.248 -33.229  1.00  0.00           C  
ATOM   1314  CG2 THR A  81      -8.158  40.819 -33.029  1.00  0.00           C  
ATOM   1315  OG1 THR A  81      -9.577  42.581 -32.196  1.00  0.00           O  
ATOM   1316  H   THR A  81      -8.840  44.785 -33.678  1.00  0.00           H  
ATOM   1317  HA  THR A  81      -7.058  43.161 -32.171  1.00  0.00           H  
ATOM   1318  HB  THR A  81      -9.141  42.325 -34.193  1.00  0.00           H  
ATOM   1319  HG1 THR A  81     -10.122  43.317 -32.484  1.00  0.00           H  
ATOM   1320 1HG2 THR A  81      -9.010  40.140 -33.064  1.00  0.00           H  
ATOM   1321 2HG2 THR A  81      -7.454  40.559 -33.820  1.00  0.00           H  
ATOM   1322 3HG2 THR A  81      -7.665  40.733 -32.062  1.00  0.00           H  
ATOM   1323  N   VAL A  82      -6.742  43.127 -35.496  1.00  0.00           N  
ATOM   1324  CA  VAL A  82      -5.859  42.818 -36.613  1.00  0.00           C  
ATOM   1325  C   VAL A  82      -4.525  43.539 -36.494  1.00  0.00           C  
ATOM   1326  O   VAL A  82      -3.468  42.930 -36.668  1.00  0.00           O  
ATOM   1327  CB  VAL A  82      -6.523  43.195 -37.952  1.00  0.00           C  
ATOM   1328  CG1 VAL A  82      -5.521  43.064 -39.097  1.00  0.00           C  
ATOM   1329  CG2 VAL A  82      -7.716  42.287 -38.203  1.00  0.00           C  
ATOM   1330  H   VAL A  82      -7.634  43.565 -35.680  1.00  0.00           H  
ATOM   1331  HA  VAL A  82      -5.662  41.746 -36.612  1.00  0.00           H  
ATOM   1332  HB  VAL A  82      -6.856  44.232 -37.903  1.00  0.00           H  
ATOM   1333 1HG1 VAL A  82      -6.004  43.334 -40.036  1.00  0.00           H  
ATOM   1334 2HG1 VAL A  82      -4.677  43.731 -38.919  1.00  0.00           H  
ATOM   1335 3HG1 VAL A  82      -5.167  42.035 -39.155  1.00  0.00           H  
ATOM   1336 1HG2 VAL A  82      -8.184  42.556 -39.149  1.00  0.00           H  
ATOM   1337 2HG2 VAL A  82      -7.382  41.250 -38.244  1.00  0.00           H  
ATOM   1338 3HG2 VAL A  82      -8.439  42.404 -37.395  1.00  0.00           H  
ATOM   1339  N   VAL A  83      -4.574  44.835 -36.195  1.00  0.00           N  
ATOM   1340  CA  VAL A  83      -3.350  45.608 -36.055  1.00  0.00           C  
ATOM   1341  C   VAL A  83      -2.505  45.109 -34.881  1.00  0.00           C  
ATOM   1342  O   VAL A  83      -1.284  45.039 -34.993  1.00  0.00           O  
ATOM   1343  CB  VAL A  83      -3.655  47.123 -35.900  1.00  0.00           C  
ATOM   1344  CG1 VAL A  83      -2.391  47.884 -35.516  1.00  0.00           C  
ATOM   1345  CG2 VAL A  83      -4.196  47.672 -37.218  1.00  0.00           C  
ATOM   1346  H   VAL A  83      -5.464  45.293 -36.062  1.00  0.00           H  
ATOM   1347  HA  VAL A  83      -2.749  45.471 -36.955  1.00  0.00           H  
ATOM   1348  HB  VAL A  83      -4.398  47.256 -35.114  1.00  0.00           H  
ATOM   1349 1HG1 VAL A  83      -2.622  48.944 -35.411  1.00  0.00           H  
ATOM   1350 2HG1 VAL A  83      -2.010  47.501 -34.569  1.00  0.00           H  
ATOM   1351 3HG1 VAL A  83      -1.637  47.753 -36.291  1.00  0.00           H  
ATOM   1352 1HG2 VAL A  83      -4.411  48.734 -37.108  1.00  0.00           H  
ATOM   1353 2HG2 VAL A  83      -3.454  47.530 -38.004  1.00  0.00           H  
ATOM   1354 3HG2 VAL A  83      -5.111  47.143 -37.484  1.00  0.00           H  
ATOM   1355  N   SER A  84      -3.132  44.739 -33.764  1.00  0.00           N  
ATOM   1356  CA  SER A  84      -2.341  44.257 -32.631  1.00  0.00           C  
ATOM   1357  C   SER A  84      -1.583  42.987 -33.014  1.00  0.00           C  
ATOM   1358  O   SER A  84      -0.459  42.770 -32.570  1.00  0.00           O  
ATOM   1359  CB  SER A  84      -3.218  44.013 -31.387  1.00  0.00           C  
ATOM   1360  OG  SER A  84      -4.017  42.849 -31.494  1.00  0.00           O  
ATOM   1361  H   SER A  84      -4.137  44.783 -33.678  1.00  0.00           H  
ATOM   1362  HA  SER A  84      -1.599  45.017 -32.379  1.00  0.00           H  
ATOM   1363 1HB  SER A  84      -2.583  43.920 -30.507  1.00  0.00           H  
ATOM   1364 2HB  SER A  84      -3.872  44.870 -31.230  1.00  0.00           H  
ATOM   1365  HG  SER A  84      -3.817  42.469 -32.352  1.00  0.00           H  
ATOM   1366  N   VAL A  85      -2.189  42.152 -33.853  1.00  0.00           N  
ATOM   1367  CA  VAL A  85      -1.524  40.927 -34.289  1.00  0.00           C  
ATOM   1368  C   VAL A  85      -0.308  41.284 -35.144  1.00  0.00           C  
ATOM   1369  O   VAL A  85       0.794  40.777 -34.929  1.00  0.00           O  
ATOM   1370  CB  VAL A  85      -2.478  40.031 -35.120  1.00  0.00           C  
ATOM   1371  CG1 VAL A  85      -1.712  38.843 -35.703  1.00  0.00           C  
ATOM   1372  CG2 VAL A  85      -3.618  39.530 -34.234  1.00  0.00           C  
ATOM   1373  H   VAL A  85      -3.117  42.357 -34.196  1.00  0.00           H  
ATOM   1374  HA  VAL A  85      -1.215  40.367 -33.406  1.00  0.00           H  
ATOM   1375  HB  VAL A  85      -2.886  40.618 -35.944  1.00  0.00           H  
ATOM   1376 1HG1 VAL A  85      -2.393  38.221 -36.285  1.00  0.00           H  
ATOM   1377 2HG1 VAL A  85      -0.913  39.206 -36.348  1.00  0.00           H  
ATOM   1378 3HG1 VAL A  85      -1.286  38.253 -34.892  1.00  0.00           H  
ATOM   1379 1HG2 VAL A  85      -4.287  38.902 -34.821  1.00  0.00           H  
ATOM   1380 2HG2 VAL A  85      -3.208  38.950 -33.407  1.00  0.00           H  
ATOM   1381 3HG2 VAL A  85      -4.174  40.381 -33.840  1.00  0.00           H  
ATOM   1382  N   LEU A  86      -0.510  42.177 -36.105  1.00  0.00           N  
ATOM   1383  CA  LEU A  86       0.560  42.586 -37.005  1.00  0.00           C  
ATOM   1384  C   LEU A  86       1.689  43.358 -36.332  1.00  0.00           C  
ATOM   1385  O   LEU A  86       2.863  43.153 -36.647  1.00  0.00           O  
ATOM   1386  CB  LEU A  86      -0.027  43.444 -38.132  1.00  0.00           C  
ATOM   1387  CG  LEU A  86      -0.934  42.707 -39.126  1.00  0.00           C  
ATOM   1388  CD1 LEU A  86      -1.575  43.713 -40.073  1.00  0.00           C  
ATOM   1389  CD2 LEU A  86      -0.117  41.678 -39.893  1.00  0.00           C  
ATOM   1390  H   LEU A  86      -1.428  42.582 -36.215  1.00  0.00           H  
ATOM   1391  HA  LEU A  86       1.027  41.692 -37.418  1.00  0.00           H  
ATOM   1392 1HB  LEU A  86      -0.609  44.250 -37.688  1.00  0.00           H  
ATOM   1393 2HB  LEU A  86       0.794  43.886 -38.697  1.00  0.00           H  
ATOM   1394  HG  LEU A  86      -1.735  42.203 -38.584  1.00  0.00           H  
ATOM   1395 1HD1 LEU A  86      -2.219  43.190 -40.779  1.00  0.00           H  
ATOM   1396 2HD1 LEU A  86      -2.169  44.425 -39.500  1.00  0.00           H  
ATOM   1397 3HD1 LEU A  86      -0.796  44.246 -40.618  1.00  0.00           H  
ATOM   1398 1HD2 LEU A  86      -0.761  41.154 -40.599  1.00  0.00           H  
ATOM   1399 2HD2 LEU A  86       0.683  42.181 -40.437  1.00  0.00           H  
ATOM   1400 3HD2 LEU A  86       0.315  40.962 -39.194  1.00  0.00           H  
ATOM   1401  N   LYS A  87       1.342  44.240 -35.403  1.00  0.00           N  
ATOM   1402  CA  LYS A  87       2.358  45.038 -34.730  1.00  0.00           C  
ATOM   1403  C   LYS A  87       3.128  44.298 -33.639  1.00  0.00           C  
ATOM   1404  O   LYS A  87       4.336  44.494 -33.493  1.00  0.00           O  
ATOM   1405  CB  LYS A  87       1.710  46.288 -34.133  1.00  0.00           C  
ATOM   1406  CG  LYS A  87       2.690  47.252 -33.476  1.00  0.00           C  
ATOM   1407  CD  LYS A  87       1.996  48.534 -33.040  1.00  0.00           C  
ATOM   1408  CE  LYS A  87       2.976  49.503 -32.397  1.00  0.00           C  
ATOM   1409  NZ  LYS A  87       2.311  50.763 -31.965  1.00  0.00           N1+
ATOM   1410  H   LYS A  87       0.372  44.368 -35.153  1.00  0.00           H  
ATOM   1411  HA  LYS A  87       3.111  45.329 -35.464  1.00  0.00           H  
ATOM   1412 1HB  LYS A  87       1.180  46.832 -34.915  1.00  0.00           H  
ATOM   1413 2HB  LYS A  87       0.976  45.995 -33.383  1.00  0.00           H  
ATOM   1414 1HG  LYS A  87       3.138  46.776 -32.603  1.00  0.00           H  
ATOM   1415 2HG  LYS A  87       3.484  47.500 -34.180  1.00  0.00           H  
ATOM   1416 1HD  LYS A  87       1.539  49.013 -33.907  1.00  0.00           H  
ATOM   1417 2HD  LYS A  87       1.211  48.296 -32.323  1.00  0.00           H  
ATOM   1418 1HE  LYS A  87       3.435  49.033 -31.528  1.00  0.00           H  
ATOM   1419 2HE  LYS A  87       3.764  49.750 -33.108  1.00  0.00           H  
ATOM   1420 1HZ  LYS A  87       2.993  51.378 -31.545  1.00  0.00           H  
ATOM   1421 2HZ  LYS A  87       1.897  51.217 -32.767  1.00  0.00           H  
ATOM   1422 3HZ  LYS A  87       1.590  50.549 -31.291  1.00  0.00           H  
ATOM   1423  N   LYS A  88       2.442  43.437 -32.892  1.00  0.00           N  
ATOM   1424  CA  LYS A  88       3.075  42.726 -31.784  1.00  0.00           C  
ATOM   1425  C   LYS A  88       3.633  41.331 -32.060  1.00  0.00           C  
ATOM   1426  O   LYS A  88       4.184  40.695 -31.157  1.00  0.00           O  
ATOM   1427  CB  LYS A  88       2.073  42.628 -30.633  1.00  0.00           C  
ATOM   1428  CG  LYS A  88       1.596  43.972 -30.098  1.00  0.00           C  
ATOM   1429  CD  LYS A  88       2.741  44.758 -29.477  1.00  0.00           C  
ATOM   1430  CE  LYS A  88       2.249  46.055 -28.853  1.00  0.00           C  
ATOM   1431  NZ  LYS A  88       3.364  46.853 -28.276  1.00  0.00           N1+
ATOM   1432  H   LYS A  88       1.466  43.268 -33.091  1.00  0.00           H  
ATOM   1433  HA  LYS A  88       3.966  43.278 -31.483  1.00  0.00           H  
ATOM   1434 1HB  LYS A  88       1.196  42.068 -30.960  1.00  0.00           H  
ATOM   1435 2HB  LYS A  88       2.522  42.079 -29.805  1.00  0.00           H  
ATOM   1436 1HG  LYS A  88       1.165  44.556 -30.912  1.00  0.00           H  
ATOM   1437 2HG  LYS A  88       0.826  43.811 -29.344  1.00  0.00           H  
ATOM   1438 1HD  LYS A  88       3.222  44.153 -28.707  1.00  0.00           H  
ATOM   1439 2HD  LYS A  88       3.479  44.992 -30.244  1.00  0.00           H  
ATOM   1440 1HE  LYS A  88       1.744  46.654 -29.610  1.00  0.00           H  
ATOM   1441 2HE  LYS A  88       1.533  45.830 -28.063  1.00  0.00           H  
ATOM   1442 1HZ  LYS A  88       2.997  47.704 -27.873  1.00  0.00           H  
ATOM   1443 2HZ  LYS A  88       3.829  46.314 -27.559  1.00  0.00           H  
ATOM   1444 3HZ  LYS A  88       4.025  47.085 -29.004  1.00  0.00           H  
ATOM   1445  N   SER A  89       3.497  40.856 -33.290  1.00  0.00           N  
ATOM   1446  CA  SER A  89       4.000  39.532 -33.647  1.00  0.00           C  
ATOM   1447  C   SER A  89       5.154  39.642 -34.637  1.00  0.00           C  
ATOM   1448  O   SER A  89       5.159  40.529 -35.488  1.00  0.00           O  
ATOM   1449  CB  SER A  89       2.888  38.690 -34.241  1.00  0.00           C  
ATOM   1450  OG  SER A  89       1.876  38.458 -33.300  1.00  0.00           O  
ATOM   1451  H   SER A  89       3.039  41.415 -33.996  1.00  0.00           H  
ATOM   1452  HA  SER A  89       4.320  39.026 -32.735  1.00  0.00           H  
ATOM   1453 1HB  SER A  89       2.471  39.198 -35.110  1.00  0.00           H  
ATOM   1454 2HB  SER A  89       3.295  37.739 -34.583  1.00  0.00           H  
ATOM   1455  HG  SER A  89       1.075  38.833 -33.674  1.00  0.00           H  
ATOM   1456  N   PRO A  90       6.160  38.757 -34.521  1.00  0.00           N  
ATOM   1457  CA  PRO A  90       7.277  38.818 -35.470  1.00  0.00           C  
ATOM   1458  C   PRO A  90       6.662  38.542 -36.844  1.00  0.00           C  
ATOM   1459  O   PRO A  90       5.944  37.558 -37.021  1.00  0.00           O  
ATOM   1460  CB  PRO A  90       8.229  37.707 -35.014  1.00  0.00           C  
ATOM   1461  CG  PRO A  90       7.896  37.508 -33.575  1.00  0.00           C  
ATOM   1462  CD  PRO A  90       6.407  37.718 -33.503  1.00  0.00           C  
ATOM   1463  HA  PRO A  90       7.788  39.786 -35.366  1.00  0.00           H  
ATOM   1464 1HB  PRO A  90       8.068  36.803 -35.619  1.00  0.00           H  
ATOM   1465 2HB  PRO A  90       9.272  38.018 -35.172  1.00  0.00           H  
ATOM   1466 1HG  PRO A  90       8.197  36.500 -33.252  1.00  0.00           H  
ATOM   1467 2HG  PRO A  90       8.455  38.222 -32.953  1.00  0.00           H  
ATOM   1468 1HD  PRO A  90       5.891  36.776 -33.738  1.00  0.00           H  
ATOM   1469 2HD  PRO A  90       6.134  38.069 -32.497  1.00  0.00           H  
ATOM   1470  N   PRO A  91       6.920  39.409 -37.832  1.00  0.00           N  
ATOM   1471  CA  PRO A  91       6.343  39.183 -39.160  1.00  0.00           C  
ATOM   1472  C   PRO A  91       6.529  37.786 -39.753  1.00  0.00           C  
ATOM   1473  O   PRO A  91       5.612  37.251 -40.375  1.00  0.00           O  
ATOM   1474  CB  PRO A  91       7.083  40.228 -40.001  1.00  0.00           C  
ATOM   1475  CG  PRO A  91       7.353  41.341 -39.046  1.00  0.00           C  
ATOM   1476  CD  PRO A  91       7.723  40.649 -37.760  1.00  0.00           C  
ATOM   1477  HA  PRO A  91       5.253  39.326 -39.110  1.00  0.00           H  
ATOM   1478 1HB  PRO A  91       8.002  39.792 -40.419  1.00  0.00           H  
ATOM   1479 2HB  PRO A  91       6.458  40.538 -40.851  1.00  0.00           H  
ATOM   1480 1HG  PRO A  91       8.160  41.982 -39.429  1.00  0.00           H  
ATOM   1481 2HG  PRO A  91       6.463  41.978 -38.943  1.00  0.00           H  
ATOM   1482 1HD  PRO A  91       8.800  40.427 -37.760  1.00  0.00           H  
ATOM   1483 2HD  PRO A  91       7.459  41.293 -36.908  1.00  0.00           H  
ATOM   1484  N   HIS A  92       7.703  37.189 -39.561  1.00  0.00           N  
ATOM   1485  CA  HIS A  92       7.951  35.867 -40.122  1.00  0.00           C  
ATOM   1486  C   HIS A  92       7.114  34.767 -39.485  1.00  0.00           C  
ATOM   1487  O   HIS A  92       7.041  33.657 -40.008  1.00  0.00           O  
ATOM   1488  CB  HIS A  92       9.443  35.499 -40.048  1.00  0.00           C  
ATOM   1489  CG  HIS A  92       9.931  35.149 -38.675  1.00  0.00           C  
ATOM   1490  CD2 HIS A  92      10.248  33.954 -38.125  1.00  0.00           C  
ATOM   1491  ND1 HIS A  92      10.175  36.095 -37.702  1.00  0.00           N  
ATOM   1492  CE1 HIS A  92      10.626  35.497 -36.614  1.00  0.00           C  
ATOM   1493  NE2 HIS A  92      10.680  34.198 -36.843  1.00  0.00           N  
ATOM   1494  H   HIS A  92       8.432  37.642 -39.028  1.00  0.00           H  
ATOM   1495  HA  HIS A  92       7.658  35.856 -41.172  1.00  0.00           H  
ATOM   1496 1HB  HIS A  92       9.640  34.646 -40.699  1.00  0.00           H  
ATOM   1497 2HB  HIS A  92      10.042  36.333 -40.411  1.00  0.00           H  
ATOM   1498  HD2 HIS A  92      10.177  32.980 -38.610  1.00  0.00           H  
ATOM   1499  HE1 HIS A  92      10.905  35.992 -35.683  1.00  0.00           H  
ATOM   1500  HE2 HIS A  92      10.988  33.494 -36.187  1.00  0.00           H  
ATOM   1501  N   LEU A  93       6.470  35.075 -38.365  1.00  0.00           N  
ATOM   1502  CA  LEU A  93       5.636  34.084 -37.693  1.00  0.00           C  
ATOM   1503  C   LEU A  93       4.159  34.226 -38.046  1.00  0.00           C  
ATOM   1504  O   LEU A  93       3.346  33.376 -37.683  1.00  0.00           O  
ATOM   1505  CB  LEU A  93       5.832  34.164 -36.177  1.00  0.00           C  
ATOM   1506  CG  LEU A  93       7.233  33.763 -35.705  1.00  0.00           C  
ATOM   1507  CD1 LEU A  93       7.324  33.902 -34.197  1.00  0.00           C  
ATOM   1508  CD2 LEU A  93       7.532  32.329 -36.132  1.00  0.00           C  
ATOM   1509  H   LEU A  93       6.551  36.000 -37.967  1.00  0.00           H  
ATOM   1510  HA  LEU A  93       5.937  33.093 -38.032  1.00  0.00           H  
ATOM   1511 1HB  LEU A  93       5.637  35.186 -35.856  1.00  0.00           H  
ATOM   1512 2HB  LEU A  93       5.104  33.510 -35.698  1.00  0.00           H  
ATOM   1513  HG  LEU A  93       7.972  34.432 -36.147  1.00  0.00           H  
ATOM   1514 1HD1 LEU A  93       8.323  33.616 -33.865  1.00  0.00           H  
ATOM   1515 2HD1 LEU A  93       7.132  34.937 -33.916  1.00  0.00           H  
ATOM   1516 3HD1 LEU A  93       6.586  33.254 -33.727  1.00  0.00           H  
ATOM   1517 1HD2 LEU A  93       8.530  32.048 -35.794  1.00  0.00           H  
ATOM   1518 2HD2 LEU A  93       6.797  31.656 -35.690  1.00  0.00           H  
ATOM   1519 3HD2 LEU A  93       7.483  32.255 -37.219  1.00  0.00           H  
ATOM   1520  N   ILE A  94       3.810  35.301 -38.745  1.00  0.00           N  
ATOM   1521  CA  ILE A  94       2.427  35.502 -39.163  1.00  0.00           C  
ATOM   1522  C   ILE A  94       2.347  35.159 -40.647  1.00  0.00           C  
ATOM   1523  O   ILE A  94       2.728  35.958 -41.505  1.00  0.00           O  
ATOM   1524  CB  ILE A  94       1.962  36.965 -38.949  1.00  0.00           C  
ATOM   1525  CG1 ILE A  94       2.047  37.324 -37.467  1.00  0.00           C  
ATOM   1526  CG2 ILE A  94       0.533  37.143 -39.458  1.00  0.00           C  
ATOM   1527  CD1 ILE A  94       1.221  36.425 -36.547  1.00  0.00           C  
ATOM   1528  H   ILE A  94       4.505  35.991 -38.993  1.00  0.00           H  
ATOM   1529  HA  ILE A  94       1.787  34.856 -38.564  1.00  0.00           H  
ATOM   1530  HB  ILE A  94       2.621  37.639 -39.494  1.00  0.00           H  
ATOM   1531 1HG1 ILE A  94       3.085  37.273 -37.140  1.00  0.00           H  
ATOM   1532 2HG1 ILE A  94       1.709  38.350 -37.321  1.00  0.00           H  
ATOM   1533 1HG2 ILE A  94       0.217  38.174 -39.301  1.00  0.00           H  
ATOM   1534 2HG2 ILE A  94       0.494  36.908 -40.521  1.00  0.00           H  
ATOM   1535 3HG2 ILE A  94      -0.133  36.472 -38.914  1.00  0.00           H  
ATOM   1536 1HD1 ILE A  94       1.341  36.753 -35.513  1.00  0.00           H  
ATOM   1537 2HD1 ILE A  94       0.169  36.486 -36.827  1.00  0.00           H  
ATOM   1538 3HD1 ILE A  94       1.562  35.395 -36.641  1.00  0.00           H  
ATOM   1539  N   LYS A  95       1.875  33.953 -40.944  1.00  0.00           N  
ATOM   1540  CA  LYS A  95       1.756  33.520 -42.330  1.00  0.00           C  
ATOM   1541  C   LYS A  95       0.636  34.294 -42.999  1.00  0.00           C  
ATOM   1542  O   LYS A  95       0.750  34.718 -44.151  1.00  0.00           O  
ATOM   1543  CB  LYS A  95       1.496  32.015 -42.418  1.00  0.00           C  
ATOM   1544  CG  LYS A  95       1.421  31.472 -43.839  1.00  0.00           C  
ATOM   1545  CD  LYS A  95       2.763  31.588 -44.546  1.00  0.00           C  
ATOM   1546  CE  LYS A  95       2.718  30.953 -45.928  1.00  0.00           C  
ATOM   1547  NZ  LYS A  95       4.014  31.091 -46.646  1.00  0.00           N1+
ATOM   1548  H   LYS A  95       1.594  33.324 -40.206  1.00  0.00           H  
ATOM   1549  HA  LYS A  95       2.704  33.711 -42.835  1.00  0.00           H  
ATOM   1550 1HB  LYS A  95       2.288  31.478 -41.895  1.00  0.00           H  
ATOM   1551 2HB  LYS A  95       0.556  31.779 -41.919  1.00  0.00           H  
ATOM   1552 1HG  LYS A  95       1.124  30.423 -43.812  1.00  0.00           H  
ATOM   1553 2HG  LYS A  95       0.673  32.029 -44.402  1.00  0.00           H  
ATOM   1554 1HD  LYS A  95       3.031  32.641 -44.648  1.00  0.00           H  
ATOM   1555 2HD  LYS A  95       3.531  31.092 -43.953  1.00  0.00           H  
ATOM   1556 1HE  LYS A  95       2.480  29.895 -45.834  1.00  0.00           H  
ATOM   1557 2HE  LYS A  95       1.936  31.427 -46.521  1.00  0.00           H  
ATOM   1558 1HZ  LYS A  95       3.943  30.658 -47.556  1.00  0.00           H  
ATOM   1559 2HZ  LYS A  95       4.236  32.071 -46.756  1.00  0.00           H  
ATOM   1560 3HZ  LYS A  95       4.744  30.638 -46.115  1.00  0.00           H  
ATOM   1561  N   GLU A  96      -0.446  34.492 -42.254  1.00  0.00           N  
ATOM   1562  CA  GLU A  96      -1.609  35.188 -42.775  1.00  0.00           C  
ATOM   1563  C   GLU A  96      -2.603  35.409 -41.643  1.00  0.00           C  
ATOM   1564  O   GLU A  96      -2.511  34.770 -40.593  1.00  0.00           O  
ATOM   1565  CB  GLU A  96      -2.275  34.318 -43.844  1.00  0.00           C  
ATOM   1566  CG  GLU A  96      -2.944  33.072 -43.255  1.00  0.00           C  
ATOM   1567  CD  GLU A  96      -3.235  31.986 -44.280  1.00  0.00           C  
ATOM   1568  OE1 GLU A  96      -3.490  32.317 -45.458  1.00  0.00           O  
ATOM   1569  OE2 GLU A  96      -3.226  30.797 -43.897  1.00  0.00           O1-
ATOM   1570  H   GLU A  96      -0.462  34.150 -41.304  1.00  0.00           H  
ATOM   1571  HA  GLU A  96      -1.278  36.125 -43.226  1.00  0.00           H  
ATOM   1572 1HB  GLU A  96      -3.026  34.904 -44.374  1.00  0.00           H  
ATOM   1573 2HB  GLU A  96      -1.529  34.005 -44.575  1.00  0.00           H  
ATOM   1574 1HG  GLU A  96      -2.295  32.653 -42.486  1.00  0.00           H  
ATOM   1575 2HG  GLU A  96      -3.879  33.365 -42.780  1.00  0.00           H  
ATOM   1576  N   ILE A  97      -3.537  36.327 -41.859  1.00  0.00           N  
ATOM   1577  CA  ILE A  97      -4.602  36.588 -40.902  1.00  0.00           C  
ATOM   1578  C   ILE A  97      -5.875  36.340 -41.702  1.00  0.00           C  
ATOM   1579  O   ILE A  97      -6.169  37.061 -42.655  1.00  0.00           O  
ATOM   1580  CB  ILE A  97      -4.563  38.020 -40.337  1.00  0.00           C  
ATOM   1581  CG1 ILE A  97      -3.226  38.282 -39.639  1.00  0.00           C  
ATOM   1582  CG2 ILE A  97      -5.721  38.244 -39.377  1.00  0.00           C  
ATOM   1583  CD1 ILE A  97      -3.026  39.721 -39.223  1.00  0.00           C  
ATOM   1584  H   ILE A  97      -3.505  36.859 -42.717  1.00  0.00           H  
ATOM   1585  HA  ILE A  97      -4.460  35.937 -40.040  1.00  0.00           H  
ATOM   1586  HB  ILE A  97      -4.637  38.736 -41.154  1.00  0.00           H  
ATOM   1587 1HG1 ILE A  97      -3.151  37.656 -38.750  1.00  0.00           H  
ATOM   1588 2HG1 ILE A  97      -2.408  38.003 -40.303  1.00  0.00           H  
ATOM   1589 1HG2 ILE A  97      -5.678  39.260 -38.986  1.00  0.00           H  
ATOM   1590 2HG2 ILE A  97      -6.663  38.097 -39.903  1.00  0.00           H  
ATOM   1591 3HG2 ILE A  97      -5.652  37.535 -38.551  1.00  0.00           H  
ATOM   1592 1HD1 ILE A  97      -2.056  39.828 -38.736  1.00  0.00           H  
ATOM   1593 2HD1 ILE A  97      -3.062  40.363 -40.104  1.00  0.00           H  
ATOM   1594 3HD1 ILE A  97      -3.814  40.012 -38.529  1.00  0.00           H  
ATOM   1595  N   ILE A  98      -6.610  35.295 -41.338  1.00  0.00           N  
ATOM   1596  CA  ILE A  98      -7.839  34.954 -42.034  1.00  0.00           C  
ATOM   1597  C   ILE A  98      -9.039  35.489 -41.269  1.00  0.00           C  
ATOM   1598  O   ILE A  98      -9.298  35.074 -40.141  1.00  0.00           O  
ATOM   1599  CB  ILE A  98      -7.984  33.421 -42.182  1.00  0.00           C  
ATOM   1600  CG1 ILE A  98      -6.789  32.866 -42.959  1.00  0.00           C  
ATOM   1601  CG2 ILE A  98      -9.295  33.080 -42.900  1.00  0.00           C  
ATOM   1602  CD1 ILE A  98      -6.739  31.335 -43.004  1.00  0.00           C  
ATOM   1603  H   ILE A  98      -6.311  34.725 -40.560  1.00  0.00           H  
ATOM   1604  HA  ILE A  98      -7.805  35.394 -43.030  1.00  0.00           H  
ATOM   1605  HB  ILE A  98      -7.987  32.959 -41.195  1.00  0.00           H  
ATOM   1606 1HG1 ILE A  98      -6.818  33.237 -43.983  1.00  0.00           H  
ATOM   1607 2HG1 ILE A  98      -5.864  33.224 -42.508  1.00  0.00           H  
ATOM   1608 1HG2 ILE A  98      -9.386  31.999 -42.999  1.00  0.00           H  
ATOM   1609 2HG2 ILE A  98     -10.135  33.464 -42.324  1.00  0.00           H  
ATOM   1610 3HG2 ILE A  98      -9.296  33.536 -43.891  1.00  0.00           H  
ATOM   1611 1HD1 ILE A  98      -5.865  31.016 -43.572  1.00  0.00           H  
ATOM   1612 2HD1 ILE A  98      -6.675  30.943 -41.989  1.00  0.00           H  
ATOM   1613 3HD1 ILE A  98      -7.640  30.956 -43.483  1.00  0.00           H  
ATOM   1614  N   LEU A  99      -9.758  36.428 -41.877  1.00  0.00           N  
ATOM   1615  CA  LEU A  99     -10.947  36.986 -41.243  1.00  0.00           C  
ATOM   1616  C   LEU A  99     -12.163  36.279 -41.815  1.00  0.00           C  
ATOM   1617  O   LEU A  99     -12.427  36.369 -43.014  1.00  0.00           O  
ATOM   1618  CB  LEU A  99     -11.045  38.498 -41.484  1.00  0.00           C  
ATOM   1619  CG  LEU A  99      -9.862  39.330 -40.974  1.00  0.00           C  
ATOM   1620  CD1 LEU A  99     -10.071  40.792 -41.347  1.00  0.00           C  
ATOM   1621  CD2 LEU A  99      -9.735  39.163 -39.467  1.00  0.00           C  
ATOM   1622  H   LEU A  99      -9.482  36.764 -42.788  1.00  0.00           H  
ATOM   1623  HA  LEU A  99     -10.864  36.843 -40.167  1.00  0.00           H  
ATOM   1624 1HB  LEU A  99     -11.136  38.674 -42.554  1.00  0.00           H  
ATOM   1625 2HB  LEU A  99     -11.946  38.870 -40.997  1.00  0.00           H  
ATOM   1626  HG  LEU A  99      -8.944  38.992 -41.455  1.00  0.00           H  
ATOM   1627 1HD1 LEU A  99      -9.230  41.384 -40.984  1.00  0.00           H  
ATOM   1628 2HD1 LEU A  99     -10.139  40.885 -42.430  1.00  0.00           H  
ATOM   1629 3HD1 LEU A  99     -10.992  41.155 -40.892  1.00  0.00           H  
ATOM   1630 1HD2 LEU A  99      -8.893  39.754 -39.105  1.00  0.00           H  
ATOM   1631 2HD2 LEU A  99     -10.651  39.503 -38.985  1.00  0.00           H  
ATOM   1632 3HD2 LEU A  99      -9.568  38.112 -39.231  1.00  0.00           H  
ATOM   1633  N   VAL A 100     -12.887  35.550 -40.971  1.00  0.00           N  
ATOM   1634  CA  VAL A 100     -14.083  34.860 -41.429  1.00  0.00           C  
ATOM   1635  C   VAL A 100     -15.288  35.708 -41.048  1.00  0.00           C  
ATOM   1636  O   VAL A 100     -15.693  35.765 -39.887  1.00  0.00           O  
ATOM   1637  CB  VAL A 100     -14.196  33.461 -40.795  1.00  0.00           C  
ATOM   1638  CG1 VAL A 100     -15.451  32.754 -41.283  1.00  0.00           C  
ATOM   1639  CG2 VAL A 100     -12.955  32.644 -41.120  1.00  0.00           C  
ATOM   1640  H   VAL A 100     -12.613  35.470 -40.003  1.00  0.00           H  
ATOM   1641  HA  VAL A 100     -14.024  34.745 -42.512  1.00  0.00           H  
ATOM   1642  HB  VAL A 100     -14.288  33.569 -39.714  1.00  0.00           H  
ATOM   1643 1HG1 VAL A 100     -15.515  31.767 -40.825  1.00  0.00           H  
ATOM   1644 2HG1 VAL A 100     -16.328  33.339 -41.008  1.00  0.00           H  
ATOM   1645 3HG1 VAL A 100     -15.410  32.648 -42.367  1.00  0.00           H  
ATOM   1646 1HG2 VAL A 100     -13.041  31.656 -40.668  1.00  0.00           H  
ATOM   1647 2HG2 VAL A 100     -12.860  32.541 -42.201  1.00  0.00           H  
ATOM   1648 3HG2 VAL A 100     -12.073  33.148 -40.725  1.00  0.00           H  
ATOM   1649  N   ASP A 101     -15.834  36.391 -42.046  1.00  0.00           N  
ATOM   1650  CA  ASP A 101     -16.992  37.255 -41.864  1.00  0.00           C  
ATOM   1651  C   ASP A 101     -18.225  36.365 -41.803  1.00  0.00           C  
ATOM   1652  O   ASP A 101     -18.710  35.903 -42.835  1.00  0.00           O  
ATOM   1653  CB  ASP A 101     -17.089  38.215 -43.052  1.00  0.00           C  
ATOM   1654  CG  ASP A 101     -18.327  39.074 -43.010  1.00  0.00           C  
ATOM   1655  OD1 ASP A 101     -18.543  39.837 -43.976  1.00  0.00           O  
ATOM   1656  OD2 ASP A 101     -19.080  38.987 -42.018  1.00  0.00           O1-
ATOM   1657  H   ASP A 101     -15.426  36.304 -42.965  1.00  0.00           H  
ATOM   1658  HA  ASP A 101     -16.856  37.828 -40.947  1.00  0.00           H  
ATOM   1659 1HB  ASP A 101     -16.213  38.865 -43.067  1.00  0.00           H  
ATOM   1660 2HB  ASP A 101     -17.090  37.645 -43.981  1.00  0.00           H  
ATOM   1661  N   ASP A 102     -18.725  36.120 -40.596  1.00  0.00           N  
ATOM   1662  CA  ASP A 102     -19.881  35.252 -40.433  1.00  0.00           C  
ATOM   1663  C   ASP A 102     -21.197  35.942 -40.769  1.00  0.00           C  
ATOM   1664  O   ASP A 102     -22.012  36.215 -39.888  1.00  0.00           O  
ATOM   1665  CB  ASP A 102     -19.942  34.700 -39.007  1.00  0.00           C  
ATOM   1666  CG  ASP A 102     -20.980  33.600 -38.857  1.00  0.00           C  
ATOM   1667  OD1 ASP A 102     -21.375  33.305 -37.713  1.00  0.00           O  
ATOM   1668  OD2 ASP A 102     -21.391  33.024 -39.889  1.00  0.00           O1-
ATOM   1669  H   ASP A 102     -18.304  36.538 -39.778  1.00  0.00           H  
ATOM   1670  HA  ASP A 102     -19.786  34.415 -41.126  1.00  0.00           H  
ATOM   1671 1HB  ASP A 102     -18.966  34.305 -38.727  1.00  0.00           H  
ATOM   1672 2HB  ASP A 102     -20.180  35.507 -38.315  1.00  0.00           H  
ATOM   1673  N   TYR A 103     -21.388  36.226 -42.053  1.00  0.00           N  
ATOM   1674  CA  TYR A 103     -22.611  36.849 -42.546  1.00  0.00           C  
ATOM   1675  C   TYR A 103     -22.981  38.150 -41.820  1.00  0.00           C  
ATOM   1676  O   TYR A 103     -24.119  38.328 -41.376  1.00  0.00           O  
ATOM   1677  CB  TYR A 103     -23.768  35.853 -42.446  1.00  0.00           C  
ATOM   1678  CG  TYR A 103     -25.085  36.393 -42.960  1.00  0.00           C  
ATOM   1679  CD1 TYR A 103     -25.324  36.453 -44.325  1.00  0.00           C  
ATOM   1680  CD2 TYR A 103     -26.052  36.828 -42.066  1.00  0.00           C  
ATOM   1681  CE1 TYR A 103     -26.526  36.946 -44.794  1.00  0.00           C  
ATOM   1682  CE2 TYR A 103     -27.254  37.321 -42.536  1.00  0.00           C  
ATOM   1683  CZ  TYR A 103     -27.492  37.381 -43.894  1.00  0.00           C  
ATOM   1684  OH  TYR A 103     -28.689  37.873 -44.361  1.00  0.00           O  
ATOM   1685  H   TYR A 103     -20.653  35.998 -42.708  1.00  0.00           H  
ATOM   1686  HA  TYR A 103     -22.461  37.125 -43.590  1.00  0.00           H  
ATOM   1687 1HB  TYR A 103     -23.525  34.953 -43.014  1.00  0.00           H  
ATOM   1688 2HB  TYR A 103     -23.905  35.557 -41.407  1.00  0.00           H  
ATOM   1689  HD1 TYR A 103     -24.564  36.111 -45.028  1.00  0.00           H  
ATOM   1690  HD2 TYR A 103     -25.863  36.782 -40.994  1.00  0.00           H  
ATOM   1691  HE1 TYR A 103     -26.715  36.994 -45.867  1.00  0.00           H  
ATOM   1692  HE2 TYR A 103     -28.014  37.664 -41.833  1.00  0.00           H  
ATOM   1693  HH  TYR A 103     -29.241  38.128 -43.618  1.00  0.00           H  
ATOM   1694  N   SER A 104     -22.018  39.059 -41.704  1.00  0.00           N  
ATOM   1695  CA  SER A 104     -22.270  40.340 -41.055  1.00  0.00           C  
ATOM   1696  C   SER A 104     -23.253  41.148 -41.898  1.00  0.00           C  
ATOM   1697  O   SER A 104     -23.341  40.969 -43.113  1.00  0.00           O  
ATOM   1698  CB  SER A 104     -20.976  41.109 -40.871  1.00  0.00           C  
ATOM   1699  OG  SER A 104     -20.100  40.423 -40.020  1.00  0.00           O  
ATOM   1700  H   SER A 104     -21.097  38.865 -42.069  1.00  0.00           H  
ATOM   1701  HA  SER A 104     -22.684  40.151 -40.064  1.00  0.00           H  
ATOM   1702 1HB  SER A 104     -20.502  41.260 -41.841  1.00  0.00           H  
ATOM   1703 2HB  SER A 104     -21.194  42.093 -40.457  1.00  0.00           H  
ATOM   1704  HG  SER A 104     -19.685  39.745 -40.559  1.00  0.00           H  
ATOM   1705  N   ASN A 105     -23.980  42.043 -41.238  1.00  0.00           N  
ATOM   1706  CA  ASN A 105     -24.968  42.894 -41.896  1.00  0.00           C  
ATOM   1707  C   ASN A 105     -24.377  43.684 -43.063  1.00  0.00           C  
ATOM   1708  O   ASN A 105     -24.978  43.774 -44.134  1.00  0.00           O  
ATOM   1709  CB  ASN A 105     -25.600  43.837 -40.889  1.00  0.00           C  
ATOM   1710  CG  ASN A 105     -26.552  43.136 -39.959  1.00  0.00           C  
ATOM   1711  ND2 ASN A 105     -26.831  43.750 -38.838  1.00  0.00           N  
ATOM   1712  OD1 ASN A 105     -27.033  42.037 -40.257  1.00  0.00           O  
ATOM   1713  H   ASN A 105     -23.839  42.132 -40.242  1.00  0.00           H  
ATOM   1714  HA  ASN A 105     -25.763  42.260 -42.292  1.00  0.00           H  
ATOM   1715 1HB  ASN A 105     -24.818  44.315 -40.297  1.00  0.00           H  
ATOM   1716 2HB  ASN A 105     -26.139  44.624 -41.416  1.00  0.00           H  
ATOM   1717 1HD2 ASN A 105     -27.459  43.331 -38.181  1.00  0.00           H  
ATOM   1718 2HD2 ASN A 105     -26.419  44.638 -38.638  1.00  0.00           H  
ATOM   1719  N   ASP A 106     -23.194  44.251 -42.848  1.00  0.00           N  
ATOM   1720  CA  ASP A 106     -22.516  45.050 -43.867  1.00  0.00           C  
ATOM   1721  C   ASP A 106     -21.234  44.350 -44.317  1.00  0.00           C  
ATOM   1722  O   ASP A 106     -20.262  44.280 -43.564  1.00  0.00           O  
ATOM   1723  CB  ASP A 106     -22.190  46.447 -43.334  1.00  0.00           C  
ATOM   1724  CG  ASP A 106     -21.540  47.343 -44.380  1.00  0.00           C  
ATOM   1725  OD1 ASP A 106     -21.230  46.856 -45.441  1.00  0.00           O  
ATOM   1726  OD2 ASP A 106     -21.359  48.506 -44.107  1.00  0.00           O1-
ATOM   1727  H   ASP A 106     -22.752  44.124 -41.949  1.00  0.00           H  
ATOM   1728  HA  ASP A 106     -23.187  45.165 -44.719  1.00  0.00           H  
ATOM   1729 1HB  ASP A 106     -23.105  46.926 -42.985  1.00  0.00           H  
ATOM   1730 2HB  ASP A 106     -21.518  46.364 -42.480  1.00  0.00           H  
ATOM   1731  N   PRO A 107     -21.216  43.827 -45.555  1.00  0.00           N  
ATOM   1732  CA  PRO A 107     -20.036  43.130 -46.081  1.00  0.00           C  
ATOM   1733  C   PRO A 107     -18.765  43.976 -46.103  1.00  0.00           C  
ATOM   1734  O   PRO A 107     -17.661  43.442 -46.156  1.00  0.00           O  
ATOM   1735  CB  PRO A 107     -20.478  42.768 -47.502  1.00  0.00           C  
ATOM   1736  CG  PRO A 107     -21.966  42.725 -47.423  1.00  0.00           C  
ATOM   1737  CD  PRO A 107     -22.321  43.825 -46.458  1.00  0.00           C  
ATOM   1738  HA  PRO A 107     -19.835  42.242 -45.463  1.00  0.00           H  
ATOM   1739 1HB  PRO A 107     -20.110  43.522 -48.214  1.00  0.00           H  
ATOM   1740 2HB  PRO A 107     -20.039  41.805 -47.799  1.00  0.00           H  
ATOM   1741 1HG  PRO A 107     -22.404  42.878 -48.420  1.00  0.00           H  
ATOM   1742 2HG  PRO A 107     -22.299  41.735 -47.078  1.00  0.00           H  
ATOM   1743 1HD  PRO A 107     -22.409  44.777 -47.003  1.00  0.00           H  
ATOM   1744 2HD  PRO A 107     -23.266  43.579 -45.952  1.00  0.00           H  
ATOM   1745  N   GLU A 108     -18.920  45.295 -46.066  1.00  0.00           N  
ATOM   1746  CA  GLU A 108     -17.766  46.181 -46.103  1.00  0.00           C  
ATOM   1747  C   GLU A 108     -16.903  46.142 -44.844  1.00  0.00           C  
ATOM   1748  O   GLU A 108     -15.711  46.441 -44.906  1.00  0.00           O  
ATOM   1749  CB  GLU A 108     -18.235  47.617 -46.346  1.00  0.00           C  
ATOM   1750  CG  GLU A 108     -18.835  47.858 -47.725  1.00  0.00           C  
ATOM   1751  CD  GLU A 108     -17.844  47.654 -48.837  1.00  0.00           C  
ATOM   1752  OE1 GLU A 108     -16.775  48.210 -48.765  1.00  0.00           O  
ATOM   1753  OE2 GLU A 108     -18.157  46.940 -49.761  1.00  0.00           O1-
ATOM   1754  H   GLU A 108     -19.847  45.695 -46.012  1.00  0.00           H  
ATOM   1755  HA  GLU A 108     -17.100  45.848 -46.900  1.00  0.00           H  
ATOM   1756 1HB  GLU A 108     -18.986  47.885 -45.603  1.00  0.00           H  
ATOM   1757 2HB  GLU A 108     -17.395  48.300 -46.223  1.00  0.00           H  
ATOM   1758 1HG  GLU A 108     -19.672  47.175 -47.869  1.00  0.00           H  
ATOM   1759 2HG  GLU A 108     -19.220  48.875 -47.769  1.00  0.00           H  
ATOM   1760  N   ASP A 109     -17.492  45.766 -43.710  1.00  0.00           N  
ATOM   1761  CA  ASP A 109     -16.730  45.691 -42.461  1.00  0.00           C  
ATOM   1762  C   ASP A 109     -15.488  44.839 -42.702  1.00  0.00           C  
ATOM   1763  O   ASP A 109     -14.377  45.207 -42.329  1.00  0.00           O  
ATOM   1764  CB  ASP A 109     -17.558  45.046 -41.343  1.00  0.00           C  
ATOM   1765  CG  ASP A 109     -18.773  45.868 -40.946  1.00  0.00           C  
ATOM   1766  OD1 ASP A 109     -18.945  46.998 -41.449  1.00  0.00           O  
ATOM   1767  OD2 ASP A 109     -19.562  45.375 -40.112  1.00  0.00           O1-
ATOM   1768  H   ASP A 109     -18.474  45.528 -43.703  1.00  0.00           H  
ATOM   1769  HA  ASP A 109     -16.471  46.704 -42.153  1.00  0.00           H  
ATOM   1770 1HB  ASP A 109     -17.897  44.061 -41.664  1.00  0.00           H  
ATOM   1771 2HB  ASP A 109     -16.932  44.905 -40.461  1.00  0.00           H  
ATOM   1772  N   GLY A 110     -15.685  43.696 -43.347  1.00  0.00           N  
ATOM   1773  CA  GLY A 110     -14.565  42.816 -43.619  1.00  0.00           C  
ATOM   1774  C   GLY A 110     -13.823  43.120 -44.908  1.00  0.00           C  
ATOM   1775  O   GLY A 110     -12.595  43.145 -44.930  1.00  0.00           O  
ATOM   1776  H   GLY A 110     -16.608  43.426 -43.653  1.00  0.00           H  
ATOM   1777 1HA  GLY A 110     -13.849  42.873 -42.798  1.00  0.00           H  
ATOM   1778 2HA  GLY A 110     -14.916  41.785 -43.669  1.00  0.00           H  
ATOM   1779  N   ALA A 111     -14.567  43.363 -45.982  1.00  0.00           N  
ATOM   1780  CA  ALA A 111     -13.963  43.629 -47.284  1.00  0.00           C  
ATOM   1781  C   ALA A 111     -12.949  44.770 -47.287  1.00  0.00           C  
ATOM   1782  O   ALA A 111     -11.911  44.687 -47.946  1.00  0.00           O  
ATOM   1783  CB  ALA A 111     -15.055  43.905 -48.307  1.00  0.00           C  
ATOM   1784  H   ALA A 111     -15.574  43.363 -45.898  1.00  0.00           H  
ATOM   1785  HA  ALA A 111     -13.404  42.744 -47.586  1.00  0.00           H  
ATOM   1786 1HB  ALA A 111     -14.601  44.104 -49.279  1.00  0.00           H  
ATOM   1787 2HB  ALA A 111     -15.709  43.037 -48.384  1.00  0.00           H  
ATOM   1788 3HB  ALA A 111     -15.636  44.772 -47.994  1.00  0.00           H  
ATOM   1789  N   LEU A 112     -13.248  45.838 -46.559  1.00  0.00           N  
ATOM   1790  CA  LEU A 112     -12.346  46.978 -46.524  1.00  0.00           C  
ATOM   1791  C   LEU A 112     -11.010  46.642 -45.869  1.00  0.00           C  
ATOM   1792  O   LEU A 112      -9.987  47.248 -46.188  1.00  0.00           O  
ATOM   1793  CB  LEU A 112     -13.011  48.155 -45.807  1.00  0.00           C  
ATOM   1794  CG  LEU A 112     -14.192  48.789 -46.553  1.00  0.00           C  
ATOM   1795  CD1 LEU A 112     -14.872  49.828 -45.674  1.00  0.00           C  
ATOM   1796  CD2 LEU A 112     -13.689  49.417 -47.844  1.00  0.00           C  
ATOM   1797  H   LEU A 112     -14.103  45.871 -46.023  1.00  0.00           H  
ATOM   1798  HA  LEU A 112     -12.120  47.273 -47.549  1.00  0.00           H  
ATOM   1799 1HB  LEU A 112     -13.371  47.813 -44.838  1.00  0.00           H  
ATOM   1800 2HB  LEU A 112     -12.262  48.930 -45.642  1.00  0.00           H  
ATOM   1801  HG  LEU A 112     -14.930  48.021 -46.784  1.00  0.00           H  
ATOM   1802 1HD1 LEU A 112     -15.709  50.270 -46.215  1.00  0.00           H  
ATOM   1803 2HD1 LEU A 112     -15.240  49.352 -44.766  1.00  0.00           H  
ATOM   1804 3HD1 LEU A 112     -14.157  50.607 -45.412  1.00  0.00           H  
ATOM   1805 1HD2 LEU A 112     -14.525  49.869 -48.379  1.00  0.00           H  
ATOM   1806 2HD2 LEU A 112     -12.951  50.185 -47.612  1.00  0.00           H  
ATOM   1807 3HD2 LEU A 112     -13.232  48.650 -48.469  1.00  0.00           H  
ATOM   1808  N   LEU A 113     -11.011  45.669 -44.961  1.00  0.00           N  
ATOM   1809  CA  LEU A 113      -9.780  45.276 -44.281  1.00  0.00           C  
ATOM   1810  C   LEU A 113      -8.931  44.355 -45.150  1.00  0.00           C  
ATOM   1811  O   LEU A 113      -7.759  44.123 -44.861  1.00  0.00           O  
ATOM   1812  CB  LEU A 113     -10.109  44.576 -42.956  1.00  0.00           C  
ATOM   1813  CG  LEU A 113     -10.802  45.445 -41.899  1.00  0.00           C  
ATOM   1814  CD1 LEU A 113     -11.173  44.585 -40.699  1.00  0.00           C  
ATOM   1815  CD2 LEU A 113      -9.878  46.583 -41.492  1.00  0.00           C  
ATOM   1816  H   LEU A 113     -11.872  45.191 -44.735  1.00  0.00           H  
ATOM   1817  HA  LEU A 113      -9.199  46.174 -44.073  1.00  0.00           H  
ATOM   1818 1HB  LEU A 113     -10.757  43.726 -43.164  1.00  0.00           H  
ATOM   1819 2HB  LEU A 113      -9.182  44.201 -42.522  1.00  0.00           H  
ATOM   1820  HG  LEU A 113     -11.723  45.855 -42.313  1.00  0.00           H  
ATOM   1821 1HD1 LEU A 113     -11.666  45.202 -39.947  1.00  0.00           H  
ATOM   1822 2HD1 LEU A 113     -11.850  43.791 -41.015  1.00  0.00           H  
ATOM   1823 3HD1 LEU A 113     -10.272  44.146 -40.273  1.00  0.00           H  
ATOM   1824 1HD2 LEU A 113     -10.371  47.201 -40.740  1.00  0.00           H  
ATOM   1825 2HD2 LEU A 113      -8.957  46.173 -41.077  1.00  0.00           H  
ATOM   1826 3HD2 LEU A 113      -9.643  47.191 -42.365  1.00  0.00           H  
ATOM   1827  N   GLY A 114      -9.534  43.842 -46.217  1.00  0.00           N  
ATOM   1828  CA  GLY A 114      -8.828  42.944 -47.115  1.00  0.00           C  
ATOM   1829  C   GLY A 114      -7.670  43.583 -47.857  1.00  0.00           C  
ATOM   1830  O   GLY A 114      -6.873  42.891 -48.486  1.00  0.00           O  
ATOM   1831  H   GLY A 114     -10.497  44.074 -46.413  1.00  0.00           H  
ATOM   1832 1HA  GLY A 114      -8.439  42.096 -46.551  1.00  0.00           H  
ATOM   1833 2HA  GLY A 114      -9.522  42.547 -47.854  1.00  0.00           H  
ATOM   1834  N   LYS A 115      -7.572  44.905 -47.788  1.00  0.00           N  
ATOM   1835  CA  LYS A 115      -6.493  45.616 -48.465  1.00  0.00           C  
ATOM   1836  C   LYS A 115      -5.235  45.647 -47.612  1.00  0.00           C  
ATOM   1837  O   LYS A 115      -4.156  45.982 -48.095  1.00  0.00           O  
ATOM   1838  CB  LYS A 115      -6.925  47.041 -48.814  1.00  0.00           C  
ATOM   1839  CG  LYS A 115      -8.043  47.125 -49.845  1.00  0.00           C  
ATOM   1840  CD  LYS A 115      -8.420  48.570 -50.136  1.00  0.00           C  
ATOM   1841  CE  LYS A 115      -9.531  48.655 -51.172  1.00  0.00           C  
ATOM   1842  NZ  LYS A 115      -9.932  50.062 -51.443  1.00  0.00           N1+
ATOM   1843  H   LYS A 115      -8.252  45.433 -47.259  1.00  0.00           H  
ATOM   1844  HA  LYS A 115      -6.265  45.094 -49.395  1.00  0.00           H  
ATOM   1845 1HB  LYS A 115      -7.264  47.550 -47.912  1.00  0.00           H  
ATOM   1846 2HB  LYS A 115      -6.070  47.596 -49.202  1.00  0.00           H  
ATOM   1847 1HG  LYS A 115      -7.720  46.649 -50.772  1.00  0.00           H  
ATOM   1848 2HG  LYS A 115      -8.921  46.597 -49.474  1.00  0.00           H  
ATOM   1849 1HD  LYS A 115      -8.755  49.051 -49.216  1.00  0.00           H  
ATOM   1850 2HD  LYS A 115      -7.547  49.105 -50.508  1.00  0.00           H  
ATOM   1851 1HE  LYS A 115      -9.196  48.200 -52.103  1.00  0.00           H  
ATOM   1852 2HE  LYS A 115     -10.402  48.104 -50.817  1.00  0.00           H  
ATOM   1853 1HZ  LYS A 115     -10.670  50.076 -52.133  1.00  0.00           H  
ATOM   1854 2HZ  LYS A 115     -10.263  50.489 -50.589  1.00  0.00           H  
ATOM   1855 3HZ  LYS A 115      -9.137  50.579 -51.790  1.00  0.00           H  
ATOM   1856  N   ILE A 116      -5.377  45.290 -46.341  1.00  0.00           N  
ATOM   1857  CA  ILE A 116      -4.238  45.289 -45.439  1.00  0.00           C  
ATOM   1858  C   ILE A 116      -3.332  44.082 -45.643  1.00  0.00           C  
ATOM   1859  O   ILE A 116      -3.799  42.967 -45.863  1.00  0.00           O  
ATOM   1860  CB  ILE A 116      -4.712  45.326 -43.975  1.00  0.00           C  
ATOM   1861  CG1 ILE A 116      -5.430  46.645 -43.679  1.00  0.00           C  
ATOM   1862  CG2 ILE A 116      -3.536  45.132 -43.030  1.00  0.00           C  
ATOM   1863  CD1 ILE A 116      -6.145  46.666 -42.347  1.00  0.00           C  
ATOM   1864  H   ILE A 116      -6.282  45.014 -45.989  1.00  0.00           H  
ATOM   1865  HA  ILE A 116      -3.642  46.180 -45.633  1.00  0.00           H  
ATOM   1866  HB  ILE A 116      -5.436  44.529 -43.807  1.00  0.00           H  
ATOM   1867 1HG1 ILE A 116      -4.709  47.462 -43.692  1.00  0.00           H  
ATOM   1868 2HG1 ILE A 116      -6.162  46.844 -44.461  1.00  0.00           H  
ATOM   1869 1HG2 ILE A 116      -3.888  45.161 -41.999  1.00  0.00           H  
ATOM   1870 2HG2 ILE A 116      -3.067  44.169 -43.226  1.00  0.00           H  
ATOM   1871 3HG2 ILE A 116      -2.808  45.929 -43.187  1.00  0.00           H  
ATOM   1872 1HD1 ILE A 116      -6.630  47.632 -42.209  1.00  0.00           H  
ATOM   1873 2HD1 ILE A 116      -6.898  45.876 -42.326  1.00  0.00           H  
ATOM   1874 3HD1 ILE A 116      -5.426  46.504 -41.545  1.00  0.00           H  
ATOM   1875  N   GLU A 117      -2.029  44.328 -45.574  1.00  0.00           N  
ATOM   1876  CA  GLU A 117      -1.019  43.285 -45.723  1.00  0.00           C  
ATOM   1877  C   GLU A 117      -1.351  42.032 -44.913  1.00  0.00           C  
ATOM   1878  O   GLU A 117      -1.657  42.117 -43.725  1.00  0.00           O  
ATOM   1879  CB  GLU A 117       0.337  43.820 -45.264  1.00  0.00           C  
ATOM   1880  CG  GLU A 117       0.281  44.500 -43.895  1.00  0.00           C  
ATOM   1881  CD  GLU A 117       1.645  44.917 -43.373  1.00  0.00           C  
ATOM   1882  OE1 GLU A 117       1.696  45.814 -42.505  1.00  0.00           O  
ATOM   1883  OE2 GLU A 117       2.662  44.341 -43.815  1.00  0.00           O1-
ATOM   1884  H   GLU A 117      -1.733  45.279 -45.411  1.00  0.00           H  
ATOM   1885  HA  GLU A 117      -0.958  43.012 -46.777  1.00  0.00           H  
ATOM   1886 1HB  GLU A 117       1.054  43.000 -45.215  1.00  0.00           H  
ATOM   1887 2HB  GLU A 117       0.711  44.539 -45.994  1.00  0.00           H  
ATOM   1888 1HG  GLU A 117      -0.350  45.386 -43.967  1.00  0.00           H  
ATOM   1889 2HG  GLU A 117      -0.178  43.818 -43.182  1.00  0.00           H  
ATOM   1890  N   LYS A 118      -1.290  40.879 -45.576  1.00  0.00           N  
ATOM   1891  CA  LYS A 118      -1.533  39.576 -44.961  1.00  0.00           C  
ATOM   1892  C   LYS A 118      -2.956  39.208 -44.565  1.00  0.00           C  
ATOM   1893  O   LYS A 118      -3.192  38.095 -44.091  1.00  0.00           O  
ATOM   1894  CB  LYS A 118      -0.623  39.398 -43.747  1.00  0.00           C  
ATOM   1895  CG  LYS A 118       0.838  39.287 -44.116  1.00  0.00           C  
ATOM   1896  CD  LYS A 118       1.679  38.949 -42.913  1.00  0.00           C  
ATOM   1897  CE  LYS A 118       3.132  38.764 -43.302  1.00  0.00           C  
ATOM   1898  NZ  LYS A 118       3.969  38.519 -42.102  1.00  0.00           N1+
ATOM   1899  H   LYS A 118      -1.061  40.924 -46.558  1.00  0.00           H  
ATOM   1900  HA  LYS A 118      -1.304  38.800 -45.692  1.00  0.00           H  
ATOM   1901 1HB  LYS A 118      -0.747  40.245 -43.071  1.00  0.00           H  
ATOM   1902 2HB  LYS A 118      -0.914  38.500 -43.202  1.00  0.00           H  
ATOM   1903 1HG  LYS A 118       0.965  38.508 -44.869  1.00  0.00           H  
ATOM   1904 2HG  LYS A 118       1.181  40.232 -44.535  1.00  0.00           H  
ATOM   1905 1HD  LYS A 118       1.603  39.752 -42.179  1.00  0.00           H  
ATOM   1906 2HD  LYS A 118       1.310  38.029 -42.459  1.00  0.00           H  
ATOM   1907 1HE  LYS A 118       3.220  37.920 -43.985  1.00  0.00           H  
ATOM   1908 2HE  LYS A 118       3.488  39.657 -43.816  1.00  0.00           H  
ATOM   1909 1HZ  LYS A 118       4.932  38.399 -42.381  1.00  0.00           H  
ATOM   1910 2HZ  LYS A 118       3.896  39.307 -41.473  1.00  0.00           H  
ATOM   1911 3HZ  LYS A 118       3.649  37.684 -41.632  1.00  0.00           H  
ATOM   1912  N   VAL A 119      -3.902  40.119 -44.759  1.00  0.00           N  
ATOM   1913  CA  VAL A 119      -5.289  39.842 -44.403  1.00  0.00           C  
ATOM   1914  C   VAL A 119      -6.065  39.198 -45.549  1.00  0.00           C  
ATOM   1915  O   VAL A 119      -6.065  39.700 -46.674  1.00  0.00           O  
ATOM   1916  CB  VAL A 119      -5.995  41.146 -43.986  1.00  0.00           C  
ATOM   1917  CG1 VAL A 119      -7.463  40.884 -43.686  1.00  0.00           C  
ATOM   1918  CG2 VAL A 119      -5.296  41.749 -42.777  1.00  0.00           C  
ATOM   1919  H   VAL A 119      -3.667  41.016 -45.158  1.00  0.00           H  
ATOM   1920  HA  VAL A 119      -5.297  39.149 -43.561  1.00  0.00           H  
ATOM   1921  HB  VAL A 119      -5.957  41.850 -44.818  1.00  0.00           H  
ATOM   1922 1HG1 VAL A 119      -7.947  41.817 -43.393  1.00  0.00           H  
ATOM   1923 2HG1 VAL A 119      -7.951  40.487 -44.576  1.00  0.00           H  
ATOM   1924 3HG1 VAL A 119      -7.545  40.163 -42.873  1.00  0.00           H  
ATOM   1925 1HG2 VAL A 119      -5.800  42.671 -42.489  1.00  0.00           H  
ATOM   1926 2HG2 VAL A 119      -5.329  41.043 -41.948  1.00  0.00           H  
ATOM   1927 3HG2 VAL A 119      -4.258  41.967 -43.028  1.00  0.00           H  
ATOM   1928  N   ARG A 120      -6.707  38.070 -45.258  1.00  0.00           N  
ATOM   1929  CA  ARG A 120      -7.528  37.369 -46.242  1.00  0.00           C  
ATOM   1930  C   ARG A 120      -8.942  37.316 -45.678  1.00  0.00           C  
ATOM   1931  O   ARG A 120      -9.143  36.882 -44.548  1.00  0.00           O  
ATOM   1932  CB  ARG A 120      -7.032  35.938 -46.471  1.00  0.00           C  
ATOM   1933  CG  ARG A 120      -7.920  35.166 -47.434  1.00  0.00           C  
ATOM   1934  CD  ARG A 120      -7.518  33.708 -47.579  1.00  0.00           C  
ATOM   1935  NE  ARG A 120      -8.329  33.055 -48.607  1.00  0.00           N  
ATOM   1936  CZ  ARG A 120      -8.260  31.762 -48.910  1.00  0.00           C  
ATOM   1937  NH1 ARG A 120      -9.036  31.264 -49.863  1.00  0.00           N1+
ATOM   1938  NH2 ARG A 120      -7.425  30.967 -48.258  1.00  0.00           N  
ATOM   1939  H   ARG A 120      -6.624  37.689 -44.327  1.00  0.00           H  
ATOM   1940  HA  ARG A 120      -7.467  37.905 -47.189  1.00  0.00           H  
ATOM   1941 1HB  ARG A 120      -6.018  35.965 -46.868  1.00  0.00           H  
ATOM   1942 2HB  ARG A 120      -6.997  35.409 -45.518  1.00  0.00           H  
ATOM   1943 1HG  ARG A 120      -8.950  35.189 -47.077  1.00  0.00           H  
ATOM   1944 2HG  ARG A 120      -7.868  35.623 -48.423  1.00  0.00           H  
ATOM   1945 1HD  ARG A 120      -6.468  33.646 -47.865  1.00  0.00           H  
ATOM   1946 2HD  ARG A 120      -7.667  33.193 -46.631  1.00  0.00           H  
ATOM   1947  HE  ARG A 120      -8.986  33.626 -49.120  1.00  0.00           H  
ATOM   1948 1HH1 ARG A 120      -9.679  31.866 -50.358  1.00  0.00           H  
ATOM   1949 2HH1 ARG A 120      -8.984  30.283 -50.095  1.00  0.00           H  
ATOM   1950 1HH2 ARG A 120      -6.837  31.343 -47.526  1.00  0.00           H  
ATOM   1951 2HH2 ARG A 120      -7.375  29.986 -48.492  1.00  0.00           H  
ATOM   1952  N   VAL A 121      -9.919  37.758 -46.460  1.00  0.00           N  
ATOM   1953  CA  VAL A 121     -11.299  37.762 -45.991  1.00  0.00           C  
ATOM   1954  C   VAL A 121     -12.171  36.681 -46.627  1.00  0.00           C  
ATOM   1955  O   VAL A 121     -12.245  36.564 -47.851  1.00  0.00           O  
ATOM   1956  CB  VAL A 121     -11.932  39.138 -46.271  1.00  0.00           C  
ATOM   1957  CG1 VAL A 121     -13.390  39.151 -45.836  1.00  0.00           C  
ATOM   1958  CG2 VAL A 121     -11.146  40.225 -45.555  1.00  0.00           C  
ATOM   1959  H   VAL A 121      -9.714  38.095 -47.389  1.00  0.00           H  
ATOM   1960  HA  VAL A 121     -11.300  37.570 -44.917  1.00  0.00           H  
ATOM   1961  HB  VAL A 121     -11.916  39.324 -47.345  1.00  0.00           H  
ATOM   1962 1HG1 VAL A 121     -13.822  40.131 -46.041  1.00  0.00           H  
ATOM   1963 2HG1 VAL A 121     -13.941  38.390 -46.387  1.00  0.00           H  
ATOM   1964 3HG1 VAL A 121     -13.453  38.945 -44.767  1.00  0.00           H  
ATOM   1965 1HG2 VAL A 121     -11.599  41.195 -45.758  1.00  0.00           H  
ATOM   1966 2HG2 VAL A 121     -11.158  40.036 -44.481  1.00  0.00           H  
ATOM   1967 3HG2 VAL A 121     -10.116  40.224 -45.912  1.00  0.00           H  
ATOM   1968  N   LEU A 122     -12.821  35.887 -45.782  1.00  0.00           N  
ATOM   1969  CA  LEU A 122     -13.723  34.834 -46.236  1.00  0.00           C  
ATOM   1970  C   LEU A 122     -15.104  35.148 -45.667  1.00  0.00           C  
ATOM   1971  O   LEU A 122     -15.313  35.062 -44.457  1.00  0.00           O  
ATOM   1972  CB  LEU A 122     -13.263  33.468 -45.722  1.00  0.00           C  
ATOM   1973  CG  LEU A 122     -11.870  33.000 -46.140  1.00  0.00           C  
ATOM   1974  CD1 LEU A 122     -11.587  31.659 -45.486  1.00  0.00           C  
ATOM   1975  CD2 LEU A 122     -11.784  32.896 -47.656  1.00  0.00           C  
ATOM   1976  H   LEU A 122     -12.681  36.022 -44.791  1.00  0.00           H  
ATOM   1977  HA  LEU A 122     -13.712  34.816 -47.326  1.00  0.00           H  
ATOM   1978 1HB  LEU A 122     -13.279  33.483 -44.633  1.00  0.00           H  
ATOM   1979 2HB  LEU A 122     -13.969  32.711 -46.065  1.00  0.00           H  
ATOM   1980  HG  LEU A 122     -11.125  33.715 -45.788  1.00  0.00           H  
ATOM   1981 1HD1 LEU A 122     -10.595  31.315 -45.777  1.00  0.00           H  
ATOM   1982 2HD1 LEU A 122     -11.629  31.767 -44.402  1.00  0.00           H  
ATOM   1983 3HD1 LEU A 122     -12.333  30.934 -45.810  1.00  0.00           H  
ATOM   1984 1HD2 LEU A 122     -10.786  32.563 -47.943  1.00  0.00           H  
ATOM   1985 2HD2 LEU A 122     -12.523  32.179 -48.014  1.00  0.00           H  
ATOM   1986 3HD2 LEU A 122     -11.981  33.873 -48.100  1.00  0.00           H  
ATOM   1987  N   ARG A 123     -16.044  35.520 -46.531  1.00  0.00           N  
ATOM   1988  CA  ARG A 123     -17.388  35.842 -46.062  1.00  0.00           C  
ATOM   1989  C   ARG A 123     -18.329  34.666 -46.244  1.00  0.00           C  
ATOM   1990  O   ARG A 123     -18.444  34.117 -47.343  1.00  0.00           O  
ATOM   1991  CB  ARG A 123     -17.955  37.056 -46.808  1.00  0.00           C  
ATOM   1992  CG  ARG A 123     -19.366  37.445 -46.356  1.00  0.00           C  
ATOM   1993  CD  ARG A 123     -19.863  38.706 -47.047  1.00  0.00           C  
ATOM   1994  NE  ARG A 123     -21.305  38.892 -46.889  1.00  0.00           N  
ATOM   1995  CZ  ARG A 123     -21.898  39.420 -45.823  1.00  0.00           C  
ATOM   1996  NH1 ARG A 123     -23.222  39.533 -45.794  1.00  0.00           N1+
ATOM   1997  NH2 ARG A 123     -21.180  39.852 -44.793  1.00  0.00           N  
ATOM   1998  H   ARG A 123     -15.839  35.583 -47.518  1.00  0.00           H  
ATOM   1999  HA  ARG A 123     -17.335  36.085 -45.001  1.00  0.00           H  
ATOM   2000 1HB  ARG A 123     -17.299  37.913 -46.660  1.00  0.00           H  
ATOM   2001 2HB  ARG A 123     -17.983  36.845 -47.877  1.00  0.00           H  
ATOM   2002 1HG  ARG A 123     -20.059  36.636 -46.590  1.00  0.00           H  
ATOM   2003 2HG  ARG A 123     -19.368  37.626 -45.281  1.00  0.00           H  
ATOM   2004 1HD  ARG A 123     -19.361  39.575 -46.623  1.00  0.00           H  
ATOM   2005 2HD  ARG A 123     -19.644  38.647 -48.113  1.00  0.00           H  
ATOM   2006  HE  ARG A 123     -21.903  38.596 -47.650  1.00  0.00           H  
ATOM   2007 1HH1 ARG A 123     -23.773  39.217 -46.579  1.00  0.00           H  
ATOM   2008 2HH1 ARG A 123     -23.677  39.934 -44.987  1.00  0.00           H  
ATOM   2009 1HH2 ARG A 123     -20.171  39.781 -44.815  1.00  0.00           H  
ATOM   2010 2HH2 ARG A 123     -21.640  40.252 -43.989  1.00  0.00           H  
ATOM   2011  N   ASN A 124     -18.990  34.270 -45.161  1.00  0.00           N  
ATOM   2012  CA  ASN A 124     -19.938  33.166 -45.227  1.00  0.00           C  
ATOM   2013  C   ASN A 124     -21.158  33.615 -46.015  1.00  0.00           C  
ATOM   2014  O   ASN A 124     -21.633  34.736 -45.841  1.00  0.00           O  
ATOM   2015  CB  ASN A 124     -20.326  32.688 -43.840  1.00  0.00           C  
ATOM   2016  CG  ASN A 124     -19.183  32.035 -43.113  1.00  0.00           C  
ATOM   2017  ND2 ASN A 124     -19.258  32.014 -41.806  1.00  0.00           N  
ATOM   2018  OD1 ASN A 124     -18.231  31.550 -43.735  1.00  0.00           O  
ATOM   2019  H   ASN A 124     -18.837  34.736 -44.277  1.00  0.00           H  
ATOM   2020  HA  ASN A 124     -19.450  32.318 -45.711  1.00  0.00           H  
ATOM   2021 1HB  ASN A 124     -20.682  33.534 -43.250  1.00  0.00           H  
ATOM   2022 2HB  ASN A 124     -21.147  31.975 -43.917  1.00  0.00           H  
ATOM   2023 1HD2 ASN A 124     -18.526  31.593 -41.270  1.00  0.00           H  
ATOM   2024 2HD2 ASN A 124     -20.047  32.419 -41.345  1.00  0.00           H  
ATOM   2025  N   ASP A 125     -21.656  32.737 -46.879  1.00  0.00           N  
ATOM   2026  CA  ASP A 125     -22.824  33.037 -47.701  1.00  0.00           C  
ATOM   2027  C   ASP A 125     -24.103  33.079 -46.870  1.00  0.00           C  
ATOM   2028  O   ASP A 125     -25.127  33.603 -47.313  1.00  0.00           O  
ATOM   2029  CB  ASP A 125     -22.971  32.000 -48.817  1.00  0.00           C  
ATOM   2030  CG  ASP A 125     -21.895  32.125 -49.888  1.00  0.00           C  
ATOM   2031  OD1 ASP A 125     -21.239  33.139 -49.927  1.00  0.00           O  
ATOM   2032  OD2 ASP A 125     -21.740  31.206 -50.656  1.00  0.00           O1-
ATOM   2033  H   ASP A 125     -21.211  31.834 -46.967  1.00  0.00           H  
ATOM   2034  HA  ASP A 125     -22.698  34.031 -48.131  1.00  0.00           H  
ATOM   2035 1HB  ASP A 125     -22.923  30.997 -48.391  1.00  0.00           H  
ATOM   2036 2HB  ASP A 125     -23.947  32.108 -49.289  1.00  0.00           H  
ATOM   2037  N   ARG A 126     -24.036  32.519 -45.665  1.00  0.00           N  
ATOM   2038  CA  ARG A 126     -25.154  32.523 -44.727  1.00  0.00           C  
ATOM   2039  C   ARG A 126     -24.564  32.464 -43.322  1.00  0.00           C  
ATOM   2040  O   ARG A 126     -23.374  32.195 -43.163  1.00  0.00           O  
ATOM   2041  CB  ARG A 126     -26.087  31.344 -44.959  1.00  0.00           C  
ATOM   2042  CG  ARG A 126     -25.477  29.980 -44.677  1.00  0.00           C  
ATOM   2043  CD  ARG A 126     -26.431  28.880 -44.971  1.00  0.00           C  
ATOM   2044  NE  ARG A 126     -25.862  27.575 -44.675  1.00  0.00           N  
ATOM   2045  CZ  ARG A 126     -26.499  26.403 -44.860  1.00  0.00           C  
ATOM   2046  NH1 ARG A 126     -27.723  26.389 -45.341  1.00  0.00           N1+
ATOM   2047  NH2 ARG A 126     -25.895  25.267 -44.559  1.00  0.00           N  
ATOM   2048  H   ARG A 126     -23.170  32.075 -45.396  1.00  0.00           H  
ATOM   2049  HA  ARG A 126     -25.730  33.436 -44.880  1.00  0.00           H  
ATOM   2050 1HB  ARG A 126     -26.968  31.448 -44.327  1.00  0.00           H  
ATOM   2051 2HB  ARG A 126     -26.425  31.346 -45.995  1.00  0.00           H  
ATOM   2052 1HG  ARG A 126     -24.592  29.842 -45.298  1.00  0.00           H  
ATOM   2053 2HG  ARG A 126     -25.196  29.918 -43.625  1.00  0.00           H  
ATOM   2054 1HD  ARG A 126     -27.329  29.006 -44.366  1.00  0.00           H  
ATOM   2055 2HD  ARG A 126     -26.699  28.904 -46.026  1.00  0.00           H  
ATOM   2056  HE  ARG A 126     -24.922  27.546 -44.303  1.00  0.00           H  
ATOM   2057 1HH1 ARG A 126     -28.185  27.257 -45.571  1.00  0.00           H  
ATOM   2058 2HH1 ARG A 126     -28.201  25.510 -45.480  1.00  0.00           H  
ATOM   2059 1HH2 ARG A 126     -24.954  25.278 -44.190  1.00  0.00           H  
ATOM   2060 2HH2 ARG A 126     -26.372  24.389 -44.699  1.00  0.00           H  
ATOM   2061  N   ARG A 127     -25.385  32.728 -42.310  1.00  0.00           N  
ATOM   2062  CA  ARG A 127     -24.922  32.690 -40.922  1.00  0.00           C  
ATOM   2063  C   ARG A 127     -24.533  31.252 -40.576  1.00  0.00           C  
ATOM   2064  O   ARG A 127     -25.324  30.324 -40.769  1.00  0.00           O  
ATOM   2065  CB  ARG A 127     -26.036  33.180 -39.988  1.00  0.00           C  
ATOM   2066  CG  ARG A 127     -25.678  33.233 -38.504  1.00  0.00           C  
ATOM   2067  CD  ARG A 127     -24.720  34.365 -38.164  1.00  0.00           C  
ATOM   2068  NE  ARG A 127     -24.533  34.494 -36.718  1.00  0.00           N  
ATOM   2069  CZ  ARG A 127     -25.450  34.974 -35.882  1.00  0.00           C  
ATOM   2070  NH1 ARG A 127     -26.624  35.386 -36.344  1.00  0.00           N1+
ATOM   2071  NH2 ARG A 127     -25.202  35.030 -34.577  1.00  0.00           N  
ATOM   2072  H   ARG A 127     -26.349  32.961 -42.501  1.00  0.00           H  
ATOM   2073  HA  ARG A 127     -24.062  33.353 -40.826  1.00  0.00           H  
ATOM   2074 1HB  ARG A 127     -26.341  34.183 -40.281  1.00  0.00           H  
ATOM   2075 2HB  ARG A 127     -26.905  32.530 -40.087  1.00  0.00           H  
ATOM   2076 1HG  ARG A 127     -26.585  33.379 -37.916  1.00  0.00           H  
ATOM   2077 2HG  ARG A 127     -25.202  32.296 -38.210  1.00  0.00           H  
ATOM   2078 1HD  ARG A 127     -23.749  34.170 -38.621  1.00  0.00           H  
ATOM   2079 2HD  ARG A 127     -25.117  35.304 -38.545  1.00  0.00           H  
ATOM   2080  HE  ARG A 127     -23.647  34.198 -36.331  1.00  0.00           H  
ATOM   2081 1HH1 ARG A 127     -26.824  35.334 -37.333  1.00  0.00           H  
ATOM   2082 2HH1 ARG A 127     -27.318  35.749 -35.706  1.00  0.00           H  
ATOM   2083 1HH2 ARG A 127     -24.314  34.708 -34.217  1.00  0.00           H  
ATOM   2084 2HH2 ARG A 127     -25.900  35.396 -33.946  1.00  0.00           H  
ATOM   2085  N   GLU A 128     -23.318  31.066 -40.060  1.00  0.00           N  
ATOM   2086  CA  GLU A 128     -22.832  29.730 -39.715  1.00  0.00           C  
ATOM   2087  C   GLU A 128     -22.519  29.524 -38.233  1.00  0.00           C  
ATOM   2088  O   GLU A 128     -22.513  28.391 -37.753  1.00  0.00           O  
ATOM   2089  CB  GLU A 128     -21.583  29.405 -40.537  1.00  0.00           C  
ATOM   2090  CG  GLU A 128     -21.828  29.315 -42.033  1.00  0.00           C  
ATOM   2091  CD  GLU A 128     -22.617  28.079 -42.434  1.00  0.00           C  
ATOM   2092  OE1 GLU A 128     -22.919  27.936 -43.641  1.00  0.00           O  
ATOM   2093  OE2 GLU A 128     -22.933  27.245 -41.555  1.00  0.00           O1-
ATOM   2094  H   GLU A 128     -22.718  31.863 -39.903  1.00  0.00           H  
ATOM   2095  HA  GLU A 128     -23.611  29.006 -39.953  1.00  0.00           H  
ATOM   2096 1HB  GLU A 128     -20.825  30.170 -40.366  1.00  0.00           H  
ATOM   2097 2HB  GLU A 128     -21.168  28.453 -40.206  1.00  0.00           H  
ATOM   2098 1HG  GLU A 128     -22.376  30.201 -42.355  1.00  0.00           H  
ATOM   2099 2HG  GLU A 128     -20.868  29.309 -42.547  1.00  0.00           H  
ATOM   2100  N   GLY A 129     -22.261  30.609 -37.510  1.00  0.00           N  
ATOM   2101  CA  GLY A 129     -21.934  30.474 -36.099  1.00  0.00           C  
ATOM   2102  C   GLY A 129     -20.425  30.383 -35.949  1.00  0.00           C  
ATOM   2103  O   GLY A 129     -19.716  30.144 -36.929  1.00  0.00           O  
ATOM   2104  H   GLY A 129     -22.287  31.531 -37.923  1.00  0.00           H  
ATOM   2105 1HA  GLY A 129     -22.328  31.330 -35.551  1.00  0.00           H  
ATOM   2106 2HA  GLY A 129     -22.418  29.585 -35.697  1.00  0.00           H  
ATOM   2107  N   LEU A 130     -19.917  30.539 -34.732  1.00  0.00           N  
ATOM   2108  CA  LEU A 130     -18.469  30.515 -34.545  1.00  0.00           C  
ATOM   2109  C   LEU A 130     -17.790  29.167 -34.763  1.00  0.00           C  
ATOM   2110  O   LEU A 130     -16.641  29.127 -35.206  1.00  0.00           O  
ATOM   2111  CB  LEU A 130     -18.144  31.001 -33.127  1.00  0.00           C  
ATOM   2112  CG  LEU A 130     -18.437  32.480 -32.847  1.00  0.00           C  
ATOM   2113  CD1 LEU A 130     -18.206  32.774 -31.371  1.00  0.00           C  
ATOM   2114  CD2 LEU A 130     -17.546  33.348 -33.723  1.00  0.00           C  
ATOM   2115  H   LEU A 130     -20.514  30.674 -33.930  1.00  0.00           H  
ATOM   2116  HA  LEU A 130     -18.011  31.139 -35.311  1.00  0.00           H  
ATOM   2117 1HB  LEU A 130     -18.720  30.409 -32.417  1.00  0.00           H  
ATOM   2118 2HB  LEU A 130     -17.085  30.830 -32.935  1.00  0.00           H  
ATOM   2119  HG  LEU A 130     -19.483  32.693 -33.069  1.00  0.00           H  
ATOM   2120 1HD1 LEU A 130     -18.414  33.825 -31.172  1.00  0.00           H  
ATOM   2121 2HD1 LEU A 130     -18.869  32.153 -30.768  1.00  0.00           H  
ATOM   2122 3HD1 LEU A 130     -17.170  32.554 -31.114  1.00  0.00           H  
ATOM   2123 1HD2 LEU A 130     -17.754  34.400 -33.524  1.00  0.00           H  
ATOM   2124 2HD2 LEU A 130     -16.500  33.138 -33.500  1.00  0.00           H  
ATOM   2125 3HD2 LEU A 130     -17.744  33.131 -34.772  1.00  0.00           H  
ATOM   2126  N   MET A 131     -18.483  28.067 -34.474  1.00  0.00           N  
ATOM   2127  CA  MET A 131     -17.869  26.755 -34.649  1.00  0.00           C  
ATOM   2128  C   MET A 131     -17.587  26.442 -36.111  1.00  0.00           C  
ATOM   2129  O   MET A 131     -16.458  26.106 -36.467  1.00  0.00           O  
ATOM   2130  CB  MET A 131     -18.743  25.645 -34.054  1.00  0.00           C  
ATOM   2131  CG  MET A 131     -19.109  25.842 -32.590  1.00  0.00           C  
ATOM   2132  SD  MET A 131     -20.599  26.844 -32.399  1.00  0.00           S  
ATOM   2133  CE  MET A 131     -20.673  26.989 -30.603  1.00  0.00           C  
ATOM   2134  H   MET A 131     -19.432  28.125 -34.134  1.00  0.00           H  
ATOM   2135  HA  MET A 131     -16.910  26.749 -34.129  1.00  0.00           H  
ATOM   2136 1HB  MET A 131     -19.670  25.570 -34.620  1.00  0.00           H  
ATOM   2137 2HB  MET A 131     -18.227  24.688 -34.141  1.00  0.00           H  
ATOM   2138 1HG  MET A 131     -19.274  24.872 -32.122  1.00  0.00           H  
ATOM   2139 2HG  MET A 131     -18.285  26.334 -32.072  1.00  0.00           H  
ATOM   2140 1HE  MET A 131     -21.542  27.583 -30.321  1.00  0.00           H  
ATOM   2141 2HE  MET A 131     -20.753  25.996 -30.160  1.00  0.00           H  
ATOM   2142 3HE  MET A 131     -19.767  27.477 -30.239  1.00  0.00           H  
ATOM   2143  N   ARG A 132     -18.606  26.537 -36.957  1.00  0.00           N  
ATOM   2144  CA  ARG A 132     -18.410  26.257 -38.372  1.00  0.00           C  
ATOM   2145  C   ARG A 132     -17.534  27.325 -39.037  1.00  0.00           C  
ATOM   2146  O   ARG A 132     -16.828  27.044 -40.008  1.00  0.00           O  
ATOM   2147  CB  ARG A 132     -19.765  26.136 -39.078  1.00  0.00           C  
ATOM   2148  CG  ARG A 132     -20.564  24.919 -38.611  1.00  0.00           C  
ATOM   2149  CD  ARG A 132     -21.764  24.649 -39.503  1.00  0.00           C  
ATOM   2150  NE  ARG A 132     -22.879  25.563 -39.259  1.00  0.00           N  
ATOM   2151  CZ  ARG A 132     -23.719  25.470 -38.234  1.00  0.00           C  
ATOM   2152  NH1 ARG A 132     -23.581  24.499 -37.337  1.00  0.00           N1+
ATOM   2153  NH2 ARG A 132     -24.715  26.341 -38.114  1.00  0.00           N  
ATOM   2154  H   ARG A 132     -19.524  26.804 -36.631  1.00  0.00           H  
ATOM   2155  HA  ARG A 132     -17.878  25.310 -38.467  1.00  0.00           H  
ATOM   2156 1HB  ARG A 132     -20.352  27.035 -38.894  1.00  0.00           H  
ATOM   2157 2HB  ARG A 132     -19.609  26.062 -40.154  1.00  0.00           H  
ATOM   2158 1HG  ARG A 132     -19.925  24.037 -38.625  1.00  0.00           H  
ATOM   2159 2HG  ARG A 132     -20.926  25.089 -37.596  1.00  0.00           H  
ATOM   2160 1HD  ARG A 132     -21.473  24.757 -40.547  1.00  0.00           H  
ATOM   2161 2HD  ARG A 132     -22.125  23.636 -39.329  1.00  0.00           H  
ATOM   2162  HE  ARG A 132     -23.021  26.318 -39.917  1.00  0.00           H  
ATOM   2163 1HH1 ARG A 132     -22.834  23.825 -37.432  1.00  0.00           H  
ATOM   2164 2HH1 ARG A 132     -24.224  24.433 -36.560  1.00  0.00           H  
ATOM   2165 1HH2 ARG A 132     -24.833  27.073 -38.801  1.00  0.00           H  
ATOM   2166 2HH2 ARG A 132     -25.354  26.272 -37.336  1.00  0.00           H  
ATOM   2167  N   SER A 133     -17.565  28.545 -38.506  1.00  0.00           N  
ATOM   2168  CA  SER A 133     -16.747  29.614 -39.064  1.00  0.00           C  
ATOM   2169  C   SER A 133     -15.275  29.372 -38.755  1.00  0.00           C  
ATOM   2170  O   SER A 133     -14.417  29.573 -39.618  1.00  0.00           O  
ATOM   2171  CB  SER A 133     -17.181  30.956 -38.507  1.00  0.00           C  
ATOM   2172  OG  SER A 133     -18.477  31.280 -38.931  1.00  0.00           O  
ATOM   2173  H   SER A 133     -18.156  28.741 -37.711  1.00  0.00           H  
ATOM   2174  HA  SER A 133     -16.879  29.622 -40.147  1.00  0.00           H  
ATOM   2175 1HB  SER A 133     -17.149  30.926 -37.419  1.00  0.00           H  
ATOM   2176 2HB  SER A 133     -16.486  31.728 -38.834  1.00  0.00           H  
ATOM   2177  HG  SER A 133     -19.073  30.788 -38.360  1.00  0.00           H  
ATOM   2178  N   ARG A 134     -14.970  28.937 -37.533  1.00  0.00           N  
ATOM   2179  CA  ARG A 134     -13.578  28.665 -37.185  1.00  0.00           C  
ATOM   2180  C   ARG A 134     -13.065  27.496 -38.013  1.00  0.00           C  
ATOM   2181  O   ARG A 134     -11.913  27.491 -38.442  1.00  0.00           O  
ATOM   2182  CB  ARG A 134     -13.423  28.346 -35.695  1.00  0.00           C  
ATOM   2183  CG  ARG A 134     -13.506  29.575 -34.803  1.00  0.00           C  
ATOM   2184  CD  ARG A 134     -13.343  29.211 -33.331  1.00  0.00           C  
ATOM   2185  NE  ARG A 134     -13.741  30.331 -32.482  1.00  0.00           N  
ATOM   2186  CZ  ARG A 134     -14.019  30.238 -31.186  1.00  0.00           C  
ATOM   2187  NH1 ARG A 134     -14.381  31.322 -30.511  1.00  0.00           N1+
ATOM   2188  NH2 ARG A 134     -13.938  29.070 -30.563  1.00  0.00           N  
ATOM   2189  H   ARG A 134     -15.689  28.790 -36.838  1.00  0.00           H  
ATOM   2190  HA  ARG A 134     -12.987  29.555 -37.405  1.00  0.00           H  
ATOM   2191 1HB  ARG A 134     -14.200  27.647 -35.390  1.00  0.00           H  
ATOM   2192 2HB  ARG A 134     -12.461  27.861 -35.525  1.00  0.00           H  
ATOM   2193 1HG  ARG A 134     -12.715  30.275 -35.073  1.00  0.00           H  
ATOM   2194 2HG  ARG A 134     -14.476  30.055 -34.934  1.00  0.00           H  
ATOM   2195 1HD  ARG A 134     -13.967  28.349 -33.097  1.00  0.00           H  
ATOM   2196 2HD  ARG A 134     -12.300  28.967 -33.129  1.00  0.00           H  
ATOM   2197  HE  ARG A 134     -13.812  31.244 -32.911  1.00  0.00           H  
ATOM   2198 1HH1 ARG A 134     -14.444  32.213 -30.983  1.00  0.00           H  
ATOM   2199 2HH1 ARG A 134     -14.593  31.256 -29.526  1.00  0.00           H  
ATOM   2200 1HH2 ARG A 134     -13.663  28.242 -31.075  1.00  0.00           H  
ATOM   2201 2HH2 ARG A 134     -14.150  29.007 -29.579  1.00  0.00           H  
ATOM   2202  N   VAL A 135     -13.918  26.503 -38.242  1.00  0.00           N  
ATOM   2203  CA  VAL A 135     -13.497  25.360 -39.038  1.00  0.00           C  
ATOM   2204  C   VAL A 135     -13.220  25.796 -40.476  1.00  0.00           C  
ATOM   2205  O   VAL A 135     -12.261  25.327 -41.088  1.00  0.00           O  
ATOM   2206  CB  VAL A 135     -14.550  24.229 -38.999  1.00  0.00           C  
ATOM   2207  CG1 VAL A 135     -14.164  23.117 -39.973  1.00  0.00           C  
ATOM   2208  CG2 VAL A 135     -14.640  23.668 -37.583  1.00  0.00           C  
ATOM   2209  H   VAL A 135     -14.857  26.527 -37.873  1.00  0.00           H  
ATOM   2210  HA  VAL A 135     -12.566  24.973 -38.622  1.00  0.00           H  
ATOM   2211  HB  VAL A 135     -15.516  24.635 -39.297  1.00  0.00           H  
ATOM   2212 1HG1 VAL A 135     -14.913  22.326 -39.938  1.00  0.00           H  
ATOM   2213 2HG1 VAL A 135     -14.112  23.521 -40.984  1.00  0.00           H  
ATOM   2214 3HG1 VAL A 135     -13.193  22.709 -39.694  1.00  0.00           H  
ATOM   2215 1HG2 VAL A 135     -15.383  22.871 -37.554  1.00  0.00           H  
ATOM   2216 2HG2 VAL A 135     -13.670  23.272 -37.285  1.00  0.00           H  
ATOM   2217 3HG2 VAL A 135     -14.934  24.462 -36.895  1.00  0.00           H  
ATOM   2218  N   ARG A 136     -14.035  26.705 -41.015  1.00  0.00           N  
ATOM   2219  CA  ARG A 136     -13.791  27.171 -42.380  1.00  0.00           C  
ATOM   2220  C   ARG A 136     -12.434  27.862 -42.443  1.00  0.00           C  
ATOM   2221  O   ARG A 136     -11.660  27.642 -43.372  1.00  0.00           O  
ATOM   2222  CB  ARG A 136     -14.863  28.161 -42.854  1.00  0.00           C  
ATOM   2223  CG  ARG A 136     -14.594  28.629 -44.287  1.00  0.00           C  
ATOM   2224  CD  ARG A 136     -15.549  29.697 -44.771  1.00  0.00           C  
ATOM   2225  NE  ARG A 136     -15.260  30.031 -46.162  1.00  0.00           N  
ATOM   2226  CZ  ARG A 136     -15.885  30.973 -46.858  1.00  0.00           C  
ATOM   2227  NH1 ARG A 136     -16.849  31.689 -46.296  1.00  0.00           N1+
ATOM   2228  NH2 ARG A 136     -15.530  31.213 -48.117  1.00  0.00           N  
ATOM   2229  H   ARG A 136     -14.819  27.079 -40.501  1.00  0.00           H  
ATOM   2230  HA  ARG A 136     -13.813  26.311 -43.048  1.00  0.00           H  
ATOM   2231 1HB  ARG A 136     -15.843  27.687 -42.804  1.00  0.00           H  
ATOM   2232 2HB  ARG A 136     -14.882  29.024 -42.187  1.00  0.00           H  
ATOM   2233 1HG  ARG A 136     -13.588  29.043 -44.352  1.00  0.00           H  
ATOM   2234 2HG  ARG A 136     -14.684  27.783 -44.969  1.00  0.00           H  
ATOM   2235 1HD  ARG A 136     -16.572  29.331 -44.695  1.00  0.00           H  
ATOM   2236 2HD  ARG A 136     -15.436  30.590 -44.158  1.00  0.00           H  
ATOM   2237  HE  ARG A 136     -14.533  29.508 -46.631  1.00  0.00           H  
ATOM   2238 1HH1 ARG A 136     -17.110  31.519 -45.334  1.00  0.00           H  
ATOM   2239 2HH1 ARG A 136     -17.322  32.405 -46.828  1.00  0.00           H  
ATOM   2240 1HH2 ARG A 136     -14.786  30.678 -48.543  1.00  0.00           H  
ATOM   2241 2HH2 ARG A 136     -16.004  31.929 -48.647  1.00  0.00           H  
ATOM   2242  N   GLY A 137     -12.153  28.703 -41.454  1.00  0.00           N  
ATOM   2243  CA  GLY A 137     -10.880  29.401 -41.420  1.00  0.00           C  
ATOM   2244  C   GLY A 137      -9.719  28.435 -41.267  1.00  0.00           C  
ATOM   2245  O   GLY A 137      -8.685  28.586 -41.915  1.00  0.00           O  
ATOM   2246  H   GLY A 137     -12.823  28.867 -40.715  1.00  0.00           H  
ATOM   2247 1HA  GLY A 137     -10.756  29.977 -42.338  1.00  0.00           H  
ATOM   2248 2HA  GLY A 137     -10.875  30.110 -40.593  1.00  0.00           H  
ATOM   2249  N   ALA A 138      -9.890  27.441 -40.402  1.00  0.00           N  
ATOM   2250  CA  ALA A 138      -8.852  26.442 -40.172  1.00  0.00           C  
ATOM   2251  C   ALA A 138      -8.581  25.625 -41.434  1.00  0.00           C  
ATOM   2252  O   ALA A 138      -7.428  25.332 -41.761  1.00  0.00           O  
ATOM   2253  CB  ALA A 138      -9.263  25.522 -39.030  1.00  0.00           C  
ATOM   2254  H   ALA A 138     -10.759  27.374 -39.892  1.00  0.00           H  
ATOM   2255  HA  ALA A 138      -7.935  26.964 -39.898  1.00  0.00           H  
ATOM   2256 1HB  ALA A 138      -8.483  24.779 -38.862  1.00  0.00           H  
ATOM   2257 2HB  ALA A 138      -9.405  26.109 -38.122  1.00  0.00           H  
ATOM   2258 3HB  ALA A 138     -10.195  25.020 -39.285  1.00  0.00           H  
ATOM   2259  N   ASP A 139      -9.647  25.257 -42.137  1.00  0.00           N  
ATOM   2260  CA  ASP A 139      -9.537  24.475 -43.368  1.00  0.00           C  
ATOM   2261  C   ASP A 139      -8.812  25.243 -44.470  1.00  0.00           C  
ATOM   2262  O   ASP A 139      -8.204  24.644 -45.359  1.00  0.00           O  
ATOM   2263  CB  ASP A 139     -10.927  24.084 -43.878  1.00  0.00           C  
ATOM   2264  CG  ASP A 139     -11.549  22.953 -43.084  1.00  0.00           C  
ATOM   2265  OD1 ASP A 139     -12.735  22.647 -43.330  1.00  0.00           O  
ATOM   2266  OD2 ASP A 139     -10.857  22.361 -42.231  1.00  0.00           O1-
ATOM   2267  H   ASP A 139     -10.562  25.528 -41.806  1.00  0.00           H  
ATOM   2268  HA  ASP A 139      -8.974  23.566 -43.151  1.00  0.00           H  
ATOM   2269 1HB  ASP A 139     -11.589  24.948 -43.830  1.00  0.00           H  
ATOM   2270 2HB  ASP A 139     -10.860  23.780 -44.923  1.00  0.00           H  
ATOM   2271  N   ALA A 140      -8.885  26.568 -44.412  1.00  0.00           N  
ATOM   2272  CA  ALA A 140      -8.248  27.417 -45.412  1.00  0.00           C  
ATOM   2273  C   ALA A 140      -6.827  27.803 -45.024  1.00  0.00           C  
ATOM   2274  O   ALA A 140      -6.072  28.315 -45.850  1.00  0.00           O  
ATOM   2275  CB  ALA A 140      -9.078  28.674 -45.624  1.00  0.00           C  
ATOM   2276  H   ALA A 140      -9.395  26.998 -43.654  1.00  0.00           H  
ATOM   2277  HA  ALA A 140      -8.198  26.860 -46.347  1.00  0.00           H  
ATOM   2278 1HB  ALA A 140      -8.598  29.306 -46.372  1.00  0.00           H  
ATOM   2279 2HB  ALA A 140     -10.076  28.399 -45.967  1.00  0.00           H  
ATOM   2280 3HB  ALA A 140      -9.156  29.220 -44.685  1.00  0.00           H  
ATOM   2281  N   ALA A 141      -6.463  27.548 -43.770  1.00  0.00           N  
ATOM   2282  CA  ALA A 141      -5.139  27.903 -43.265  1.00  0.00           C  
ATOM   2283  C   ALA A 141      -3.999  27.189 -43.980  1.00  0.00           C  
ATOM   2284  O   ALA A 141      -4.052  25.979 -44.210  1.00  0.00           O  
ATOM   2285  CB  ALA A 141      -5.069  27.624 -41.770  1.00  0.00           C  
ATOM   2286  H   ALA A 141      -7.120  27.097 -43.150  1.00  0.00           H  
ATOM   2287  HA  ALA A 141      -4.984  28.969 -43.439  1.00  0.00           H  
ATOM   2288 1HB  ALA A 141      -4.080  27.889 -41.396  1.00  0.00           H  
ATOM   2289 2HB  ALA A 141      -5.824  28.217 -41.253  1.00  0.00           H  
ATOM   2290 3HB  ALA A 141      -5.252  26.566 -41.589  1.00  0.00           H  
ATOM   2291  N   GLN A 142      -2.958  27.942 -44.313  1.00  0.00           N  
ATOM   2292  CA  GLN A 142      -1.814  27.369 -45.002  1.00  0.00           C  
ATOM   2293  C   GLN A 142      -0.632  27.079 -44.085  1.00  0.00           C  
ATOM   2294  O   GLN A 142       0.288  26.353 -44.466  1.00  0.00           O  
ATOM   2295  CB  GLN A 142      -1.368  28.305 -46.128  1.00  0.00           C  
ATOM   2296  CG  GLN A 142      -2.427  28.553 -47.189  1.00  0.00           C  
ATOM   2297  CD  GLN A 142      -2.836  27.280 -47.906  1.00  0.00           C  
ATOM   2298  NE2 GLN A 142      -4.129  26.978 -47.883  1.00  0.00           N  
ATOM   2299  OE1 GLN A 142      -1.998  26.576 -48.476  1.00  0.00           O  
ATOM   2300  H   GLN A 142      -2.955  28.927 -44.088  1.00  0.00           H  
ATOM   2301  HA  GLN A 142      -2.104  26.401 -45.410  1.00  0.00           H  
ATOM   2302 1HB  GLN A 142      -1.080  29.269 -45.708  1.00  0.00           H  
ATOM   2303 2HB  GLN A 142      -0.489  27.888 -46.621  1.00  0.00           H  
ATOM   2304 1HG  GLN A 142      -3.311  28.976 -46.712  1.00  0.00           H  
ATOM   2305 2HG  GLN A 142      -2.031  29.250 -47.927  1.00  0.00           H  
ATOM   2306 1HE2 GLN A 142      -4.458  26.151 -48.339  1.00  0.00           H  
ATOM   2307 2HE2 GLN A 142      -4.774  27.577 -47.409  1.00  0.00           H  
ATOM   2308  N   ALA A 143      -0.658  27.634 -42.876  1.00  0.00           N  
ATOM   2309  CA  ALA A 143       0.425  27.402 -41.926  1.00  0.00           C  
ATOM   2310  C   ALA A 143       0.233  26.054 -41.238  1.00  0.00           C  
ATOM   2311  O   ALA A 143      -0.807  25.407 -41.391  1.00  0.00           O  
ATOM   2312  CB  ALA A 143       0.470  28.520 -40.889  1.00  0.00           C  
ATOM   2313  H   ALA A 143      -1.432  28.222 -42.605  1.00  0.00           H  
ATOM   2314  HA  ALA A 143       1.364  27.395 -42.479  1.00  0.00           H  
ATOM   2315 1HB  ALA A 143       1.283  28.333 -40.188  1.00  0.00           H  
ATOM   2316 2HB  ALA A 143       0.634  29.474 -41.389  1.00  0.00           H  
ATOM   2317 3HB  ALA A 143      -0.474  28.552 -40.349  1.00  0.00           H  
ATOM   2318  N   LYS A 144       1.240  25.635 -40.477  1.00  0.00           N  
ATOM   2319  CA  LYS A 144       1.174  24.365 -39.766  1.00  0.00           C  
ATOM   2320  C   LYS A 144       0.613  24.502 -38.355  1.00  0.00           C  
ATOM   2321  O   LYS A 144       0.358  23.505 -37.685  1.00  0.00           O  
ATOM   2322  CB  LYS A 144       2.562  23.726 -39.709  1.00  0.00           C  
ATOM   2323  CG  LYS A 144       3.124  23.320 -41.065  1.00  0.00           C  
ATOM   2324  CD  LYS A 144       4.500  22.686 -40.926  1.00  0.00           C  
ATOM   2325  CE  LYS A 144       5.058  22.271 -42.280  1.00  0.00           C  
ATOM   2326  NZ  LYS A 144       6.418  21.677 -42.162  1.00  0.00           N1+
ATOM   2327  H   LYS A 144       2.068  26.207 -40.388  1.00  0.00           H  
ATOM   2328  HA  LYS A 144       0.496  23.702 -40.305  1.00  0.00           H  
ATOM   2329 1HB  LYS A 144       3.265  24.422 -39.251  1.00  0.00           H  
ATOM   2330 2HB  LYS A 144       2.528  22.835 -39.081  1.00  0.00           H  
ATOM   2331 1HG  LYS A 144       2.450  22.605 -41.539  1.00  0.00           H  
ATOM   2332 2HG  LYS A 144       3.201  24.199 -41.705  1.00  0.00           H  
ATOM   2333 1HD  LYS A 144       5.184  23.398 -40.463  1.00  0.00           H  
ATOM   2334 2HD  LYS A 144       4.432  21.806 -40.287  1.00  0.00           H  
ATOM   2335 1HE  LYS A 144       4.394  21.540 -42.738  1.00  0.00           H  
ATOM   2336 2HE  LYS A 144       5.112  23.141 -42.933  1.00  0.00           H  
ATOM   2337 1HZ  LYS A 144       6.752  21.416 -43.079  1.00  0.00           H  
ATOM   2338 2HZ  LYS A 144       7.048  22.353 -41.754  1.00  0.00           H  
ATOM   2339 3HZ  LYS A 144       6.378  20.857 -41.573  1.00  0.00           H  
ATOM   2340  N   VAL A 145       0.429  25.739 -37.908  1.00  0.00           N  
ATOM   2341  CA  VAL A 145      -0.115  26.003 -36.580  1.00  0.00           C  
ATOM   2342  C   VAL A 145      -1.253  27.003 -36.688  1.00  0.00           C  
ATOM   2343  O   VAL A 145      -1.152  27.988 -37.420  1.00  0.00           O  
ATOM   2344  CB  VAL A 145       0.974  26.555 -35.641  1.00  0.00           C  
ATOM   2345  CG1 VAL A 145       0.387  26.874 -34.274  1.00  0.00           C  
ATOM   2346  CG2 VAL A 145       2.111  25.552 -35.520  1.00  0.00           C  
ATOM   2347  H   VAL A 145       0.672  26.517 -38.504  1.00  0.00           H  
ATOM   2348  HA  VAL A 145      -0.476  25.064 -36.160  1.00  0.00           H  
ATOM   2349  HB  VAL A 145       1.356  27.490 -36.053  1.00  0.00           H  
ATOM   2350 1HG1 VAL A 145       1.171  27.264 -33.624  1.00  0.00           H  
ATOM   2351 2HG1 VAL A 145      -0.399  27.621 -34.381  1.00  0.00           H  
ATOM   2352 3HG1 VAL A 145      -0.029  25.967 -33.836  1.00  0.00           H  
ATOM   2353 1HG2 VAL A 145       2.877  25.949 -34.855  1.00  0.00           H  
ATOM   2354 2HG2 VAL A 145       1.727  24.615 -35.114  1.00  0.00           H  
ATOM   2355 3HG2 VAL A 145       2.543  25.370 -36.504  1.00  0.00           H  
ATOM   2356  N   LEU A 146      -2.335  26.733 -35.963  1.00  0.00           N  
ATOM   2357  CA  LEU A 146      -3.510  27.595 -35.965  1.00  0.00           C  
ATOM   2358  C   LEU A 146      -3.565  28.454 -34.711  1.00  0.00           C  
ATOM   2359  O   LEU A 146      -3.187  28.015 -33.626  1.00  0.00           O  
ATOM   2360  CB  LEU A 146      -4.789  26.755 -36.009  1.00  0.00           C  
ATOM   2361  CG  LEU A 146      -5.011  25.801 -37.180  1.00  0.00           C  
ATOM   2362  CD1 LEU A 146      -6.209  24.898 -36.889  1.00  0.00           C  
ATOM   2363  CD2 LEU A 146      -5.239  26.611 -38.440  1.00  0.00           C  
ATOM   2364  H   LEU A 146      -2.337  25.899 -35.392  1.00  0.00           H  
ATOM   2365  HA  LEU A 146      -3.476  28.224 -36.853  1.00  0.00           H  
ATOM   2366 1HB  LEU A 146      -4.831  26.140 -35.112  1.00  0.00           H  
ATOM   2367 2HB  LEU A 146      -5.647  27.428 -36.004  1.00  0.00           H  
ATOM   2368  HG  LEU A 146      -4.133  25.167 -37.304  1.00  0.00           H  
ATOM   2369 1HD1 LEU A 146      -6.364  24.218 -37.726  1.00  0.00           H  
ATOM   2370 2HD1 LEU A 146      -6.017  24.320 -35.985  1.00  0.00           H  
ATOM   2371 3HD1 LEU A 146      -7.099  25.510 -36.747  1.00  0.00           H  
ATOM   2372 1HD2 LEU A 146      -5.398  25.936 -39.283  1.00  0.00           H  
ATOM   2373 2HD2 LEU A 146      -6.117  27.245 -38.312  1.00  0.00           H  
ATOM   2374 3HD2 LEU A 146      -4.366  27.235 -38.635  1.00  0.00           H  
ATOM   2375  N   THR A 147      -4.028  29.687 -34.875  1.00  0.00           N  
ATOM   2376  CA  THR A 147      -4.201  30.596 -33.751  1.00  0.00           C  
ATOM   2377  C   THR A 147      -5.590  31.188 -33.913  1.00  0.00           C  
ATOM   2378  O   THR A 147      -5.913  31.733 -34.968  1.00  0.00           O  
ATOM   2379  CB  THR A 147      -3.131  31.702 -33.712  1.00  0.00           C  
ATOM   2380  CG2 THR A 147      -3.348  32.612 -32.512  1.00  0.00           C  
ATOM   2381  OG1 THR A 147      -1.829  31.108 -33.627  1.00  0.00           O  
ATOM   2382  H   THR A 147      -4.267  30.002 -35.803  1.00  0.00           H  
ATOM   2383  HA  THR A 147      -4.052  30.036 -32.827  1.00  0.00           H  
ATOM   2384  HB  THR A 147      -3.186  32.295 -34.624  1.00  0.00           H  
ATOM   2385  HG1 THR A 147      -1.554  30.816 -34.500  1.00  0.00           H  
ATOM   2386 1HG2 THR A 147      -2.582  33.388 -32.501  1.00  0.00           H  
ATOM   2387 2HG2 THR A 147      -4.332  33.074 -32.579  1.00  0.00           H  
ATOM   2388 3HG2 THR A 147      -3.284  32.027 -31.596  1.00  0.00           H  
ATOM   2389  N   PHE A 148      -6.422  31.056 -32.886  1.00  0.00           N  
ATOM   2390  CA  PHE A 148      -7.763  31.619 -32.936  1.00  0.00           C  
ATOM   2391  C   PHE A 148      -7.843  32.775 -31.951  1.00  0.00           C  
ATOM   2392  O   PHE A 148      -7.313  32.683 -30.845  1.00  0.00           O  
ATOM   2393  CB  PHE A 148      -8.822  30.575 -32.552  1.00  0.00           C  
ATOM   2394  CG  PHE A 148      -8.880  29.389 -33.474  1.00  0.00           C  
ATOM   2395  CD1 PHE A 148      -8.147  28.239 -33.192  1.00  0.00           C  
ATOM   2396  CD2 PHE A 148      -9.673  29.417 -34.616  1.00  0.00           C  
ATOM   2397  CE1 PHE A 148      -8.202  27.127 -34.036  1.00  0.00           C  
ATOM   2398  CE2 PHE A 148      -9.736  28.310 -35.470  1.00  0.00           C  
ATOM   2399  CZ  PHE A 148      -8.999  27.163 -35.176  1.00  0.00           C  
ATOM   2400  H   PHE A 148      -6.127  30.560 -32.057  1.00  0.00           H  
ATOM   2401  HA  PHE A 148      -7.963  31.948 -33.957  1.00  0.00           H  
ATOM   2402 1HB  PHE A 148      -8.624  30.209 -31.545  1.00  0.00           H  
ATOM   2403 2HB  PHE A 148      -9.806  31.042 -32.542  1.00  0.00           H  
ATOM   2404  HD1 PHE A 148      -7.523  28.211 -32.298  1.00  0.00           H  
ATOM   2405  HD2 PHE A 148     -10.248  30.315 -34.844  1.00  0.00           H  
ATOM   2406  HE1 PHE A 148      -7.621  26.235 -33.799  1.00  0.00           H  
ATOM   2407  HE2 PHE A 148     -10.359  28.343 -36.363  1.00  0.00           H  
ATOM   2408  HZ  PHE A 148      -9.046  26.298 -35.836  1.00  0.00           H  
ATOM   2409  N   LEU A 149      -8.490  33.862 -32.364  1.00  0.00           N  
ATOM   2410  CA  LEU A 149      -8.671  35.035 -31.510  1.00  0.00           C  
ATOM   2411  C   LEU A 149     -10.039  35.647 -31.786  1.00  0.00           C  
ATOM   2412  O   LEU A 149     -10.494  35.646 -32.933  1.00  0.00           O  
ATOM   2413  CB  LEU A 149      -7.611  36.105 -31.804  1.00  0.00           C  
ATOM   2414  CG  LEU A 149      -6.138  35.831 -31.502  1.00  0.00           C  
ATOM   2415  CD1 LEU A 149      -5.296  36.972 -32.053  1.00  0.00           C  
ATOM   2416  CD2 LEU A 149      -5.926  35.684 -30.005  1.00  0.00           C  
ATOM   2417  H   LEU A 149      -8.867  33.872 -33.300  1.00  0.00           H  
ATOM   2418  HA  LEU A 149      -8.565  34.726 -30.471  1.00  0.00           H  
ATOM   2419 1HB  LEU A 149      -7.647  36.348 -32.864  1.00  0.00           H  
ATOM   2420 2HB  LEU A 149      -7.858  37.003 -31.238  1.00  0.00           H  
ATOM   2421  HG  LEU A 149      -5.831  34.909 -31.997  1.00  0.00           H  
ATOM   2422 1HD1 LEU A 149      -4.244  36.783 -31.842  1.00  0.00           H  
ATOM   2423 2HD1 LEU A 149      -5.442  37.045 -33.131  1.00  0.00           H  
ATOM   2424 3HD1 LEU A 149      -5.598  37.907 -31.581  1.00  0.00           H  
ATOM   2425 1HD2 LEU A 149      -4.872  35.488 -29.804  1.00  0.00           H  
ATOM   2426 2HD2 LEU A 149      -6.226  36.604 -29.504  1.00  0.00           H  
ATOM   2427 3HD2 LEU A 149      -6.527  34.854 -29.632  1.00  0.00           H  
ATOM   2428  N   ASP A 150     -10.694  36.159 -30.745  1.00  0.00           N  
ATOM   2429  CA  ASP A 150     -11.982  36.819 -30.923  1.00  0.00           C  
ATOM   2430  C   ASP A 150     -11.710  38.127 -31.668  1.00  0.00           C  
ATOM   2431  O   ASP A 150     -10.565  38.590 -31.728  1.00  0.00           O  
ATOM   2432  CB  ASP A 150     -12.669  37.088 -29.582  1.00  0.00           C  
ATOM   2433  CG  ASP A 150     -13.160  35.816 -28.902  1.00  0.00           C  
ATOM   2434  OD1 ASP A 150     -13.212  34.800 -29.554  1.00  0.00           O  
ATOM   2435  OD2 ASP A 150     -13.478  35.874 -27.739  1.00  0.00           O1-
ATOM   2436  H   ASP A 150     -10.299  36.092 -29.818  1.00  0.00           H  
ATOM   2437  HA  ASP A 150     -12.640  36.152 -31.482  1.00  0.00           H  
ATOM   2438 1HB  ASP A 150     -11.975  37.595 -28.912  1.00  0.00           H  
ATOM   2439 2HB  ASP A 150     -13.520  37.752 -29.736  1.00  0.00           H  
ATOM   2440  N   SER A 151     -12.758  38.721 -32.228  1.00  0.00           N  
ATOM   2441  CA  SER A 151     -12.620  39.953 -33.001  1.00  0.00           C  
ATOM   2442  C   SER A 151     -12.594  41.256 -32.198  1.00  0.00           C  
ATOM   2443  O   SER A 151     -12.871  42.328 -32.735  1.00  0.00           O  
ATOM   2444  CB  SER A 151     -13.754  40.024 -34.005  1.00  0.00           C  
ATOM   2445  OG  SER A 151     -14.994  40.101 -33.357  1.00  0.00           O  
ATOM   2446  H   SER A 151     -13.674  38.311 -32.115  1.00  0.00           H  
ATOM   2447  HA  SER A 151     -11.650  39.943 -33.498  1.00  0.00           H  
ATOM   2448 1HB  SER A 151     -13.620  40.897 -34.644  1.00  0.00           H  
ATOM   2449 2HB  SER A 151     -13.729  39.144 -34.645  1.00  0.00           H  
ATOM   2450  HG  SER A 151     -14.966  40.899 -32.824  1.00  0.00           H  
ATOM   2451  N   HIS A 152     -12.248  41.162 -30.919  1.00  0.00           N  
ATOM   2452  CA  HIS A 152     -12.153  42.326 -30.039  1.00  0.00           C  
ATOM   2453  C   HIS A 152     -11.076  42.042 -28.993  1.00  0.00           C  
ATOM   2454  O   HIS A 152     -11.320  42.076 -27.780  1.00  0.00           O  
ATOM   2455  CB  HIS A 152     -13.493  42.608 -29.357  1.00  0.00           C  
ATOM   2456  CG  HIS A 152     -14.279  41.373 -29.045  1.00  0.00           C  
ATOM   2457  CD2 HIS A 152     -14.164  40.476 -28.039  1.00  0.00           C  
ATOM   2458  ND1 HIS A 152     -15.287  40.912 -29.862  1.00  0.00           N  
ATOM   2459  CE1 HIS A 152     -15.760  39.778 -29.373  1.00  0.00           C  
ATOM   2460  NE2 HIS A 152     -15.097  39.491 -28.268  1.00  0.00           N  
ATOM   2461  H   HIS A 152     -12.045  40.246 -30.546  1.00  0.00           H  
ATOM   2462  HA  HIS A 152     -11.884  43.205 -30.625  1.00  0.00           H  
ATOM   2463 1HB  HIS A 152     -13.321  43.148 -28.426  1.00  0.00           H  
ATOM   2464 2HB  HIS A 152     -14.099  43.247 -29.999  1.00  0.00           H  
ATOM   2465  HD2 HIS A 152     -13.464  40.523 -27.205  1.00  0.00           H  
ATOM   2466  HE1 HIS A 152     -16.561  39.181 -29.809  1.00  0.00           H  
ATOM   2467  HE2 HIS A 152     -15.246  38.681 -27.682  1.00  0.00           H  
ATOM   2468  N   CYS A 153      -9.878  41.758 -29.491  1.00  0.00           N  
ATOM   2469  CA  CYS A 153      -8.729  41.456 -28.645  1.00  0.00           C  
ATOM   2470  C   CYS A 153      -7.575  42.392 -28.967  1.00  0.00           C  
ATOM   2471  O   CYS A 153      -7.568  43.053 -30.000  1.00  0.00           O  
ATOM   2472  CB  CYS A 153      -8.278  40.008 -28.835  1.00  0.00           C  
ATOM   2473  SG  CYS A 153      -9.544  38.781 -28.434  1.00  0.00           S  
ATOM   2474  H   CYS A 153      -9.765  41.753 -30.495  1.00  0.00           H  
ATOM   2475  HA  CYS A 153      -9.031  41.560 -27.603  1.00  0.00           H  
ATOM   2476 1HB  CYS A 153      -7.975  39.854 -29.871  1.00  0.00           H  
ATOM   2477 2HB  CYS A 153      -7.408  39.812 -28.208  1.00  0.00           H  
ATOM   2478  HG  CYS A 153      -9.693  39.145 -27.164  1.00  0.00           H  
ATOM   2479  N   GLU A 154      -6.596  42.440 -28.070  1.00  0.00           N  
ATOM   2480  CA  GLU A 154      -5.402  43.249 -28.277  1.00  0.00           C  
ATOM   2481  C   GLU A 154      -4.209  42.467 -27.740  1.00  0.00           C  
ATOM   2482  O   GLU A 154      -4.062  42.285 -26.532  1.00  0.00           O  
ATOM   2483  CB  GLU A 154      -5.520  44.605 -27.577  1.00  0.00           C  
ATOM   2484  CG  GLU A 154      -6.619  45.503 -28.127  1.00  0.00           C  
ATOM   2485  CD  GLU A 154      -6.701  46.827 -27.420  1.00  0.00           C  
ATOM   2486  OE1 GLU A 154      -5.716  47.239 -26.855  1.00  0.00           O  
ATOM   2487  OE2 GLU A 154      -7.749  47.428 -27.445  1.00  0.00           O1-
ATOM   2488  H   GLU A 154      -6.683  41.899 -27.221  1.00  0.00           H  
ATOM   2489  HA  GLU A 154      -5.284  43.425 -29.346  1.00  0.00           H  
ATOM   2490 1HB  GLU A 154      -5.714  44.450 -26.515  1.00  0.00           H  
ATOM   2491 2HB  GLU A 154      -4.576  45.142 -27.662  1.00  0.00           H  
ATOM   2492 1HG  GLU A 154      -6.434  45.680 -29.186  1.00  0.00           H  
ATOM   2493 2HG  GLU A 154      -7.575  44.988 -28.035  1.00  0.00           H  
ATOM   2494  N   CYS A 155      -3.373  41.994 -28.656  1.00  0.00           N  
ATOM   2495  CA  CYS A 155      -2.192  41.207 -28.301  1.00  0.00           C  
ATOM   2496  C   CYS A 155      -1.153  42.069 -27.604  1.00  0.00           C  
ATOM   2497  O   CYS A 155      -0.971  43.237 -27.953  1.00  0.00           O  
ATOM   2498  CB  CYS A 155      -1.568  40.574 -29.545  1.00  0.00           C  
ATOM   2499  SG  CYS A 155      -2.628  39.362 -30.368  1.00  0.00           S  
ATOM   2500  H   CYS A 155      -3.563  42.184 -29.629  1.00  0.00           H  
ATOM   2501  HA  CYS A 155      -2.500  40.403 -27.633  1.00  0.00           H  
ATOM   2502 1HB  CYS A 155      -1.324  41.354 -30.266  1.00  0.00           H  
ATOM   2503 2HB  CYS A 155      -0.636  40.078 -29.272  1.00  0.00           H  
ATOM   2504  HG  CYS A 155      -1.771  39.066 -31.340  1.00  0.00           H  
ATOM   2505  N   ASN A 156      -0.469  41.489 -26.622  1.00  0.00           N  
ATOM   2506  CA  ASN A 156       0.548  42.225 -25.883  1.00  0.00           C  
ATOM   2507  C   ASN A 156       1.954  41.842 -26.337  1.00  0.00           C  
ATOM   2508  O   ASN A 156       2.128  41.058 -27.270  1.00  0.00           O  
ATOM   2509  CB  ASN A 156       0.395  41.972 -24.379  1.00  0.00           C  
ATOM   2510  CG  ASN A 156       0.714  43.202 -23.551  1.00  0.00           C  
ATOM   2511  ND2 ASN A 156      -0.184  43.551 -22.636  1.00  0.00           N  
ATOM   2512  OD1 ASN A 156       1.755  43.831 -23.732  1.00  0.00           O  
ATOM   2513  H   ASN A 156      -0.651  40.525 -26.381  1.00  0.00           H  
ATOM   2514  HA  ASN A 156       0.413  43.290 -26.076  1.00  0.00           H  
ATOM   2515 1HB  ASN A 156      -0.627  41.657 -24.165  1.00  0.00           H  
ATOM   2516 2HB  ASN A 156       1.059  41.161 -24.078  1.00  0.00           H  
ATOM   2517 1HD2 ASN A 156      -0.025  44.354 -22.061  1.00  0.00           H  
ATOM   2518 2HD2 ASN A 156      -1.017  43.012 -22.523  1.00  0.00           H  
ATOM   2519  N   GLU A 157       2.947  42.402 -25.652  1.00  0.00           N  
ATOM   2520  CA  GLU A 157       4.359  42.183 -25.951  1.00  0.00           C  
ATOM   2521  C   GLU A 157       4.770  40.709 -25.970  1.00  0.00           C  
ATOM   2522  O   GLU A 157       4.560  39.982 -24.997  1.00  0.00           O  
ATOM   2523  CB  GLU A 157       5.220  42.933 -24.932  1.00  0.00           C  
ATOM   2524  CG  GLU A 157       5.164  44.449 -25.057  1.00  0.00           C  
ATOM   2525  CD  GLU A 157       5.853  44.960 -26.291  1.00  0.00           C  
ATOM   2526  OE1 GLU A 157       6.978  44.587 -26.518  1.00  0.00           O  
ATOM   2527  OE2 GLU A 157       5.253  45.726 -27.008  1.00  0.00           O1-
ATOM   2528  H   GLU A 157       2.695  43.010 -24.887  1.00  0.00           H  
ATOM   2529  HA  GLU A 157       4.567  42.573 -26.949  1.00  0.00           H  
ATOM   2530 1HB  GLU A 157       4.903  42.669 -23.923  1.00  0.00           H  
ATOM   2531 2HB  GLU A 157       6.260  42.627 -25.039  1.00  0.00           H  
ATOM   2532 1HG  GLU A 157       4.121  44.763 -25.080  1.00  0.00           H  
ATOM   2533 2HG  GLU A 157       5.628  44.893 -24.177  1.00  0.00           H  
ATOM   2534  N   HIS A 158       5.363  40.279 -27.082  1.00  0.00           N  
ATOM   2535  CA  HIS A 158       5.819  38.898 -27.248  1.00  0.00           C  
ATOM   2536  C   HIS A 158       4.785  37.869 -26.809  1.00  0.00           C  
ATOM   2537  O   HIS A 158       5.106  36.891 -26.132  1.00  0.00           O  
ATOM   2538  CB  HIS A 158       7.116  38.671 -26.465  1.00  0.00           C  
ATOM   2539  CG  HIS A 158       8.248  39.545 -26.909  1.00  0.00           C  
ATOM   2540  CD2 HIS A 158       8.855  40.593 -26.305  1.00  0.00           C  
ATOM   2541  ND1 HIS A 158       8.886  39.377 -28.120  1.00  0.00           N  
ATOM   2542  CE1 HIS A 158       9.838  40.286 -28.242  1.00  0.00           C  
ATOM   2543  NE2 HIS A 158       9.840  41.035 -27.154  1.00  0.00           N  
ATOM   2544  H   HIS A 158       5.500  40.937 -27.836  1.00  0.00           H  
ATOM   2545  HA  HIS A 158       6.008  38.702 -28.303  1.00  0.00           H  
ATOM   2546 1HB  HIS A 158       6.939  38.855 -25.405  1.00  0.00           H  
ATOM   2547 2HB  HIS A 158       7.427  37.632 -26.569  1.00  0.00           H  
ATOM   2548  HD1 HIS A 158       8.629  38.732 -28.840  1.00  0.00           H  
ATOM   2549  HD2 HIS A 158       8.698  41.088 -25.346  1.00  0.00           H  
ATOM   2550  HE1 HIS A 158      10.461  40.322 -29.135  1.00  0.00           H  
ATOM   2551  N   TRP A 159       3.542  38.087 -27.221  1.00  0.00           N  
ATOM   2552  CA  TRP A 159       2.446  37.188 -26.870  1.00  0.00           C  
ATOM   2553  C   TRP A 159       2.480  35.856 -27.614  1.00  0.00           C  
ATOM   2554  O   TRP A 159       2.081  34.822 -27.076  1.00  0.00           O  
ATOM   2555  CB  TRP A 159       1.106  37.855 -27.189  1.00  0.00           C  
ATOM   2556  CG  TRP A 159       0.862  37.957 -28.673  1.00  0.00           C  
ATOM   2557  CD1 TRP A 159       1.468  38.813 -29.551  1.00  0.00           C  
ATOM   2558  CD2 TRP A 159       0.005  37.120 -29.460  1.00  0.00           C  
ATOM   2559  CE2 TRP A 159       0.146  37.523 -30.807  1.00  0.00           C  
ATOM   2560  CE3 TRP A 159      -0.866  36.063 -29.158  1.00  0.00           C  
ATOM   2561  NE1 TRP A 159       1.043  38.557 -30.834  1.00  0.00           N  
ATOM   2562  CZ2 TRP A 159      -0.556  36.910 -31.854  1.00  0.00           C  
ATOM   2563  CZ3 TRP A 159      -1.568  35.449 -30.203  1.00  0.00           C  
ATOM   2564  CH2 TRP A 159      -1.404  35.878 -31.533  1.00  0.00           C  
ATOM   2565  H   TRP A 159       3.350  38.897 -27.794  1.00  0.00           H  
ATOM   2566  HA  TRP A 159       2.495  36.985 -25.801  1.00  0.00           H  
ATOM   2567 1HB  TRP A 159       0.297  37.284 -26.733  1.00  0.00           H  
ATOM   2568 2HB  TRP A 159       1.085  38.855 -26.756  1.00  0.00           H  
ATOM   2569  HD1 TRP A 159       2.183  39.585 -29.275  1.00  0.00           H  
ATOM   2570  HE1 TRP A 159       1.342  39.052 -31.662  1.00  0.00           H  
ATOM   2571  HE3 TRP A 159      -0.991  35.725 -28.130  1.00  0.00           H  
ATOM   2572  HZ2 TRP A 159      -0.443  37.225 -32.890  1.00  0.00           H  
ATOM   2573  HZ3 TRP A 159      -2.245  34.630 -29.961  1.00  0.00           H  
ATOM   2574  HH2 TRP A 159      -1.965  35.381 -32.325  1.00  0.00           H  
ATOM   2575  N   LEU A 160       2.966  35.881 -28.850  1.00  0.00           N  
ATOM   2576  CA  LEU A 160       2.956  34.686 -29.687  1.00  0.00           C  
ATOM   2577  C   LEU A 160       4.046  33.646 -29.479  1.00  0.00           C  
ATOM   2578  O   LEU A 160       3.757  32.451 -29.388  1.00  0.00           O  
ATOM   2579  CB  LEU A 160       2.936  35.109 -31.161  1.00  0.00           C  
ATOM   2580  CG  LEU A 160       2.746  34.006 -32.201  1.00  0.00           C  
ATOM   2581  CD1 LEU A 160       1.436  33.268 -31.937  1.00  0.00           C  
ATOM   2582  CD2 LEU A 160       2.755  34.615 -33.595  1.00  0.00           C  
ATOM   2583  H   LEU A 160       3.349  36.738 -29.221  1.00  0.00           H  
ATOM   2584  HA  LEU A 160       2.056  34.114 -29.464  1.00  0.00           H  
ATOM   2585 1HB  LEU A 160       2.128  35.825 -31.307  1.00  0.00           H  
ATOM   2586 2HB  LEU A 160       3.879  35.605 -31.393  1.00  0.00           H  
ATOM   2587  HG  LEU A 160       3.556  33.282 -32.116  1.00  0.00           H  
ATOM   2588 1HD1 LEU A 160       1.305  32.482 -32.681  1.00  0.00           H  
ATOM   2589 2HD1 LEU A 160       1.463  32.824 -30.942  1.00  0.00           H  
ATOM   2590 3HD1 LEU A 160       0.605  33.970 -32.000  1.00  0.00           H  
ATOM   2591 1HD2 LEU A 160       2.620  33.828 -34.337  1.00  0.00           H  
ATOM   2592 2HD2 LEU A 160       1.944  35.338 -33.682  1.00  0.00           H  
ATOM   2593 3HD2 LEU A 160       3.708  35.115 -33.767  1.00  0.00           H  
ATOM   2594  N   GLU A 161       5.292  34.087 -29.394  1.00  0.00           N  
ATOM   2595  CA  GLU A 161       6.392  33.147 -29.240  1.00  0.00           C  
ATOM   2596  C   GLU A 161       6.236  32.121 -28.115  1.00  0.00           C  
ATOM   2597  O   GLU A 161       6.500  30.938 -28.316  1.00  0.00           O  
ATOM   2598  CB  GLU A 161       7.713  33.904 -29.082  1.00  0.00           C  
ATOM   2599  CG  GLU A 161       8.032  34.841 -30.239  1.00  0.00           C  
ATOM   2600  CD  GLU A 161       7.677  36.285 -29.936  1.00  0.00           C  
ATOM   2601  OE1 GLU A 161       6.557  36.532 -29.437  1.00  0.00           O  
ATOM   2602  OE2 GLU A 161       8.518  37.172 -30.206  1.00  0.00           O1-
ATOM   2603  H   GLU A 161       5.492  35.076 -29.434  1.00  0.00           H  
ATOM   2604  HA  GLU A 161       6.446  32.528 -30.137  1.00  0.00           H  
ATOM   2605 1HB  GLU A 161       7.686  34.494 -28.166  1.00  0.00           H  
ATOM   2606 2HB  GLU A 161       8.532  33.192 -28.988  1.00  0.00           H  
ATOM   2607 1HG  GLU A 161       9.097  34.777 -30.460  1.00  0.00           H  
ATOM   2608 2HG  GLU A 161       7.485  34.512 -31.121  1.00  0.00           H  
ATOM   2609  N   PRO A 162       5.808  32.553 -26.919  1.00  0.00           N  
ATOM   2610  CA  PRO A 162       5.657  31.582 -25.829  1.00  0.00           C  
ATOM   2611  C   PRO A 162       4.640  30.489 -26.147  1.00  0.00           C  
ATOM   2612  O   PRO A 162       4.781  29.348 -25.700  1.00  0.00           O  
ATOM   2613  CB  PRO A 162       5.212  32.445 -24.654  1.00  0.00           C  
ATOM   2614  CG  PRO A 162       5.838  33.771 -24.954  1.00  0.00           C  
ATOM   2615  CD  PRO A 162       5.590  33.926 -26.430  1.00  0.00           C  
ATOM   2616  HA  PRO A 162       6.630  31.112 -25.624  1.00  0.00           H  
ATOM   2617 1HB  PRO A 162       4.113  32.480 -24.608  1.00  0.00           H  
ATOM   2618 2HB  PRO A 162       5.558  32.003 -23.708  1.00  0.00           H  
ATOM   2619 1HG  PRO A 162       5.369  34.559 -24.346  1.00  0.00           H  
ATOM   2620 2HG  PRO A 162       6.904  33.757 -24.687  1.00  0.00           H  
ATOM   2621 1HD  PRO A 162       4.557  34.267 -26.595  1.00  0.00           H  
ATOM   2622 2HD  PRO A 162       6.308  34.645 -26.850  1.00  0.00           H  
ATOM   2623  N   LEU A 163       3.608  30.843 -26.907  1.00  0.00           N  
ATOM   2624  CA  LEU A 163       2.582  29.870 -27.264  1.00  0.00           C  
ATOM   2625  C   LEU A 163       3.127  28.892 -28.295  1.00  0.00           C  
ATOM   2626  O   LEU A 163       2.991  27.675 -28.144  1.00  0.00           O  
ATOM   2627  CB  LEU A 163       1.340  30.579 -27.819  1.00  0.00           C  
ATOM   2628  CG  LEU A 163       0.653  31.575 -26.874  1.00  0.00           C  
ATOM   2629  CD1 LEU A 163      -0.522  32.216 -27.595  1.00  0.00           C  
ATOM   2630  CD2 LEU A 163       0.175  30.870 -25.602  1.00  0.00           C  
ATOM   2631  H   LEU A 163       3.526  31.791 -27.245  1.00  0.00           H  
ATOM   2632  HA  LEU A 163       2.299  29.323 -26.366  1.00  0.00           H  
ATOM   2633 1HB  LEU A 163       1.623  31.123 -28.719  1.00  0.00           H  
ATOM   2634 2HB  LEU A 163       0.603  29.825 -28.093  1.00  0.00           H  
ATOM   2635  HG  LEU A 163       1.357  32.361 -26.598  1.00  0.00           H  
ATOM   2636 1HD1 LEU A 163      -1.015  32.925 -26.930  1.00  0.00           H  
ATOM   2637 2HD1 LEU A 163      -0.163  32.741 -28.480  1.00  0.00           H  
ATOM   2638 3HD1 LEU A 163      -1.232  31.445 -27.893  1.00  0.00           H  
ATOM   2639 1HD2 LEU A 163      -0.308  31.593 -24.945  1.00  0.00           H  
ATOM   2640 2HD2 LEU A 163      -0.535  30.086 -25.865  1.00  0.00           H  
ATOM   2641 3HD2 LEU A 163       1.029  30.427 -25.089  1.00  0.00           H  
ATOM   2642  N   LEU A 164       3.755  29.426 -29.337  1.00  0.00           N  
ATOM   2643  CA  LEU A 164       4.314  28.590 -30.393  1.00  0.00           C  
ATOM   2644  C   LEU A 164       5.368  27.618 -29.860  1.00  0.00           C  
ATOM   2645  O   LEU A 164       5.442  26.476 -30.311  1.00  0.00           O  
ATOM   2646  CB  LEU A 164       4.933  29.473 -31.484  1.00  0.00           C  
ATOM   2647  CG  LEU A 164       3.941  30.289 -32.322  1.00  0.00           C  
ATOM   2648  CD1 LEU A 164       4.707  31.236 -33.236  1.00  0.00           C  
ATOM   2649  CD2 LEU A 164       3.059  29.346 -33.126  1.00  0.00           C  
ATOM   2650  H   LEU A 164       3.848  30.429 -29.402  1.00  0.00           H  
ATOM   2651  HA  LEU A 164       3.511  27.990 -30.820  1.00  0.00           H  
ATOM   2652 1HB  LEU A 164       5.624  30.171 -31.015  1.00  0.00           H  
ATOM   2653 2HB  LEU A 164       5.499  28.837 -32.166  1.00  0.00           H  
ATOM   2654  HG  LEU A 164       3.319  30.894 -31.662  1.00  0.00           H  
ATOM   2655 1HD1 LEU A 164       4.002  31.816 -33.832  1.00  0.00           H  
ATOM   2656 2HD1 LEU A 164       5.314  31.911 -32.634  1.00  0.00           H  
ATOM   2657 3HD1 LEU A 164       5.352  30.659 -33.898  1.00  0.00           H  
ATOM   2658 1HD2 LEU A 164       2.353  29.926 -33.721  1.00  0.00           H  
ATOM   2659 2HD2 LEU A 164       3.680  28.742 -33.787  1.00  0.00           H  
ATOM   2660 3HD2 LEU A 164       2.510  28.693 -32.447  1.00  0.00           H  
ATOM   2661  N   GLU A 165       6.172  28.070 -28.899  1.00  0.00           N  
ATOM   2662  CA  GLU A 165       7.214  27.229 -28.317  1.00  0.00           C  
ATOM   2663  C   GLU A 165       6.635  25.964 -27.689  1.00  0.00           C  
ATOM   2664  O   GLU A 165       7.207  24.880 -27.815  1.00  0.00           O  
ATOM   2665  CB  GLU A 165       8.005  28.014 -27.262  1.00  0.00           C  
ATOM   2666  CG  GLU A 165       8.960  27.163 -26.432  1.00  0.00           C  
ATOM   2667  CD  GLU A 165      10.116  26.585 -27.238  1.00  0.00           C  
ATOM   2668  OE1 GLU A 165      10.803  25.678 -26.719  1.00  0.00           O  
ATOM   2669  OE2 GLU A 165      10.344  27.039 -28.379  1.00  0.00           O1-
ATOM   2670  H   GLU A 165       6.059  29.017 -28.565  1.00  0.00           H  
ATOM   2671  HA  GLU A 165       7.896  26.924 -29.111  1.00  0.00           H  
ATOM   2672 1HB  GLU A 165       8.590  28.793 -27.752  1.00  0.00           H  
ATOM   2673 2HB  GLU A 165       7.312  28.505 -26.578  1.00  0.00           H  
ATOM   2674 1HG  GLU A 165       9.369  27.775 -25.628  1.00  0.00           H  
ATOM   2675 2HG  GLU A 165       8.401  26.346 -25.979  1.00  0.00           H  
ATOM   2676  N   ARG A 166       5.502  26.102 -27.008  1.00  0.00           N  
ATOM   2677  CA  ARG A 166       4.866  24.954 -26.373  1.00  0.00           C  
ATOM   2678  C   ARG A 166       4.435  23.923 -27.410  1.00  0.00           C  
ATOM   2679  O   ARG A 166       4.688  22.728 -27.254  1.00  0.00           O  
ATOM   2680  CB  ARG A 166       3.644  25.395 -25.559  1.00  0.00           C  
ATOM   2681  CG  ARG A 166       3.963  26.353 -24.418  1.00  0.00           C  
ATOM   2682  CD  ARG A 166       4.964  25.748 -23.451  1.00  0.00           C  
ATOM   2683  NE  ARG A 166       4.453  24.546 -22.797  1.00  0.00           N  
ATOM   2684  CZ  ARG A 166       5.207  23.696 -22.105  1.00  0.00           C  
ATOM   2685  NH1 ARG A 166       4.661  22.631 -21.538  1.00  0.00           N1+
ATOM   2686  NH2 ARG A 166       6.515  23.904 -21.995  1.00  0.00           N  
ATOM   2687  H   ARG A 166       5.071  27.012 -26.924  1.00  0.00           H  
ATOM   2688  HA  ARG A 166       5.585  24.490 -25.697  1.00  0.00           H  
ATOM   2689 1HB  ARG A 166       2.927  25.885 -26.216  1.00  0.00           H  
ATOM   2690 2HB  ARG A 166       3.154  24.519 -25.134  1.00  0.00           H  
ATOM   2691 1HG  ARG A 166       4.385  27.273 -24.823  1.00  0.00           H  
ATOM   2692 2HG  ARG A 166       3.048  26.583 -23.870  1.00  0.00           H  
ATOM   2693 1HD  ARG A 166       5.872  25.477 -23.990  1.00  0.00           H  
ATOM   2694 2HD  ARG A 166       5.206  26.474 -22.677  1.00  0.00           H  
ATOM   2695  HE  ARG A 166       3.464  24.352 -22.878  1.00  0.00           H  
ATOM   2696 1HH1 ARG A 166       3.669  22.462 -21.629  1.00  0.00           H  
ATOM   2697 2HH1 ARG A 166       5.236  21.988 -21.014  1.00  0.00           H  
ATOM   2698 1HH2 ARG A 166       6.938  24.708 -22.436  1.00  0.00           H  
ATOM   2699 2HH2 ARG A 166       7.085  23.258 -21.469  1.00  0.00           H  
ATOM   2700  N   VAL A 167       3.790  24.395 -28.474  1.00  0.00           N  
ATOM   2701  CA  VAL A 167       3.297  23.521 -29.529  1.00  0.00           C  
ATOM   2702  C   VAL A 167       4.412  22.920 -30.375  1.00  0.00           C  
ATOM   2703  O   VAL A 167       4.296  21.791 -30.855  1.00  0.00           O  
ATOM   2704  CB  VAL A 167       2.339  24.303 -30.447  1.00  0.00           C  
ATOM   2705  CG1 VAL A 167       1.939  23.455 -31.645  1.00  0.00           C  
ATOM   2706  CG2 VAL A 167       1.112  24.741 -29.663  1.00  0.00           C  
ATOM   2707  H   VAL A 167       3.638  25.391 -28.550  1.00  0.00           H  
ATOM   2708  HA  VAL A 167       2.768  22.686 -29.067  1.00  0.00           H  
ATOM   2709  HB  VAL A 167       2.857  25.181 -30.834  1.00  0.00           H  
ATOM   2710 1HG1 VAL A 167       1.262  24.023 -32.284  1.00  0.00           H  
ATOM   2711 2HG1 VAL A 167       2.829  23.183 -32.212  1.00  0.00           H  
ATOM   2712 3HG1 VAL A 167       1.437  22.551 -31.300  1.00  0.00           H  
ATOM   2713 1HG2 VAL A 167       0.439  25.294 -30.318  1.00  0.00           H  
ATOM   2714 2HG2 VAL A 167       0.598  23.863 -29.271  1.00  0.00           H  
ATOM   2715 3HG2 VAL A 167       1.418  25.381 -28.835  1.00  0.00           H  
ATOM   2716  N   ALA A 168       5.491  23.673 -30.560  1.00  0.00           N  
ATOM   2717  CA  ALA A 168       6.619  23.189 -31.346  1.00  0.00           C  
ATOM   2718  C   ALA A 168       7.307  22.042 -30.610  1.00  0.00           C  
ATOM   2719  O   ALA A 168       7.854  21.131 -31.231  1.00  0.00           O  
ATOM   2720  CB  ALA A 168       7.609  24.321 -31.592  1.00  0.00           C  
ATOM   2721  H   ALA A 168       5.535  24.596 -30.152  1.00  0.00           H  
ATOM   2722  HA  ALA A 168       6.238  22.836 -32.303  1.00  0.00           H  
ATOM   2723 1HB  ALA A 168       8.448  23.948 -32.180  1.00  0.00           H  
ATOM   2724 2HB  ALA A 168       7.114  25.125 -32.136  1.00  0.00           H  
ATOM   2725 3HB  ALA A 168       7.973  24.699 -30.639  1.00  0.00           H  
ATOM   2726  N   GLU A 169       7.268  22.092 -29.284  1.00  0.00           N  
ATOM   2727  CA  GLU A 169       7.888  21.059 -28.465  1.00  0.00           C  
ATOM   2728  C   GLU A 169       7.026  19.803 -28.390  1.00  0.00           C  
ATOM   2729  O   GLU A 169       7.540  18.681 -28.429  1.00  0.00           O  
ATOM   2730  CB  GLU A 169       8.151  21.591 -27.054  1.00  0.00           C  
ATOM   2731  CG  GLU A 169       8.882  20.616 -26.141  1.00  0.00           C  
ATOM   2732  CD  GLU A 169       9.244  21.219 -24.813  1.00  0.00           C  
ATOM   2733  OE1 GLU A 169       8.358  21.656 -24.118  1.00  0.00           O  
ATOM   2734  OE2 GLU A 169      10.409  21.243 -24.491  1.00  0.00           O1-
ATOM   2735  H   GLU A 169       6.799  22.864 -28.831  1.00  0.00           H  
ATOM   2736  HA  GLU A 169       8.838  20.778 -28.922  1.00  0.00           H  
ATOM   2737 1HB  GLU A 169       8.745  22.503 -27.114  1.00  0.00           H  
ATOM   2738 2HB  GLU A 169       7.204  21.848 -26.580  1.00  0.00           H  
ATOM   2739 1HG  GLU A 169       8.247  19.747 -25.972  1.00  0.00           H  
ATOM   2740 2HG  GLU A 169       9.789  20.277 -26.641  1.00  0.00           H  
ATOM   2741  N   ASP A 170       5.716  19.998 -28.277  1.00  0.00           N  
ATOM   2742  CA  ASP A 170       4.768  18.887 -28.205  1.00  0.00           C  
ATOM   2743  C   ASP A 170       3.499  19.303 -28.948  1.00  0.00           C  
ATOM   2744  O   ASP A 170       2.758  20.175 -28.488  1.00  0.00           O  
ATOM   2745  CB  ASP A 170       4.445  18.520 -26.754  1.00  0.00           C  
ATOM   2746  CG  ASP A 170       3.532  17.307 -26.639  1.00  0.00           C  
ATOM   2747  OD1 ASP A 170       3.051  16.852 -27.650  1.00  0.00           O  
ATOM   2748  OD2 ASP A 170       3.325  16.847 -25.542  1.00  0.00           O1-
ATOM   2749  H   ASP A 170       5.369  20.945 -28.239  1.00  0.00           H  
ATOM   2750  HA  ASP A 170       5.230  18.008 -28.656  1.00  0.00           H  
ATOM   2751 1HB  ASP A 170       5.371  18.313 -26.217  1.00  0.00           H  
ATOM   2752 2HB  ASP A 170       3.964  19.367 -26.263  1.00  0.00           H  
ATOM   2753  N   ARG A 171       3.250  18.662 -30.085  1.00  0.00           N  
ATOM   2754  CA  ARG A 171       2.097  18.970 -30.929  1.00  0.00           C  
ATOM   2755  C   ARG A 171       0.730  18.703 -30.301  1.00  0.00           C  
ATOM   2756  O   ARG A 171      -0.287  19.167 -30.819  1.00  0.00           O  
ATOM   2757  CB  ARG A 171       2.192  18.173 -32.222  1.00  0.00           C  
ATOM   2758  CG  ARG A 171       3.309  18.605 -33.159  1.00  0.00           C  
ATOM   2759  CD  ARG A 171       3.357  17.762 -34.381  1.00  0.00           C  
ATOM   2760  NE  ARG A 171       4.424  18.171 -35.280  1.00  0.00           N  
ATOM   2761  CZ  ARG A 171       4.774  17.512 -36.401  1.00  0.00           C  
ATOM   2762  NH1 ARG A 171       4.135  16.417 -36.747  1.00  0.00           N1+
ATOM   2763  NH2 ARG A 171       5.761  17.966 -37.154  1.00  0.00           N  
ATOM   2764  H   ARG A 171       3.888  17.933 -30.371  1.00  0.00           H  
ATOM   2765  HA  ARG A 171       2.109  20.037 -31.156  1.00  0.00           H  
ATOM   2766 1HB  ARG A 171       2.345  17.120 -31.990  1.00  0.00           H  
ATOM   2767 2HB  ARG A 171       1.252  18.255 -32.769  1.00  0.00           H  
ATOM   2768 1HG  ARG A 171       3.150  19.640 -33.462  1.00  0.00           H  
ATOM   2769 2HG  ARG A 171       4.267  18.520 -32.646  1.00  0.00           H  
ATOM   2770 1HD  ARG A 171       3.527  16.724 -34.100  1.00  0.00           H  
ATOM   2771 2HD  ARG A 171       2.411  17.843 -34.916  1.00  0.00           H  
ATOM   2772  HE  ARG A 171       4.939  19.010 -35.047  1.00  0.00           H  
ATOM   2773 1HH1 ARG A 171       3.380  16.069 -36.172  1.00  0.00           H  
ATOM   2774 2HH1 ARG A 171       4.398  15.922 -37.587  1.00  0.00           H  
ATOM   2775 1HH2 ARG A 171       6.253  18.808 -36.888  1.00  0.00           H  
ATOM   2776 2HH2 ARG A 171       6.024  17.472 -37.994  1.00  0.00           H  
ATOM   2777  N   THR A 172       0.697  17.954 -29.203  1.00  0.00           N  
ATOM   2778  CA  THR A 172      -0.572  17.631 -28.558  1.00  0.00           C  
ATOM   2779  C   THR A 172      -0.976  18.624 -27.483  1.00  0.00           C  
ATOM   2780  O   THR A 172      -1.972  18.423 -26.787  1.00  0.00           O  
ATOM   2781  CB  THR A 172      -0.521  16.221 -27.941  1.00  0.00           C  
ATOM   2782  CG2 THR A 172      -0.262  15.177 -29.017  1.00  0.00           C  
ATOM   2783  OG1 THR A 172       0.525  16.162 -26.962  1.00  0.00           O  
ATOM   2784  H   THR A 172       1.555  17.602 -28.804  1.00  0.00           H  
ATOM   2785  HA  THR A 172      -1.360  17.662 -29.311  1.00  0.00           H  
ATOM   2786  HB  THR A 172      -1.470  16.003 -27.453  1.00  0.00           H  
ATOM   2787  HG1 THR A 172       1.366  16.359 -27.381  1.00  0.00           H  
ATOM   2788 1HG2 THR A 172      -0.229  14.187 -28.563  1.00  0.00           H  
ATOM   2789 2HG2 THR A 172      -1.061  15.211 -29.757  1.00  0.00           H  
ATOM   2790 3HG2 THR A 172       0.691  15.385 -29.503  1.00  0.00           H  
ATOM   2791  N   ARG A 173      -0.199  19.693 -27.348  1.00  0.00           N  
ATOM   2792  CA  ARG A 173      -0.503  20.741 -26.379  1.00  0.00           C  
ATOM   2793  C   ARG A 173      -1.344  21.825 -27.038  1.00  0.00           C  
ATOM   2794  O   ARG A 173      -0.963  22.372 -28.072  1.00  0.00           O  
ATOM   2795  CB  ARG A 173       0.778  21.396 -25.863  1.00  0.00           C  
ATOM   2796  CG  ARG A 173       1.624  20.534 -24.955  1.00  0.00           C  
ATOM   2797  CD  ARG A 173       2.975  21.191 -24.766  1.00  0.00           C  
ATOM   2798  NE  ARG A 173       3.871  20.405 -23.926  1.00  0.00           N  
ATOM   2799  CZ  ARG A 173       5.174  20.637 -23.826  1.00  0.00           C  
ATOM   2800  NH1 ARG A 173       5.930  19.883 -23.038  1.00  0.00           N1+
ATOM   2801  NH2 ARG A 173       5.725  21.619 -24.525  1.00  0.00           N  
ATOM   2802  H   ARG A 173       0.622  19.784 -27.930  1.00  0.00           H  
ATOM   2803  HA  ARG A 173      -1.024  20.291 -25.534  1.00  0.00           H  
ATOM   2804 1HB  ARG A 173       1.402  21.689 -26.706  1.00  0.00           H  
ATOM   2805 2HB  ARG A 173       0.527  22.301 -25.310  1.00  0.00           H  
ATOM   2806 1HG  ARG A 173       1.129  20.429 -23.989  1.00  0.00           H  
ATOM   2807 2HG  ARG A 173       1.754  19.550 -25.405  1.00  0.00           H  
ATOM   2808 1HD  ARG A 173       3.455  21.320 -25.736  1.00  0.00           H  
ATOM   2809 2HD  ARG A 173       2.843  22.164 -24.294  1.00  0.00           H  
ATOM   2810  HE  ARG A 173       3.470  19.643 -23.394  1.00  0.00           H  
ATOM   2811 1HH1 ARG A 173       5.514  19.128 -22.510  1.00  0.00           H  
ATOM   2812 2HH1 ARG A 173       6.920  20.063 -22.964  1.00  0.00           H  
ATOM   2813 1HH2 ARG A 173       5.154  22.190 -25.133  1.00  0.00           H  
ATOM   2814 2HH2 ARG A 173       6.715  21.797 -24.450  1.00  0.00           H  
ATOM   2815  N   VAL A 174      -2.488  22.127 -26.434  1.00  0.00           N  
ATOM   2816  CA  VAL A 174      -3.369  23.180 -26.927  1.00  0.00           C  
ATOM   2817  C   VAL A 174      -3.171  24.273 -25.886  1.00  0.00           C  
ATOM   2818  O   VAL A 174      -3.494  24.087 -24.713  1.00  0.00           O  
ATOM   2819  CB  VAL A 174      -4.842  22.736 -27.009  1.00  0.00           C  
ATOM   2820  CG1 VAL A 174      -5.716  23.880 -27.502  1.00  0.00           C  
ATOM   2821  CG2 VAL A 174      -4.968  21.528 -27.924  1.00  0.00           C  
ATOM   2822  H   VAL A 174      -2.753  21.610 -25.608  1.00  0.00           H  
ATOM   2823  HA  VAL A 174      -3.066  23.439 -27.942  1.00  0.00           H  
ATOM   2824  HB  VAL A 174      -5.189  22.474 -26.010  1.00  0.00           H  
ATOM   2825 1HG1 VAL A 174      -6.753  23.549 -27.553  1.00  0.00           H  
ATOM   2826 2HG1 VAL A 174      -5.638  24.720 -26.812  1.00  0.00           H  
ATOM   2827 3HG1 VAL A 174      -5.385  24.190 -28.493  1.00  0.00           H  
ATOM   2828 1HG2 VAL A 174      -6.012  21.219 -27.977  1.00  0.00           H  
ATOM   2829 2HG2 VAL A 174      -4.615  21.789 -28.922  1.00  0.00           H  
ATOM   2830 3HG2 VAL A 174      -4.367  20.708 -27.530  1.00  0.00           H  
ATOM   2831  N   VAL A 175      -2.632  25.410 -26.316  1.00  0.00           N  
ATOM   2832  CA  VAL A 175      -2.310  26.472 -25.377  1.00  0.00           C  
ATOM   2833  C   VAL A 175      -3.024  27.790 -25.570  1.00  0.00           C  
ATOM   2834  O   VAL A 175      -3.418  28.158 -26.675  1.00  0.00           O  
ATOM   2835  CB  VAL A 175      -0.786  26.722 -25.381  1.00  0.00           C  
ATOM   2836  CG1 VAL A 175      -0.061  25.419 -25.091  1.00  0.00           C  
ATOM   2837  CG2 VAL A 175      -0.340  27.281 -26.731  1.00  0.00           C  
ATOM   2838  H   VAL A 175      -2.441  25.547 -27.298  1.00  0.00           H  
ATOM   2839  HA  VAL A 175      -2.617  26.156 -24.379  1.00  0.00           H  
ATOM   2840  HB  VAL A 175      -0.543  27.440 -24.597  1.00  0.00           H  
ATOM   2841 1HG1 VAL A 175       1.016  25.593 -25.093  1.00  0.00           H  
ATOM   2842 2HG1 VAL A 175      -0.365  25.042 -24.114  1.00  0.00           H  
ATOM   2843 3HG1 VAL A 175      -0.310  24.686 -25.858  1.00  0.00           H  
ATOM   2844 1HG2 VAL A 175       0.737  27.452 -26.717  1.00  0.00           H  
ATOM   2845 2HG2 VAL A 175      -0.583  26.567 -27.519  1.00  0.00           H  
ATOM   2846 3HG2 VAL A 175      -0.854  28.223 -26.923  1.00  0.00           H  
ATOM   2847  N   SER A 176      -3.166  28.511 -24.467  1.00  0.00           N  
ATOM   2848  CA  SER A 176      -3.844  29.791 -24.484  1.00  0.00           C  
ATOM   2849  C   SER A 176      -3.100  30.804 -23.636  1.00  0.00           C  
ATOM   2850  O   SER A 176      -2.382  30.449 -22.702  1.00  0.00           O  
ATOM   2851  CB  SER A 176      -5.266  29.638 -23.980  1.00  0.00           C  
ATOM   2852  OG  SER A 176      -5.283  29.239 -22.637  1.00  0.00           O  
ATOM   2853  H   SER A 176      -2.794  28.162 -23.595  1.00  0.00           H  
ATOM   2854  HA  SER A 176      -3.889  30.145 -25.515  1.00  0.00           H  
ATOM   2855 1HB  SER A 176      -5.794  30.585 -24.088  1.00  0.00           H  
ATOM   2856 2HB  SER A 176      -5.789  28.901 -24.588  1.00  0.00           H  
ATOM   2857  HG  SER A 176      -4.822  28.397 -22.606  1.00  0.00           H  
ATOM   2858  N   PRO A 177      -3.250  32.088 -23.966  1.00  0.00           N  
ATOM   2859  CA  PRO A 177      -2.580  33.140 -23.206  1.00  0.00           C  
ATOM   2860  C   PRO A 177      -3.407  33.458 -21.974  1.00  0.00           C  
ATOM   2861  O   PRO A 177      -4.540  32.981 -21.837  1.00  0.00           O  
ATOM   2862  CB  PRO A 177      -2.588  34.311 -24.181  1.00  0.00           C  
ATOM   2863  CG  PRO A 177      -3.932  34.141 -24.855  1.00  0.00           C  
ATOM   2864  CD  PRO A 177      -3.986  32.644 -25.118  1.00  0.00           C  
ATOM   2865  HA  PRO A 177      -1.552  32.822 -22.976  1.00  0.00           H  
ATOM   2866 1HB  PRO A 177      -2.479  35.258 -23.632  1.00  0.00           H  
ATOM   2867 2HB  PRO A 177      -1.730  34.236 -24.866  1.00  0.00           H  
ATOM   2868 1HG  PRO A 177      -4.736  34.501 -24.196  1.00  0.00           H  
ATOM   2869 2HG  PRO A 177      -3.978  34.750 -25.771  1.00  0.00           H  
ATOM   2870 1HD  PRO A 177      -5.034  32.312 -25.135  1.00  0.00           H  
ATOM   2871 2HD  PRO A 177      -3.493  32.421 -26.075  1.00  0.00           H  
ATOM   2872  N   ILE A 178      -2.828  34.220 -21.055  1.00  0.00           N  
ATOM   2873  CA  ILE A 178      -3.598  34.676 -19.914  1.00  0.00           C  
ATOM   2874  C   ILE A 178      -4.390  35.779 -20.614  1.00  0.00           C  
ATOM   2875  O   ILE A 178      -3.818  36.597 -21.341  1.00  0.00           O  
ATOM   2876  CB  ILE A 178      -2.708  35.266 -18.796  1.00  0.00           C  
ATOM   2877  CG1 ILE A 178      -2.168  34.130 -17.920  1.00  0.00           C  
ATOM   2878  CG2 ILE A 178      -3.509  36.260 -17.954  1.00  0.00           C  
ATOM   2879  CD1 ILE A 178      -1.370  34.599 -16.715  1.00  0.00           C  
ATOM   2880  H   ILE A 178      -1.858  34.489 -21.133  1.00  0.00           H  
ATOM   2881  HA  ILE A 178      -4.134  33.824 -19.495  1.00  0.00           H  
ATOM   2882  HB  ILE A 178      -1.857  35.780 -19.241  1.00  0.00           H  
ATOM   2883 1HG1 ILE A 178      -2.998  33.523 -17.560  1.00  0.00           H  
ATOM   2884 2HG1 ILE A 178      -1.526  33.484 -18.518  1.00  0.00           H  
ATOM   2885 1HG2 ILE A 178      -2.873  36.669 -17.170  1.00  0.00           H  
ATOM   2886 2HG2 ILE A 178      -3.866  37.069 -18.590  1.00  0.00           H  
ATOM   2887 3HG2 ILE A 178      -4.360  35.751 -17.502  1.00  0.00           H  
ATOM   2888 1HD1 ILE A 178      -1.024  33.735 -16.149  1.00  0.00           H  
ATOM   2889 2HD1 ILE A 178      -0.510  35.180 -17.052  1.00  0.00           H  
ATOM   2890 3HD1 ILE A 178      -2.002  35.219 -16.080  1.00  0.00           H  
ATOM   2891  N   ILE A 179      -5.702  35.778 -20.429  1.00  0.00           N  
ATOM   2892  CA  ILE A 179      -6.563  36.753 -21.093  1.00  0.00           C  
ATOM   2893  C   ILE A 179      -6.827  37.982 -20.226  1.00  0.00           C  
ATOM   2894  O   ILE A 179      -7.572  37.921 -19.248  1.00  0.00           O  
ATOM   2895  CB  ILE A 179      -7.900  36.092 -21.497  1.00  0.00           C  
ATOM   2896  CG1 ILE A 179      -7.619  34.814 -22.299  1.00  0.00           C  
ATOM   2897  CG2 ILE A 179      -8.723  37.055 -22.342  1.00  0.00           C  
ATOM   2898  CD1 ILE A 179      -8.855  33.980 -22.604  1.00  0.00           C  
ATOM   2899  H   ILE A 179      -6.119  35.090 -19.818  1.00  0.00           H  
ATOM   2900  HA  ILE A 179      -6.058  37.105 -21.992  1.00  0.00           H  
ATOM   2901  HB  ILE A 179      -8.463  35.832 -20.601  1.00  0.00           H  
ATOM   2902 1HG1 ILE A 179      -7.147  35.076 -23.246  1.00  0.00           H  
ATOM   2903 2HG1 ILE A 179      -6.918  34.185 -21.749  1.00  0.00           H  
ATOM   2904 1HG2 ILE A 179      -9.663  36.581 -22.624  1.00  0.00           H  
ATOM   2905 2HG2 ILE A 179      -8.930  37.958 -21.769  1.00  0.00           H  
ATOM   2906 3HG2 ILE A 179      -8.165  37.316 -23.242  1.00  0.00           H  
ATOM   2907 1HD1 ILE A 179      -8.567  33.096 -23.173  1.00  0.00           H  
ATOM   2908 2HD1 ILE A 179      -9.327  33.672 -21.670  1.00  0.00           H  
ATOM   2909 3HD1 ILE A 179      -9.559  34.572 -23.187  1.00  0.00           H  
ATOM   2910  N   ASP A 180      -6.201  39.094 -20.605  1.00  0.00           N  
ATOM   2911  CA  ASP A 180      -6.343  40.350 -19.886  1.00  0.00           C  
ATOM   2912  C   ASP A 180      -7.705  40.965 -20.160  1.00  0.00           C  
ATOM   2913  O   ASP A 180      -8.403  40.565 -21.092  1.00  0.00           O  
ATOM   2914  CB  ASP A 180      -5.220  41.309 -20.290  1.00  0.00           C  
ATOM   2915  CG  ASP A 180      -3.866  40.877 -19.744  1.00  0.00           C  
ATOM   2916  OD1 ASP A 180      -2.835  41.235 -20.351  1.00  0.00           O  
ATOM   2917  OD2 ASP A 180      -3.834  40.191 -18.698  1.00  0.00           O1-
ATOM   2918  H   ASP A 180      -5.607  39.058 -21.420  1.00  0.00           H  
ATOM   2919  HA  ASP A 180      -6.268  40.148 -18.818  1.00  0.00           H  
ATOM   2920 1HB  ASP A 180      -5.162  41.364 -21.378  1.00  0.00           H  
ATOM   2921 2HB  ASP A 180      -5.447  42.310 -19.923  1.00  0.00           H  
ATOM   2922  N   VAL A 181      -8.072  41.950 -19.350  1.00  0.00           N  
ATOM   2923  CA  VAL A 181      -9.374  42.586 -19.475  1.00  0.00           C  
ATOM   2924  C   VAL A 181      -9.393  43.963 -20.131  1.00  0.00           C  
ATOM   2925  O   VAL A 181      -8.693  44.878 -19.700  1.00  0.00           O  
ATOM   2926  CB  VAL A 181     -10.000  42.712 -18.073  1.00  0.00           C  
ATOM   2927  CG1 VAL A 181     -11.346  43.418 -18.153  1.00  0.00           C  
ATOM   2928  CG2 VAL A 181     -10.150  41.334 -17.447  1.00  0.00           C  
ATOM   2929  H   VAL A 181      -7.437  42.266 -18.630  1.00  0.00           H  
ATOM   2930  HA  VAL A 181     -10.008  41.957 -20.102  1.00  0.00           H  
ATOM   2931  HB  VAL A 181      -9.351  43.327 -17.450  1.00  0.00           H  
ATOM   2932 1HG1 VAL A 181     -11.774  43.499 -17.153  1.00  0.00           H  
ATOM   2933 2HG1 VAL A 181     -11.209  44.416 -18.571  1.00  0.00           H  
ATOM   2934 3HG1 VAL A 181     -12.020  42.846 -18.790  1.00  0.00           H  
ATOM   2935 1HG2 VAL A 181     -10.593  41.432 -16.456  1.00  0.00           H  
ATOM   2936 2HG2 VAL A 181     -10.794  40.718 -18.074  1.00  0.00           H  
ATOM   2937 3HG2 VAL A 181      -9.170  40.865 -17.362  1.00  0.00           H  
ATOM   2938  N   ILE A 182     -10.198  44.100 -21.181  1.00  0.00           N  
ATOM   2939  CA  ILE A 182     -10.372  45.391 -21.833  1.00  0.00           C  
ATOM   2940  C   ILE A 182     -11.781  45.777 -21.377  1.00  0.00           C  
ATOM   2941  O   ILE A 182     -12.772  45.152 -21.769  1.00  0.00           O  
ATOM   2942  CB  ILE A 182     -10.260  45.323 -23.367  1.00  0.00           C  
ATOM   2943  CG1 ILE A 182      -8.892  44.774 -23.779  1.00  0.00           C  
ATOM   2944  CG2 ILE A 182     -10.492  46.696 -23.979  1.00  0.00           C  
ATOM   2945  CD1 ILE A 182      -8.770  44.482 -25.257  1.00  0.00           C  
ATOM   2946  H   ILE A 182     -10.700  43.298 -21.535  1.00  0.00           H  
ATOM   2947  HA  ILE A 182      -9.573  46.053 -21.507  1.00  0.00           H  
ATOM   2948  HB  ILE A 182     -11.007  44.631 -23.756  1.00  0.00           H  
ATOM   2949 1HG1 ILE A 182      -8.116  45.489 -23.507  1.00  0.00           H  
ATOM   2950 2HG1 ILE A 182      -8.691  43.851 -23.233  1.00  0.00           H  
ATOM   2951 1HG2 ILE A 182     -10.409  46.630 -25.063  1.00  0.00           H  
ATOM   2952 2HG2 ILE A 182     -11.487  47.049 -23.712  1.00  0.00           H  
ATOM   2953 3HG2 ILE A 182      -9.746  47.394 -23.599  1.00  0.00           H  
ATOM   2954 1HD1 ILE A 182      -7.773  44.096 -25.471  1.00  0.00           H  
ATOM   2955 2HD1 ILE A 182      -9.516  43.741 -25.545  1.00  0.00           H  
ATOM   2956 3HD1 ILE A 182      -8.933  45.399 -25.823  1.00  0.00           H  
ATOM   2957  N   ASN A 183     -11.849  46.790 -20.516  1.00  0.00           N  
ATOM   2958  CA  ASN A 183     -13.107  47.259 -19.936  1.00  0.00           C  
ATOM   2959  C   ASN A 183     -14.152  47.622 -20.993  1.00  0.00           C  
ATOM   2960  O   ASN A 183     -13.884  48.402 -21.903  1.00  0.00           O  
ATOM   2961  CB  ASN A 183     -12.850  48.445 -19.025  1.00  0.00           C  
ATOM   2962  CG  ASN A 183     -14.076  48.864 -18.262  1.00  0.00           C  
ATOM   2963  ND2 ASN A 183     -14.201  48.386 -17.050  1.00  0.00           N  
ATOM   2964  OD1 ASN A 183     -14.910  49.622 -18.771  1.00  0.00           O  
ATOM   2965  H   ASN A 183     -10.987  47.248 -20.258  1.00  0.00           H  
ATOM   2966  HA  ASN A 183     -13.547  46.444 -19.359  1.00  0.00           H  
ATOM   2967 1HB  ASN A 183     -12.062  48.194 -18.313  1.00  0.00           H  
ATOM   2968 2HB  ASN A 183     -12.498  49.290 -19.617  1.00  0.00           H  
ATOM   2969 1HD2 ASN A 183     -14.997  48.631 -16.496  1.00  0.00           H  
ATOM   2970 2HD2 ASN A 183     -13.502  47.777 -16.678  1.00  0.00           H  
ATOM   2971  N   MET A 184     -15.351  47.055 -20.861  1.00  0.00           N  
ATOM   2972  CA  MET A 184     -16.415  47.294 -21.839  1.00  0.00           C  
ATOM   2973  C   MET A 184     -17.040  48.684 -21.787  1.00  0.00           C  
ATOM   2974  O   MET A 184     -17.803  49.060 -22.680  1.00  0.00           O  
ATOM   2975  CB  MET A 184     -17.518  46.239 -21.693  1.00  0.00           C  
ATOM   2976  CG  MET A 184     -18.255  46.274 -20.366  1.00  0.00           C  
ATOM   2977  SD  MET A 184     -19.709  45.209 -20.378  1.00  0.00           S  
ATOM   2978  CE  MET A 184     -20.943  46.361 -20.941  1.00  0.00           C  
ATOM   2979  H   MET A 184     -15.533  46.449 -20.074  1.00  0.00           H  
ATOM   2980  HA  MET A 184     -15.989  47.219 -22.839  1.00  0.00           H  
ATOM   2981 1HB  MET A 184     -18.253  46.369 -22.485  1.00  0.00           H  
ATOM   2982 2HB  MET A 184     -17.087  45.243 -21.807  1.00  0.00           H  
ATOM   2983 1HG  MET A 184     -17.586  45.950 -19.569  1.00  0.00           H  
ATOM   2984 2HG  MET A 184     -18.569  47.296 -20.151  1.00  0.00           H  
ATOM   2985 1HE  MET A 184     -21.908  45.859 -21.004  1.00  0.00           H  
ATOM   2986 2HE  MET A 184     -21.011  47.193 -20.239  1.00  0.00           H  
ATOM   2987 3HE  MET A 184     -20.666  46.740 -21.925  1.00  0.00           H  
ATOM   2988  N   ASP A 185     -16.725  49.448 -20.746  1.00  0.00           N  
ATOM   2989  CA  ASP A 185     -17.277  50.790 -20.617  1.00  0.00           C  
ATOM   2990  C   ASP A 185     -16.328  51.891 -21.063  1.00  0.00           C  
ATOM   2991  O   ASP A 185     -16.756  52.849 -21.701  1.00  0.00           O  
ATOM   2992  CB  ASP A 185     -17.717  51.050 -19.177  1.00  0.00           C  
ATOM   2993  CG  ASP A 185     -18.909  50.208 -18.777  1.00  0.00           C  
ATOM   2994  OD1 ASP A 185     -19.875  50.148 -19.569  1.00  0.00           O  
ATOM   2995  OD2 ASP A 185     -18.885  49.614 -17.678  1.00  0.00           O1-
ATOM   2996  H   ASP A 185     -16.099  49.103 -20.033  1.00  0.00           H  
ATOM   2997  HA  ASP A 185     -18.148  50.870 -21.268  1.00  0.00           H  
ATOM   2998 1HB  ASP A 185     -16.890  50.836 -18.500  1.00  0.00           H  
ATOM   2999 2HB  ASP A 185     -17.973  52.103 -19.059  1.00  0.00           H  
ATOM   3000  N   ASN A 186     -15.043  51.767 -20.734  1.00  0.00           N  
ATOM   3001  CA  ASN A 186     -14.090  52.796 -21.125  1.00  0.00           C  
ATOM   3002  C   ASN A 186     -12.820  52.266 -21.777  1.00  0.00           C  
ATOM   3003  O   ASN A 186     -11.843  52.993 -21.954  1.00  0.00           O  
ATOM   3004  CB  ASN A 186     -13.732  53.644 -19.918  1.00  0.00           C  
ATOM   3005  CG  ASN A 186     -13.042  52.853 -18.841  1.00  0.00           C  
ATOM   3006  ND2 ASN A 186     -12.920  53.437 -17.676  1.00  0.00           N  
ATOM   3007  OD1 ASN A 186     -12.618  51.714 -19.065  1.00  0.00           O  
ATOM   3008  H   ASN A 186     -14.717  50.966 -20.213  1.00  0.00           H  
ATOM   3009  HA  ASN A 186     -14.539  53.403 -21.912  1.00  0.00           H  
ATOM   3010 1HB  ASN A 186     -13.080  54.461 -20.227  1.00  0.00           H  
ATOM   3011 2HB  ASN A 186     -14.638  54.086 -19.503  1.00  0.00           H  
ATOM   3012 1HD2 ASN A 186     -12.469  52.958 -16.922  1.00  0.00           H  
ATOM   3013 2HD2 ASN A 186     -13.278  54.360 -17.539  1.00  0.00           H  
ATOM   3014  N   PHE A 187     -12.862  50.988 -22.136  1.00  0.00           N  
ATOM   3015  CA  PHE A 187     -11.778  50.301 -22.824  1.00  0.00           C  
ATOM   3016  C   PHE A 187     -10.357  50.318 -22.277  1.00  0.00           C  
ATOM   3017  O   PHE A 187      -9.399  50.123 -23.032  1.00  0.00           O  
ATOM   3018  CB  PHE A 187     -11.775  50.725 -24.299  1.00  0.00           C  
ATOM   3019  CG  PHE A 187     -12.741  49.943 -25.141  1.00  0.00           C  
ATOM   3020  CD1 PHE A 187     -14.112  50.038 -24.922  1.00  0.00           C  
ATOM   3021  CD2 PHE A 187     -12.274  49.034 -26.085  1.00  0.00           C  
ATOM   3022  CE1 PHE A 187     -15.006  49.228 -25.630  1.00  0.00           C  
ATOM   3023  CE2 PHE A 187     -13.159  48.221 -26.795  1.00  0.00           C  
ATOM   3024  CZ  PHE A 187     -14.525  48.316 -26.566  1.00  0.00           C  
ATOM   3025  H   PHE A 187     -13.704  50.478 -21.910  1.00  0.00           H  
ATOM   3026  HA  PHE A 187     -11.951  49.226 -22.758  1.00  0.00           H  
ATOM   3027 1HB  PHE A 187     -12.028  51.782 -24.374  1.00  0.00           H  
ATOM   3028 2HB  PHE A 187     -10.775  50.598 -24.712  1.00  0.00           H  
ATOM   3029  HD1 PHE A 187     -14.486  50.755 -24.190  1.00  0.00           H  
ATOM   3030  HD2 PHE A 187     -11.201  48.959 -26.264  1.00  0.00           H  
ATOM   3031  HE1 PHE A 187     -16.077  49.311 -25.449  1.00  0.00           H  
ATOM   3032  HE2 PHE A 187     -12.778  47.510 -27.529  1.00  0.00           H  
ATOM   3033  HZ  PHE A 187     -15.218  47.680 -27.114  1.00  0.00           H  
ATOM   3034  N   GLN A 188     -10.204  50.535 -20.976  1.00  0.00           N  
ATOM   3035  CA  GLN A 188      -8.865  50.503 -20.397  1.00  0.00           C  
ATOM   3036  C   GLN A 188      -8.421  49.040 -20.397  1.00  0.00           C  
ATOM   3037  O   GLN A 188      -9.241  48.135 -20.217  1.00  0.00           O  
ATOM   3038  CB  GLN A 188      -8.846  51.086 -18.981  1.00  0.00           C  
ATOM   3039  CG  GLN A 188      -9.213  52.558 -18.909  1.00  0.00           C  
ATOM   3040  CD  GLN A 188      -8.203  53.442 -19.617  1.00  0.00           C  
ATOM   3041  NE2 GLN A 188      -8.676  54.226 -20.578  1.00  0.00           N  
ATOM   3042  OE1 GLN A 188      -7.010  53.419 -19.302  1.00  0.00           O  
ATOM   3043  H   GLN A 188     -10.994  50.722 -20.377  1.00  0.00           H  
ATOM   3044  HA  GLN A 188      -8.214  51.138 -20.998  1.00  0.00           H  
ATOM   3045 1HB  GLN A 188      -9.544  50.534 -18.351  1.00  0.00           H  
ATOM   3046 2HB  GLN A 188      -7.852  50.966 -18.552  1.00  0.00           H  
ATOM   3047 1HG  GLN A 188     -10.185  52.703 -19.382  1.00  0.00           H  
ATOM   3048 2HG  GLN A 188      -9.258  52.860 -17.863  1.00  0.00           H  
ATOM   3049 1HE2 GLN A 188      -8.056  54.832 -21.081  1.00  0.00           H  
ATOM   3050 2HE2 GLN A 188      -9.651  54.213 -20.801  1.00  0.00           H  
ATOM   3051  N   TYR A 189      -7.126  48.817 -20.608  1.00  0.00           N  
ATOM   3052  CA  TYR A 189      -6.542  47.477 -20.643  1.00  0.00           C  
ATOM   3053  C   TYR A 189      -5.957  47.184 -19.260  1.00  0.00           C  
ATOM   3054  O   TYR A 189      -5.094  47.921 -18.781  1.00  0.00           O  
ATOM   3055  CB  TYR A 189      -5.450  47.445 -21.725  1.00  0.00           C  
ATOM   3056  CG  TYR A 189      -4.815  46.094 -21.975  1.00  0.00           C  
ATOM   3057  CD1 TYR A 189      -3.984  45.504 -21.024  1.00  0.00           C  
ATOM   3058  CD2 TYR A 189      -5.031  45.414 -23.176  1.00  0.00           C  
ATOM   3059  CE1 TYR A 189      -3.385  44.272 -21.258  1.00  0.00           C  
ATOM   3060  CE2 TYR A 189      -4.434  44.183 -23.423  1.00  0.00           C  
ATOM   3061  CZ  TYR A 189      -3.613  43.618 -22.458  1.00  0.00           C  
ATOM   3062  OH  TYR A 189      -3.027  42.396 -22.687  1.00  0.00           O  
ATOM   3063  H   TYR A 189      -6.527  49.618 -20.749  1.00  0.00           H  
ATOM   3064  HA  TYR A 189      -7.327  46.765 -20.896  1.00  0.00           H  
ATOM   3065 1HB  TYR A 189      -5.868  47.786 -22.674  1.00  0.00           H  
ATOM   3066 2HB  TYR A 189      -4.649  48.132 -21.454  1.00  0.00           H  
ATOM   3067  HD1 TYR A 189      -3.795  46.009 -20.077  1.00  0.00           H  
ATOM   3068  HD2 TYR A 189      -5.677  45.847 -23.939  1.00  0.00           H  
ATOM   3069  HE1 TYR A 189      -2.739  43.828 -20.501  1.00  0.00           H  
ATOM   3070  HE2 TYR A 189      -4.613  43.671 -24.369  1.00  0.00           H  
ATOM   3071  HH  TYR A 189      -3.288  42.075 -23.554  1.00  0.00           H  
ATOM   3072  N   VAL A 190      -6.430  46.114 -18.620  1.00  0.00           N  
ATOM   3073  CA  VAL A 190      -5.968  45.744 -17.284  1.00  0.00           C  
ATOM   3074  C   VAL A 190      -5.557  44.278 -17.189  1.00  0.00           C  
ATOM   3075  O   VAL A 190      -6.244  43.391 -17.686  1.00  0.00           O  
ATOM   3076  CB  VAL A 190      -7.076  46.021 -16.251  1.00  0.00           C  
ATOM   3077  CG1 VAL A 190      -6.627  45.594 -14.861  1.00  0.00           C  
ATOM   3078  CG2 VAL A 190      -7.443  47.497 -16.269  1.00  0.00           C  
ATOM   3079  H   VAL A 190      -7.128  45.543 -19.075  1.00  0.00           H  
ATOM   3080  HA  VAL A 190      -5.091  46.345 -17.042  1.00  0.00           H  
ATOM   3081  HB  VAL A 190      -7.953  45.425 -16.503  1.00  0.00           H  
ATOM   3082 1HG1 VAL A 190      -7.422  45.797 -14.143  1.00  0.00           H  
ATOM   3083 2HG1 VAL A 190      -6.404  44.527 -14.863  1.00  0.00           H  
ATOM   3084 3HG1 VAL A 190      -5.734  46.152 -14.580  1.00  0.00           H  
ATOM   3085 1HG2 VAL A 190      -8.227  47.686 -15.536  1.00  0.00           H  
ATOM   3086 2HG2 VAL A 190      -6.564  48.094 -16.022  1.00  0.00           H  
ATOM   3087 3HG2 VAL A 190      -7.801  47.771 -17.262  1.00  0.00           H  
ATOM   3088  N   GLY A 191      -4.434  44.034 -16.528  1.00  0.00           N  
ATOM   3089  CA  GLY A 191      -3.942  42.680 -16.393  1.00  0.00           C  
ATOM   3090  C   GLY A 191      -4.787  41.760 -15.539  1.00  0.00           C  
ATOM   3091  O   GLY A 191      -5.261  42.132 -14.464  1.00  0.00           O  
ATOM   3092  H   GLY A 191      -3.910  44.792 -16.113  1.00  0.00           H  
ATOM   3093 1HA  GLY A 191      -3.858  42.223 -17.379  1.00  0.00           H  
ATOM   3094 2HA  GLY A 191      -2.942  42.698 -15.962  1.00  0.00           H  
ATOM   3095  N   ALA A 192      -4.972  40.542 -16.029  1.00  0.00           N  
ATOM   3096  CA  ALA A 192      -5.735  39.536 -15.313  1.00  0.00           C  
ATOM   3097  C   ALA A 192      -4.834  38.918 -14.256  1.00  0.00           C  
ATOM   3098  O   ALA A 192      -3.607  38.981 -14.356  1.00  0.00           O  
ATOM   3099  CB  ALA A 192      -6.207  38.450 -16.280  1.00  0.00           C  
ATOM   3100  H   ALA A 192      -4.571  40.308 -16.926  1.00  0.00           H  
ATOM   3101  HA  ALA A 192      -6.606  40.022 -14.872  1.00  0.00           H  
ATOM   3102 1HB  ALA A 192      -6.779  37.700 -15.733  1.00  0.00           H  
ATOM   3103 2HB  ALA A 192      -6.837  38.897 -17.050  1.00  0.00           H  
ATOM   3104 3HB  ALA A 192      -5.345  37.978 -16.747  1.00  0.00           H  
ATOM   3105  N   SER A 193      -5.451  38.326 -13.240  1.00  0.00           N  
ATOM   3106  CA  SER A 193      -4.704  37.650 -12.191  1.00  0.00           C  
ATOM   3107  C   SER A 193      -4.048  36.428 -12.839  1.00  0.00           C  
ATOM   3108  O   SER A 193      -4.616  35.823 -13.749  1.00  0.00           O  
ATOM   3109  CB  SER A 193      -5.610  37.241 -11.046  1.00  0.00           C  
ATOM   3110  OG  SER A 193      -4.902  36.514 -10.080  1.00  0.00           O  
ATOM   3111  H   SER A 193      -6.460  38.345 -13.193  1.00  0.00           H  
ATOM   3112  HA  SER A 193      -3.987  38.354 -11.768  1.00  0.00           H  
ATOM   3113 1HB  SER A 193      -6.045  38.130 -10.590  1.00  0.00           H  
ATOM   3114 2HB  SER A 193      -6.431  36.637 -11.431  1.00  0.00           H  
ATOM   3115  HG  SER A 193      -4.702  37.135  -9.376  1.00  0.00           H  
ATOM   3116  N   ALA A 194      -2.857  36.069 -12.378  1.00  0.00           N  
ATOM   3117  CA  ALA A 194      -2.155  34.917 -12.932  1.00  0.00           C  
ATOM   3118  C   ALA A 194      -2.392  33.659 -12.097  1.00  0.00           C  
ATOM   3119  O   ALA A 194      -1.839  32.599 -12.389  1.00  0.00           O  
ATOM   3120  CB  ALA A 194      -0.660  35.213 -13.027  1.00  0.00           C  
ATOM   3121  H   ALA A 194      -2.425  36.597 -11.634  1.00  0.00           H  
ATOM   3122  HA  ALA A 194      -2.546  34.732 -13.933  1.00  0.00           H  
ATOM   3123 1HB  ALA A 194      -0.143  34.347 -13.441  1.00  0.00           H  
ATOM   3124 2HB  ALA A 194      -0.501  36.075 -13.673  1.00  0.00           H  
ATOM   3125 3HB  ALA A 194      -0.268  35.428 -12.034  1.00  0.00           H  
ATOM   3126  N   ASP A 195      -3.218  33.780 -11.060  1.00  0.00           N  
ATOM   3127  CA  ASP A 195      -3.531  32.656 -10.171  1.00  0.00           C  
ATOM   3128  C   ASP A 195      -4.819  31.951 -10.581  1.00  0.00           C  
ATOM   3129  O   ASP A 195      -5.484  31.346  -9.739  1.00  0.00           O  
ATOM   3130  CB  ASP A 195      -3.652  33.137  -8.723  1.00  0.00           C  
ATOM   3131  CG  ASP A 195      -2.326  33.607  -8.142  1.00  0.00           C  
ATOM   3132  OD1 ASP A 195      -1.336  32.950  -8.362  1.00  0.00           O  
ATOM   3133  OD2 ASP A 195      -2.315  34.620  -7.483  1.00  0.00           O1-
ATOM   3134  H   ASP A 195      -3.642  34.680 -10.883  1.00  0.00           H  
ATOM   3135  HA  ASP A 195      -2.697  31.954 -10.196  1.00  0.00           H  
ATOM   3136 1HB  ASP A 195      -4.367  33.958  -8.671  1.00  0.00           H  
ATOM   3137 2HB  ASP A 195      -4.038  32.328  -8.102  1.00  0.00           H  
ATOM   3138  N   LEU A 196      -5.167  32.010 -11.862  1.00  0.00           N  
ATOM   3139  CA  LEU A 196      -6.411  31.401 -12.320  1.00  0.00           C  
ATOM   3140  C   LEU A 196      -6.263  30.218 -13.264  1.00  0.00           C  
ATOM   3141  O   LEU A 196      -5.236  30.040 -13.915  1.00  0.00           O  
ATOM   3142  CB  LEU A 196      -7.264  32.472 -13.012  1.00  0.00           C  
ATOM   3143  CG  LEU A 196      -7.602  33.704 -12.163  1.00  0.00           C  
ATOM   3144  CD1 LEU A 196      -8.356  34.716 -13.014  1.00  0.00           C  
ATOM   3145  CD2 LEU A 196      -8.429  33.279 -10.958  1.00  0.00           C  
ATOM   3146  H   LEU A 196      -4.573  32.478 -12.531  1.00  0.00           H  
ATOM   3147  HA  LEU A 196      -6.950  31.020 -11.454  1.00  0.00           H  
ATOM   3148 1HB  LEU A 196      -6.736  32.816 -13.899  1.00  0.00           H  
ATOM   3149 2HB  LEU A 196      -8.203  32.018 -13.327  1.00  0.00           H  
ATOM   3150  HG  LEU A 196      -6.680  34.175 -11.822  1.00  0.00           H  
ATOM   3151 1HD1 LEU A 196      -8.597  35.592 -12.411  1.00  0.00           H  
ATOM   3152 2HD1 LEU A 196      -7.734  35.017 -13.857  1.00  0.00           H  
ATOM   3153 3HD1 LEU A 196      -9.277  34.267 -13.384  1.00  0.00           H  
ATOM   3154 1HD2 LEU A 196      -8.669  34.154 -10.354  1.00  0.00           H  
ATOM   3155 2HD2 LEU A 196      -9.352  32.808 -11.298  1.00  0.00           H  
ATOM   3156 3HD2 LEU A 196      -7.860  32.569 -10.358  1.00  0.00           H  
ATOM   3157  N   LYS A 197      -7.325  29.421 -13.324  1.00  0.00           N  
ATOM   3158  CA  LYS A 197      -7.413  28.264 -14.208  1.00  0.00           C  
ATOM   3159  C   LYS A 197      -8.828  28.285 -14.779  1.00  0.00           C  
ATOM   3160  O   LYS A 197      -9.708  28.958 -14.238  1.00  0.00           O  
ATOM   3161  CB  LYS A 197      -7.219  26.954 -13.442  1.00  0.00           C  
ATOM   3162  CG  LYS A 197      -5.839  26.744 -12.865  1.00  0.00           C  
ATOM   3163  CD  LYS A 197      -5.606  25.271 -12.579  1.00  0.00           C  
ATOM   3164  CE  LYS A 197      -4.154  25.006 -12.197  1.00  0.00           C  
ATOM   3165  NZ  LYS A 197      -3.767  23.581 -12.410  1.00  0.00           N1+
ATOM   3166  H   LYS A 197      -8.106  29.640 -12.721  1.00  0.00           H  
ATOM   3167  HA  LYS A 197      -6.623  28.340 -14.956  1.00  0.00           H  
ATOM   3168 1HB  LYS A 197      -7.929  26.908 -12.615  1.00  0.00           H  
ATOM   3169 2HB  LYS A 197      -7.429  26.112 -14.101  1.00  0.00           H  
ATOM   3170 1HG  LYS A 197      -5.090  27.101 -13.573  1.00  0.00           H  
ATOM   3171 2HG  LYS A 197      -5.740  27.315 -11.941  1.00  0.00           H  
ATOM   3172 1HD  LYS A 197      -6.254  24.952 -11.762  1.00  0.00           H  
ATOM   3173 2HD  LYS A 197      -5.853  24.686 -13.465  1.00  0.00           H  
ATOM   3174 1HE  LYS A 197      -3.500  25.639 -12.794  1.00  0.00           H  
ATOM   3175 2HE  LYS A 197      -4.004  25.254 -11.146  1.00  0.00           H  
ATOM   3176 1HZ  LYS A 197      -2.801  23.451 -12.144  1.00  0.00           H  
ATOM   3177 2HZ  LYS A 197      -4.354  22.984 -11.846  1.00  0.00           H  
ATOM   3178 3HZ  LYS A 197      -3.882  23.343 -13.385  1.00  0.00           H  
ATOM   3179  N   GLY A 198      -9.047  27.561 -15.870  1.00  0.00           N  
ATOM   3180  CA  GLY A 198     -10.378  27.502 -16.448  1.00  0.00           C  
ATOM   3181  C   GLY A 198     -11.152  26.384 -15.772  1.00  0.00           C  
ATOM   3182  O   GLY A 198     -10.590  25.329 -15.485  1.00  0.00           O  
ATOM   3183  H   GLY A 198      -8.296  27.046 -16.309  1.00  0.00           H  
ATOM   3184 1HA  GLY A 198     -10.877  28.462 -16.312  1.00  0.00           H  
ATOM   3185 2HA  GLY A 198     -10.301  27.330 -17.522  1.00  0.00           H  
ATOM   3186  N   GLY A 199     -12.437  26.603 -15.513  1.00  0.00           N  
ATOM   3187  CA  GLY A 199     -13.227  25.574 -14.861  1.00  0.00           C  
ATOM   3188  C   GLY A 199     -14.705  25.690 -15.153  1.00  0.00           C  
ATOM   3189  O   GLY A 199     -15.113  26.423 -16.054  1.00  0.00           O  
ATOM   3190  H   GLY A 199     -12.878  27.476 -15.762  1.00  0.00           H  
ATOM   3191 1HA  GLY A 199     -12.882  24.591 -15.184  1.00  0.00           H  
ATOM   3192 2HA  GLY A 199     -13.076  25.629 -13.783  1.00  0.00           H  
ATOM   3193  N   PHE A 200     -15.511  24.967 -14.382  1.00  0.00           N  
ATOM   3194  CA  PHE A 200     -16.958  24.989 -14.562  1.00  0.00           C  
ATOM   3195  C   PHE A 200     -17.665  24.240 -13.437  1.00  0.00           C  
ATOM   3196  O   PHE A 200     -17.079  23.369 -12.791  1.00  0.00           O  
ATOM   3197  CB  PHE A 200     -17.324  24.344 -15.905  1.00  0.00           C  
ATOM   3198  CG  PHE A 200     -16.870  22.909 -16.032  1.00  0.00           C  
ATOM   3199  CD1 PHE A 200     -17.674  21.863 -15.584  1.00  0.00           C  
ATOM   3200  CD2 PHE A 200     -15.620  22.608 -16.566  1.00  0.00           C  
ATOM   3201  CE1 PHE A 200     -17.236  20.540 -15.666  1.00  0.00           C  
ATOM   3202  CE2 PHE A 200     -15.174  21.292 -16.650  1.00  0.00           C  
ATOM   3203  CZ  PHE A 200     -15.986  20.254 -16.199  1.00  0.00           C  
ATOM   3204  H   PHE A 200     -15.115  24.389 -13.655  1.00  0.00           H  
ATOM   3205  HA  PHE A 200     -17.291  26.028 -14.565  1.00  0.00           H  
ATOM   3206 1HB  PHE A 200     -18.404  24.373 -16.040  1.00  0.00           H  
ATOM   3207 2HB  PHE A 200     -16.877  24.915 -16.717  1.00  0.00           H  
ATOM   3208  HD1 PHE A 200     -18.656  22.088 -15.166  1.00  0.00           H  
ATOM   3209  HD2 PHE A 200     -14.985  23.422 -16.918  1.00  0.00           H  
ATOM   3210  HE1 PHE A 200     -17.876  19.732 -15.312  1.00  0.00           H  
ATOM   3211  HE2 PHE A 200     -14.191  21.073 -17.068  1.00  0.00           H  
ATOM   3212  HZ  PHE A 200     -15.642  19.222 -16.262  1.00  0.00           H  
ATOM   3213  N   ASP A 201     -18.919  24.607 -13.200  1.00  0.00           N  
ATOM   3214  CA  ASP A 201     -19.748  23.939 -12.206  1.00  0.00           C  
ATOM   3215  C   ASP A 201     -20.569  22.962 -13.037  1.00  0.00           C  
ATOM   3216  O   ASP A 201     -20.427  22.922 -14.260  1.00  0.00           O  
ATOM   3217  CB  ASP A 201     -20.640  24.911 -11.430  1.00  0.00           C  
ATOM   3218  CG  ASP A 201     -19.849  25.859 -10.539  1.00  0.00           C  
ATOM   3219  OD1 ASP A 201     -18.926  25.411  -9.900  1.00  0.00           O  
ATOM   3220  OD2 ASP A 201     -20.175  27.021 -10.505  1.00  0.00           O1-
ATOM   3221  H   ASP A 201     -19.308  25.374 -13.729  1.00  0.00           H  
ATOM   3222  HA  ASP A 201     -19.095  23.477 -11.466  1.00  0.00           H  
ATOM   3223 1HB  ASP A 201     -21.229  25.503 -12.132  1.00  0.00           H  
ATOM   3224 2HB  ASP A 201     -21.338  24.349 -10.809  1.00  0.00           H  
ATOM   3225  N   TRP A 202     -21.433  22.180 -12.401  1.00  0.00           N  
ATOM   3226  CA  TRP A 202     -22.216  21.207 -13.147  1.00  0.00           C  
ATOM   3227  C   TRP A 202     -23.235  21.759 -14.137  1.00  0.00           C  
ATOM   3228  O   TRP A 202     -23.822  20.998 -14.906  1.00  0.00           O  
ATOM   3229  CB  TRP A 202     -22.877  20.220 -12.185  1.00  0.00           C  
ATOM   3230  CG  TRP A 202     -21.863  19.371 -11.494  1.00  0.00           C  
ATOM   3231  CD1 TRP A 202     -21.424  19.494 -10.204  1.00  0.00           C  
ATOM   3232  CD2 TRP A 202     -21.089  18.318 -12.081  1.00  0.00           C  
ATOM   3233  CE2 TRP A 202     -20.198  17.851 -11.089  1.00  0.00           C  
ATOM   3234  CE3 TRP A 202     -21.063  17.724 -13.352  1.00  0.00           C  
ATOM   3235  NE1 TRP A 202     -20.424  18.585  -9.955  1.00  0.00           N  
ATOM   3236  CZ2 TRP A 202     -19.284  16.819 -11.329  1.00  0.00           C  
ATOM   3237  CZ3 TRP A 202     -20.156  16.697 -13.590  1.00  0.00           C  
ATOM   3238  CH2 TRP A 202     -19.279  16.255 -12.582  1.00  0.00           C  
ATOM   3239  H   TRP A 202     -21.557  22.249 -11.401  1.00  0.00           H  
ATOM   3240  HA  TRP A 202     -21.546  20.659 -13.810  1.00  0.00           H  
ATOM   3241 1HB  TRP A 202     -23.458  20.768 -11.443  1.00  0.00           H  
ATOM   3242 2HB  TRP A 202     -23.569  19.582 -12.735  1.00  0.00           H  
ATOM   3243  HD1 TRP A 202     -21.811  20.209  -9.481  1.00  0.00           H  
ATOM   3244  HE1 TRP A 202     -19.934  18.475  -9.079  1.00  0.00           H  
ATOM   3245  HE3 TRP A 202     -21.742  18.059 -14.136  1.00  0.00           H  
ATOM   3246  HZ2 TRP A 202     -18.597  16.463 -10.560  1.00  0.00           H  
ATOM   3247  HZ3 TRP A 202     -20.140  16.243 -14.580  1.00  0.00           H  
ATOM   3248  HH2 TRP A 202     -18.580  15.447 -12.802  1.00  0.00           H  
ATOM   3249  N   ASN A 203     -23.447  23.072 -14.135  1.00  0.00           N  
ATOM   3250  CA  ASN A 203     -24.373  23.665 -15.094  1.00  0.00           C  
ATOM   3251  C   ASN A 203     -23.612  23.852 -16.405  1.00  0.00           C  
ATOM   3252  O   ASN A 203     -24.135  24.387 -17.379  1.00  0.00           O  
ATOM   3253  CB  ASN A 203     -24.943  24.977 -14.586  1.00  0.00           C  
ATOM   3254  CG  ASN A 203     -23.895  26.046 -14.446  1.00  0.00           C  
ATOM   3255  ND2 ASN A 203     -24.253  27.134 -13.812  1.00  0.00           N  
ATOM   3256  OD1 ASN A 203     -22.761  25.886 -14.912  1.00  0.00           O  
ATOM   3257  H   ASN A 203     -22.974  23.671 -13.474  1.00  0.00           H  
ATOM   3258  HA  ASN A 203     -25.212  22.981 -15.234  1.00  0.00           H  
ATOM   3259 1HB  ASN A 203     -25.715  25.328 -15.272  1.00  0.00           H  
ATOM   3260 2HB  ASN A 203     -25.414  24.817 -13.616  1.00  0.00           H  
ATOM   3261 1HD2 ASN A 203     -23.597  27.879 -13.689  1.00  0.00           H  
ATOM   3262 2HD2 ASN A 203     -25.181  27.220 -13.451  1.00  0.00           H  
ATOM   3263  N   LEU A 204     -22.362  23.393 -16.397  1.00  0.00           N  
ATOM   3264  CA  LEU A 204     -21.470  23.435 -17.551  1.00  0.00           C  
ATOM   3265  C   LEU A 204     -21.206  24.806 -18.172  1.00  0.00           C  
ATOM   3266  O   LEU A 204     -21.029  24.934 -19.387  1.00  0.00           O  
ATOM   3267  CB  LEU A 204     -21.968  22.451 -18.616  1.00  0.00           C  
ATOM   3268  CG  LEU A 204     -22.095  21.008 -18.100  1.00  0.00           C  
ATOM   3269  CD1 LEU A 204     -22.471  20.075 -19.244  1.00  0.00           C  
ATOM   3270  CD2 LEU A 204     -20.779  20.563 -17.464  1.00  0.00           C  
ATOM   3271  H   LEU A 204     -22.029  22.995 -15.531  1.00  0.00           H  
ATOM   3272  HA  LEU A 204     -20.473  23.138 -17.228  1.00  0.00           H  
ATOM   3273 1HB  LEU A 204     -22.941  22.786 -18.971  1.00  0.00           H  
ATOM   3274 2HB  LEU A 204     -21.272  22.466 -19.456  1.00  0.00           H  
ATOM   3275  HG  LEU A 204     -22.888  20.957 -17.353  1.00  0.00           H  
ATOM   3276 1HD1 LEU A 204     -22.559  19.055 -18.868  1.00  0.00           H  
ATOM   3277 2HD1 LEU A 204     -23.425  20.387 -19.670  1.00  0.00           H  
ATOM   3278 3HD1 LEU A 204     -21.700  20.112 -20.013  1.00  0.00           H  
ATOM   3279 1HD2 LEU A 204     -20.879  19.540 -17.101  1.00  0.00           H  
ATOM   3280 2HD2 LEU A 204     -19.983  20.608 -18.207  1.00  0.00           H  
ATOM   3281 3HD2 LEU A 204     -20.536  21.222 -16.631  1.00  0.00           H  
ATOM   3282  N   VAL A 205     -21.160  25.826 -17.322  1.00  0.00           N  
ATOM   3283  CA  VAL A 205     -20.885  27.188 -17.760  1.00  0.00           C  
ATOM   3284  C   VAL A 205     -19.440  27.507 -17.393  1.00  0.00           C  
ATOM   3285  O   VAL A 205     -19.051  27.394 -16.230  1.00  0.00           O  
ATOM   3286  CB  VAL A 205     -21.840  28.194 -17.090  1.00  0.00           C  
ATOM   3287  CG1 VAL A 205     -21.508  29.614 -17.524  1.00  0.00           C  
ATOM   3288  CG2 VAL A 205     -23.281  27.847 -17.435  1.00  0.00           C  
ATOM   3289  H   VAL A 205     -21.321  25.647 -16.342  1.00  0.00           H  
ATOM   3290  HA  VAL A 205     -21.064  27.249 -18.834  1.00  0.00           H  
ATOM   3291  HB  VAL A 205     -21.703  28.147 -16.010  1.00  0.00           H  
ATOM   3292 1HG1 VAL A 205     -22.193  30.311 -17.041  1.00  0.00           H  
ATOM   3293 2HG1 VAL A 205     -20.485  29.852 -17.236  1.00  0.00           H  
ATOM   3294 3HG1 VAL A 205     -21.610  29.697 -18.606  1.00  0.00           H  
ATOM   3295 1HG2 VAL A 205     -23.951  28.562 -16.957  1.00  0.00           H  
ATOM   3296 2HG2 VAL A 205     -23.417  27.888 -18.516  1.00  0.00           H  
ATOM   3297 3HG2 VAL A 205     -23.509  26.843 -17.078  1.00  0.00           H  
ATOM   3298  N   PHE A 206     -18.643  27.896 -18.383  1.00  0.00           N  
ATOM   3299  CA  PHE A 206     -17.246  28.229 -18.124  1.00  0.00           C  
ATOM   3300  C   PHE A 206     -17.100  29.337 -17.085  1.00  0.00           C  
ATOM   3301  O   PHE A 206     -17.829  30.328 -17.108  1.00  0.00           O  
ATOM   3302  CB  PHE A 206     -16.550  28.686 -19.407  1.00  0.00           C  
ATOM   3303  CG  PHE A 206     -15.150  29.183 -19.184  1.00  0.00           C  
ATOM   3304  CD1 PHE A 206     -14.089  28.293 -19.074  1.00  0.00           C  
ATOM   3305  CD2 PHE A 206     -14.900  30.543 -19.023  1.00  0.00           C  
ATOM   3306  CE1 PHE A 206     -12.798  28.748 -18.803  1.00  0.00           C  
ATOM   3307  CE2 PHE A 206     -13.613  31.007 -18.750  1.00  0.00           C  
ATOM   3308  CZ  PHE A 206     -12.562  30.106 -18.640  1.00  0.00           C  
ATOM   3309  H   PHE A 206     -18.999  27.964 -19.325  1.00  0.00           H  
ATOM   3310  HA  PHE A 206     -16.740  27.335 -17.756  1.00  0.00           H  
ATOM   3311 1HB  PHE A 206     -16.511  27.858 -20.115  1.00  0.00           H  
ATOM   3312 2HB  PHE A 206     -17.128  29.484 -19.870  1.00  0.00           H  
ATOM   3313  HD1 PHE A 206     -14.274  27.226 -19.201  1.00  0.00           H  
ATOM   3314  HD2 PHE A 206     -15.726  31.249 -19.109  1.00  0.00           H  
ATOM   3315  HE1 PHE A 206     -11.976  28.036 -18.721  1.00  0.00           H  
ATOM   3316  HE2 PHE A 206     -13.432  32.073 -18.623  1.00  0.00           H  
ATOM   3317  HZ  PHE A 206     -11.556  30.463 -18.424  1.00  0.00           H  
ATOM   3318  N   LYS A 207     -16.143  29.160 -16.180  1.00  0.00           N  
ATOM   3319  CA  LYS A 207     -15.863  30.146 -15.147  1.00  0.00           C  
ATOM   3320  C   LYS A 207     -14.377  30.077 -14.833  1.00  0.00           C  
ATOM   3321  O   LYS A 207     -13.699  29.129 -15.225  1.00  0.00           O  
ATOM   3322  CB  LYS A 207     -16.700  29.896 -13.892  1.00  0.00           C  
ATOM   3323  CG  LYS A 207     -16.378  28.595 -13.169  1.00  0.00           C  
ATOM   3324  CD  LYS A 207     -17.332  28.356 -12.008  1.00  0.00           C  
ATOM   3325  CE  LYS A 207     -17.078  29.336 -10.872  1.00  0.00           C  
ATOM   3326  NZ  LYS A 207     -17.910  29.029  -9.677  1.00  0.00           N1+
ATOM   3327  H   LYS A 207     -15.595  28.313 -16.212  1.00  0.00           H  
ATOM   3328  HA  LYS A 207     -16.147  31.130 -15.522  1.00  0.00           H  
ATOM   3329 1HB  LYS A 207     -16.554  30.715 -13.187  1.00  0.00           H  
ATOM   3330 2HB  LYS A 207     -17.758  29.878 -14.157  1.00  0.00           H  
ATOM   3331 1HG  LYS A 207     -16.455  27.761 -13.868  1.00  0.00           H  
ATOM   3332 2HG  LYS A 207     -15.358  28.634 -12.788  1.00  0.00           H  
ATOM   3333 1HD  LYS A 207     -18.361  28.471 -12.353  1.00  0.00           H  
ATOM   3334 2HD  LYS A 207     -17.204  27.340 -11.635  1.00  0.00           H  
ATOM   3335 1HE  LYS A 207     -16.027  29.298 -10.587  1.00  0.00           H  
ATOM   3336 2HE  LYS A 207     -17.305  30.348 -11.206  1.00  0.00           H  
ATOM   3337 1HZ  LYS A 207     -17.712  29.700  -8.947  1.00  0.00           H  
ATOM   3338 2HZ  LYS A 207     -18.888  29.080  -9.925  1.00  0.00           H  
ATOM   3339 3HZ  LYS A 207     -17.695  28.100  -9.345  1.00  0.00           H  
ATOM   3340  N   TRP A 208     -13.869  31.091 -14.145  1.00  0.00           N  
ATOM   3341  CA  TRP A 208     -12.465  31.105 -13.763  1.00  0.00           C  
ATOM   3342  C   TRP A 208     -12.349  30.578 -12.341  1.00  0.00           C  
ATOM   3343  O   TRP A 208     -13.086  31.013 -11.450  1.00  0.00           O  
ATOM   3344  CB  TRP A 208     -11.881  32.516 -13.856  1.00  0.00           C  
ATOM   3345  CG  TRP A 208     -11.748  33.017 -15.262  1.00  0.00           C  
ATOM   3346  CD1 TRP A 208     -12.608  33.846 -15.917  1.00  0.00           C  
ATOM   3347  CD2 TRP A 208     -10.684  32.722 -16.199  1.00  0.00           C  
ATOM   3348  CE2 TRP A 208     -10.980  33.407 -17.381  1.00  0.00           C  
ATOM   3349  CE3 TRP A 208      -9.521  31.944 -16.131  1.00  0.00           C  
ATOM   3350  NE1 TRP A 208     -12.157  34.086 -17.191  1.00  0.00           N  
ATOM   3351  CZ2 TRP A 208     -10.156  33.340 -18.494  1.00  0.00           C  
ATOM   3352  CZ3 TRP A 208      -8.695  31.878 -17.246  1.00  0.00           C  
ATOM   3353  CH2 TRP A 208      -9.005  32.558 -18.397  1.00  0.00           C  
ATOM   3354  H   TRP A 208     -14.459  31.865 -13.878  1.00  0.00           H  
ATOM   3355  HA  TRP A 208     -11.910  30.478 -14.461  1.00  0.00           H  
ATOM   3356 1HB  TRP A 208     -12.516  33.209 -13.304  1.00  0.00           H  
ATOM   3357 2HB  TRP A 208     -10.896  32.533 -13.391  1.00  0.00           H  
ATOM   3358  HD1 TRP A 208     -13.522  34.258 -15.491  1.00  0.00           H  
ATOM   3359  HE1 TRP A 208     -12.616  34.666 -17.879  1.00  0.00           H  
ATOM   3360  HE3 TRP A 208      -9.269  31.404 -15.219  1.00  0.00           H  
ATOM   3361  HZ2 TRP A 208     -10.385  33.873 -19.417  1.00  0.00           H  
ATOM   3362  HZ3 TRP A 208      -7.792  31.269 -17.185  1.00  0.00           H  
ATOM   3363  HH2 TRP A 208      -8.334  32.484 -19.253  1.00  0.00           H  
ATOM   3364  N   ASP A 209     -11.440  29.626 -12.140  1.00  0.00           N  
ATOM   3365  CA  ASP A 209     -11.198  29.034 -10.825  1.00  0.00           C  
ATOM   3366  C   ASP A 209      -9.908  29.605 -10.258  1.00  0.00           C  
ATOM   3367  O   ASP A 209      -8.929  29.768 -10.983  1.00  0.00           O  
ATOM   3368  CB  ASP A 209     -11.107  27.508 -10.911  1.00  0.00           C  
ATOM   3369  CG  ASP A 209     -12.435  26.855 -11.272  1.00  0.00           C  
ATOM   3370  OD1 ASP A 209     -13.444  27.513 -11.174  1.00  0.00           O  
ATOM   3371  OD2 ASP A 209     -12.427  25.706 -11.641  1.00  0.00           O1-
ATOM   3372  H   ASP A 209     -10.900  29.306 -12.931  1.00  0.00           H  
ATOM   3373  HA  ASP A 209     -12.058  29.242 -10.189  1.00  0.00           H  
ATOM   3374 1HB  ASP A 209     -10.366  27.230 -11.660  1.00  0.00           H  
ATOM   3375 2HB  ASP A 209     -10.771  27.109  -9.954  1.00  0.00           H  
ATOM   3376  N   TYR A 210      -9.902  29.901  -8.965  1.00  0.00           N  
ATOM   3377  CA  TYR A 210      -8.699  30.425  -8.332  1.00  0.00           C  
ATOM   3378  C   TYR A 210      -7.900  29.222  -7.846  1.00  0.00           C  
ATOM   3379  O   TYR A 210      -8.469  28.264  -7.320  1.00  0.00           O  
ATOM   3380  CB  TYR A 210      -9.033  31.377  -7.181  1.00  0.00           C  
ATOM   3381  CG  TYR A 210      -9.656  32.680  -7.628  1.00  0.00           C  
ATOM   3382  CD1 TYR A 210     -10.929  32.687  -8.179  1.00  0.00           C  
ATOM   3383  CD2 TYR A 210      -8.955  33.869  -7.487  1.00  0.00           C  
ATOM   3384  CE1 TYR A 210     -11.498  33.878  -8.588  1.00  0.00           C  
ATOM   3385  CE2 TYR A 210      -9.524  35.059  -7.895  1.00  0.00           C  
ATOM   3386  CZ  TYR A 210     -10.791  35.066  -8.443  1.00  0.00           C  
ATOM   3387  OH  TYR A 210     -11.358  36.252  -8.850  1.00  0.00           O  
ATOM   3388  H   TYR A 210     -10.733  29.767  -8.406  1.00  0.00           H  
ATOM   3389  HA  TYR A 210      -8.156  31.025  -9.064  1.00  0.00           H  
ATOM   3390 1HB  TYR A 210      -9.724  30.887  -6.493  1.00  0.00           H  
ATOM   3391 2HB  TYR A 210      -8.125  31.607  -6.624  1.00  0.00           H  
ATOM   3392  HD1 TYR A 210     -11.480  31.753  -8.290  1.00  0.00           H  
ATOM   3393  HD2 TYR A 210      -7.955  33.863  -7.054  1.00  0.00           H  
ATOM   3394  HE1 TYR A 210     -12.498  33.884  -9.021  1.00  0.00           H  
ATOM   3395  HE2 TYR A 210      -8.973  35.993  -7.785  1.00  0.00           H  
ATOM   3396  HH  TYR A 210     -12.233  36.082  -9.205  1.00  0.00           H  
ATOM   3397  N   MET A 211      -6.586  29.266  -8.029  1.00  0.00           N  
ATOM   3398  CA  MET A 211      -5.732  28.165  -7.607  1.00  0.00           C  
ATOM   3399  C   MET A 211      -5.833  27.898  -6.116  1.00  0.00           C  
ATOM   3400  O   MET A 211      -6.045  28.814  -5.326  1.00  0.00           O  
ATOM   3401  CB  MET A 211      -4.282  28.453  -7.992  1.00  0.00           C  
ATOM   3402  CG  MET A 211      -4.008  28.411  -9.489  1.00  0.00           C  
ATOM   3403  SD  MET A 211      -2.275  28.713  -9.887  1.00  0.00           S  
ATOM   3404  CE  MET A 211      -2.314  28.606 -11.674  1.00  0.00           C  
ATOM   3405  H   MET A 211      -6.167  30.073  -8.468  1.00  0.00           H  
ATOM   3406  HA  MET A 211      -6.038  27.266  -8.140  1.00  0.00           H  
ATOM   3407 1HB  MET A 211      -3.998  29.439  -7.628  1.00  0.00           H  
ATOM   3408 2HB  MET A 211      -3.627  27.725  -7.512  1.00  0.00           H  
ATOM   3409 1HG  MET A 211      -4.287  27.434  -9.882  1.00  0.00           H  
ATOM   3410 2HG  MET A 211      -4.613  29.165  -9.991  1.00  0.00           H  
ATOM   3411 1HE  MET A 211      -1.313  28.772 -12.071  1.00  0.00           H  
ATOM   3412 2HE  MET A 211      -2.665  27.617 -11.972  1.00  0.00           H  
ATOM   3413 3HE  MET A 211      -2.991  29.365 -12.069  1.00  0.00           H  
ATOM   3414  N   THR A 212      -5.681  26.631  -5.744  1.00  0.00           N  
ATOM   3415  CA  THR A 212      -5.720  26.239  -4.345  1.00  0.00           C  
ATOM   3416  C   THR A 212      -4.408  26.704  -3.726  1.00  0.00           C  
ATOM   3417  O   THR A 212      -3.459  27.036  -4.437  1.00  0.00           O  
ATOM   3418  CB  THR A 212      -5.902  24.720  -4.168  1.00  0.00           C  
ATOM   3419  CG2 THR A 212      -7.181  24.254  -4.846  1.00  0.00           C  
ATOM   3420  OG1 THR A 212      -4.783  24.032  -4.741  1.00  0.00           O  
ATOM   3421  H   THR A 212      -5.536  25.923  -6.449  1.00  0.00           H  
ATOM   3422  HA  THR A 212      -6.613  26.669  -3.889  1.00  0.00           H  
ATOM   3423  HB  THR A 212      -5.953  24.481  -3.106  1.00  0.00           H  
ATOM   3424  HG1 THR A 212      -5.091  23.245  -5.196  1.00  0.00           H  
ATOM   3425 1HG2 THR A 212      -7.293  23.178  -4.710  1.00  0.00           H  
ATOM   3426 2HG2 THR A 212      -8.035  24.767  -4.404  1.00  0.00           H  
ATOM   3427 3HG2 THR A 212      -7.132  24.481  -5.910  1.00  0.00           H  
ATOM   3428  N   PRO A 213      -4.336  26.736  -2.389  1.00  0.00           N  
ATOM   3429  CA  PRO A 213      -3.108  27.167  -1.715  1.00  0.00           C  
ATOM   3430  C   PRO A 213      -1.881  26.375  -2.171  1.00  0.00           C  
ATOM   3431  O   PRO A 213      -0.824  26.946  -2.440  1.00  0.00           O  
ATOM   3432  CB  PRO A 213      -3.425  26.905  -0.239  1.00  0.00           C  
ATOM   3433  CG  PRO A 213      -4.903  27.072  -0.147  1.00  0.00           C  
ATOM   3434  CD  PRO A 213      -5.430  26.450  -1.414  1.00  0.00           C  
ATOM   3435  HA  PRO A 213      -2.942  28.236  -1.914  1.00  0.00           H  
ATOM   3436 1HB  PRO A 213      -3.091  25.896   0.044  1.00  0.00           H  
ATOM   3437 2HB  PRO A 213      -2.875  27.615   0.396  1.00  0.00           H  
ATOM   3438 1HG  PRO A 213      -5.286  26.577   0.758  1.00  0.00           H  
ATOM   3439 2HG  PRO A 213      -5.161  28.138  -0.060  1.00  0.00           H  
ATOM   3440 1HD  PRO A 213      -5.562  25.368  -1.264  1.00  0.00           H  
ATOM   3441 2HD  PRO A 213      -6.384  26.924  -1.686  1.00  0.00           H  
ATOM   3442  N   GLU A 214      -2.032  25.058  -2.269  1.00  0.00           N  
ATOM   3443  CA  GLU A 214      -0.927  24.198  -2.677  1.00  0.00           C  
ATOM   3444  C   GLU A 214      -0.495  24.433  -4.119  1.00  0.00           C  
ATOM   3445  O   GLU A 214       0.693  24.375  -4.431  1.00  0.00           O  
ATOM   3446  CB  GLU A 214      -1.315  22.729  -2.495  1.00  0.00           C  
ATOM   3447  CG  GLU A 214      -1.457  22.292  -1.044  1.00  0.00           C  
ATOM   3448  CD  GLU A 214      -1.889  20.858  -0.904  1.00  0.00           C  
ATOM   3449  OE1 GLU A 214      -2.162  20.240  -1.905  1.00  0.00           O  
ATOM   3450  OE2 GLU A 214      -1.945  20.381   0.204  1.00  0.00           O1-
ATOM   3451  H   GLU A 214      -2.928  24.643  -2.058  1.00  0.00           H  
ATOM   3452  HA  GLU A 214      -0.063  24.425  -2.053  1.00  0.00           H  
ATOM   3453 1HB  GLU A 214      -2.264  22.538  -2.996  1.00  0.00           H  
ATOM   3454 2HB  GLU A 214      -0.563  22.093  -2.963  1.00  0.00           H  
ATOM   3455 1HG  GLU A 214      -0.500  22.421  -0.540  1.00  0.00           H  
ATOM   3456 2HG  GLU A 214      -2.186  22.935  -0.551  1.00  0.00           H  
ATOM   3457  N   GLN A 215      -1.453  24.695  -5.001  1.00  0.00           N  
ATOM   3458  CA  GLN A 215      -1.120  24.938  -6.399  1.00  0.00           C  
ATOM   3459  C   GLN A 215      -0.292  26.215  -6.525  1.00  0.00           C  
ATOM   3460  O   GLN A 215       0.681  26.259  -7.278  1.00  0.00           O  
ATOM   3461  CB  GLN A 215      -2.389  25.041  -7.249  1.00  0.00           C  
ATOM   3462  CG  GLN A 215      -3.139  23.730  -7.409  1.00  0.00           C  
ATOM   3463  CD  GLN A 215      -4.431  23.894  -8.187  1.00  0.00           C  
ATOM   3464  NE2 GLN A 215      -4.848  22.834  -8.871  1.00  0.00           N  
ATOM   3465  OE1 GLN A 215      -5.049  24.962  -8.174  1.00  0.00           O  
ATOM   3466  H   GLN A 215      -2.421  24.730  -4.713  1.00  0.00           H  
ATOM   3467  HA  GLN A 215      -0.520  24.102  -6.761  1.00  0.00           H  
ATOM   3468 1HB  GLN A 215      -3.069  25.765  -6.801  1.00  0.00           H  
ATOM   3469 2HB  GLN A 215      -2.132  25.405  -8.244  1.00  0.00           H  
ATOM   3470 1HG  GLN A 215      -2.505  23.024  -7.945  1.00  0.00           H  
ATOM   3471 2HG  GLN A 215      -3.382  23.339  -6.421  1.00  0.00           H  
ATOM   3472 1HE2 GLN A 215      -5.694  22.882  -9.403  1.00  0.00           H  
ATOM   3473 2HE2 GLN A 215      -4.316  21.987  -8.854  1.00  0.00           H  
ATOM   3474  N   ARG A 216      -0.663  27.246  -5.773  1.00  0.00           N  
ATOM   3475  CA  ARG A 216       0.076  28.504  -5.815  1.00  0.00           C  
ATOM   3476  C   ARG A 216       1.470  28.308  -5.215  1.00  0.00           C  
ATOM   3477  O   ARG A 216       2.464  28.814  -5.738  1.00  0.00           O  
ATOM   3478  CB  ARG A 216      -0.664  29.594  -5.053  1.00  0.00           C  
ATOM   3479  CG  ARG A 216      -1.948  30.074  -5.709  1.00  0.00           C  
ATOM   3480  CD  ARG A 216      -2.643  31.090  -4.878  1.00  0.00           C  
ATOM   3481  NE  ARG A 216      -1.875  32.320  -4.771  1.00  0.00           N  
ATOM   3482  CZ  ARG A 216      -2.124  33.302  -3.882  1.00  0.00           C  
ATOM   3483  NH1 ARG A 216      -3.120  33.184  -3.033  1.00  0.00           N1+
ATOM   3484  NH2 ARG A 216      -1.365  34.384  -3.865  1.00  0.00           N  
ATOM   3485  H   ARG A 216      -1.465  27.166  -5.164  1.00  0.00           H  
ATOM   3486  HA  ARG A 216       0.170  28.816  -6.856  1.00  0.00           H  
ATOM   3487 1HB  ARG A 216      -0.917  29.233  -4.057  1.00  0.00           H  
ATOM   3488 2HB  ARG A 216      -0.012  30.459  -4.932  1.00  0.00           H  
ATOM   3489 1HG  ARG A 216      -1.718  30.521  -6.676  1.00  0.00           H  
ATOM   3490 2HG  ARG A 216      -2.622  29.228  -5.851  1.00  0.00           H  
ATOM   3491 1HD  ARG A 216      -3.608  31.328  -5.326  1.00  0.00           H  
ATOM   3492 2HD  ARG A 216      -2.797  30.695  -3.874  1.00  0.00           H  
ATOM   3493  HE  ARG A 216      -1.100  32.447  -5.408  1.00  0.00           H  
ATOM   3494 1HH1 ARG A 216      -3.700  32.356  -3.047  1.00  0.00           H  
ATOM   3495 2HH1 ARG A 216      -3.306  33.919  -2.368  1.00  0.00           H  
ATOM   3496 1HH2 ARG A 216      -0.599  34.475  -4.518  1.00  0.00           H  
ATOM   3497 2HH2 ARG A 216      -1.551  35.120  -3.199  1.00  0.00           H  
ATOM   3498  N   ARG A 217       1.540  27.564  -4.115  1.00  0.00           N  
ATOM   3499  CA  ARG A 217       2.815  27.304  -3.457  1.00  0.00           C  
ATOM   3500  C   ARG A 217       3.785  26.576  -4.386  1.00  0.00           C  
ATOM   3501  O   ARG A 217       4.967  26.916  -4.455  1.00  0.00           O  
ATOM   3502  CB  ARG A 217       2.605  26.476  -2.198  1.00  0.00           C  
ATOM   3503  CG  ARG A 217       3.869  26.185  -1.405  1.00  0.00           C  
ATOM   3504  CD  ARG A 217       3.584  25.372  -0.196  1.00  0.00           C  
ATOM   3505  NE  ARG A 217       3.102  24.043  -0.537  1.00  0.00           N  
ATOM   3506  CZ  ARG A 217       2.491  23.206   0.324  1.00  0.00           C  
ATOM   3507  NH1 ARG A 217       2.295  23.574   1.571  1.00  0.00           N1+
ATOM   3508  NH2 ARG A 217       2.088  22.016  -0.084  1.00  0.00           N  
ATOM   3509  H   ARG A 217       0.695  27.170  -3.726  1.00  0.00           H  
ATOM   3510  HA  ARG A 217       3.253  28.258  -3.160  1.00  0.00           H  
ATOM   3511 1HB  ARG A 217       1.911  26.991  -1.535  1.00  0.00           H  
ATOM   3512 2HB  ARG A 217       2.155  25.519  -2.463  1.00  0.00           H  
ATOM   3513 1HG  ARG A 217       4.573  25.635  -2.030  1.00  0.00           H  
ATOM   3514 2HG  ARG A 217       4.324  27.124  -1.088  1.00  0.00           H  
ATOM   3515 1HD  ARG A 217       4.495  25.263   0.393  1.00  0.00           H  
ATOM   3516 2HD  ARG A 217       2.822  25.867   0.405  1.00  0.00           H  
ATOM   3517  HE  ARG A 217       3.235  23.724  -1.487  1.00  0.00           H  
ATOM   3518 1HH1 ARG A 217       2.602  24.485   1.883  1.00  0.00           H  
ATOM   3519 2HH1 ARG A 217       1.836  22.948   2.216  1.00  0.00           H  
ATOM   3520 1HH2 ARG A 217       2.239  21.733  -1.043  1.00  0.00           H  
ATOM   3521 2HH2 ARG A 217       1.630  21.389   0.561  1.00  0.00           H  
ATOM   3522  N   SER A 218       3.277  25.577  -5.100  1.00  0.00           N  
ATOM   3523  CA  SER A 218       4.097  24.779  -6.006  1.00  0.00           C  
ATOM   3524  C   SER A 218       4.672  25.554  -7.186  1.00  0.00           C  
ATOM   3525  O   SER A 218       5.621  25.097  -7.827  1.00  0.00           O  
ATOM   3526  CB  SER A 218       3.275  23.618  -6.531  1.00  0.00           C  
ATOM   3527  OG  SER A 218       2.240  24.070  -7.360  1.00  0.00           O  
ATOM   3528  H   SER A 218       2.293  25.365  -5.011  1.00  0.00           H  
ATOM   3529  HA  SER A 218       4.955  24.398  -5.452  1.00  0.00           H  
ATOM   3530 1HB  SER A 218       3.920  22.938  -7.087  1.00  0.00           H  
ATOM   3531 2HB  SER A 218       2.857  23.061  -5.694  1.00  0.00           H  
ATOM   3532  HG  SER A 218       1.896  24.864  -6.945  1.00  0.00           H  
ATOM   3533  N   ARG A 219       4.104  26.721  -7.474  1.00  0.00           N  
ATOM   3534  CA  ARG A 219       4.562  27.528  -8.604  1.00  0.00           C  
ATOM   3535  C   ARG A 219       5.562  28.618  -8.235  1.00  0.00           C  
ATOM   3536  O   ARG A 219       5.962  29.415  -9.086  1.00  0.00           O  
ATOM   3537  CB  ARG A 219       3.356  28.155  -9.311  1.00  0.00           C  
ATOM   3538  CG  ARG A 219       2.409  27.135  -9.934  1.00  0.00           C  
ATOM   3539  CD  ARG A 219       1.198  27.814 -10.562  1.00  0.00           C  
ATOM   3540  NE  ARG A 219       1.588  28.811 -11.556  1.00  0.00           N  
ATOM   3541  CZ  ARG A 219       2.095  28.525 -12.752  1.00  0.00           C  
ATOM   3542  NH1 ARG A 219       2.425  29.505 -13.581  1.00  0.00           N1+
ATOM   3543  NH2 ARG A 219       2.263  27.259 -13.125  1.00  0.00           N  
ATOM   3544  H   ARG A 219       3.342  27.060  -6.904  1.00  0.00           H  
ATOM   3545  HA  ARG A 219       5.086  26.876  -9.304  1.00  0.00           H  
ATOM   3546 1HB  ARG A 219       2.790  28.755  -8.601  1.00  0.00           H  
ATOM   3547 2HB  ARG A 219       3.704  28.822 -10.099  1.00  0.00           H  
ATOM   3548 1HG  ARG A 219       2.934  26.577 -10.710  1.00  0.00           H  
ATOM   3549 2HG  ARG A 219       2.059  26.446  -9.165  1.00  0.00           H  
ATOM   3550 1HD  ARG A 219       0.577  27.067 -11.055  1.00  0.00           H  
ATOM   3551 2HD  ARG A 219       0.619  28.315  -9.787  1.00  0.00           H  
ATOM   3552  HE  ARG A 219       1.463  29.787 -11.319  1.00  0.00           H  
ATOM   3553 1HH1 ARG A 219       2.291  30.468 -13.303  1.00  0.00           H  
ATOM   3554 2HH1 ARG A 219       2.810  29.290 -14.489  1.00  0.00           H  
ATOM   3555 1HH2 ARG A 219       2.005  26.509 -12.499  1.00  0.00           H  
ATOM   3556 2HH2 ARG A 219       2.648  27.048 -14.034  1.00  0.00           H  
ATOM   3557  N   GLN A 220       5.978  28.643  -6.974  1.00  0.00           N  
ATOM   3558  CA  GLN A 220       6.926  29.648  -6.506  1.00  0.00           C  
ATOM   3559  C   GLN A 220       8.312  29.520  -7.136  1.00  0.00           C  
ATOM   3560  O   GLN A 220       9.016  30.519  -7.303  1.00  0.00           O  
ATOM   3561  CB  GLN A 220       7.050  29.572  -4.982  1.00  0.00           C  
ATOM   3562  CG  GLN A 220       5.807  30.020  -4.233  1.00  0.00           C  
ATOM   3563  CD  GLN A 220       5.959  29.890  -2.729  1.00  0.00           C  
ATOM   3564  NE2 GLN A 220       5.254  30.737  -1.989  1.00  0.00           N  
ATOM   3565  OE1 GLN A 220       6.703  29.036  -2.238  1.00  0.00           O  
ATOM   3566  H   GLN A 220       5.632  27.953  -6.322  1.00  0.00           H  
ATOM   3567  HA  GLN A 220       6.565  30.631  -6.807  1.00  0.00           H  
ATOM   3568 1HB  GLN A 220       7.271  28.547  -4.686  1.00  0.00           H  
ATOM   3569 2HB  GLN A 220       7.883  30.194  -4.653  1.00  0.00           H  
ATOM   3570 1HG  GLN A 220       5.611  31.066  -4.467  1.00  0.00           H  
ATOM   3571 2HG  GLN A 220       4.965  29.402  -4.546  1.00  0.00           H  
ATOM   3572 1HE2 GLN A 220       5.313  30.699  -0.990  1.00  0.00           H  
ATOM   3573 2HE2 GLN A 220       4.663  31.414  -2.428  1.00  0.00           H  
ATOM   3574  N   GLY A 221       8.699  28.300  -7.496  1.00  0.00           N  
ATOM   3575  CA  GLY A 221       9.994  28.095  -8.119  1.00  0.00           C  
ATOM   3576  C   GLY A 221      10.044  28.772  -9.476  1.00  0.00           C  
ATOM   3577  O   GLY A 221      11.062  29.351  -9.858  1.00  0.00           O  
ATOM   3578  H   GLY A 221       8.096  27.504  -7.340  1.00  0.00           H  
ATOM   3579 1HA  GLY A 221      10.776  28.495  -7.472  1.00  0.00           H  
ATOM   3580 2HA  GLY A 221      10.182  27.028  -8.226  1.00  0.00           H  
ATOM   3581  N   ASN A 222       8.937  28.687 -10.208  1.00  0.00           N  
ATOM   3582  CA  ASN A 222       8.829  29.299 -11.524  1.00  0.00           C  
ATOM   3583  C   ASN A 222       7.401  29.791 -11.730  1.00  0.00           C  
ATOM   3584  O   ASN A 222       6.545  29.060 -12.224  1.00  0.00           O  
ATOM   3585  CB  ASN A 222       9.237  28.327 -12.615  1.00  0.00           C  
ATOM   3586  CG  ASN A 222       9.245  28.959 -13.979  1.00  0.00           C  
ATOM   3587  ND2 ASN A 222      10.042  28.422 -14.867  1.00  0.00           N  
ATOM   3588  OD1 ASN A 222       8.528  29.935 -14.229  1.00  0.00           O  
ATOM   3589  H   ASN A 222       8.148  28.179  -9.835  1.00  0.00           H  
ATOM   3590  HA  ASN A 222       9.523  30.139 -11.573  1.00  0.00           H  
ATOM   3591 1HB  ASN A 222      10.234  27.937 -12.402  1.00  0.00           H  
ATOM   3592 2HB  ASN A 222       8.550  27.480 -12.624  1.00  0.00           H  
ATOM   3593 1HD2 ASN A 222      10.089  28.801 -15.792  1.00  0.00           H  
ATOM   3594 2HD2 ASN A 222      10.604  27.633 -14.622  1.00  0.00           H  
ATOM   3595  N   PRO A 223       7.127  31.045 -11.347  1.00  0.00           N  
ATOM   3596  CA  PRO A 223       5.793  31.639 -11.486  1.00  0.00           C  
ATOM   3597  C   PRO A 223       5.260  31.760 -12.912  1.00  0.00           C  
ATOM   3598  O   PRO A 223       4.064  31.978 -13.107  1.00  0.00           O  
ATOM   3599  CB  PRO A 223       6.001  33.025 -10.868  1.00  0.00           C  
ATOM   3600  CG  PRO A 223       7.176  32.851  -9.967  1.00  0.00           C  
ATOM   3601  CD  PRO A 223       8.059  31.869 -10.689  1.00  0.00           C  
ATOM   3602  HA  PRO A 223       5.061  31.012 -10.957  1.00  0.00           H  
ATOM   3603 1HB  PRO A 223       6.178  33.768 -11.660  1.00  0.00           H  
ATOM   3604 2HB  PRO A 223       5.094  33.338 -10.331  1.00  0.00           H  
ATOM   3605 1HG  PRO A 223       7.670  33.819  -9.795  1.00  0.00           H  
ATOM   3606 2HG  PRO A 223       6.849  32.483  -8.983  1.00  0.00           H  
ATOM   3607 1HD  PRO A 223       8.708  32.409 -11.395  1.00  0.00           H  
ATOM   3608 2HD  PRO A 223       8.662  31.310  -9.958  1.00  0.00           H  
ATOM   3609  N   VAL A 224       6.131  31.621 -13.907  1.00  0.00           N  
ATOM   3610  CA  VAL A 224       5.676  31.727 -15.289  1.00  0.00           C  
ATOM   3611  C   VAL A 224       5.619  30.381 -16.003  1.00  0.00           C  
ATOM   3612  O   VAL A 224       5.560  30.321 -17.232  1.00  0.00           O  
ATOM   3613  CB  VAL A 224       6.559  32.700 -16.105  1.00  0.00           C  
ATOM   3614  CG1 VAL A 224       6.351  34.120 -15.610  1.00  0.00           C  
ATOM   3615  CG2 VAL A 224       8.022  32.309 -15.987  1.00  0.00           C  
ATOM   3616  H   VAL A 224       7.108  31.442 -13.724  1.00  0.00           H  
ATOM   3617  HA  VAL A 224       4.656  32.113 -15.289  1.00  0.00           H  
ATOM   3618  HB  VAL A 224       6.258  32.658 -17.153  1.00  0.00           H  
ATOM   3619 1HG1 VAL A 224       6.975  34.801 -16.187  1.00  0.00           H  
ATOM   3620 2HG1 VAL A 224       5.304  34.397 -15.729  1.00  0.00           H  
ATOM   3621 3HG1 VAL A 224       6.627  34.181 -14.557  1.00  0.00           H  
ATOM   3622 1HG2 VAL A 224       8.631  33.002 -16.566  1.00  0.00           H  
ATOM   3623 2HG2 VAL A 224       8.325  32.347 -14.941  1.00  0.00           H  
ATOM   3624 3HG2 VAL A 224       8.161  31.297 -16.368  1.00  0.00           H  
ATOM   3625  N   ALA A 225       5.638  29.300 -15.230  1.00  0.00           N  
ATOM   3626  CA  ALA A 225       5.565  27.966 -15.808  1.00  0.00           C  
ATOM   3627  C   ALA A 225       4.166  27.780 -16.390  1.00  0.00           C  
ATOM   3628  O   ALA A 225       3.219  28.459 -15.984  1.00  0.00           O  
ATOM   3629  CB  ALA A 225       5.826  26.911 -14.733  1.00  0.00           C  
ATOM   3630  H   ALA A 225       5.702  29.399 -14.227  1.00  0.00           H  
ATOM   3631  HA  ALA A 225       6.335  27.887 -16.576  1.00  0.00           H  
ATOM   3632 1HB  ALA A 225       5.769  25.917 -15.177  1.00  0.00           H  
ATOM   3633 2HB  ALA A 225       6.818  27.062 -14.308  1.00  0.00           H  
ATOM   3634 3HB  ALA A 225       5.077  27.000 -13.947  1.00  0.00           H  
ATOM   3635  N   PRO A 226       4.017  26.869 -17.364  1.00  0.00           N  
ATOM   3636  CA  PRO A 226       2.689  26.655 -17.943  1.00  0.00           C  
ATOM   3637  C   PRO A 226       1.715  26.226 -16.849  1.00  0.00           C  
ATOM   3638  O   PRO A 226       2.109  25.608 -15.855  1.00  0.00           O  
ATOM   3639  CB  PRO A 226       2.923  25.541 -18.969  1.00  0.00           C  
ATOM   3640  CG  PRO A 226       4.372  25.648 -19.300  1.00  0.00           C  
ATOM   3641  CD  PRO A 226       5.024  26.051 -18.005  1.00  0.00           C  
ATOM   3642  HA  PRO A 226       2.361  27.576 -18.447  1.00  0.00           H  
ATOM   3643 1HB  PRO A 226       2.655  24.567 -18.534  1.00  0.00           H  
ATOM   3644 2HB  PRO A 226       2.272  25.693 -19.843  1.00  0.00           H  
ATOM   3645 1HG  PRO A 226       4.746  24.686 -19.681  1.00  0.00           H  
ATOM   3646 2HG  PRO A 226       4.525  26.387 -20.100  1.00  0.00           H  
ATOM   3647 1HD  PRO A 226       5.259  25.152 -17.417  1.00  0.00           H  
ATOM   3648 2HD  PRO A 226       5.937  26.627 -18.217  1.00  0.00           H  
ATOM   3649  N   ILE A 227       0.442  26.555 -17.038  1.00  0.00           N  
ATOM   3650  CA  ILE A 227      -0.590  26.194 -16.079  1.00  0.00           C  
ATOM   3651  C   ILE A 227      -1.523  25.197 -16.748  1.00  0.00           C  
ATOM   3652  O   ILE A 227      -2.224  25.548 -17.692  1.00  0.00           O  
ATOM   3653  CB  ILE A 227      -1.438  27.421 -15.659  1.00  0.00           C  
ATOM   3654  CG1 ILE A 227      -0.545  28.481 -15.015  1.00  0.00           C  
ATOM   3655  CG2 ILE A 227      -2.530  26.996 -14.679  1.00  0.00           C  
ATOM   3656  CD1 ILE A 227      -1.239  29.825 -14.834  1.00  0.00           C  
ATOM   3657  H   ILE A 227       0.185  27.070 -17.868  1.00  0.00           H  
ATOM   3658  HA  ILE A 227      -0.109  25.794 -15.188  1.00  0.00           H  
ATOM   3659  HB  ILE A 227      -1.902  27.861 -16.541  1.00  0.00           H  
ATOM   3660 1HG1 ILE A 227      -0.211  28.131 -14.039  1.00  0.00           H  
ATOM   3661 2HG1 ILE A 227       0.343  28.631 -15.630  1.00  0.00           H  
ATOM   3662 1HG2 ILE A 227      -3.119  27.866 -14.390  1.00  0.00           H  
ATOM   3663 2HG2 ILE A 227      -3.178  26.261 -15.153  1.00  0.00           H  
ATOM   3664 3HG2 ILE A 227      -2.073  26.557 -13.791  1.00  0.00           H  
ATOM   3665 1HD1 ILE A 227      -0.548  30.532 -14.371  1.00  0.00           H  
ATOM   3666 2HD1 ILE A 227      -1.551  30.208 -15.806  1.00  0.00           H  
ATOM   3667 3HD1 ILE A 227      -2.111  29.701 -14.194  1.00  0.00           H  
ATOM   3668  N   LYS A 228      -1.523  23.952 -16.283  1.00  0.00           N  
ATOM   3669  CA  LYS A 228      -2.435  22.977 -16.862  1.00  0.00           C  
ATOM   3670  C   LYS A 228      -3.831  23.411 -16.436  1.00  0.00           C  
ATOM   3671  O   LYS A 228      -4.060  23.730 -15.270  1.00  0.00           O  
ATOM   3672  CB  LYS A 228      -2.124  21.554 -16.396  1.00  0.00           C  
ATOM   3673  CG  LYS A 228      -0.802  20.998 -16.910  1.00  0.00           C  
ATOM   3674  CD  LYS A 228      -0.580  19.571 -16.433  1.00  0.00           C  
ATOM   3675  CE  LYS A 228       0.744  19.018 -16.938  1.00  0.00           C  
ATOM   3676  NZ  LYS A 228       0.987  17.630 -16.459  1.00  0.00           N1+
ATOM   3677  H   LYS A 228      -0.906  23.665 -15.536  1.00  0.00           H  
ATOM   3678  HA  LYS A 228      -2.316  22.992 -17.947  1.00  0.00           H  
ATOM   3679 1HB  LYS A 228      -2.097  21.526 -15.306  1.00  0.00           H  
ATOM   3680 2HB  LYS A 228      -2.918  20.882 -16.721  1.00  0.00           H  
ATOM   3681 1HG  LYS A 228      -0.801  21.012 -18.001  1.00  0.00           H  
ATOM   3682 2HG  LYS A 228       0.017  21.623 -16.556  1.00  0.00           H  
ATOM   3683 1HD  LYS A 228      -0.582  19.547 -15.342  1.00  0.00           H  
ATOM   3684 2HD  LYS A 228      -1.389  18.936 -16.794  1.00  0.00           H  
ATOM   3685 1HE  LYS A 228       0.747  19.019 -18.027  1.00  0.00           H  
ATOM   3686 2HE  LYS A 228       1.560  19.654 -16.594  1.00  0.00           H  
ATOM   3687 1HZ  LYS A 228       1.873  17.300 -16.815  1.00  0.00           H  
ATOM   3688 2HZ  LYS A 228       1.005  17.620 -15.449  1.00  0.00           H  
ATOM   3689 3HZ  LYS A 228       0.247  17.026 -16.788  1.00  0.00           H  
ATOM   3690  N   THR A 229      -4.761  23.435 -17.384  1.00  0.00           N  
ATOM   3691  CA  THR A 229      -6.114  23.879 -17.088  1.00  0.00           C  
ATOM   3692  C   THR A 229      -7.168  22.876 -17.551  1.00  0.00           C  
ATOM   3693  O   THR A 229      -7.064  22.309 -18.637  1.00  0.00           O  
ATOM   3694  CB  THR A 229      -6.390  25.249 -17.735  1.00  0.00           C  
ATOM   3695  CG2 THR A 229      -6.253  25.162 -19.247  1.00  0.00           C  
ATOM   3696  OG1 THR A 229      -7.717  25.678 -17.403  1.00  0.00           O  
ATOM   3697  H   THR A 229      -4.533  23.144 -18.323  1.00  0.00           H  
ATOM   3698  HA  THR A 229      -6.215  23.986 -16.008  1.00  0.00           H  
ATOM   3699  HB  THR A 229      -5.680  25.982 -17.353  1.00  0.00           H  
ATOM   3700  HG1 THR A 229      -8.226  25.795 -18.209  1.00  0.00           H  
ATOM   3701 1HG2 THR A 229      -6.451  26.139 -19.688  1.00  0.00           H  
ATOM   3702 2HG2 THR A 229      -5.242  24.846 -19.503  1.00  0.00           H  
ATOM   3703 3HG2 THR A 229      -6.968  24.437 -19.636  1.00  0.00           H  
ATOM   3704  N   PRO A 230      -8.191  22.633 -16.718  1.00  0.00           N  
ATOM   3705  CA  PRO A 230      -9.258  21.690 -17.067  1.00  0.00           C  
ATOM   3706  C   PRO A 230     -10.020  22.138 -18.305  1.00  0.00           C  
ATOM   3707  O   PRO A 230     -10.540  21.320 -19.061  1.00  0.00           O  
ATOM   3708  CB  PRO A 230     -10.152  21.705 -15.823  1.00  0.00           C  
ATOM   3709  CG  PRO A 230      -9.242  22.133 -14.723  1.00  0.00           C  
ATOM   3710  CD  PRO A 230      -8.307  23.118 -15.373  1.00  0.00           C  
ATOM   3711  HA  PRO A 230      -8.822  20.691 -17.221  1.00  0.00           H  
ATOM   3712 1HB  PRO A 230     -10.994  22.396 -15.974  1.00  0.00           H  
ATOM   3713 2HB  PRO A 230     -10.583  20.706 -15.657  1.00  0.00           H  
ATOM   3714 1HG  PRO A 230      -9.822  22.578 -13.902  1.00  0.00           H  
ATOM   3715 2HG  PRO A 230      -8.716  21.262 -14.305  1.00  0.00           H  
ATOM   3716 1HD  PRO A 230      -8.750  24.124 -15.336  1.00  0.00           H  
ATOM   3717 2HD  PRO A 230      -7.338  23.101 -14.853  1.00  0.00           H  
ATOM   3718  N   MET A 231     -10.076  23.448 -18.509  1.00  0.00           N  
ATOM   3719  CA  MET A 231     -10.815  23.998 -19.635  1.00  0.00           C  
ATOM   3720  C   MET A 231     -10.193  25.313 -20.103  1.00  0.00           C  
ATOM   3721  O   MET A 231      -9.614  26.053 -19.306  1.00  0.00           O  
ATOM   3722  CB  MET A 231     -12.256  24.239 -19.179  1.00  0.00           C  
ATOM   3723  CG  MET A 231     -13.280  24.453 -20.261  1.00  0.00           C  
ATOM   3724  SD  MET A 231     -14.912  24.656 -19.505  1.00  0.00           S  
ATOM   3725  CE  MET A 231     -15.811  25.431 -20.867  1.00  0.00           C  
ATOM   3726  H   MET A 231      -9.601  24.077 -17.878  1.00  0.00           H  
ATOM   3727  HA  MET A 231     -10.800  23.271 -20.447  1.00  0.00           H  
ATOM   3728 1HB  MET A 231     -12.598  23.390 -18.590  1.00  0.00           H  
ATOM   3729 2HB  MET A 231     -12.293  25.119 -18.535  1.00  0.00           H  
ATOM   3730 1HG  MET A 231     -13.019  25.340 -20.837  1.00  0.00           H  
ATOM   3731 2HG  MET A 231     -13.282  23.598 -20.936  1.00  0.00           H  
ATOM   3732 1HE  MET A 231     -16.840  25.622 -20.561  1.00  0.00           H  
ATOM   3733 2HE  MET A 231     -15.330  26.372 -21.133  1.00  0.00           H  
ATOM   3734 3HE  MET A 231     -15.807  24.766 -21.731  1.00  0.00           H  
ATOM   3735  N   ILE A 232     -10.281  25.595 -21.400  1.00  0.00           N  
ATOM   3736  CA  ILE A 232      -9.773  26.865 -21.905  1.00  0.00           C  
ATOM   3737  C   ILE A 232     -10.991  27.766 -22.066  1.00  0.00           C  
ATOM   3738  O   ILE A 232     -12.103  27.274 -22.265  1.00  0.00           O  
ATOM   3739  CB  ILE A 232      -9.068  26.739 -23.283  1.00  0.00           C  
ATOM   3740  CG1 ILE A 232      -9.991  26.054 -24.292  1.00  0.00           C  
ATOM   3741  CG2 ILE A 232      -7.740  26.013 -23.123  1.00  0.00           C  
ATOM   3742  CD1 ILE A 232      -9.470  26.103 -25.726  1.00  0.00           C  
ATOM   3743  H   ILE A 232     -10.697  24.939 -22.047  1.00  0.00           H  
ATOM   3744  HA  ILE A 232      -9.038  27.247 -21.198  1.00  0.00           H  
ATOM   3745  HB  ILE A 232      -8.887  27.732 -23.690  1.00  0.00           H  
ATOM   3746 1HG1 ILE A 232     -10.126  25.010 -24.011  1.00  0.00           H  
ATOM   3747 2HG1 ILE A 232     -10.973  26.528 -24.268  1.00  0.00           H  
ATOM   3748 1HG2 ILE A 232      -7.253  25.929 -24.095  1.00  0.00           H  
ATOM   3749 2HG2 ILE A 232      -7.099  26.572 -22.443  1.00  0.00           H  
ATOM   3750 3HG2 ILE A 232      -7.917  25.017 -22.718  1.00  0.00           H  
ATOM   3751 1HD1 ILE A 232     -10.175  25.598 -26.387  1.00  0.00           H  
ATOM   3752 2HD1 ILE A 232      -9.359  27.142 -26.038  1.00  0.00           H  
ATOM   3753 3HD1 ILE A 232      -8.503  25.603 -25.779  1.00  0.00           H  
ATOM   3754  N   ALA A 233     -10.788  29.076 -21.957  1.00  0.00           N  
ATOM   3755  CA  ALA A 233     -11.887  30.026 -22.112  1.00  0.00           C  
ATOM   3756  C   ALA A 233     -12.597  29.749 -23.436  1.00  0.00           C  
ATOM   3757  O   ALA A 233     -13.821  29.844 -23.524  1.00  0.00           O  
ATOM   3758  CB  ALA A 233     -11.360  31.448 -22.075  1.00  0.00           C  
ATOM   3759  H   ALA A 233      -9.860  29.425 -21.763  1.00  0.00           H  
ATOM   3760  HA  ALA A 233     -12.579  29.883 -21.282  1.00  0.00           H  
ATOM   3761 1HB  ALA A 233     -12.189  32.146 -22.192  1.00  0.00           H  
ATOM   3762 2HB  ALA A 233     -10.866  31.628 -21.120  1.00  0.00           H  
ATOM   3763 3HB  ALA A 233     -10.647  31.592 -22.886  1.00  0.00           H  
ATOM   3764  N   GLY A 234     -11.819  29.427 -24.468  1.00  0.00           N  
ATOM   3765  CA  GLY A 234     -12.412  29.083 -25.749  1.00  0.00           C  
ATOM   3766  C   GLY A 234     -12.112  29.898 -26.990  1.00  0.00           C  
ATOM   3767  O   GLY A 234     -11.904  29.329 -28.062  1.00  0.00           O  
ATOM   3768  H   GLY A 234     -10.814  29.418 -24.373  1.00  0.00           H  
ATOM   3769 1HA  GLY A 234     -12.131  28.064 -26.018  1.00  0.00           H  
ATOM   3770 2HA  GLY A 234     -13.498  29.106 -25.665  1.00  0.00           H  
ATOM   3771  N   GLY A 235     -12.093  31.220 -26.863  1.00  0.00           N  
ATOM   3772  CA  GLY A 235     -11.856  32.060 -28.026  1.00  0.00           C  
ATOM   3773  C   GLY A 235     -10.424  32.445 -28.347  1.00  0.00           C  
ATOM   3774  O   GLY A 235     -10.162  33.000 -29.414  1.00  0.00           O  
ATOM   3775  H   GLY A 235     -12.243  31.655 -25.964  1.00  0.00           H  
ATOM   3776 1HA  GLY A 235     -12.243  31.566 -28.918  1.00  0.00           H  
ATOM   3777 2HA  GLY A 235     -12.402  32.996 -27.916  1.00  0.00           H  
ATOM   3778  N   LEU A 236      -9.494  32.161 -27.443  1.00  0.00           N  
ATOM   3779  CA  LEU A 236      -8.100  32.511 -27.678  1.00  0.00           C  
ATOM   3780  C   LEU A 236      -7.185  31.329 -27.405  1.00  0.00           C  
ATOM   3781  O   LEU A 236      -6.936  30.994 -26.253  1.00  0.00           O  
ATOM   3782  CB  LEU A 236      -7.683  33.679 -26.778  1.00  0.00           C  
ATOM   3783  CG  LEU A 236      -8.305  35.060 -27.016  1.00  0.00           C  
ATOM   3784  CD1 LEU A 236      -9.755  35.076 -26.550  1.00  0.00           C  
ATOM   3785  CD2 LEU A 236      -7.508  36.100 -26.248  1.00  0.00           C  
ATOM   3786  H   LEU A 236      -9.746  31.697 -26.582  1.00  0.00           H  
ATOM   3787  HA  LEU A 236      -7.992  32.816 -28.718  1.00  0.00           H  
ATOM   3788 1HB  LEU A 236      -7.913  33.421 -25.746  1.00  0.00           H  
ATOM   3789 2HB  LEU A 236      -6.606  33.817 -26.865  1.00  0.00           H  
ATOM   3790  HG  LEU A 236      -8.282  35.291 -28.080  1.00  0.00           H  
ATOM   3791 1HD1 LEU A 236     -10.182  36.063 -26.726  1.00  0.00           H  
ATOM   3792 2HD1 LEU A 236     -10.324  34.331 -27.106  1.00  0.00           H  
ATOM   3793 3HD1 LEU A 236      -9.799  34.846 -25.486  1.00  0.00           H  
ATOM   3794 1HD2 LEU A 236      -7.943  37.086 -26.412  1.00  0.00           H  
ATOM   3795 2HD2 LEU A 236      -7.533  35.864 -25.184  1.00  0.00           H  
ATOM   3796 3HD2 LEU A 236      -6.475  36.096 -26.597  1.00  0.00           H  
ATOM   3797  N   PHE A 237      -6.687  30.697 -28.464  1.00  0.00           N  
ATOM   3798  CA  PHE A 237      -5.792  29.565 -28.278  1.00  0.00           C  
ATOM   3799  C   PHE A 237      -5.000  29.230 -29.530  1.00  0.00           C  
ATOM   3800  O   PHE A 237      -5.317  29.694 -30.624  1.00  0.00           O  
ATOM   3801  CB  PHE A 237      -6.566  28.335 -27.760  1.00  0.00           C  
ATOM   3802  CG  PHE A 237      -7.495  27.696 -28.764  1.00  0.00           C  
ATOM   3803  CD1 PHE A 237      -7.095  26.570 -29.486  1.00  0.00           C  
ATOM   3804  CD2 PHE A 237      -8.787  28.183 -28.948  1.00  0.00           C  
ATOM   3805  CE1 PHE A 237      -7.969  25.933 -30.367  1.00  0.00           C  
ATOM   3806  CE2 PHE A 237      -9.671  27.554 -29.829  1.00  0.00           C  
ATOM   3807  CZ  PHE A 237      -9.263  26.427 -30.538  1.00  0.00           C  
ATOM   3808  H   PHE A 237      -6.922  30.991 -29.401  1.00  0.00           H  
ATOM   3809  HA  PHE A 237      -5.039  29.837 -27.537  1.00  0.00           H  
ATOM   3810 1HB  PHE A 237      -5.860  27.572 -27.434  1.00  0.00           H  
ATOM   3811 2HB  PHE A 237      -7.162  28.619 -26.894  1.00  0.00           H  
ATOM   3812  HD1 PHE A 237      -6.083  26.185 -29.354  1.00  0.00           H  
ATOM   3813  HD2 PHE A 237      -9.106  29.066 -28.393  1.00  0.00           H  
ATOM   3814  HE1 PHE A 237      -7.640  25.054 -30.920  1.00  0.00           H  
ATOM   3815  HE2 PHE A 237     -10.680  27.945 -29.962  1.00  0.00           H  
ATOM   3816  HZ  PHE A 237      -9.948  25.932 -31.225  1.00  0.00           H  
ATOM   3817  N   VAL A 238      -3.953  28.436 -29.341  1.00  0.00           N  
ATOM   3818  CA  VAL A 238      -3.060  28.017 -30.412  1.00  0.00           C  
ATOM   3819  C   VAL A 238      -3.050  26.492 -30.435  1.00  0.00           C  
ATOM   3820  O   VAL A 238      -3.001  25.849 -29.385  1.00  0.00           O  
ATOM   3821  CB  VAL A 238      -1.637  28.557 -30.150  1.00  0.00           C  
ATOM   3822  CG1 VAL A 238      -0.652  27.989 -31.153  1.00  0.00           C  
ATOM   3823  CG2 VAL A 238      -1.651  30.079 -30.230  1.00  0.00           C  
ATOM   3824  H   VAL A 238      -3.777  28.112 -28.401  1.00  0.00           H  
ATOM   3825  HA  VAL A 238      -3.429  28.432 -31.351  1.00  0.00           H  
ATOM   3826  HB  VAL A 238      -1.316  28.242 -29.157  1.00  0.00           H  
ATOM   3827 1HG1 VAL A 238       0.343  28.383 -30.948  1.00  0.00           H  
ATOM   3828 2HG1 VAL A 238      -0.636  26.902 -31.072  1.00  0.00           H  
ATOM   3829 3HG1 VAL A 238      -0.954  28.275 -32.160  1.00  0.00           H  
ATOM   3830 1HG2 VAL A 238      -0.648  30.462 -30.044  1.00  0.00           H  
ATOM   3831 2HG2 VAL A 238      -1.981  30.389 -31.221  1.00  0.00           H  
ATOM   3832 3HG2 VAL A 238      -2.335  30.476 -29.480  1.00  0.00           H  
ATOM   3833  N   MET A 239      -3.091  25.916 -31.630  1.00  0.00           N  
ATOM   3834  CA  MET A 239      -3.117  24.465 -31.765  1.00  0.00           C  
ATOM   3835  C   MET A 239      -2.502  24.012 -33.080  1.00  0.00           C  
ATOM   3836  O   MET A 239      -2.711  24.635 -34.119  1.00  0.00           O  
ATOM   3837  CB  MET A 239      -4.571  23.984 -31.672  1.00  0.00           C  
ATOM   3838  CG  MET A 239      -4.792  22.475 -31.700  1.00  0.00           C  
ATOM   3839  SD  MET A 239      -6.537  22.111 -31.344  1.00  0.00           S  
ATOM   3840  CE  MET A 239      -6.466  20.337 -30.978  1.00  0.00           C  
ATOM   3841  H   MET A 239      -3.106  26.489 -32.463  1.00  0.00           H  
ATOM   3842  HA  MET A 239      -2.539  24.031 -30.950  1.00  0.00           H  
ATOM   3843 1HB  MET A 239      -5.016  24.348 -30.749  1.00  0.00           H  
ATOM   3844 2HB  MET A 239      -5.146  24.401 -32.500  1.00  0.00           H  
ATOM   3845 1HG  MET A 239      -4.521  22.086 -32.681  1.00  0.00           H  
ATOM   3846 2HG  MET A 239      -4.152  22.000 -30.957  1.00  0.00           H  
ATOM   3847 1HE  MET A 239      -7.466  19.974 -30.742  1.00  0.00           H  
ATOM   3848 2HE  MET A 239      -6.080  19.801 -31.847  1.00  0.00           H  
ATOM   3849 3HE  MET A 239      -5.807  20.167 -30.126  1.00  0.00           H  
ATOM   3850  N   ASP A 240      -1.726  22.934 -33.020  1.00  0.00           N  
ATOM   3851  CA  ASP A 240      -1.103  22.366 -34.210  1.00  0.00           C  
ATOM   3852  C   ASP A 240      -2.240  22.082 -35.195  1.00  0.00           C  
ATOM   3853  O   ASP A 240      -3.254  21.492 -34.816  1.00  0.00           O  
ATOM   3854  CB  ASP A 240      -0.368  21.083 -33.819  1.00  0.00           C  
ATOM   3855  CG  ASP A 240       0.265  20.380 -34.998  1.00  0.00           C  
ATOM   3856  OD1 ASP A 240      -0.453  19.649 -35.705  1.00  0.00           O  
ATOM   3857  OD2 ASP A 240       1.482  20.560 -35.215  1.00  0.00           O1-
ATOM   3858  H   ASP A 240      -1.565  22.497 -32.124  1.00  0.00           H  
ATOM   3859  HA  ASP A 240      -0.387  23.089 -34.604  1.00  0.00           H  
ATOM   3860 1HB  ASP A 240       0.413  21.318 -33.095  1.00  0.00           H  
ATOM   3861 2HB  ASP A 240      -1.065  20.396 -33.338  1.00  0.00           H  
ATOM   3862  N   LYS A 241      -2.086  22.498 -36.450  1.00  0.00           N  
ATOM   3863  CA  LYS A 241      -3.154  22.303 -37.428  1.00  0.00           C  
ATOM   3864  C   LYS A 241      -3.534  20.849 -37.686  1.00  0.00           C  
ATOM   3865  O   LYS A 241      -4.714  20.506 -37.666  1.00  0.00           O  
ATOM   3866  CB  LYS A 241      -2.758  22.960 -38.751  1.00  0.00           C  
ATOM   3867  CG  LYS A 241      -3.808  22.846 -39.849  1.00  0.00           C  
ATOM   3868  CD  LYS A 241      -3.360  23.556 -41.118  1.00  0.00           C  
ATOM   3869  CE  LYS A 241      -4.446  23.526 -42.183  1.00  0.00           C  
ATOM   3870  NZ  LYS A 241      -4.689  22.149 -42.691  1.00  0.00           N1+
ATOM   3871  H   LYS A 241      -1.231  22.952 -36.738  1.00  0.00           H  
ATOM   3872  HA  LYS A 241      -4.067  22.755 -37.040  1.00  0.00           H  
ATOM   3873 1HB  LYS A 241      -2.559  24.019 -38.587  1.00  0.00           H  
ATOM   3874 2HB  LYS A 241      -1.837  22.507 -39.121  1.00  0.00           H  
ATOM   3875 1HG  LYS A 241      -3.986  21.794 -40.075  1.00  0.00           H  
ATOM   3876 2HG  LYS A 241      -4.742  23.289 -39.506  1.00  0.00           H  
ATOM   3877 1HD  LYS A 241      -3.117  24.594 -40.887  1.00  0.00           H  
ATOM   3878 2HD  LYS A 241      -2.466  23.071 -41.510  1.00  0.00           H  
ATOM   3879 1HE  LYS A 241      -5.374  23.916 -41.767  1.00  0.00           H  
ATOM   3880 2HE  LYS A 241      -4.153  24.161 -43.019  1.00  0.00           H  
ATOM   3881 1HZ  LYS A 241      -5.414  22.172 -43.394  1.00  0.00           H  
ATOM   3882 2HZ  LYS A 241      -3.838  21.784 -43.096  1.00  0.00           H  
ATOM   3883 3HZ  LYS A 241      -4.979  21.554 -41.928  1.00  0.00           H  
ATOM   3884  N   PHE A 242      -2.550  19.990 -37.932  1.00  0.00           N  
ATOM   3885  CA  PHE A 242      -2.860  18.587 -38.179  1.00  0.00           C  
ATOM   3886  C   PHE A 242      -3.540  17.975 -36.960  1.00  0.00           C  
ATOM   3887  O   PHE A 242      -4.429  17.137 -37.092  1.00  0.00           O  
ATOM   3888  CB  PHE A 242      -1.590  17.804 -38.516  1.00  0.00           C  
ATOM   3889  CG  PHE A 242      -1.056  18.077 -39.893  1.00  0.00           C  
ATOM   3890  CD1 PHE A 242       0.228  18.569 -40.070  1.00  0.00           C  
ATOM   3891  CD2 PHE A 242      -1.838  17.844 -41.015  1.00  0.00           C  
ATOM   3892  CE1 PHE A 242       0.720  18.821 -41.337  1.00  0.00           C  
ATOM   3893  CE2 PHE A 242      -1.348  18.094 -42.282  1.00  0.00           C  
ATOM   3894  CZ  PHE A 242      -0.068  18.584 -42.443  1.00  0.00           C  
ATOM   3895  H   PHE A 242      -1.586  20.291 -37.952  1.00  0.00           H  
ATOM   3896  HA  PHE A 242      -3.538  18.527 -39.032  1.00  0.00           H  
ATOM   3897 1HB  PHE A 242      -0.811  18.047 -37.795  1.00  0.00           H  
ATOM   3898 2HB  PHE A 242      -1.789  16.736 -38.434  1.00  0.00           H  
ATOM   3899  HD1 PHE A 242       0.852  18.756 -39.195  1.00  0.00           H  
ATOM   3900  HD2 PHE A 242      -2.850  17.457 -40.888  1.00  0.00           H  
ATOM   3901  HE1 PHE A 242       1.731  19.208 -41.461  1.00  0.00           H  
ATOM   3902  HE2 PHE A 242      -1.973  17.906 -43.155  1.00  0.00           H  
ATOM   3903  HZ  PHE A 242       0.319  18.783 -43.441  1.00  0.00           H  
ATOM   3904  N   TYR A 243      -3.116  18.395 -35.771  1.00  0.00           N  
ATOM   3905  CA  TYR A 243      -3.699  17.876 -34.540  1.00  0.00           C  
ATOM   3906  C   TYR A 243      -5.160  18.326 -34.442  1.00  0.00           C  
ATOM   3907  O   TYR A 243      -6.039  17.540 -34.099  1.00  0.00           O  
ATOM   3908  CB  TYR A 243      -2.907  18.376 -33.326  1.00  0.00           C  
ATOM   3909  CG  TYR A 243      -3.265  17.671 -32.031  1.00  0.00           C  
ATOM   3910  CD1 TYR A 243      -3.604  18.399 -30.892  1.00  0.00           C  
ATOM   3911  CD2 TYR A 243      -3.264  16.273 -31.946  1.00  0.00           C  
ATOM   3912  CE1 TYR A 243      -3.935  17.757 -29.695  1.00  0.00           C  
ATOM   3913  CE2 TYR A 243      -3.593  15.625 -30.754  1.00  0.00           C  
ATOM   3914  CZ  TYR A 243      -3.929  16.376 -29.635  1.00  0.00           C  
ATOM   3915  OH  TYR A 243      -4.259  15.749 -28.451  1.00  0.00           O  
ATOM   3916  H   TYR A 243      -2.378  19.083 -35.715  1.00  0.00           H  
ATOM   3917  HA  TYR A 243      -3.652  16.787 -34.566  1.00  0.00           H  
ATOM   3918 1HB  TYR A 243      -1.840  18.238 -33.506  1.00  0.00           H  
ATOM   3919 2HB  TYR A 243      -3.082  19.443 -33.192  1.00  0.00           H  
ATOM   3920  HD1 TYR A 243      -3.613  19.488 -30.923  1.00  0.00           H  
ATOM   3921  HD2 TYR A 243      -3.002  15.675 -32.818  1.00  0.00           H  
ATOM   3922  HE1 TYR A 243      -4.196  18.346 -28.816  1.00  0.00           H  
ATOM   3923  HE2 TYR A 243      -3.585  14.536 -30.706  1.00  0.00           H  
ATOM   3924  HH  TYR A 243      -4.209  14.797 -28.569  1.00  0.00           H  
ATOM   3925  N   PHE A 244      -5.414  19.595 -34.748  1.00  0.00           N  
ATOM   3926  CA  PHE A 244      -6.776  20.119 -34.712  1.00  0.00           C  
ATOM   3927  C   PHE A 244      -7.666  19.273 -35.620  1.00  0.00           C  
ATOM   3928  O   PHE A 244      -8.785  18.907 -35.258  1.00  0.00           O  
ATOM   3929  CB  PHE A 244      -6.787  21.573 -35.196  1.00  0.00           C  
ATOM   3930  CG  PHE A 244      -8.164  22.165 -35.337  1.00  0.00           C  
ATOM   3931  CD1 PHE A 244      -8.863  22.612 -34.220  1.00  0.00           C  
ATOM   3932  CD2 PHE A 244      -8.753  22.289 -36.591  1.00  0.00           C  
ATOM   3933  CE1 PHE A 244     -10.136  23.181 -34.351  1.00  0.00           C  
ATOM   3934  CE2 PHE A 244     -10.021  22.856 -36.738  1.00  0.00           C  
ATOM   3935  CZ  PHE A 244     -10.712  23.302 -35.613  1.00  0.00           C  
ATOM   3936  H   PHE A 244      -4.656  20.209 -35.011  1.00  0.00           H  
ATOM   3937  HA  PHE A 244      -7.132  20.087 -33.681  1.00  0.00           H  
ATOM   3938 1HB  PHE A 244      -6.224  22.193 -34.499  1.00  0.00           H  
ATOM   3939 2HB  PHE A 244      -6.293  21.637 -36.164  1.00  0.00           H  
ATOM   3940  HD1 PHE A 244      -8.409  22.515 -33.233  1.00  0.00           H  
ATOM   3941  HD2 PHE A 244      -8.208  21.939 -37.469  1.00  0.00           H  
ATOM   3942  HE1 PHE A 244     -10.670  23.528 -33.468  1.00  0.00           H  
ATOM   3943  HE2 PHE A 244     -10.471  22.949 -37.725  1.00  0.00           H  
ATOM   3944  HZ  PHE A 244     -11.702  23.743 -35.721  1.00  0.00           H  
ATOM   3945  N   GLU A 245      -7.151  18.961 -36.802  1.00  0.00           N  
ATOM   3946  CA  GLU A 245      -7.891  18.177 -37.777  1.00  0.00           C  
ATOM   3947  C   GLU A 245      -8.129  16.731 -37.353  1.00  0.00           C  
ATOM   3948  O   GLU A 245      -9.270  16.272 -37.292  1.00  0.00           O  
ATOM   3949  CB  GLU A 245      -7.158  18.207 -39.122  1.00  0.00           C  
ATOM   3950  CG  GLU A 245      -7.100  19.592 -39.739  1.00  0.00           C  
ATOM   3951  CD  GLU A 245      -6.262  19.646 -41.000  1.00  0.00           C  
ATOM   3952  OE1 GLU A 245      -6.216  20.723 -41.637  1.00  0.00           O  
ATOM   3953  OE2 GLU A 245      -5.649  18.620 -41.359  1.00  0.00           O1-
ATOM   3954  H   GLU A 245      -6.220  19.277 -37.030  1.00  0.00           H  
ATOM   3955  HA  GLU A 245      -8.879  18.621 -37.899  1.00  0.00           H  
ATOM   3956 1HB  GLU A 245      -6.138  17.845 -38.988  1.00  0.00           H  
ATOM   3957 2HB  GLU A 245      -7.654  17.536 -39.822  1.00  0.00           H  
ATOM   3958 1HG  GLU A 245      -8.113  19.914 -39.978  1.00  0.00           H  
ATOM   3959 2HG  GLU A 245      -6.691  20.287 -39.007  1.00  0.00           H  
ATOM   3960  N   GLU A 246      -7.053  16.014 -37.055  1.00  0.00           N  
ATOM   3961  CA  GLU A 246      -7.182  14.614 -36.679  1.00  0.00           C  
ATOM   3962  C   GLU A 246      -7.955  14.379 -35.388  1.00  0.00           C  
ATOM   3963  O   GLU A 246      -8.575  13.323 -35.219  1.00  0.00           O  
ATOM   3964  CB  GLU A 246      -5.790  13.991 -36.550  1.00  0.00           C  
ATOM   3965  CG  GLU A 246      -5.041  13.847 -37.867  1.00  0.00           C  
ATOM   3966  CD  GLU A 246      -3.669  13.256 -37.698  1.00  0.00           C  
ATOM   3967  OE1 GLU A 246      -3.269  13.040 -36.579  1.00  0.00           O  
ATOM   3968  OE2 GLU A 246      -3.020  13.019 -38.690  1.00  0.00           O1-
ATOM   3969  H   GLU A 246      -6.134  16.433 -37.086  1.00  0.00           H  
ATOM   3970  HA  GLU A 246      -7.765  14.103 -37.445  1.00  0.00           H  
ATOM   3971 1HB  GLU A 246      -5.180  14.599 -35.882  1.00  0.00           H  
ATOM   3972 2HB  GLU A 246      -5.875  13.000 -36.104  1.00  0.00           H  
ATOM   3973 1HG  GLU A 246      -5.620  13.209 -38.535  1.00  0.00           H  
ATOM   3974 2HG  GLU A 246      -4.954  14.828 -38.332  1.00  0.00           H  
ATOM   3975  N   LEU A 247      -7.937  15.350 -34.480  1.00  0.00           N  
ATOM   3976  CA  LEU A 247      -8.657  15.184 -33.224  1.00  0.00           C  
ATOM   3977  C   LEU A 247     -10.153  15.436 -33.398  1.00  0.00           C  
ATOM   3978  O   LEU A 247     -10.941  15.156 -32.494  1.00  0.00           O  
ATOM   3979  CB  LEU A 247      -8.091  16.110 -32.140  1.00  0.00           C  
ATOM   3980  CG  LEU A 247      -8.187  15.527 -30.726  1.00  0.00           C  
ATOM   3981  CD1 LEU A 247      -7.254  14.324 -30.615  1.00  0.00           C  
ATOM   3982  CD2 LEU A 247      -7.814  16.576 -29.692  1.00  0.00           C  
ATOM   3983  H   LEU A 247      -7.429  16.207 -34.647  1.00  0.00           H  
ATOM   3984  HA  LEU A 247      -8.541  14.153 -32.892  1.00  0.00           H  
ATOM   3985 1HB  LEU A 247      -7.046  16.312 -32.367  1.00  0.00           H  
ATOM   3986 2HB  LEU A 247      -8.636  17.053 -32.169  1.00  0.00           H  
ATOM   3987  HG  LEU A 247      -9.208  15.192 -30.539  1.00  0.00           H  
ATOM   3988 1HD1 LEU A 247      -7.318  13.906 -29.610  1.00  0.00           H  
ATOM   3989 2HD1 LEU A 247      -7.547  13.568 -31.342  1.00  0.00           H  
ATOM   3990 3HD1 LEU A 247      -6.230  14.640 -30.811  1.00  0.00           H  
ATOM   3991 1HD2 LEU A 247      -7.887  16.145 -28.693  1.00  0.00           H  
ATOM   3992 2HD2 LEU A 247      -6.791  16.912 -29.868  1.00  0.00           H  
ATOM   3993 3HD2 LEU A 247      -8.493  17.424 -29.773  1.00  0.00           H  
ATOM   3994  N   GLY A 248     -10.547  15.969 -34.552  1.00  0.00           N  
ATOM   3995  CA  GLY A 248     -11.960  16.202 -34.791  1.00  0.00           C  
ATOM   3996  C   GLY A 248     -12.434  17.636 -34.952  1.00  0.00           C  
ATOM   3997  O   GLY A 248     -13.625  17.904 -34.790  1.00  0.00           O  
ATOM   3998  H   GLY A 248      -9.879  16.215 -35.268  1.00  0.00           H  
ATOM   3999 1HA  GLY A 248     -12.266  15.682 -35.698  1.00  0.00           H  
ATOM   4000 2HA  GLY A 248     -12.542  15.785 -33.970  1.00  0.00           H  
ATOM   4001  N   LYS A 249     -11.519  18.554 -35.254  1.00  0.00           N  
ATOM   4002  CA  LYS A 249     -11.867  19.963 -35.453  1.00  0.00           C  
ATOM   4003  C   LYS A 249     -12.853  20.432 -34.377  1.00  0.00           C  
ATOM   4004  O   LYS A 249     -12.489  20.528 -33.206  1.00  0.00           O  
ATOM   4005  CB  LYS A 249     -12.454  20.134 -36.858  1.00  0.00           C  
ATOM   4006  CG  LYS A 249     -11.533  19.590 -37.945  1.00  0.00           C  
ATOM   4007  CD  LYS A 249     -12.064  19.844 -39.352  1.00  0.00           C  
ATOM   4008  CE  LYS A 249     -11.094  19.295 -40.394  1.00  0.00           C  
ATOM   4009  NZ  LYS A 249     -11.519  19.583 -41.795  1.00  0.00           N1+
ATOM   4010  H   LYS A 249     -10.555  18.268 -35.350  1.00  0.00           H  
ATOM   4011  HA  LYS A 249     -10.958  20.560 -35.364  1.00  0.00           H  
ATOM   4012 1HB  LYS A 249     -13.412  19.619 -36.917  1.00  0.00           H  
ATOM   4013 2HB  LYS A 249     -12.639  21.192 -37.050  1.00  0.00           H  
ATOM   4014 1HG  LYS A 249     -10.552  20.061 -37.861  1.00  0.00           H  
ATOM   4015 2HG  LYS A 249     -11.413  18.515 -37.815  1.00  0.00           H  
ATOM   4016 1HD  LYS A 249     -13.035  19.361 -39.468  1.00  0.00           H  
ATOM   4017 2HD  LYS A 249     -12.191  20.916 -39.505  1.00  0.00           H  
ATOM   4018 1HE  LYS A 249     -10.108  19.732 -40.240  1.00  0.00           H  
ATOM   4019 2HE  LYS A 249     -11.010  18.214 -40.279  1.00  0.00           H  
ATOM   4020 1HZ  LYS A 249     -10.842  19.198 -42.438  1.00  0.00           H  
ATOM   4021 2HZ  LYS A 249     -12.423  19.165 -41.966  1.00  0.00           H  
ATOM   4022 3HZ  LYS A 249     -11.579  20.582 -41.930  1.00  0.00           H  
ATOM   4023  N   TYR A 250     -14.077  20.755 -34.786  1.00  0.00           N  
ATOM   4024  CA  TYR A 250     -15.149  21.145 -33.867  1.00  0.00           C  
ATOM   4025  C   TYR A 250     -16.366  20.334 -34.298  1.00  0.00           C  
ATOM   4026  O   TYR A 250     -16.422  19.840 -35.428  1.00  0.00           O  
ATOM   4027  CB  TYR A 250     -15.515  22.632 -33.988  1.00  0.00           C  
ATOM   4028  CG  TYR A 250     -14.675  23.598 -33.176  1.00  0.00           C  
ATOM   4029  CD1 TYR A 250     -14.077  24.697 -33.787  1.00  0.00           C  
ATOM   4030  CD2 TYR A 250     -14.515  23.442 -31.796  1.00  0.00           C  
ATOM   4031  CE1 TYR A 250     -13.344  25.622 -33.053  1.00  0.00           C  
ATOM   4032  CE2 TYR A 250     -13.774  24.372 -31.046  1.00  0.00           C  
ATOM   4033  CZ  TYR A 250     -13.197  25.456 -31.688  1.00  0.00           C  
ATOM   4034  OH  TYR A 250     -12.482  26.394 -30.981  1.00  0.00           O  
ATOM   4035  H   TYR A 250     -14.265  20.728 -35.778  1.00  0.00           H  
ATOM   4036  HA  TYR A 250     -14.811  20.969 -32.846  1.00  0.00           H  
ATOM   4037 1HB  TYR A 250     -15.437  22.943 -35.031  1.00  0.00           H  
ATOM   4038 2HB  TYR A 250     -16.550  22.778 -33.681  1.00  0.00           H  
ATOM   4039  HD1 TYR A 250     -14.179  24.845 -34.862  1.00  0.00           H  
ATOM   4040  HD2 TYR A 250     -14.968  22.591 -31.288  1.00  0.00           H  
ATOM   4041  HE1 TYR A 250     -12.885  26.475 -33.553  1.00  0.00           H  
ATOM   4042  HE2 TYR A 250     -13.657  24.236 -29.970  1.00  0.00           H  
ATOM   4043  HH  TYR A 250     -12.475  26.155 -30.051  1.00  0.00           H  
ATOM   4044  N   ASP A 251     -17.329  20.185 -33.398  1.00  0.00           N  
ATOM   4045  CA  ASP A 251     -18.566  19.478 -33.712  1.00  0.00           C  
ATOM   4046  C   ASP A 251     -19.319  20.439 -34.634  1.00  0.00           C  
ATOM   4047  O   ASP A 251     -19.823  21.471 -34.191  1.00  0.00           O  
ATOM   4048  CB  ASP A 251     -19.343  19.231 -32.417  1.00  0.00           C  
ATOM   4049  CG  ASP A 251     -20.684  18.565 -32.645  1.00  0.00           C  
ATOM   4050  OD1 ASP A 251     -21.335  18.235 -31.635  1.00  0.00           O  
ATOM   4051  OD2 ASP A 251     -21.085  18.372 -33.814  1.00  0.00           O1-
ATOM   4052  H   ASP A 251     -17.204  20.571 -32.473  1.00  0.00           H  
ATOM   4053  HA  ASP A 251     -18.311  18.521 -34.168  1.00  0.00           H  
ATOM   4054 1HB  ASP A 251     -18.752  18.602 -31.752  1.00  0.00           H  
ATOM   4055 2HB  ASP A 251     -19.511  20.180 -31.907  1.00  0.00           H  
ATOM   4056  N   MET A 252     -19.383  20.099 -35.917  1.00  0.00           N  
ATOM   4057  CA  MET A 252     -20.030  20.948 -36.912  1.00  0.00           C  
ATOM   4058  C   MET A 252     -21.516  21.206 -36.706  1.00  0.00           C  
ATOM   4059  O   MET A 252     -22.086  22.100 -37.336  1.00  0.00           O  
ATOM   4060  CB  MET A 252     -19.801  20.366 -38.307  1.00  0.00           C  
ATOM   4061  CG  MET A 252     -18.343  20.387 -38.752  1.00  0.00           C  
ATOM   4062  SD  MET A 252     -17.705  22.062 -38.966  1.00  0.00           S  
ATOM   4063  CE  MET A 252     -18.333  22.459 -40.585  1.00  0.00           C  
ATOM   4064  H   MET A 252     -18.969  19.225 -36.210  1.00  0.00           H  
ATOM   4065  HA  MET A 252     -19.585  21.940 -36.862  1.00  0.00           H  
ATOM   4066 1HB  MET A 252     -20.149  19.334 -38.332  1.00  0.00           H  
ATOM   4067 2HB  MET A 252     -20.387  20.927 -39.036  1.00  0.00           H  
ATOM   4068 1HG  MET A 252     -17.729  19.876 -38.011  1.00  0.00           H  
ATOM   4069 2HG  MET A 252     -18.244  19.858 -39.700  1.00  0.00           H  
ATOM   4070 1HE  MET A 252     -18.022  23.466 -40.860  1.00  0.00           H  
ATOM   4071 2HE  MET A 252     -17.941  21.746 -41.312  1.00  0.00           H  
ATOM   4072 3HE  MET A 252     -19.422  22.406 -40.576  1.00  0.00           H  
ATOM   4073  N   MET A 253     -22.152  20.434 -35.834  1.00  0.00           N  
ATOM   4074  CA  MET A 253     -23.569  20.639 -35.588  1.00  0.00           C  
ATOM   4075  C   MET A 253     -23.846  21.654 -34.484  1.00  0.00           C  
ATOM   4076  O   MET A 253     -25.003  21.951 -34.189  1.00  0.00           O  
ATOM   4077  CB  MET A 253     -24.226  19.303 -35.248  1.00  0.00           C  
ATOM   4078  CG  MET A 253     -24.206  18.285 -36.379  1.00  0.00           C  
ATOM   4079  SD  MET A 253     -25.073  18.864 -37.851  1.00  0.00           S  
ATOM   4080  CE  MET A 253     -26.767  18.849 -37.274  1.00  0.00           C  
ATOM   4081  H   MET A 253     -21.667  19.702 -35.334  1.00  0.00           H  
ATOM   4082  HA  MET A 253     -24.016  21.072 -36.483  1.00  0.00           H  
ATOM   4083 1HB  MET A 253     -23.724  18.859 -34.390  1.00  0.00           H  
ATOM   4084 2HB  MET A 253     -25.267  19.470 -34.967  1.00  0.00           H  
ATOM   4085 1HG  MET A 253     -23.175  18.064 -36.651  1.00  0.00           H  
ATOM   4086 2HG  MET A 253     -24.676  17.360 -36.045  1.00  0.00           H  
ATOM   4087 1HE  MET A 253     -27.428  19.184 -38.075  1.00  0.00           H  
ATOM   4088 2HE  MET A 253     -27.042  17.837 -36.976  1.00  0.00           H  
ATOM   4089 3HE  MET A 253     -26.866  19.519 -36.419  1.00  0.00           H  
ATOM   4090  N   MET A 254     -22.788  22.183 -33.872  1.00  0.00           N  
ATOM   4091  CA  MET A 254     -22.965  23.200 -32.839  1.00  0.00           C  
ATOM   4092  C   MET A 254     -23.227  24.524 -33.547  1.00  0.00           C  
ATOM   4093  O   MET A 254     -22.597  24.828 -34.564  1.00  0.00           O  
ATOM   4094  CB  MET A 254     -21.729  23.299 -31.947  1.00  0.00           C  
ATOM   4095  CG  MET A 254     -21.578  22.092 -31.034  1.00  0.00           C  
ATOM   4096  SD  MET A 254     -20.328  22.338 -29.772  1.00  0.00           S  
ATOM   4097  CE  MET A 254     -20.613  20.896 -28.745  1.00  0.00           C  
ATOM   4098  H   MET A 254     -21.854  21.887 -34.117  1.00  0.00           H  
ATOM   4099  HA  MET A 254     -23.814  22.918 -32.218  1.00  0.00           H  
ATOM   4100 1HB  MET A 254     -20.838  23.387 -32.568  1.00  0.00           H  
ATOM   4101 2HB  MET A 254     -21.791  24.201 -31.337  1.00  0.00           H  
ATOM   4102 1HG  MET A 254     -22.529  21.886 -30.544  1.00  0.00           H  
ATOM   4103 2HG  MET A 254     -21.304  21.219 -31.627  1.00  0.00           H  
ATOM   4104 1HE  MET A 254     -19.912  20.901 -27.910  1.00  0.00           H  
ATOM   4105 2HE  MET A 254     -21.634  20.917 -28.362  1.00  0.00           H  
ATOM   4106 3HE  MET A 254     -20.466  19.993 -29.337  1.00  0.00           H  
ATOM   4107  N   ASP A 255     -24.158  25.297 -32.998  1.00  0.00           N  
ATOM   4108  CA  ASP A 255     -24.576  26.563 -33.587  1.00  0.00           C  
ATOM   4109  C   ASP A 255     -24.251  27.824 -32.787  1.00  0.00           C  
ATOM   4110  O   ASP A 255     -24.240  27.817 -31.557  1.00  0.00           O  
ATOM   4111  CB  ASP A 255     -26.097  26.543 -33.805  1.00  0.00           C  
ATOM   4112  CG  ASP A 255     -26.554  25.439 -34.741  1.00  0.00           C  
ATOM   4113  OD1 ASP A 255     -27.688  24.948 -34.557  1.00  0.00           O  
ATOM   4114  OD2 ASP A 255     -25.803  25.071 -35.665  1.00  0.00           O1-
ATOM   4115  H   ASP A 255     -24.589  24.989 -32.139  1.00  0.00           H  
ATOM   4116  HA  ASP A 255     -24.077  26.679 -34.549  1.00  0.00           H  
ATOM   4117 1HB  ASP A 255     -26.601  26.415 -32.846  1.00  0.00           H  
ATOM   4118 2HB  ASP A 255     -26.418  27.500 -34.217  1.00  0.00           H  
ATOM   4119  N   VAL A 256     -24.013  28.904 -33.525  1.00  0.00           N  
ATOM   4120  CA  VAL A 256     -23.752  30.238 -32.989  1.00  0.00           C  
ATOM   4121  C   VAL A 256     -22.625  30.446 -31.982  1.00  0.00           C  
ATOM   4122  O   VAL A 256     -21.545  30.923 -32.336  1.00  0.00           O  
ATOM   4123  CB  VAL A 256     -25.047  30.825 -32.368  1.00  0.00           C  
ATOM   4124  CG1 VAL A 256     -24.859  32.299 -32.059  1.00  0.00           C  
ATOM   4125  CG2 VAL A 256     -26.226  30.618 -33.321  1.00  0.00           C  
ATOM   4126  H   VAL A 256     -24.018  28.768 -34.526  1.00  0.00           H  
ATOM   4127  HA  VAL A 256     -23.430  30.883 -33.808  1.00  0.00           H  
ATOM   4128  HB  VAL A 256     -25.249  30.317 -31.424  1.00  0.00           H  
ATOM   4129 1HG1 VAL A 256     -25.775  32.700 -31.622  1.00  0.00           H  
ATOM   4130 2HG1 VAL A 256     -24.039  32.422 -31.352  1.00  0.00           H  
ATOM   4131 3HG1 VAL A 256     -24.631  32.838 -32.978  1.00  0.00           H  
ATOM   4132 1HG2 VAL A 256     -27.131  31.032 -32.876  1.00  0.00           H  
ATOM   4133 2HG2 VAL A 256     -26.023  31.122 -34.266  1.00  0.00           H  
ATOM   4134 3HG2 VAL A 256     -26.366  29.552 -33.500  1.00  0.00           H  
ATOM   4135  N   TRP A 257     -22.896  30.106 -30.729  1.00  0.00           N  
ATOM   4136  CA  TRP A 257     -21.944  30.299 -29.643  1.00  0.00           C  
ATOM   4137  C   TRP A 257     -22.283  29.360 -28.493  1.00  0.00           C  
ATOM   4138  O   TRP A 257     -23.427  28.932 -28.353  1.00  0.00           O  
ATOM   4139  CB  TRP A 257     -22.047  31.754 -29.154  1.00  0.00           C  
ATOM   4140  CG  TRP A 257     -21.239  32.091 -27.931  1.00  0.00           C  
ATOM   4141  CD1 TRP A 257     -20.011  32.680 -27.900  1.00  0.00           C  
ATOM   4142  CD2 TRP A 257     -21.609  31.864 -26.563  1.00  0.00           C  
ATOM   4143  CE2 TRP A 257     -20.550  32.342 -25.759  1.00  0.00           C  
ATOM   4144  CE3 TRP A 257     -22.729  31.300 -25.940  1.00  0.00           C  
ATOM   4145  NE1 TRP A 257     -19.589  32.837 -26.600  1.00  0.00           N  
ATOM   4146  CZ2 TRP A 257     -20.577  32.277 -24.362  1.00  0.00           C  
ATOM   4147  CZ3 TRP A 257     -22.758  31.235 -24.547  1.00  0.00           C  
ATOM   4148  CH2 TRP A 257     -21.685  31.720 -23.777  1.00  0.00           C  
ATOM   4149  H   TRP A 257     -23.798  29.699 -30.526  1.00  0.00           H  
ATOM   4150  HA  TRP A 257     -20.940  30.114 -30.025  1.00  0.00           H  
ATOM   4151 1HB  TRP A 257     -21.727  32.429 -29.948  1.00  0.00           H  
ATOM   4152 2HB  TRP A 257     -23.088  31.989 -28.926  1.00  0.00           H  
ATOM   4153  HD1 TRP A 257     -19.443  32.983 -28.778  1.00  0.00           H  
ATOM   4154  HE1 TRP A 257     -18.715  33.250 -26.311  1.00  0.00           H  
ATOM   4155  HE3 TRP A 257     -23.563  30.923 -26.531  1.00  0.00           H  
ATOM   4156  HZ2 TRP A 257     -19.758  32.652 -23.748  1.00  0.00           H  
ATOM   4157  HZ3 TRP A 257     -23.634  30.797 -24.069  1.00  0.00           H  
ATOM   4158  HH2 TRP A 257     -21.738  31.649 -22.690  1.00  0.00           H  
ATOM   4159  N   GLY A 258     -21.282  29.045 -27.677  1.00  0.00           N  
ATOM   4160  CA  GLY A 258     -21.500  28.202 -26.515  1.00  0.00           C  
ATOM   4161  C   GLY A 258     -21.292  26.709 -26.636  1.00  0.00           C  
ATOM   4162  O   GLY A 258     -21.693  26.086 -27.620  1.00  0.00           O  
ATOM   4163  H   GLY A 258     -20.353  29.394 -27.864  1.00  0.00           H  
ATOM   4164 1HA  GLY A 258     -20.843  28.521 -25.705  1.00  0.00           H  
ATOM   4165 2HA  GLY A 258     -22.525  28.324 -26.165  1.00  0.00           H  
ATOM   4166  N   GLY A 259     -20.651  26.139 -25.619  1.00  0.00           N  
ATOM   4167  CA  GLY A 259     -20.433  24.707 -25.574  1.00  0.00           C  
ATOM   4168  C   GLY A 259     -19.267  24.108 -26.331  1.00  0.00           C  
ATOM   4169  O   GLY A 259     -18.855  22.987 -26.024  1.00  0.00           O  
ATOM   4170  H   GLY A 259     -20.309  26.712 -24.861  1.00  0.00           H  
ATOM   4171 1HA  GLY A 259     -20.297  24.392 -24.539  1.00  0.00           H  
ATOM   4172 2HA  GLY A 259     -21.315  24.192 -25.953  1.00  0.00           H  
ATOM   4173  N   GLU A 260     -18.715  24.825 -27.304  1.00  0.00           N  
ATOM   4174  CA  GLU A 260     -17.613  24.253 -28.062  1.00  0.00           C  
ATOM   4175  C   GLU A 260     -16.317  24.279 -27.266  1.00  0.00           C  
ATOM   4176  O   GLU A 260     -15.438  23.445 -27.485  1.00  0.00           O  
ATOM   4177  CB  GLU A 260     -17.427  25.008 -29.380  1.00  0.00           C  
ATOM   4178  CG  GLU A 260     -16.916  26.433 -29.222  1.00  0.00           C  
ATOM   4179  CD  GLU A 260     -16.826  27.170 -30.529  1.00  0.00           C  
ATOM   4180  OE1 GLU A 260     -16.100  26.732 -31.388  1.00  0.00           O  
ATOM   4181  OE2 GLU A 260     -17.486  28.173 -30.669  1.00  0.00           O1-
ATOM   4182  H   GLU A 260     -19.039  25.754 -27.531  1.00  0.00           H  
ATOM   4183  HA  GLU A 260     -17.833  23.203 -28.254  1.00  0.00           H  
ATOM   4184 1HB  GLU A 260     -16.722  24.468 -30.012  1.00  0.00           H  
ATOM   4185 2HB  GLU A 260     -18.377  25.050 -29.912  1.00  0.00           H  
ATOM   4186 1HG  GLU A 260     -17.586  26.976 -28.557  1.00  0.00           H  
ATOM   4187 2HG  GLU A 260     -15.931  26.405 -28.758  1.00  0.00           H  
ATOM   4188  N   ASN A 261     -16.198  25.228 -26.340  1.00  0.00           N  
ATOM   4189  CA  ASN A 261     -15.000  25.298 -25.520  1.00  0.00           C  
ATOM   4190  C   ASN A 261     -14.996  24.122 -24.544  1.00  0.00           C  
ATOM   4191  O   ASN A 261     -13.947  23.548 -24.248  1.00  0.00           O  
ATOM   4192  CB  ASN A 261     -14.912  26.649 -24.782  1.00  0.00           C  
ATOM   4193  CG  ASN A 261     -16.175  27.003 -24.002  1.00  0.00           C  
ATOM   4194  ND2 ASN A 261     -16.096  28.074 -23.218  1.00  0.00           N  
ATOM   4195  OD1 ASN A 261     -17.203  26.342 -24.109  1.00  0.00           O  
ATOM   4196  H   ASN A 261     -16.933  25.907 -26.197  1.00  0.00           H  
ATOM   4197  HA  ASN A 261     -14.129  25.204 -26.170  1.00  0.00           H  
ATOM   4198 1HB  ASN A 261     -14.073  26.628 -24.084  1.00  0.00           H  
ATOM   4199 2HB  ASN A 261     -14.720  27.444 -25.502  1.00  0.00           H  
ATOM   4200 1HD2 ASN A 261     -16.890  28.360 -22.680  1.00  0.00           H  
ATOM   4201 2HD2 ASN A 261     -15.244  28.593 -23.166  1.00  0.00           H  
ATOM   4202  N   LEU A 262     -16.181  23.747 -24.074  1.00  0.00           N  
ATOM   4203  CA  LEU A 262     -16.316  22.621 -23.159  1.00  0.00           C  
ATOM   4204  C   LEU A 262     -15.980  21.338 -23.919  1.00  0.00           C  
ATOM   4205  O   LEU A 262     -15.228  20.490 -23.436  1.00  0.00           O  
ATOM   4206  CB  LEU A 262     -17.755  22.532 -22.635  1.00  0.00           C  
ATOM   4207  CG  LEU A 262     -18.018  21.523 -21.515  1.00  0.00           C  
ATOM   4208  CD1 LEU A 262     -17.658  22.152 -20.179  1.00  0.00           C  
ATOM   4209  CD2 LEU A 262     -19.485  21.113 -21.516  1.00  0.00           C  
ATOM   4210  H   LEU A 262     -17.008  24.252 -24.357  1.00  0.00           H  
ATOM   4211  HA  LEU A 262     -15.646  22.780 -22.314  1.00  0.00           H  
ATOM   4212 1HB  LEU A 262     -18.050  23.512 -22.261  1.00  0.00           H  
ATOM   4213 2HB  LEU A 262     -18.411  22.270 -23.465  1.00  0.00           H  
ATOM   4214  HG  LEU A 262     -17.399  20.639 -21.670  1.00  0.00           H  
ATOM   4215 1HD1 LEU A 262     -17.843  21.437 -19.378  1.00  0.00           H  
ATOM   4216 2HD1 LEU A 262     -16.604  22.430 -20.179  1.00  0.00           H  
ATOM   4217 3HD1 LEU A 262     -18.268  23.040 -20.020  1.00  0.00           H  
ATOM   4218 1HD2 LEU A 262     -19.663  20.394 -20.715  1.00  0.00           H  
ATOM   4219 2HD2 LEU A 262     -20.109  21.994 -21.357  1.00  0.00           H  
ATOM   4220 3HD2 LEU A 262     -19.736  20.659 -22.474  1.00  0.00           H  
ATOM   4221  N   GLU A 263     -16.538  21.218 -25.120  1.00  0.00           N  
ATOM   4222  CA  GLU A 263     -16.335  20.037 -25.950  1.00  0.00           C  
ATOM   4223  C   GLU A 263     -14.881  19.786 -26.336  1.00  0.00           C  
ATOM   4224  O   GLU A 263     -14.385  18.668 -26.189  1.00  0.00           O  
ATOM   4225  CB  GLU A 263     -17.207  20.146 -27.207  1.00  0.00           C  
ATOM   4226  CG  GLU A 263     -17.276  18.890 -28.082  1.00  0.00           C  
ATOM   4227  CD  GLU A 263     -16.150  18.802 -29.095  1.00  0.00           C  
ATOM   4228  OE1 GLU A 263     -15.556  19.851 -29.426  1.00  0.00           O  
ATOM   4229  OE2 GLU A 263     -15.870  17.685 -29.579  1.00  0.00           O1-
ATOM   4230  H   GLU A 263     -17.119  21.968 -25.467  1.00  0.00           H  
ATOM   4231  HA  GLU A 263     -16.635  19.158 -25.379  1.00  0.00           H  
ATOM   4232 1HB  GLU A 263     -18.229  20.393 -26.920  1.00  0.00           H  
ATOM   4233 2HB  GLU A 263     -16.840  20.956 -27.836  1.00  0.00           H  
ATOM   4234 1HG  GLU A 263     -17.237  18.009 -27.441  1.00  0.00           H  
ATOM   4235 2HG  GLU A 263     -18.229  18.879 -28.609  1.00  0.00           H  
ATOM   4236  N   ILE A 264     -14.189  20.810 -26.827  1.00  0.00           N  
ATOM   4237  CA  ILE A 264     -12.803  20.611 -27.229  1.00  0.00           C  
ATOM   4238  C   ILE A 264     -11.898  20.398 -26.019  1.00  0.00           C  
ATOM   4239  O   ILE A 264     -10.895  19.690 -26.108  1.00  0.00           O  
ATOM   4240  CB  ILE A 264     -12.274  21.788 -28.106  1.00  0.00           C  
ATOM   4241  CG1 ILE A 264     -10.956  21.376 -28.773  1.00  0.00           C  
ATOM   4242  CG2 ILE A 264     -12.086  23.052 -27.265  1.00  0.00           C  
ATOM   4243  CD1 ILE A 264     -10.451  22.363 -29.815  1.00  0.00           C  
ATOM   4244  H   ILE A 264     -14.603  21.725 -26.928  1.00  0.00           H  
ATOM   4245  HA  ILE A 264     -12.743  19.698 -27.820  1.00  0.00           H  
ATOM   4246  HB  ILE A 264     -12.989  21.998 -28.901  1.00  0.00           H  
ATOM   4247 1HG1 ILE A 264     -10.184  21.261 -28.013  1.00  0.00           H  
ATOM   4248 2HG1 ILE A 264     -11.082  20.407 -29.260  1.00  0.00           H  
ATOM   4249 1HG2 ILE A 264     -11.716  23.858 -27.898  1.00  0.00           H  
ATOM   4250 2HG2 ILE A 264     -13.041  23.343 -26.828  1.00  0.00           H  
ATOM   4251 3HG2 ILE A 264     -11.368  22.857 -26.469  1.00  0.00           H  
ATOM   4252 1HD1 ILE A 264      -9.516  21.997 -30.239  1.00  0.00           H  
ATOM   4253 2HD1 ILE A 264     -11.193  22.466 -30.607  1.00  0.00           H  
ATOM   4254 3HD1 ILE A 264     -10.283  23.331 -29.346  1.00  0.00           H  
ATOM   4255  N   SER A 265     -12.262  20.989 -24.884  1.00  0.00           N  
ATOM   4256  CA  SER A 265     -11.463  20.830 -23.672  1.00  0.00           C  
ATOM   4257  C   SER A 265     -11.536  19.382 -23.194  1.00  0.00           C  
ATOM   4258  O   SER A 265     -10.511  18.777 -22.873  1.00  0.00           O  
ATOM   4259  CB  SER A 265     -11.953  21.766 -22.585  1.00  0.00           C  
ATOM   4260  OG  SER A 265     -11.745  23.105 -22.943  1.00  0.00           O  
ATOM   4261  H   SER A 265     -13.098  21.555 -24.851  1.00  0.00           H  
ATOM   4262  HA  SER A 265     -10.426  21.080 -23.906  1.00  0.00           H  
ATOM   4263 1HB  SER A 265     -13.015  21.595 -22.408  1.00  0.00           H  
ATOM   4264 2HB  SER A 265     -11.428  21.550 -21.655  1.00  0.00           H  
ATOM   4265  HG  SER A 265     -12.462  23.333 -23.540  1.00  0.00           H  
ATOM   4266  N   PHE A 266     -12.747  18.827 -23.148  1.00  0.00           N  
ATOM   4267  CA  PHE A 266     -12.922  17.441 -22.721  1.00  0.00           C  
ATOM   4268  C   PHE A 266     -12.233  16.510 -23.712  1.00  0.00           C  
ATOM   4269  O   PHE A 266     -11.516  15.589 -23.327  1.00  0.00           O  
ATOM   4270  CB  PHE A 266     -14.406  17.048 -22.665  1.00  0.00           C  
ATOM   4271  CG  PHE A 266     -15.193  17.710 -21.561  1.00  0.00           C  
ATOM   4272  CD1 PHE A 266     -14.559  18.325 -20.484  1.00  0.00           C  
ATOM   4273  CD2 PHE A 266     -16.586  17.683 -21.589  1.00  0.00           C  
ATOM   4274  CE1 PHE A 266     -15.304  18.902 -19.451  1.00  0.00           C  
ATOM   4275  CE2 PHE A 266     -17.339  18.254 -20.560  1.00  0.00           C  
ATOM   4276  CZ  PHE A 266     -16.698  18.865 -19.491  1.00  0.00           C  
ATOM   4277  H   PHE A 266     -13.559  19.368 -23.411  1.00  0.00           H  
ATOM   4278  HA  PHE A 266     -12.505  17.332 -21.719  1.00  0.00           H  
ATOM   4279 1HB  PHE A 266     -14.885  17.301 -23.611  1.00  0.00           H  
ATOM   4280 2HB  PHE A 266     -14.492  15.970 -22.533  1.00  0.00           H  
ATOM   4281  HD1 PHE A 266     -13.470  18.354 -20.454  1.00  0.00           H  
ATOM   4282  HD2 PHE A 266     -17.088  17.202 -22.429  1.00  0.00           H  
ATOM   4283  HE1 PHE A 266     -14.795  19.380 -18.614  1.00  0.00           H  
ATOM   4284  HE2 PHE A 266     -18.427  18.221 -20.595  1.00  0.00           H  
ATOM   4285  HZ  PHE A 266     -17.279  19.313 -18.687  1.00  0.00           H  
ATOM   4286  N   ARG A 267     -12.461  16.758 -24.998  1.00  0.00           N  
ATOM   4287  CA  ARG A 267     -11.884  15.928 -26.049  1.00  0.00           C  
ATOM   4288  C   ARG A 267     -10.358  15.898 -26.034  1.00  0.00           C  
ATOM   4289  O   ARG A 267      -9.755  14.830 -26.131  1.00  0.00           O  
ATOM   4290  CB  ARG A 267     -12.399  16.401 -27.413  1.00  0.00           C  
ATOM   4291  CG  ARG A 267     -11.767  15.721 -28.614  1.00  0.00           C  
ATOM   4292  CD  ARG A 267     -12.710  15.782 -29.811  1.00  0.00           C  
ATOM   4293  NE  ARG A 267     -13.219  17.131 -30.045  1.00  0.00           N  
ATOM   4294  CZ  ARG A 267     -12.557  18.087 -30.687  1.00  0.00           C  
ATOM   4295  NH1 ARG A 267     -13.107  19.286 -30.841  1.00  0.00           N1+
ATOM   4296  NH2 ARG A 267     -11.357  17.842 -31.197  1.00  0.00           N  
ATOM   4297  H   ARG A 267     -13.048  17.539 -25.254  1.00  0.00           H  
ATOM   4298  HA  ARG A 267     -12.197  14.897 -25.886  1.00  0.00           H  
ATOM   4299 1HB  ARG A 267     -13.474  16.237 -27.473  1.00  0.00           H  
ATOM   4300 2HB  ARG A 267     -12.225  17.472 -27.516  1.00  0.00           H  
ATOM   4301 1HG  ARG A 267     -10.834  16.224 -28.870  1.00  0.00           H  
ATOM   4302 2HG  ARG A 267     -11.562  14.677 -28.374  1.00  0.00           H  
ATOM   4303 1HD  ARG A 267     -12.182  15.460 -30.708  1.00  0.00           H  
ATOM   4304 2HD  ARG A 267     -13.562  15.125 -29.636  1.00  0.00           H  
ATOM   4305  HE  ARG A 267     -14.140  17.353 -29.694  1.00  0.00           H  
ATOM   4306 1HH1 ARG A 267     -14.029  19.470 -30.470  1.00  0.00           H  
ATOM   4307 2HH1 ARG A 267     -12.605  20.014 -31.328  1.00  0.00           H  
ATOM   4308 1HH2 ARG A 267     -10.943  16.925 -31.097  1.00  0.00           H  
ATOM   4309 2HH2 ARG A 267     -10.857  18.572 -31.684  1.00  0.00           H  
ATOM   4310  N   VAL A 268      -9.728  17.061 -25.903  1.00  0.00           N  
ATOM   4311  CA  VAL A 268      -8.271  17.106 -25.881  1.00  0.00           C  
ATOM   4312  C   VAL A 268      -7.704  16.328 -24.695  1.00  0.00           C  
ATOM   4313  O   VAL A 268      -6.801  15.511 -24.861  1.00  0.00           O  
ATOM   4314  CB  VAL A 268      -7.754  18.558 -25.832  1.00  0.00           C  
ATOM   4315  CG1 VAL A 268      -6.248  18.576 -25.557  1.00  0.00           C  
ATOM   4316  CG2 VAL A 268      -8.037  19.241 -27.161  1.00  0.00           C  
ATOM   4317  H   VAL A 268     -10.248  17.923 -25.818  1.00  0.00           H  
ATOM   4318  HA  VAL A 268      -7.896  16.641 -26.794  1.00  0.00           H  
ATOM   4319  HB  VAL A 268      -8.264  19.089 -25.028  1.00  0.00           H  
ATOM   4320 1HG1 VAL A 268      -5.897  19.607 -25.525  1.00  0.00           H  
ATOM   4321 2HG1 VAL A 268      -6.047  18.095 -24.600  1.00  0.00           H  
ATOM   4322 3HG1 VAL A 268      -5.727  18.041 -26.350  1.00  0.00           H  
ATOM   4323 1HG2 VAL A 268      -7.673  20.268 -27.128  1.00  0.00           H  
ATOM   4324 2HG2 VAL A 268      -7.532  18.703 -27.963  1.00  0.00           H  
ATOM   4325 3HG2 VAL A 268      -9.112  19.244 -27.347  1.00  0.00           H  
ATOM   4326  N   TRP A 269      -8.234  16.573 -23.501  1.00  0.00           N  
ATOM   4327  CA  TRP A 269      -7.744  15.876 -22.314  1.00  0.00           C  
ATOM   4328  C   TRP A 269      -8.038  14.382 -22.357  1.00  0.00           C  
ATOM   4329  O   TRP A 269      -7.140  13.556 -22.195  1.00  0.00           O  
ATOM   4330  CB  TRP A 269      -8.362  16.470 -21.046  1.00  0.00           C  
ATOM   4331  CG  TRP A 269      -7.661  17.689 -20.540  1.00  0.00           C  
ATOM   4332  CD1 TRP A 269      -8.188  18.941 -20.397  1.00  0.00           C  
ATOM   4333  CD2 TRP A 269      -6.315  17.763 -20.051  1.00  0.00           C  
ATOM   4334  CE2 TRP A 269      -6.098  19.091 -19.621  1.00  0.00           C  
ATOM   4335  CE3 TRP A 269      -5.273  16.834 -19.933  1.00  0.00           C  
ATOM   4336  NE1 TRP A 269      -7.256  19.788 -19.844  1.00  0.00           N  
ATOM   4337  CZ2 TRP A 269      -4.879  19.517 -19.078  1.00  0.00           C  
ATOM   4338  CZ3 TRP A 269      -4.059  17.257 -19.392  1.00  0.00           C  
ATOM   4339  CH2 TRP A 269      -3.875  18.588 -18.971  1.00  0.00           C  
ATOM   4340  H   TRP A 269      -8.982  17.245 -23.403  1.00  0.00           H  
ATOM   4341  HA  TRP A 269      -6.662  15.997 -22.263  1.00  0.00           H  
ATOM   4342 1HB  TRP A 269      -9.403  16.733 -21.236  1.00  0.00           H  
ATOM   4343 2HB  TRP A 269      -8.353  15.723 -20.253  1.00  0.00           H  
ATOM   4344  HD1 TRP A 269      -9.199  19.227 -20.681  1.00  0.00           H  
ATOM   4345  HE1 TRP A 269      -7.400  20.766 -19.636  1.00  0.00           H  
ATOM   4346  HE3 TRP A 269      -5.409  15.801 -20.254  1.00  0.00           H  
ATOM   4347  HZ2 TRP A 269      -4.719  20.544 -18.748  1.00  0.00           H  
ATOM   4348  HZ3 TRP A 269      -3.252  16.528 -19.305  1.00  0.00           H  
ATOM   4349  HH2 TRP A 269      -2.913  18.885 -18.551  1.00  0.00           H  
ATOM   4350  N   GLN A 270      -9.295  14.028 -22.582  1.00  0.00           N  
ATOM   4351  CA  GLN A 270      -9.659  12.620 -22.612  1.00  0.00           C  
ATOM   4352  C   GLN A 270      -8.992  11.836 -23.734  1.00  0.00           C  
ATOM   4353  O   GLN A 270      -8.753  10.633 -23.601  1.00  0.00           O  
ATOM   4354  CB  GLN A 270     -11.180  12.478 -22.706  1.00  0.00           C  
ATOM   4355  CG  GLN A 270     -11.896  12.847 -21.410  1.00  0.00           C  
ATOM   4356  CD  GLN A 270     -13.405  12.901 -21.550  1.00  0.00           C  
ATOM   4357  NE2 GLN A 270     -14.043  11.734 -21.546  1.00  0.00           N  
ATOM   4358  OE1 GLN A 270     -13.991  13.976 -21.659  1.00  0.00           O  
ATOM   4359  H   GLN A 270     -10.010  14.725 -22.734  1.00  0.00           H  
ATOM   4360  HA  GLN A 270      -9.317  12.155 -21.688  1.00  0.00           H  
ATOM   4361 1HB  GLN A 270     -11.556  13.117 -23.505  1.00  0.00           H  
ATOM   4362 2HB  GLN A 270     -11.435  11.449 -22.964  1.00  0.00           H  
ATOM   4363 1HG  GLN A 270     -11.658  12.101 -20.652  1.00  0.00           H  
ATOM   4364 2HG  GLN A 270     -11.556  13.830 -21.087  1.00  0.00           H  
ATOM   4365 1HE2 GLN A 270     -15.040  11.709 -21.635  1.00  0.00           H  
ATOM   4366 2HE2 GLN A 270     -13.528  10.882 -21.454  1.00  0.00           H  
ATOM   4367  N   CYS A 271      -8.658  12.511 -24.828  1.00 12.57           N  
ATOM   4368  CA  CYS A 271      -8.058  11.802 -25.940  1.00 12.79           C  
ATOM   4369  C   CYS A 271      -6.540  11.883 -26.082  1.00 12.22           C  
ATOM   4370  O   CYS A 271      -6.001  11.632 -27.160  1.00 12.82           O  
ATOM   4371  CB  CYS A 271      -8.756  12.212 -27.235  1.00 15.55           C  
ATOM   4372  SG  CYS A 271     -10.579  12.137 -27.123  1.00 15.07           S  
ATOM   4373  H   CYS A 271      -8.812  13.506 -24.905  1.00  0.00           H  
ATOM   4374  HA  CYS A 271      -8.195  10.732 -25.782  1.00  0.00           H  
ATOM   4375 1HB  CYS A 271      -8.467  13.230 -27.498  1.00  0.00           H  
ATOM   4376 2HB  CYS A 271      -8.432  11.560 -28.046  1.00  0.00           H  
ATOM   4377  N   GLY A 272      -5.849  12.240 -25.001  1.00  0.00           N  
ATOM   4378  CA  GLY A 272      -4.395  12.242 -25.045  1.00  0.00           C  
ATOM   4379  C   GLY A 272      -3.571  13.516 -25.030  1.00  0.00           C  
ATOM   4380  O   GLY A 272      -2.373  13.459 -24.753  1.00  0.00           O  
ATOM   4381  H   GLY A 272      -6.316  12.511 -24.147  1.00  0.00           H  
ATOM   4382 1HA  GLY A 272      -4.004  11.680 -24.196  1.00  0.00           H  
ATOM   4383 2HA  GLY A 272      -4.058  11.736 -25.948  1.00  0.00           H  
ATOM   4384  N   GLY A 273      -4.181  14.654 -25.335  1.00  0.00           N  
ATOM   4385  CA  GLY A 273      -3.428  15.897 -25.335  1.00  0.00           C  
ATOM   4386  C   GLY A 273      -3.523  16.610 -24.002  1.00  0.00           C  
ATOM   4387  O   GLY A 273      -3.822  15.988 -22.985  1.00  0.00           O  
ATOM   4388  H   GLY A 273      -5.164  14.673 -25.568  1.00  0.00           H  
ATOM   4389 1HA  GLY A 273      -2.383  15.690 -25.562  1.00  0.00           H  
ATOM   4390 2HA  GLY A 273      -3.805  16.549 -26.123  1.00  0.00           H  
ATOM   4391  N   SER A 274      -3.261  17.914 -24.001  1.00  0.00           N  
ATOM   4392  CA  SER A 274      -3.348  18.697 -22.772  1.00  0.00           C  
ATOM   4393  C   SER A 274      -3.691  20.147 -23.085  1.00  0.00           C  
ATOM   4394  O   SER A 274      -3.514  20.604 -24.213  1.00  0.00           O  
ATOM   4395  CB  SER A 274      -2.040  18.628 -22.009  1.00  0.00           C  
ATOM   4396  OG  SER A 274      -1.009  19.254 -22.722  1.00  0.00           O  
ATOM   4397  H   SER A 274      -2.998  18.374 -24.860  1.00  0.00           H  
ATOM   4398  HA  SER A 274      -4.141  18.279 -22.151  1.00  0.00           H  
ATOM   4399 1HB  SER A 274      -2.159  19.109 -21.038  1.00  0.00           H  
ATOM   4400 2HB  SER A 274      -1.779  17.586 -21.828  1.00  0.00           H  
ATOM   4401  HG  SER A 274      -0.908  20.125 -22.330  1.00  0.00           H  
ATOM   4402  N   LEU A 275      -4.206  20.851 -22.081  1.00  0.00           N  
ATOM   4403  CA  LEU A 275      -4.568  22.258 -22.220  1.00  0.00           C  
ATOM   4404  C   LEU A 275      -3.692  23.056 -21.267  1.00  0.00           C  
ATOM   4405  O   LEU A 275      -3.510  22.665 -20.114  1.00  0.00           O  
ATOM   4406  CB  LEU A 275      -6.036  22.484 -21.849  1.00  0.00           C  
ATOM   4407  CG  LEU A 275      -7.117  21.699 -22.593  1.00  0.00           C  
ATOM   4408  CD1 LEU A 275      -8.478  22.067 -22.021  1.00  0.00           C  
ATOM   4409  CD2 LEU A 275      -7.061  22.017 -24.086  1.00  0.00           C  
ATOM   4410  H   LEU A 275      -4.349  20.392 -21.193  1.00  0.00           H  
ATOM   4411  HA  LEU A 275      -4.427  22.550 -23.261  1.00  0.00           H  
ATOM   4412 1HB  LEU A 275      -6.167  22.250 -20.793  1.00  0.00           H  
ATOM   4413 2HB  LEU A 275      -6.274  23.537 -21.997  1.00  0.00           H  
ATOM   4414  HG  LEU A 275      -6.955  20.630 -22.448  1.00  0.00           H  
ATOM   4415 1HD1 LEU A 275      -9.256  21.511 -22.545  1.00  0.00           H  
ATOM   4416 2HD1 LEU A 275      -8.507  21.816 -20.960  1.00  0.00           H  
ATOM   4417 3HD1 LEU A 275      -8.649  23.135 -22.147  1.00  0.00           H  
ATOM   4418 1HD2 LEU A 275      -7.835  21.453 -24.608  1.00  0.00           H  
ATOM   4419 2HD2 LEU A 275      -7.227  23.084 -24.236  1.00  0.00           H  
ATOM   4420 3HD2 LEU A 275      -6.082  21.743 -24.480  1.00  0.00           H  
ATOM   4421  N   GLU A 276      -3.162  24.177 -21.743  1.00  0.00           N  
ATOM   4422  CA  GLU A 276      -2.307  25.019 -20.918  1.00  0.00           C  
ATOM   4423  C   GLU A 276      -2.607  26.500 -21.053  1.00  0.00           C  
ATOM   4424  O   GLU A 276      -3.039  26.968 -22.111  1.00  0.00           O  
ATOM   4425  CB  GLU A 276      -0.832  24.823 -21.285  1.00  0.00           C  
ATOM   4426  CG  GLU A 276      -0.272  23.439 -21.053  1.00  0.00           C  
ATOM   4427  CD  GLU A 276       1.187  23.338 -21.472  1.00  0.00           C  
ATOM   4428  OE1 GLU A 276       1.856  22.369 -21.057  1.00  0.00           O  
ATOM   4429  OE2 GLU A 276       1.661  24.225 -22.216  1.00  0.00           O1-
ATOM   4430  H   GLU A 276      -3.353  24.454 -22.695  1.00  0.00           H  
ATOM   4431  HA  GLU A 276      -2.444  24.733 -19.875  1.00  0.00           H  
ATOM   4432 1HB  GLU A 276      -0.685  25.055 -22.340  1.00  0.00           H  
ATOM   4433 2HB  GLU A 276      -0.218  25.517 -20.711  1.00  0.00           H  
ATOM   4434 1HG  GLU A 276      -0.360  23.195 -19.995  1.00  0.00           H  
ATOM   4435 2HG  GLU A 276      -0.866  22.719 -21.615  1.00  0.00           H  
ATOM   4436  N   ILE A 277      -2.380  27.223 -19.962  1.00  0.00           N  
ATOM   4437  CA  ILE A 277      -2.507  28.674 -19.938  1.00  0.00           C  
ATOM   4438  C   ILE A 277      -1.043  29.074 -19.770  1.00  0.00           C  
ATOM   4439  O   ILE A 277      -0.362  28.588 -18.867  1.00  0.00           O  
ATOM   4440  CB  ILE A 277      -3.391  29.202 -18.794  1.00  0.00           C  
ATOM   4441  CG1 ILE A 277      -4.816  28.658 -18.925  1.00  0.00           C  
ATOM   4442  CG2 ILE A 277      -3.398  30.723 -18.785  1.00  0.00           C  
ATOM   4443  CD1 ILE A 277      -5.690  28.937 -17.724  1.00  0.00           C  
ATOM   4444  H   ILE A 277      -2.108  26.739 -19.118  1.00  0.00           H  
ATOM   4445  HA  ILE A 277      -3.018  28.991 -20.846  1.00  0.00           H  
ATOM   4446  HB  ILE A 277      -3.004  28.844 -17.840  1.00  0.00           H  
ATOM   4447 1HG1 ILE A 277      -5.293  29.094 -19.802  1.00  0.00           H  
ATOM   4448 2HG1 ILE A 277      -4.781  27.578 -19.076  1.00  0.00           H  
ATOM   4449 1HG2 ILE A 277      -4.028  31.079 -17.970  1.00  0.00           H  
ATOM   4450 2HG2 ILE A 277      -2.382  31.090 -18.645  1.00  0.00           H  
ATOM   4451 3HG2 ILE A 277      -3.789  31.091 -19.734  1.00  0.00           H  
ATOM   4452 1HD1 ILE A 277      -6.683  28.520 -17.891  1.00  0.00           H  
ATOM   4453 2HD1 ILE A 277      -5.249  28.478 -16.838  1.00  0.00           H  
ATOM   4454 3HD1 ILE A 277      -5.769  30.013 -17.574  1.00  0.00           H  
ATOM   4455  N   ILE A 278      -0.556  29.934 -20.656  1.00  0.00           N  
ATOM   4456  CA  ILE A 278       0.842  30.354 -20.638  1.00  0.00           C  
ATOM   4457  C   ILE A 278       0.972  31.778 -20.104  1.00  0.00           C  
ATOM   4458  O   ILE A 278       0.684  32.738 -20.807  1.00  0.00           O  
ATOM   4459  CB  ILE A 278       1.427  30.258 -22.067  1.00  0.00           C  
ATOM   4460  CG1 ILE A 278       1.185  28.849 -22.625  1.00  0.00           C  
ATOM   4461  CG2 ILE A 278       2.915  30.569 -22.061  1.00  0.00           C  
ATOM   4462  CD1 ILE A 278       1.731  27.725 -21.753  1.00  0.00           C  
ATOM   4463  H   ILE A 278      -1.173  30.308 -21.363  1.00  0.00           H  
ATOM   4464  HA  ILE A 278       1.396  29.686 -19.980  1.00  0.00           H  
ATOM   4465  HB  ILE A 278       0.921  30.972 -22.715  1.00  0.00           H  
ATOM   4466 1HG1 ILE A 278       0.115  28.687 -22.751  1.00  0.00           H  
ATOM   4467 2HG1 ILE A 278       1.646  28.763 -23.609  1.00  0.00           H  
ATOM   4468 1HG2 ILE A 278       3.307  30.495 -23.075  1.00  0.00           H  
ATOM   4469 2HG2 ILE A 278       3.074  31.578 -21.684  1.00  0.00           H  
ATOM   4470 3HG2 ILE A 278       3.433  29.856 -21.420  1.00  0.00           H  
ATOM   4471 1HD1 ILE A 278       1.517  26.764 -22.221  1.00  0.00           H  
ATOM   4472 2HD1 ILE A 278       2.809  27.843 -21.641  1.00  0.00           H  
ATOM   4473 3HD1 ILE A 278       1.258  27.764 -20.772  1.00  0.00           H  
ATOM   4474  N   PRO A 279       1.431  31.929 -18.853  1.00  0.00           N  
ATOM   4475  CA  PRO A 279       1.590  33.241 -18.217  1.00  0.00           C  
ATOM   4476  C   PRO A 279       2.320  34.321 -19.001  1.00  0.00           C  
ATOM   4477  O   PRO A 279       1.945  35.493 -18.942  1.00  0.00           O  
ATOM   4478  CB  PRO A 279       2.381  32.874 -16.957  1.00  0.00           C  
ATOM   4479  CG  PRO A 279       1.933  31.490 -16.634  1.00  0.00           C  
ATOM   4480  CD  PRO A 279       1.816  30.812 -17.974  1.00  0.00           C  
ATOM   4481  HA  PRO A 279       0.597  33.673 -18.024  1.00  0.00           H  
ATOM   4482 1HB  PRO A 279       3.461  32.938 -17.159  1.00  0.00           H  
ATOM   4483 2HB  PRO A 279       2.162  33.592 -16.153  1.00  0.00           H  
ATOM   4484 1HG  PRO A 279       2.663  30.999 -15.974  1.00  0.00           H  
ATOM   4485 2HG  PRO A 279       0.978  31.517 -16.089  1.00  0.00           H  
ATOM   4486 1HD  PRO A 279       2.793  30.396 -18.261  1.00  0.00           H  
ATOM   4487 2HD  PRO A 279       1.057  30.018 -17.918  1.00  0.00           H  
ATOM   4488  N   CYS A 280       3.365  33.953 -19.730  1.00 10.74           N  
ATOM   4489  CA  CYS A 280       4.093  34.971 -20.472  1.00 11.97           C  
ATOM   4490  C   CYS A 280       3.420  35.442 -21.751  1.00 10.64           C  
ATOM   4491  O   CYS A 280       3.897  36.372 -22.401  1.00 11.23           O  
ATOM   4492  CB  CYS A 280       5.519  34.510 -20.744  1.00 12.94           C  
ATOM   4493  SG  CYS A 280       6.510  34.499 -19.212  1.00 17.49           S  
ATOM   4494  H   CYS A 280       3.671  32.991 -19.784  1.00  0.00           H  
ATOM   4495  HA  CYS A 280       4.129  35.879 -19.870  1.00  0.00           H  
ATOM   4496 1HB  CYS A 280       5.501  33.509 -21.176  1.00  0.00           H  
ATOM   4497 2HB  CYS A 280       5.982  35.172 -21.474  1.00  0.00           H  
ATOM   4498  N   SER A 281       2.305  34.807 -22.108  1.00  0.00           N  
ATOM   4499  CA  SER A 281       1.550  35.211 -23.290  1.00  0.00           C  
ATOM   4500  C   SER A 281       0.340  35.981 -22.777  1.00  0.00           C  
ATOM   4501  O   SER A 281      -0.456  35.445 -22.011  1.00  0.00           O  
ATOM   4502  CB  SER A 281       1.123  34.013 -24.115  1.00  0.00           C  
ATOM   4503  OG  SER A 281       0.303  34.404 -25.182  1.00  0.00           O  
ATOM   4504  H   SER A 281       1.973  34.031 -21.553  1.00  0.00           H  
ATOM   4505  HA  SER A 281       2.201  35.812 -23.928  1.00  0.00           H  
ATOM   4506 1HB  SER A 281       2.007  33.503 -24.500  1.00  0.00           H  
ATOM   4507 2HB  SER A 281       0.588  33.307 -23.482  1.00  0.00           H  
ATOM   4508  HG  SER A 281       0.894  34.732 -25.864  1.00  0.00           H  
ATOM   4509  N   ARG A 282       0.210  37.237 -23.191  1.00  0.00           N  
ATOM   4510  CA  ARG A 282      -0.899  38.075 -22.743  1.00  0.00           C  
ATOM   4511  C   ARG A 282      -1.658  38.668 -23.913  1.00  0.00           C  
ATOM   4512  O   ARG A 282      -1.057  39.206 -24.836  1.00  0.00           O  
ATOM   4513  CB  ARG A 282      -0.373  39.222 -21.881  1.00  0.00           C  
ATOM   4514  CG  ARG A 282       0.247  38.792 -20.573  1.00  0.00           C  
ATOM   4515  CD  ARG A 282      -0.810  38.560 -19.507  1.00  0.00           C  
ATOM   4516  NE  ARG A 282      -0.201  38.132 -18.255  1.00  0.00           N  
ATOM   4517  CZ  ARG A 282      -0.670  38.436 -17.051  1.00  0.00           C  
ATOM   4518  NH1 ARG A 282      -0.038  37.996 -15.971  1.00  0.00           N1+
ATOM   4519  NH2 ARG A 282      -1.764  39.182 -16.926  1.00  0.00           N  
ATOM   4520  H   ARG A 282       0.891  37.622 -23.829  1.00  0.00           H  
ATOM   4521  HA  ARG A 282      -1.574  37.464 -22.143  1.00  0.00           H  
ATOM   4522 1HB  ARG A 282       0.379  39.780 -22.437  1.00  0.00           H  
ATOM   4523 2HB  ARG A 282      -1.187  39.909 -21.652  1.00  0.00           H  
ATOM   4524 1HG  ARG A 282       0.799  37.864 -20.720  1.00  0.00           H  
ATOM   4525 2HG  ARG A 282       0.929  39.567 -20.220  1.00  0.00           H  
ATOM   4526 1HD  ARG A 282      -1.358  39.485 -19.331  1.00  0.00           H  
ATOM   4527 2HD  ARG A 282      -1.500  37.787 -19.842  1.00  0.00           H  
ATOM   4528  HE  ARG A 282       0.635  37.565 -18.308  1.00  0.00           H  
ATOM   4529 1HH1 ARG A 282       0.795  37.433 -16.066  1.00  0.00           H  
ATOM   4530 2HH1 ARG A 282      -0.391  38.225 -15.053  1.00  0.00           H  
ATOM   4531 1HH2 ARG A 282      -2.242  39.519 -17.750  1.00  0.00           H  
ATOM   4532 2HH2 ARG A 282      -2.116  39.410 -16.009  1.00  0.00           H  
ATOM   4533  N   VAL A 283      -2.981  38.550 -23.872  1.00  0.00           N  
ATOM   4534  CA  VAL A 283      -3.840  39.098 -24.911  1.00  0.00           C  
ATOM   4535  C   VAL A 283      -5.078  39.631 -24.211  1.00  0.00           C  
ATOM   4536  O   VAL A 283      -5.710  38.914 -23.434  1.00  0.00           O  
ATOM   4537  CB  VAL A 283      -4.290  38.023 -25.931  1.00  0.00           C  
ATOM   4538  CG1 VAL A 283      -5.160  38.663 -27.014  1.00  0.00           C  
ATOM   4539  CG2 VAL A 283      -3.076  37.351 -26.565  1.00  0.00           C  
ATOM   4540  H   VAL A 283      -3.400  38.063 -23.092  1.00  0.00           H  
ATOM   4541  HA  VAL A 283      -3.281  39.859 -25.457  1.00  0.00           H  
ATOM   4542  HB  VAL A 283      -4.888  37.272 -25.413  1.00  0.00           H  
ATOM   4543 1HG1 VAL A 283      -5.473  37.901 -27.727  1.00  0.00           H  
ATOM   4544 2HG1 VAL A 283      -6.040  39.113 -26.555  1.00  0.00           H  
ATOM   4545 3HG1 VAL A 283      -4.588  39.431 -27.534  1.00  0.00           H  
ATOM   4546 1HG2 VAL A 283      -3.409  36.597 -27.278  1.00  0.00           H  
ATOM   4547 2HG2 VAL A 283      -2.474  38.099 -27.081  1.00  0.00           H  
ATOM   4548 3HG2 VAL A 283      -2.476  36.875 -25.788  1.00  0.00           H  
ATOM   4549  N   GLY A 284      -5.409  40.889 -24.475  1.00  0.00           N  
ATOM   4550  CA  GLY A 284      -6.581  41.491 -23.863  1.00  0.00           C  
ATOM   4551  C   GLY A 284      -7.833  41.134 -24.638  1.00  0.00           C  
ATOM   4552  O   GLY A 284      -7.782  40.912 -25.849  1.00  0.00           O  
ATOM   4553  H   GLY A 284      -4.846  41.441 -25.106  1.00  0.00           H  
ATOM   4554 1HA  GLY A 284      -6.671  41.147 -22.833  1.00  0.00           H  
ATOM   4555 2HA  GLY A 284      -6.460  42.573 -23.831  1.00  0.00           H  
ATOM   4556  N   HIS A 285      -8.963  41.082 -23.939  1.00  0.00           N  
ATOM   4557  CA  HIS A 285     -10.236  40.730 -24.561  1.00  0.00           C  
ATOM   4558  C   HIS A 285     -11.365  41.559 -23.959  1.00  0.00           C  
ATOM   4559  O   HIS A 285     -11.406  41.772 -22.748  1.00  0.00           O  
ATOM   4560  CB  HIS A 285     -10.504  39.232 -24.346  1.00  0.00           C  
ATOM   4561  CG  HIS A 285     -11.888  38.790 -24.717  1.00  0.00           C  
ATOM   4562  CD2 HIS A 285     -12.426  38.443 -25.910  1.00  0.00           C  
ATOM   4563  ND1 HIS A 285     -12.899  38.656 -23.788  1.00  0.00           N  
ATOM   4564  CE1 HIS A 285     -13.999  38.245 -24.394  1.00  0.00           C  
ATOM   4565  NE2 HIS A 285     -13.740  38.108 -25.681  1.00  0.00           N  
ATOM   4566  H   HIS A 285      -8.939  41.291 -22.952  1.00  0.00           H  
ATOM   4567  HA  HIS A 285     -10.185  40.928 -25.631  1.00  0.00           H  
ATOM   4568 1HB  HIS A 285      -9.796  38.648 -24.935  1.00  0.00           H  
ATOM   4569 2HB  HIS A 285     -10.343  38.981 -23.298  1.00  0.00           H  
ATOM   4570  HD2 HIS A 285     -11.912  38.431 -26.872  1.00  0.00           H  
ATOM   4571  HE1 HIS A 285     -14.958  38.052 -23.913  1.00  0.00           H  
ATOM   4572  HE2 HIS A 285     -14.396  37.806 -26.387  1.00  0.00           H  
ATOM   4573  N   VAL A 286     -12.271  42.033 -24.811  1.00  0.00           N  
ATOM   4574  CA  VAL A 286     -13.403  42.809 -24.331  1.00  0.00           C  
ATOM   4575  C   VAL A 286     -14.488  41.860 -23.835  1.00  0.00           C  
ATOM   4576  O   VAL A 286     -15.296  41.355 -24.621  1.00  0.00           O  
ATOM   4577  CB  VAL A 286     -14.016  43.704 -25.439  1.00  0.00           C  
ATOM   4578  CG1 VAL A 286     -15.173  44.529 -24.854  1.00  0.00           C  
ATOM   4579  CG2 VAL A 286     -12.955  44.614 -26.032  1.00  0.00           C  
ATOM   4580  H   VAL A 286     -12.180  41.857 -25.801  1.00  0.00           H  
ATOM   4581  HA  VAL A 286     -13.058  43.463 -23.528  1.00  0.00           H  
ATOM   4582  HB  VAL A 286     -14.423  43.066 -26.224  1.00  0.00           H  
ATOM   4583 1HG1 VAL A 286     -15.602  45.157 -25.635  1.00  0.00           H  
ATOM   4584 2HG1 VAL A 286     -15.939  43.858 -24.466  1.00  0.00           H  
ATOM   4585 3HG1 VAL A 286     -14.798  45.160 -24.048  1.00  0.00           H  
ATOM   4586 1HG2 VAL A 286     -13.401  45.234 -26.809  1.00  0.00           H  
ATOM   4587 2HG2 VAL A 286     -12.544  45.252 -25.250  1.00  0.00           H  
ATOM   4588 3HG2 VAL A 286     -12.158  44.010 -26.464  1.00  0.00           H  
ATOM   4589  N   PHE A 287     -14.497  41.599 -22.532  1.00  0.00           N  
ATOM   4590  CA  PHE A 287     -15.516  40.734 -21.958  1.00  0.00           C  
ATOM   4591  C   PHE A 287     -16.812  41.528 -21.869  1.00  0.00           C  
ATOM   4592  O   PHE A 287     -16.801  42.711 -21.529  1.00  0.00           O  
ATOM   4593  CB  PHE A 287     -15.101  40.229 -20.575  1.00  0.00           C  
ATOM   4594  CG  PHE A 287     -13.994  39.214 -20.607  1.00  0.00           C  
ATOM   4595  CD1 PHE A 287     -12.673  39.603 -20.444  1.00  0.00           C  
ATOM   4596  CD2 PHE A 287     -14.270  37.870 -20.803  1.00  0.00           C  
ATOM   4597  CE1 PHE A 287     -11.653  38.671 -20.473  1.00  0.00           C  
ATOM   4598  CE2 PHE A 287     -13.253  36.935 -20.832  1.00  0.00           C  
ATOM   4599  CZ  PHE A 287     -11.943  37.337 -20.667  1.00  0.00           C  
ATOM   4600  H   PHE A 287     -13.794  42.001 -21.927  1.00  0.00           H  
ATOM   4601  HA  PHE A 287     -15.634  39.863 -22.604  1.00  0.00           H  
ATOM   4602 1HB  PHE A 287     -14.775  41.069 -19.964  1.00  0.00           H  
ATOM   4603 2HB  PHE A 287     -15.960  39.778 -20.080  1.00  0.00           H  
ATOM   4604  HD1 PHE A 287     -12.443  40.658 -20.289  1.00  0.00           H  
ATOM   4605  HD2 PHE A 287     -15.306  37.552 -20.933  1.00  0.00           H  
ATOM   4606  HE1 PHE A 287     -10.619  38.990 -20.344  1.00  0.00           H  
ATOM   4607  HE2 PHE A 287     -13.484  35.881 -20.985  1.00  0.00           H  
ATOM   4608  HZ  PHE A 287     -11.140  36.602 -20.692  1.00  0.00           H  
ATOM   4609  N   ARG A 288     -17.921  40.868 -22.191  1.00  0.00           N  
ATOM   4610  CA  ARG A 288     -19.241  41.486 -22.167  1.00  0.00           C  
ATOM   4611  C   ARG A 288     -20.151  40.718 -21.227  1.00  0.00           C  
ATOM   4612  O   ARG A 288     -19.795  39.649 -20.735  1.00  0.00           O  
ATOM   4613  CB  ARG A 288     -19.867  41.472 -23.566  1.00  0.00           C  
ATOM   4614  CG  ARG A 288     -19.195  42.382 -24.568  1.00  0.00           C  
ATOM   4615  CD  ARG A 288     -19.897  42.342 -25.922  1.00  0.00           C  
ATOM   4616  NE  ARG A 288     -19.568  41.155 -26.711  1.00  0.00           N  
ATOM   4617  CZ  ARG A 288     -19.527  41.149 -28.040  1.00  0.00           C  
ATOM   4618  NH1 ARG A 288     -19.791  42.262 -28.706  1.00  0.00           N1+
ATOM   4619  NH2 ARG A 288     -19.214  40.045 -28.705  1.00  0.00           N  
ATOM   4620  H   ARG A 288     -17.836  39.899 -22.464  1.00  0.00           H  
ATOM   4621  HA  ARG A 288     -19.132  42.522 -21.845  1.00  0.00           H  
ATOM   4622 1HB  ARG A 288     -19.838  40.460 -23.967  1.00  0.00           H  
ATOM   4623 2HB  ARG A 288     -20.913  41.768 -23.499  1.00  0.00           H  
ATOM   4624 1HG  ARG A 288     -19.217  43.407 -24.200  1.00  0.00           H  
ATOM   4625 2HG  ARG A 288     -18.160  42.068 -24.708  1.00  0.00           H  
ATOM   4626 1HD  ARG A 288     -20.976  42.347 -25.774  1.00  0.00           H  
ATOM   4627 2HD  ARG A 288     -19.607  43.214 -26.507  1.00  0.00           H  
ATOM   4628  HE  ARG A 288     -19.363  40.301 -26.211  1.00  0.00           H  
ATOM   4629 1HH1 ARG A 288     -20.022  43.109 -28.204  1.00  0.00           H  
ATOM   4630 2HH1 ARG A 288     -19.762  42.266 -29.715  1.00  0.00           H  
ATOM   4631 1HH2 ARG A 288     -19.004  39.194 -28.202  1.00  0.00           H  
ATOM   4632 2HH2 ARG A 288     -19.187  40.054 -29.714  1.00  0.00           H  
ATOM   4633  N   LYS A 289     -21.335  41.268 -20.987  1.00  0.00           N  
ATOM   4634  CA  LYS A 289     -22.308  40.625 -20.116  1.00  0.00           C  
ATOM   4635  C   LYS A 289     -23.460  40.043 -20.927  1.00  0.00           C  
ATOM   4636  O   LYS A 289     -24.245  39.243 -20.418  1.00  0.00           O  
ATOM   4637  CB  LYS A 289     -22.838  41.616 -19.079  1.00  0.00           C  
ATOM   4638  CG  LYS A 289     -21.787  42.129 -18.104  1.00  0.00           C  
ATOM   4639  CD  LYS A 289     -22.395  43.081 -17.085  1.00  0.00           C  
ATOM   4640  CE  LYS A 289     -21.348  43.586 -16.103  1.00  0.00           C  
ATOM   4641  NZ  LYS A 289     -21.923  44.543 -15.120  1.00  0.00           N1+
ATOM   4642  H   LYS A 289     -21.568  42.151 -21.418  1.00  0.00           H  
ATOM   4643  HA  LYS A 289     -21.811  39.816 -19.579  1.00  0.00           H  
ATOM   4644 1HB  LYS A 289     -23.272  42.478 -19.587  1.00  0.00           H  
ATOM   4645 2HB  LYS A 289     -23.632  41.146 -18.498  1.00  0.00           H  
ATOM   4646 1HG  LYS A 289     -21.336  41.287 -17.578  1.00  0.00           H  
ATOM   4647 2HG  LYS A 289     -21.005  42.652 -18.654  1.00  0.00           H  
ATOM   4648 1HD  LYS A 289     -22.838  43.934 -17.602  1.00  0.00           H  
ATOM   4649 2HD  LYS A 289     -23.181  42.567 -16.531  1.00  0.00           H  
ATOM   4650 1HE  LYS A 289     -20.920  42.743 -15.563  1.00  0.00           H  
ATOM   4651 2HE  LYS A 289     -20.547  44.084 -16.650  1.00  0.00           H  
ATOM   4652 1HZ  LYS A 289     -21.198  44.854 -14.488  1.00  0.00           H  
ATOM   4653 2HZ  LYS A 289     -22.307  45.340 -15.609  1.00  0.00           H  
ATOM   4654 3HZ  LYS A 289     -22.654  44.087 -14.593  1.00  0.00           H  
ATOM   4655  N   GLN A 290     -23.557  40.442 -22.189  1.00  0.00           N  
ATOM   4656  CA  GLN A 290     -24.621  39.953 -23.055  1.00  0.00           C  
ATOM   4657  C   GLN A 290     -24.175  39.947 -24.514  1.00  0.00           C  
ATOM   4658  O   GLN A 290     -23.411  40.812 -24.935  1.00  0.00           O  
ATOM   4659  CB  GLN A 290     -25.881  40.808 -22.893  1.00  0.00           C  
ATOM   4660  CG  GLN A 290     -27.081  40.303 -23.676  1.00  0.00           C  
ATOM   4661  CD  GLN A 290     -28.330  41.122 -23.412  1.00  0.00           C  
ATOM   4662  NE2 GLN A 290     -29.476  40.606 -23.840  1.00  0.00           N  
ATOM   4663  OE1 GLN A 290     -28.266  42.207 -22.828  1.00  0.00           O  
ATOM   4664  H   GLN A 290     -22.883  41.097 -22.559  1.00  0.00           H  
ATOM   4665  HA  GLN A 290     -24.868  38.933 -22.759  1.00  0.00           H  
ATOM   4666 1HB  GLN A 290     -26.159  40.851 -21.840  1.00  0.00           H  
ATOM   4667 2HB  GLN A 290     -25.672  41.828 -23.218  1.00  0.00           H  
ATOM   4668 1HG  GLN A 290     -26.856  40.357 -24.741  1.00  0.00           H  
ATOM   4669 2HG  GLN A 290     -27.281  39.271 -23.388  1.00  0.00           H  
ATOM   4670 1HE2 GLN A 290     -30.334  41.100 -23.694  1.00  0.00           H  
ATOM   4671 2HE2 GLN A 290     -29.483  39.722 -24.309  1.00  0.00           H  
ATOM   4672  N   HIS A 291     -24.636  38.959 -25.278  1.00  0.00           N  
ATOM   4673  CA  HIS A 291     -24.291  38.879 -26.696  1.00  0.00           C  
ATOM   4674  C   HIS A 291     -25.184  39.863 -27.447  1.00  0.00           C  
ATOM   4675  O   HIS A 291     -26.388  39.935 -27.198  1.00  0.00           O  
ATOM   4676  CB  HIS A 291     -24.523  37.462 -27.226  1.00  0.00           C  
ATOM   4677  CG  HIS A 291     -23.673  36.421 -26.564  1.00  0.00           C  
ATOM   4678  CD2 HIS A 291     -24.008  35.316 -25.857  1.00  0.00           C  
ATOM   4679  ND1 HIS A 291     -22.297  36.451 -26.599  1.00  0.00           N  
ATOM   4680  CE1 HIS A 291     -21.819  35.408 -25.941  1.00  0.00           C  
ATOM   4681  NE2 HIS A 291     -22.837  34.705 -25.482  1.00  0.00           N  
ATOM   4682  H   HIS A 291     -25.233  38.252 -24.877  1.00  0.00           H  
ATOM   4683  HA  HIS A 291     -23.237  39.122 -26.829  1.00  0.00           H  
ATOM   4684 1HB  HIS A 291     -25.569  37.187 -27.085  1.00  0.00           H  
ATOM   4685 2HB  HIS A 291     -24.320  37.436 -28.297  1.00  0.00           H  
ATOM   4686  HD2 HIS A 291     -25.019  34.975 -25.628  1.00  0.00           H  
ATOM   4687  HE1 HIS A 291     -20.764  35.172 -25.803  1.00  0.00           H  
ATOM   4688  HE2 HIS A 291     -22.771  33.854 -24.941  1.00  0.00           H  
ATOM   4689  N   PRO A 292     -24.611  40.634 -28.382  1.00  0.00           N  
ATOM   4690  CA  PRO A 292     -25.424  41.598 -29.126  1.00  0.00           C  
ATOM   4691  C   PRO A 292     -26.161  41.010 -30.323  1.00  0.00           C  
ATOM   4692  O   PRO A 292     -27.100  41.614 -30.842  1.00  0.00           O  
ATOM   4693  CB  PRO A 292     -24.379  42.624 -29.576  1.00  0.00           C  
ATOM   4694  CG  PRO A 292     -23.134  41.825 -29.752  1.00  0.00           C  
ATOM   4695  CD  PRO A 292     -23.188  40.800 -28.650  1.00  0.00           C  
ATOM   4696  HA  PRO A 292     -26.179  42.030 -28.452  1.00  0.00           H  
ATOM   4697 1HB  PRO A 292     -24.708  43.112 -30.506  1.00  0.00           H  
ATOM   4698 2HB  PRO A 292     -24.276  43.414 -28.818  1.00  0.00           H  
ATOM   4699 1HG  PRO A 292     -23.112  41.370 -30.753  1.00  0.00           H  
ATOM   4700 2HG  PRO A 292     -22.251  42.477 -29.680  1.00  0.00           H  
ATOM   4701 1HD  PRO A 292     -22.741  39.858 -29.002  1.00  0.00           H  
ATOM   4702 2HD  PRO A 292     -22.650  41.179 -27.769  1.00  0.00           H  
ATOM   4703  N   TYR A 293     -25.745  39.822 -30.740  1.00  0.00           N  
ATOM   4704  CA  TYR A 293     -26.326  39.175 -31.907  1.00  0.00           C  
ATOM   4705  C   TYR A 293     -27.422  38.155 -31.639  1.00  0.00           C  
ATOM   4706  O   TYR A 293     -27.630  37.719 -30.511  1.00  0.00           O  
ATOM   4707  CB  TYR A 293     -25.209  38.537 -32.729  1.00  0.00           C  
ATOM   4708  CG  TYR A 293     -24.245  37.733 -31.897  1.00  0.00           C  
ATOM   4709  CD1 TYR A 293     -24.612  36.494 -31.372  1.00  0.00           C  
ATOM   4710  CD2 TYR A 293     -22.963  38.212 -31.623  1.00  0.00           C  
ATOM   4711  CE1 TYR A 293     -23.729  35.750 -30.600  1.00  0.00           C  
ATOM   4712  CE2 TYR A 293     -22.071  37.475 -30.848  1.00  0.00           C  
ATOM   4713  CZ  TYR A 293     -22.461  36.242 -30.344  1.00  0.00           C  
ATOM   4714  OH  TYR A 293     -21.574  35.481 -29.616  1.00  0.00           O  
ATOM   4715  H   TYR A 293     -25.005  39.354 -30.236  1.00  0.00           H  
ATOM   4716  HA  TYR A 293     -26.830  39.932 -32.508  1.00  0.00           H  
ATOM   4717 1HB  TYR A 293     -25.642  37.882 -33.487  1.00  0.00           H  
ATOM   4718 2HB  TYR A 293     -24.651  39.315 -33.250  1.00  0.00           H  
ATOM   4719  HD1 TYR A 293     -25.607  36.092 -31.566  1.00  0.00           H  
ATOM   4720  HD2 TYR A 293     -22.647  39.178 -32.018  1.00  0.00           H  
ATOM   4721  HE1 TYR A 293     -24.035  34.785 -30.199  1.00  0.00           H  
ATOM   4722  HE2 TYR A 293     -21.076  37.867 -30.640  1.00  0.00           H  
ATOM   4723  HH  TYR A 293     -20.735  35.944 -29.550  1.00  0.00           H  
ATOM   4724  N   THR A 294     -28.108  37.769 -32.709  1.00  0.00           N  
ATOM   4725  CA  THR A 294     -29.216  36.826 -32.634  1.00  0.00           C  
ATOM   4726  C   THR A 294     -28.857  35.347 -32.540  1.00  0.00           C  
ATOM   4727  O   THR A 294     -27.837  34.896 -33.065  1.00  0.00           O  
ATOM   4728  CB  THR A 294     -30.137  37.021 -33.852  1.00  0.00           C  
ATOM   4729  CG2 THR A 294     -30.688  38.439 -33.883  1.00  0.00           C  
ATOM   4730  OG1 THR A 294     -29.401  36.773 -35.057  1.00  0.00           O  
ATOM   4731  H   THR A 294     -27.845  38.149 -33.607  1.00  0.00           H  
ATOM   4732  HA  THR A 294     -29.788  37.035 -31.729  1.00  0.00           H  
ATOM   4733  HB  THR A 294     -30.967  36.317 -33.798  1.00  0.00           H  
ATOM   4734  HG1 THR A 294     -29.852  37.183 -35.798  1.00  0.00           H  
ATOM   4735 1HG2 THR A 294     -31.337  38.558 -34.750  1.00  0.00           H  
ATOM   4736 2HG2 THR A 294     -31.258  38.627 -32.974  1.00  0.00           H  
ATOM   4737 3HG2 THR A 294     -29.864  39.148 -33.947  1.00  0.00           H  
ATOM   4738  N   PHE A 295     -29.726  34.605 -31.861  1.00  0.00           N  
ATOM   4739  CA  PHE A 295     -29.608  33.161 -31.685  1.00  0.00           C  
ATOM   4740  C   PHE A 295     -30.937  32.580 -32.169  1.00  0.00           C  
ATOM   4741  O   PHE A 295     -31.973  32.817 -31.552  1.00  0.00           O  
ATOM   4742  CB  PHE A 295     -29.458  32.795 -30.205  1.00  0.00           C  
ATOM   4743  CG  PHE A 295     -28.069  32.958 -29.656  1.00  0.00           C  
ATOM   4744  CD1 PHE A 295     -27.255  31.845 -29.453  1.00  0.00           C  
ATOM   4745  CD2 PHE A 295     -27.596  34.210 -29.279  1.00  0.00           C  
ATOM   4746  CE1 PHE A 295     -25.991  31.976 -28.876  1.00  0.00           C  
ATOM   4747  CE2 PHE A 295     -26.335  34.353 -28.702  1.00  0.00           C  
ATOM   4748  CZ  PHE A 295     -25.532  33.233 -28.498  1.00  0.00           C  
ATOM   4749  H   PHE A 295     -30.512  35.088 -31.448  1.00  0.00           H  
ATOM   4750  HA  PHE A 295     -28.717  32.819 -32.214  1.00  0.00           H  
ATOM   4751 1HB  PHE A 295     -30.125  33.414 -29.608  1.00  0.00           H  
ATOM   4752 2HB  PHE A 295     -29.754  31.757 -30.056  1.00  0.00           H  
ATOM   4753  HD1 PHE A 295     -27.615  30.861 -29.753  1.00  0.00           H  
ATOM   4754  HD2 PHE A 295     -28.225  35.087 -29.438  1.00  0.00           H  
ATOM   4755  HE1 PHE A 295     -25.369  31.095 -28.722  1.00  0.00           H  
ATOM   4756  HE2 PHE A 295     -25.977  35.339 -28.410  1.00  0.00           H  
ATOM   4757  HZ  PHE A 295     -24.549  33.341 -28.044  1.00  0.00           H  
ATOM   4758  N   PRO A 296     -30.936  31.833 -33.285  1.00  0.00           N  
ATOM   4759  CA  PRO A 296     -32.216  31.277 -33.735  1.00  0.00           C  
ATOM   4760  C   PRO A 296     -32.775  30.344 -32.665  1.00  0.00           C  
ATOM   4761  O   PRO A 296     -32.148  29.346 -32.314  1.00  0.00           O  
ATOM   4762  CB  PRO A 296     -31.847  30.521 -35.015  1.00  0.00           C  
ATOM   4763  CG  PRO A 296     -30.597  31.184 -35.484  1.00  0.00           C  
ATOM   4764  CD  PRO A 296     -29.857  31.526 -34.219  1.00  0.00           C  
ATOM   4765  HA  PRO A 296     -32.912  32.100 -33.950  1.00  0.00           H  
ATOM   4766 1HB  PRO A 296     -31.706  29.453 -34.794  1.00  0.00           H  
ATOM   4767 2HB  PRO A 296     -32.667  30.592 -35.744  1.00  0.00           H  
ATOM   4768 1HG  PRO A 296     -30.029  30.505 -36.137  1.00  0.00           H  
ATOM   4769 2HG  PRO A 296     -30.841  32.073 -36.085  1.00  0.00           H  
ATOM   4770 1HD  PRO A 296     -29.264  30.658 -33.895  1.00  0.00           H  
ATOM   4771 2HD  PRO A 296     -29.207  32.396 -34.397  1.00  0.00           H  
ATOM   4772  N   GLY A 297     -33.951  30.674 -32.141  1.00  0.00           N  
ATOM   4773  CA  GLY A 297     -34.553  29.852 -31.108  1.00  0.00           C  
ATOM   4774  C   GLY A 297     -34.216  30.341 -29.710  1.00  0.00           C  
ATOM   4775  O   GLY A 297     -34.734  29.820 -28.721  1.00  0.00           O  
ATOM   4776  H   GLY A 297     -34.437  31.502 -32.456  1.00  0.00           H  
ATOM   4777 1HA  GLY A 297     -35.636  29.845 -31.233  1.00  0.00           H  
ATOM   4778 2HA  GLY A 297     -34.211  28.823 -31.217  1.00  0.00           H  
ATOM   4779  N   GLY A 298     -33.347  31.343 -29.626  1.00  0.00           N  
ATOM   4780  CA  GLY A 298     -32.964  31.886 -28.334  1.00  0.00           C  
ATOM   4781  C   GLY A 298     -31.641  31.343 -27.828  1.00  0.00           C  
ATOM   4782  O   GLY A 298     -31.330  30.171 -28.029  1.00  0.00           O  
ATOM   4783  H   GLY A 298     -32.945  31.737 -30.464  1.00  0.00           H  
ATOM   4784 1HA  GLY A 298     -32.893  32.972 -28.402  1.00  0.00           H  
ATOM   4785 2HA  GLY A 298     -33.737  31.660 -27.601  1.00  0.00           H  
ATOM   4786  N   SER A 299     -30.869  32.196 -27.162  1.00  0.00           N  
ATOM   4787  CA  SER A 299     -29.569  31.808 -26.629  1.00  0.00           C  
ATOM   4788  C   SER A 299     -29.666  30.654 -25.637  1.00  0.00           C  
ATOM   4789  O   SER A 299     -28.796  29.784 -25.598  1.00  0.00           O  
ATOM   4790  CB  SER A 299     -28.914  33.000 -25.958  1.00  0.00           C  
ATOM   4791  OG  SER A 299     -29.643  33.406 -24.832  1.00  0.00           O  
ATOM   4792  H   SER A 299     -31.196  33.141 -27.022  1.00  0.00           H  
ATOM   4793  HA  SER A 299     -28.942  31.473 -27.456  1.00  0.00           H  
ATOM   4794 1HB  SER A 299     -27.898  32.738 -25.663  1.00  0.00           H  
ATOM   4795 2HB  SER A 299     -28.846  33.824 -26.667  1.00  0.00           H  
ATOM   4796  HG  SER A 299     -29.184  33.037 -24.074  1.00  0.00           H  
ATOM   4797  N   GLY A 300     -30.726  30.651 -24.833  1.00  0.00           N  
ATOM   4798  CA  GLY A 300     -30.900  29.589 -23.859  1.00  0.00           C  
ATOM   4799  C   GLY A 300     -31.017  28.218 -24.502  1.00  0.00           C  
ATOM   4800  O   GLY A 300     -30.391  27.253 -24.057  1.00  0.00           O  
ATOM   4801  H   GLY A 300     -31.418  31.384 -24.891  1.00  0.00           H  
ATOM   4802 1HA  GLY A 300     -30.055  29.586 -23.170  1.00  0.00           H  
ATOM   4803 2HA  GLY A 300     -31.794  29.781 -23.268  1.00  0.00           H  
ATOM   4804  N   THR A 301     -31.821  28.127 -25.555  1.00  0.00           N  
ATOM   4805  CA  THR A 301     -32.013  26.861 -26.250  1.00  0.00           C  
ATOM   4806  C   THR A 301     -30.745  26.438 -26.981  1.00  0.00           C  
ATOM   4807  O   THR A 301     -30.316  25.291 -26.873  1.00  0.00           O  
ATOM   4808  CB  THR A 301     -33.182  26.948 -27.248  1.00  0.00           C  
ATOM   4809  CG2 THR A 301     -33.346  25.633 -27.994  1.00  0.00           C  
ATOM   4810  OG1 THR A 301     -34.394  27.249 -26.544  1.00  0.00           O  
ATOM   4811  H   THR A 301     -32.311  28.947 -25.884  1.00  0.00           H  
ATOM   4812  HA  THR A 301     -32.261  26.096 -25.514  1.00  0.00           H  
ATOM   4813  HB  THR A 301     -32.988  27.745 -27.967  1.00  0.00           H  
ATOM   4814  HG1 THR A 301     -34.756  28.076 -26.871  1.00  0.00           H  
ATOM   4815 1HG2 THR A 301     -34.177  25.713 -28.695  1.00  0.00           H  
ATOM   4816 2HG2 THR A 301     -32.430  25.407 -28.540  1.00  0.00           H  
ATOM   4817 3HG2 THR A 301     -33.550  24.834 -27.282  1.00  0.00           H  
ATOM   4818  N   VAL A 302     -30.151  27.364 -27.726  1.00  0.00           N  
ATOM   4819  CA  VAL A 302     -28.933  27.062 -28.469  1.00  0.00           C  
ATOM   4820  C   VAL A 302     -27.816  26.599 -27.541  1.00  0.00           C  
ATOM   4821  O   VAL A 302     -27.157  25.597 -27.813  1.00  0.00           O  
ATOM   4822  CB  VAL A 302     -28.464  28.304 -29.251  1.00  0.00           C  
ATOM   4823  CG1 VAL A 302     -27.087  28.066 -29.852  1.00  0.00           C  
ATOM   4824  CG2 VAL A 302     -29.474  28.644 -30.335  1.00  0.00           C  
ATOM   4825  H   VAL A 302     -30.544  28.293 -27.783  1.00  0.00           H  
ATOM   4826  HA  VAL A 302     -29.147  26.257 -29.173  1.00  0.00           H  
ATOM   4827  HB  VAL A 302     -28.373  29.143 -28.561  1.00  0.00           H  
ATOM   4828 1HG1 VAL A 302     -26.771  28.953 -30.400  1.00  0.00           H  
ATOM   4829 2HG1 VAL A 302     -26.373  27.859 -29.055  1.00  0.00           H  
ATOM   4830 3HG1 VAL A 302     -27.129  27.215 -30.532  1.00  0.00           H  
ATOM   4831 1HG2 VAL A 302     -29.137  29.524 -30.884  1.00  0.00           H  
ATOM   4832 2HG2 VAL A 302     -29.570  27.803 -31.022  1.00  0.00           H  
ATOM   4833 3HG2 VAL A 302     -30.443  28.851 -29.879  1.00  0.00           H  
ATOM   4834  N   PHE A 303     -27.600  27.323 -26.445  1.00  0.00           N  
ATOM   4835  CA  PHE A 303     -26.551  26.938 -25.505  1.00  0.00           C  
ATOM   4836  C   PHE A 303     -26.815  25.537 -24.959  1.00  0.00           C  
ATOM   4837  O   PHE A 303     -25.902  24.715 -24.848  1.00  0.00           O  
ATOM   4838  CB  PHE A 303     -26.467  27.916 -24.330  1.00  0.00           C  
ATOM   4839  CG  PHE A 303     -25.347  27.608 -23.375  1.00  0.00           C  
ATOM   4840  CD1 PHE A 303     -24.040  27.973 -23.678  1.00  0.00           C  
ATOM   4841  CD2 PHE A 303     -25.592  26.909 -22.194  1.00  0.00           C  
ATOM   4842  CE1 PHE A 303     -22.989  27.646 -22.822  1.00  0.00           C  
ATOM   4843  CE2 PHE A 303     -24.545  26.577 -21.330  1.00  0.00           C  
ATOM   4844  CZ  PHE A 303     -23.243  26.946 -21.648  1.00  0.00           C  
ATOM   4845  H   PHE A 303     -28.157  28.144 -26.252  1.00  0.00           H  
ATOM   4846  HA  PHE A 303     -25.594  26.952 -26.029  1.00  0.00           H  
ATOM   4847 1HB  PHE A 303     -26.328  28.928 -24.709  1.00  0.00           H  
ATOM   4848 2HB  PHE A 303     -27.405  27.900 -23.776  1.00  0.00           H  
ATOM   4849  HD1 PHE A 303     -23.840  28.522 -24.599  1.00  0.00           H  
ATOM   4850  HD2 PHE A 303     -26.615  26.620 -21.947  1.00  0.00           H  
ATOM   4851  HE1 PHE A 303     -21.970  27.938 -23.075  1.00  0.00           H  
ATOM   4852  HE2 PHE A 303     -24.747  26.029 -20.409  1.00  0.00           H  
ATOM   4853  HZ  PHE A 303     -22.423  26.688 -20.979  1.00  0.00           H  
ATOM   4854  N   ALA A 304     -28.070  25.268 -24.616  1.00  0.00           N  
ATOM   4855  CA  ALA A 304     -28.434  23.963 -24.088  1.00  0.00           C  
ATOM   4856  C   ALA A 304     -28.211  22.877 -25.136  1.00  0.00           C  
ATOM   4857  O   ALA A 304     -27.708  21.802 -24.828  1.00  0.00           O  
ATOM   4858  CB  ALA A 304     -29.894  23.972 -23.631  1.00  0.00           C  
ATOM   4859  H   ALA A 304     -28.786  25.974 -24.720  1.00  0.00           H  
ATOM   4860  HA  ALA A 304     -27.793  23.754 -23.231  1.00  0.00           H  
ATOM   4861 1HB  ALA A 304     -30.158  22.990 -23.237  1.00  0.00           H  
ATOM   4862 2HB  ALA A 304     -30.027  24.723 -22.853  1.00  0.00           H  
ATOM   4863 3HB  ALA A 304     -30.538  24.208 -24.477  1.00  0.00           H  
ATOM   4864  N   ARG A 305     -28.587  23.150 -26.382  1.00  0.00           N  
ATOM   4865  CA  ARG A 305     -28.401  22.155 -27.425  1.00  0.00           C  
ATOM   4866  C   ARG A 305     -26.923  21.841 -27.626  1.00  0.00           C  
ATOM   4867  O   ARG A 305     -26.531  20.676 -27.672  1.00  0.00           O  
ATOM   4868  CB  ARG A 305     -29.017  22.618 -28.750  1.00  0.00           C  
ATOM   4869  CG  ARG A 305     -28.854  21.585 -29.860  1.00  0.00           C  
ATOM   4870  CD  ARG A 305     -29.694  21.904 -31.089  1.00  0.00           C  
ATOM   4871  NE  ARG A 305     -29.284  23.135 -31.760  1.00  0.00           N  
ATOM   4872  CZ  ARG A 305     -30.001  24.255 -31.781  1.00  0.00           C  
ATOM   4873  NH1 ARG A 305     -29.544  25.324 -32.420  1.00  0.00           N1+
ATOM   4874  NH2 ARG A 305     -31.175  24.312 -31.164  1.00  0.00           N  
ATOM   4875  H   ARG A 305     -29.001  24.040 -26.619  1.00  0.00           H  
ATOM   4876  HA  ARG A 305     -28.900  21.235 -27.119  1.00  0.00           H  
ATOM   4877 1HB  ARG A 305     -30.078  22.818 -28.607  1.00  0.00           H  
ATOM   4878 2HB  ARG A 305     -28.547  23.551 -29.063  1.00  0.00           H  
ATOM   4879 1HG  ARG A 305     -27.809  21.547 -30.172  1.00  0.00           H  
ATOM   4880 2HG  ARG A 305     -29.159  20.605 -29.493  1.00  0.00           H  
ATOM   4881 1HD  ARG A 305     -29.609  21.092 -31.809  1.00  0.00           H  
ATOM   4882 2HD  ARG A 305     -30.737  22.021 -30.796  1.00  0.00           H  
ATOM   4883  HE  ARG A 305     -28.394  23.135 -32.241  1.00  0.00           H  
ATOM   4884 1HH1 ARG A 305     -28.651  25.287 -32.891  1.00  0.00           H  
ATOM   4885 2HH1 ARG A 305     -30.088  26.175 -32.436  1.00  0.00           H  
ATOM   4886 1HH2 ARG A 305     -31.529  23.501 -30.675  1.00  0.00           H  
ATOM   4887 2HH2 ARG A 305     -31.715  25.164 -31.184  1.00  0.00           H  
ATOM   4888  N   ASN A 306     -26.096  22.875 -27.744  1.00  0.00           N  
ATOM   4889  CA  ASN A 306     -24.669  22.647 -27.940  1.00  0.00           C  
ATOM   4890  C   ASN A 306     -24.073  21.903 -26.756  1.00  0.00           C  
ATOM   4891  O   ASN A 306     -23.298  20.968 -26.926  1.00  0.00           O  
ATOM   4892  CB  ASN A 306     -23.921  23.972 -28.119  1.00  0.00           C  
ATOM   4893  CG  ASN A 306     -24.309  24.695 -29.390  1.00  0.00           C  
ATOM   4894  ND2 ASN A 306     -24.004  25.986 -29.447  1.00  0.00           N  
ATOM   4895  OD1 ASN A 306     -24.857  24.099 -30.319  1.00  0.00           O  
ATOM   4896  H   ASN A 306     -26.442  23.823 -27.701  1.00  0.00           H  
ATOM   4897  HA  ASN A 306     -24.534  22.051 -28.844  1.00  0.00           H  
ATOM   4898 1HB  ASN A 306     -24.127  24.623 -27.268  1.00  0.00           H  
ATOM   4899 2HB  ASN A 306     -22.847  23.784 -28.136  1.00  0.00           H  
ATOM   4900 1HD2 ASN A 306     -24.235  26.518 -30.262  1.00  0.00           H  
ATOM   4901 2HD2 ASN A 306     -23.544  26.425 -28.676  1.00  0.00           H  
ATOM   4902  N   THR A 307     -24.439  22.331 -25.554  1.00  0.00           N  
ATOM   4903  CA  THR A 307     -23.922  21.707 -24.347  1.00  0.00           C  
ATOM   4904  C   THR A 307     -24.356  20.249 -24.233  1.00  0.00           C  
ATOM   4905  O   THR A 307     -23.562  19.404 -23.826  1.00  0.00           O  
ATOM   4906  CB  THR A 307     -24.375  22.480 -23.094  1.00  0.00           C  
ATOM   4907  CG2 THR A 307     -23.862  21.800 -21.834  1.00  0.00           C  
ATOM   4908  OG1 THR A 307     -23.870  23.820 -23.148  1.00  0.00           O  
ATOM   4909  H   THR A 307     -25.085  23.102 -25.471  1.00  0.00           H  
ATOM   4910  HA  THR A 307     -22.833  21.695 -24.403  1.00  0.00           H  
ATOM   4911  HB  THR A 307     -25.463  22.516 -23.063  1.00  0.00           H  
ATOM   4912  HG1 THR A 307     -24.427  24.348 -23.725  1.00  0.00           H  
ATOM   4913 1HG2 THR A 307     -24.191  22.359 -20.959  1.00  0.00           H  
ATOM   4914 2HG2 THR A 307     -24.253  20.783 -21.784  1.00  0.00           H  
ATOM   4915 3HG2 THR A 307     -22.773  21.768 -21.855  1.00  0.00           H  
ATOM   4916  N   ARG A 308     -25.601  19.947 -24.603  1.00  0.00           N  
ATOM   4917  CA  ARG A 308     -26.075  18.563 -24.556  1.00  0.00           C  
ATOM   4918  C   ARG A 308     -25.241  17.702 -25.497  1.00  0.00           C  
ATOM   4919  O   ARG A 308     -24.912  16.559 -25.180  1.00  0.00           O  
ATOM   4920  CB  ARG A 308     -27.546  18.451 -24.975  1.00  0.00           C  
ATOM   4921  CG  ARG A 308     -28.552  18.947 -23.957  1.00  0.00           C  
ATOM   4922  CD  ARG A 308     -29.918  18.303 -24.200  1.00  0.00           C  
ATOM   4923  NE  ARG A 308     -29.934  16.896 -23.801  1.00  0.00           N  
ATOM   4924  CZ  ARG A 308     -30.954  16.066 -24.006  1.00  0.00           C  
ATOM   4925  NH1 ARG A 308     -30.875  14.805 -23.602  1.00  0.00           N1+
ATOM   4926  NH2 ARG A 308     -32.048  16.491 -24.624  1.00  0.00           N  
ATOM   4927  H   ARG A 308     -26.229  20.671 -24.919  1.00  0.00           H  
ATOM   4928  HA  ARG A 308     -25.988  18.203 -23.530  1.00  0.00           H  
ATOM   4929 1HB  ARG A 308     -27.706  19.016 -25.892  1.00  0.00           H  
ATOM   4930 2HB  ARG A 308     -27.785  17.409 -25.188  1.00  0.00           H  
ATOM   4931 1HG  ARG A 308     -28.211  18.689 -22.954  1.00  0.00           H  
ATOM   4932 2HG  ARG A 308     -28.650  20.031 -24.040  1.00  0.00           H  
ATOM   4933 1HD  ARG A 308     -30.678  18.830 -23.623  1.00  0.00           H  
ATOM   4934 2HD  ARG A 308     -30.164  18.360 -25.260  1.00  0.00           H  
ATOM   4935  HE  ARG A 308     -29.113  16.530 -23.337  1.00  0.00           H  
ATOM   4936 1HH1 ARG A 308     -30.041  14.474 -23.139  1.00  0.00           H  
ATOM   4937 2HH1 ARG A 308     -31.649  14.176 -23.758  1.00  0.00           H  
ATOM   4938 1HH2 ARG A 308     -32.110  17.448 -24.941  1.00  0.00           H
```
